# Prognostic relevance of correlated co-expression of coding and noncoding RNAs in cervical cancers

**DOI:** 10.1101/2023.12.28.573593

**Authors:** Abhisikta Ghosh, Abarna Sinha, Arnab Ghosh, Somrita Roy, Sumana Mallick, Vinoth Kumar, Sonia Mathai, Jaydip Bhaumik, Asima Mukhopadhyay, Saugata Sen, Aditi Chandra, Arindam Maitra, Nidhan K. Biswas, Partha P. Majumder, Sharmila Sengupta

## Abstract

Human papillomavirus (HPV) drives cervical cancer (CaCx) pathogenesis and viral oncoproteins jeopardize global gene expression in such cancers. We aimed to identify differentially expressed coding (DEcGs) and long noncoding (sense intronic and Natural Antisense Transcripts) RNA genes (DElncGs) in HPV16-positive CaCx patients (N=44) compared to HPV-negative normal individuals (N=34). Thereby, employing strand-specific RNA-seq, we determined the relationships between DEcGs and DElncGs and their clinical implications. Gene set enrichment and protein-protein interaction analyses of DEcGs revealed enrichment of processes crucial for abortive virus life cycle and cancer progression. The DEcGs formed 16 gene clusters, portraying cancer-related functions. We recorded significantly correlated co-expression of 79 DElncGs with DEcGs at proximal genomic loci, and 24 such pairs portrayed significantly altered correlation coefficients among patients, compared to normal individuals. Of these, 6 DEcGs belonged to 5 gene clusters, one of which was survival-associated. Out of the 24 correlated DEcG: DElncG pairs, 3 pairs were identified, where expression of both members was significantly associated with patient overall survival. Besides being prognostically relevant, disruption of the significant correlative relationships of such gene pairs in CaCx bears immense potential for patient-targeted therapy.

**Graphical Abstract:**
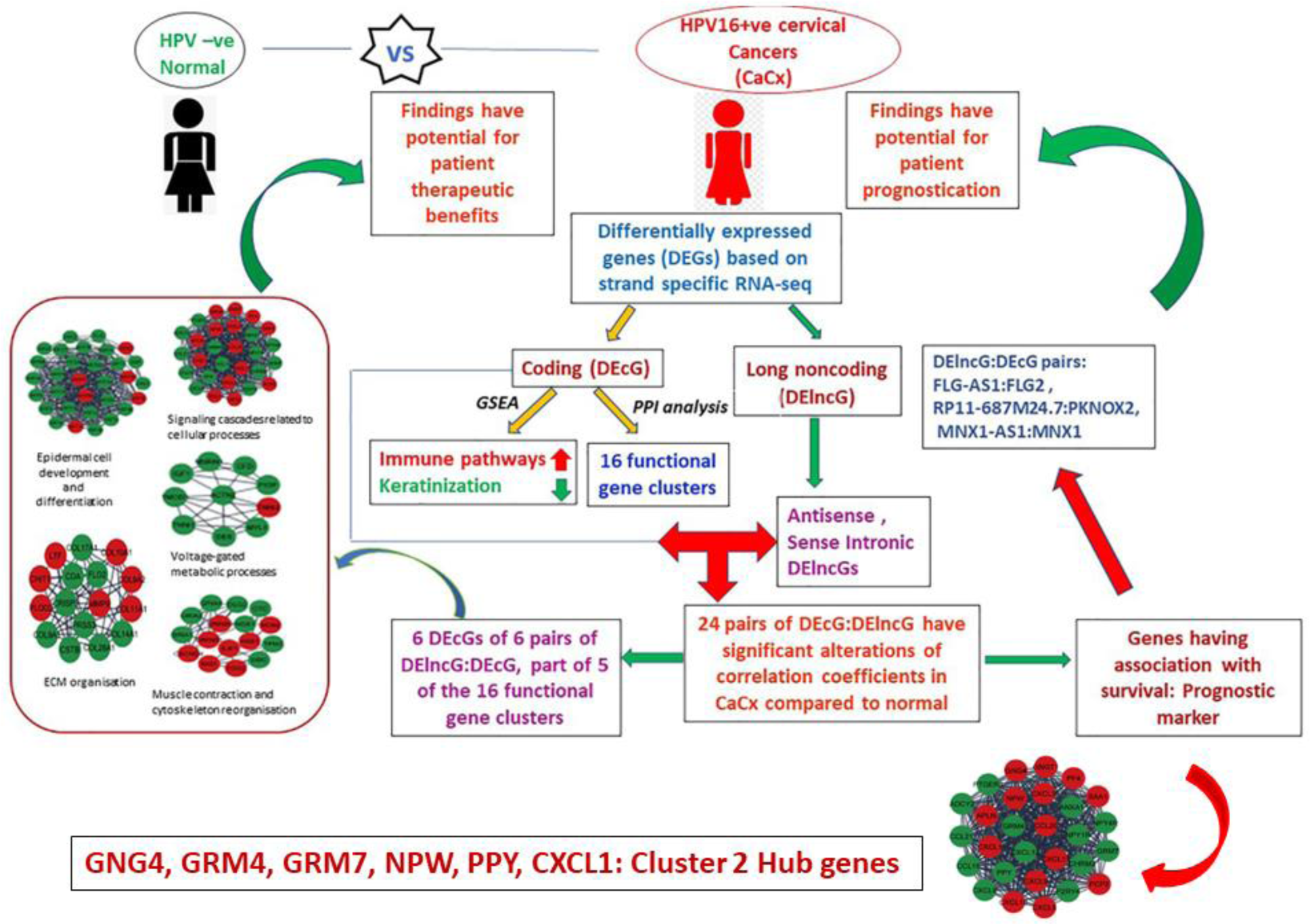
A schematic representation of the analyses undertaken to draw insights on the biological relevance of joint analysis of co-expressed and significantly correlated DElncGs and DEcGs considering the ncNATs and sense intronic DElncGs, in HPV16-positive CaCx patients compared with HPV-negative normal individuals.

## 1. Introduction

Cervical cancer (CaCx) continues to be the second most common cancer among Indian women with high mortality rates (http://cancerindia.org.in/globocan-2020-india-factsheet/). Majority of such cancers (>70%) are caused by persistent infection with high-risk human papillomavirus (HPV), especially HPV16 (1). Enhanced expression of HPV16 encoded oncoproteins E6 and E7 in the cervical epithelium, perturbs the p53 and pRb pathways in the host, leading to genomic instability and causing transformation of the cervical epithelium, that ultimately leads to invasive cancer (2,3). Thus, crosstalk between viral and host factors that impact upon host gene expressions are likely to influence HPV related CaCx pathogenesis.

In an earlier study (4), employing microarray-based assay, we showed that HPV16 E7 orchestrates the gene expression profiles of HPV16-positive CaCx cases by interacting with a long noncoding RNA (lncRNA), HOTAIR. A report on HPV16 positive CaCx (5) employing 3 such tumor tissues and matched adjacent normal samples identified the differential expression of lncRNAs in HPV-driven CaCx, with the potential of further being developed as diagnostic, prognostic and therapeutic markers of such cancers. The involvement of lncRNAs in HPV-driven cancers is also clearly depicted in a review by Casarotto et al (6). Current evidence also indicates that lncRNAs play a regulatory role in cancer progression (7). Specifically, lncRNAs, epigenetically regulate gene expression, support macromolecular complex assembly through scaffold formation, bind and inactivate miRNAs through a sponging effect and regulate mRNA stability (8). A proportion of lncRNAs also function as Natural Antisense Transcripts (ncNATs) and regulate the expression of sense coding gene at the same locus by various mechanisms (9). This prompted us to undertake this study to dissect the gene expression profiles (both coding and lncRNAs) associated with HPV16-positive CaCx patients, employing high-throughput strand-specific RNA sequencing (ssRNA-seq) technology, through which we can estimate the precise expression of various lncRNA types.

Some recent genome-wide studies (9,10) have provided evidence that within the same tissues or cells, there is a positive correlation between expression of sense genes and the corresponding antisense genes rather than a negative correlation. Therefore, perturbation of the expression of one or both partners of the ncNATs: protein coding gene pairs, associated with disease pathogenesis, could be of immense therapeutic relevance for various diseases including cancers (11).

Thus, we hypothesize that harnessing the expression profiles of lncRNAs using ssRNA-seq, together with the protein coding transcripts in CaCx tissues, is likely to provide novel insights on CaCx pathogenesis, and to pinpoint clinically relevant targets. To test this, we determined the biologically relevant genes and pathways portraying dysregulated expression in HPV16-positive CaCx patients, in comparison to HPV-negative normal individuals. We also investigated the nature of correlation in the expression levels between co-expressed noncoding (ncNAT/sense intronic) and protein coding gene pairs in patients and normal individuals. Subsequently, we considered only those gene pairs that revealed significantly altered correlation co-efficient in patients, as opposed to the normal individuals, for further interpretation of their biological relevance. We, thereby, identified putative actionable target genes of the correlated noncoding: protein coding gene pairs in patients, which belonged to biologically relevant gene clusters, identified through protein-protein interaction (PPI) analysis of our dataset. Finally, we determined the impact of the correlated genes on patient survival using the survival data available in our cohort.

## 2. Materials and Methods

### 2.1 Subjects and Samples

Tissue samples from married non-pregnant women, aged 28-50 years (median age: 43 years), who underwent hysterectomy and did not have any prior history of cervical dysplasia or malignancy were considered as healthy individuals or controls. Tissues from CaCx patients were collected from women aged 35-78 years (median age: 54 years) were considered as patients or cases. All The cervical biopsies were histopathologically confirmed as malignant or non-malignant.

### 2.2 Sample Processing and Sequencing

Cervical biopsy tissues were collected in RNAlater. Genomic DNA and total RNA were isolated from these tissues using QIAamp DNA mini kit and RNeasy mini kit (Qiagen) respectively, following the manufacturer’s protocol. Quality and integrity of RNA was determined using Agilent Bioanalyzer 2100. Details regarding DNA isolation, HPV screening and type identification, are described in our earlier studies (4,12). All the samples were tested for the presence of HPV and classified as HPV-negative or positive. Of the HPV positive samples, we selected only those with HPV16 infection as HPV16 is the most prevalent type in CaCx cases in India. Samples showing presence of both HPV18 and HPV16 were excluded from the study. For this study, we compared the HPV-negative normal samples (n=34) with HPV16-positive CaCx samples (n=44). We excluded the HPV positive normal samples because of uncertainty in ascertaining the transient or persistent nature of the infection.

About 500ng of total RNA isolated from these samples with RNA Integrity Number (RIN) ≥ 5.5, was considered for library preparation using TruSeq Stranded Total RNA Library Prep kit (Illumina). The library quality was assessed using Agilent Bioanalyzer 2100 system and then sequenced in Nova-Seq 6000 (Illumina) to generate paired-end reads of 100 bases. The details of the sequencing quality and coverage are provided in **Supplementary Table S1**.

### 2.3 Alignment of RNA sequencing data and identification of differentially expressed genes (DEGs)

A community standard approach was adopted for analysis that included using FastQC-0.11.7 (http://www.bioinformatics.bbsrc.ac.uk/projects/fastqc/) for QC analysis, STAR aligner (STAR-2.6.0c) (13) for alignment of RNA-sequence reads against human reference sequence (GRCh37 primary assembly) and HTSeq (HTSeq-0.11.0) (14) for generation of raw expression count data for all coding and noncoding genes, with stranded mode using Homo sapiens GRCh37.87 GTF (Ensembl) as the base sequence. To filter out genes with low count, only genes that were expressed (at least one transcript) in at least 50% samples in any of the two groups (a) HPV negative normal individuals, and (b) HPV16 positive CaCx patients or both, were included in further analysis. Differential gene expression analysis and generation of normalised expression counts were performed with DESeq2 (15) package using HTSeq-count data.

For the present analysis, to select the lncRNA genes, we took into consideration 3’ overlapping noncoding RNA, antisense, lincRNA, sense intronic, sense overlapping, and restricted to considering only those transcripts that were of length > 200 nucleotides, as identified from LNCipedia and Ensembl database. The remaining transcripts that did not fulfil these criteria were removed from the study. Single exon transcripts, if identified, were removed because most of such noncoding genes, by virtue of being novel, remain to be experimentally validated and may represent background noise from DNA contamination because these transcripts lack a splice junction (16).

For our analysis, we considered only the genes (coding and long noncoding) for which there was significant (p<0.05 after correction using the Benjamini-Hochberg [BH] procedure) differential expression and a high expression fold-change (FC), |log2(FC)| ≥ 2. TPM (transcripts per million) values were calculated using the TPMCalculator tool (17).

### 2.4 Expression analysis of DEGs in HPV16-positive patients compared to HPV-negative normal individuals using quantitative Real time PCR (qRT-PCR)

cDNA was generated from RNA (about 400ng) as described in our earlier study (4). The gene expression of the DEGs was determined in the HPV-negative healthy individuals (N=26) as well as in HPV16-positive CaCx patient tissue (N=26) samples using SYBR Green based qRT-PCR on ABI-Quant Studio 5. Glyceraldehyde 3-phosphate dehydrogenase (GAPDH) was used as internal control. The fold change was calculated by 2^(-ΔΔCt)^ method. The list of the primers used for the assay is provided in **Supplementary Table S2**.

### 2.5 Visualisation and characterisation of DEGs

For the visualisation of DEGs, volcano plot was generated using ggplot in R and the pheatmap R package (https://cran.r-project.org/web/packages/pheatmap/pheatmap.pdf) was used to perform unsupervised hierarchical clustering with normalised counts, which were log2 transformed after adding a constant (=1) to all values. Pathway analysis was performed using Gene Set Enrichment Analysis (GSEA) (18) and pathways were considered as significant if their FDR corrected *p* was < 0.05. PPI network of the Differentially Expressed coding genes (DEcGs) were predicted using the STRING (Search Tool for the Retrieval of Interacting Genes/Proteins) database Version 11.0 (19), considering an interaction score ≥ 0.7 (high confidence). The PPI network was visualised using Cytoscape (version 3.8.2) and the plug-in MCODE (Molecular Complex Detection) was used to extract functional gene clusters in the PPI network. Hub genes of the functional clusters were identified through Cytoscape with highest degree of connectivity. Functional enrichment of the gene members of the clusters was carried out using DAVID (Database for Annotation, Visualisation and Integrated Discovery) (20,21). Noncoding genes and their coding gene partners were identified from the Ensembl database. Survival analysis of the gene pairs was done employing KM Plotter (22), using default settings.

### 2.6 Statistical Analysis

Statistical analyses were done using R and online available tools. Pairwise Pearson’s correlation coefficients (r) between genes (antisense or sense lncRNA genes and their sense coding gene partners) were calculated using their normalised expression, i.e., TPM values. Differential correlation analysis of the gene pairs between cancer patients and healthy individuals was performed using Fisher’s z transformation of r (http://vassarstats.net/rdiff.html). The crude p values thus generated were further used to determine the adjusted p-values (after FDR correction) using an in-house developed R code. All the statistical tests were two-sided and a result was considered as significant when the adjusted *p* value was < 0.05 (after FDR correction).

## 3. Results

### 3.1 Expression patterns of coding and long noncoding RNA genes in HPV16-positive CaCx patients compared to HPV-negative healthy individuals

About 100 million RNA sequencing reads per sample mapped to a total of 55,638 genes (coding and noncoding). Of these, 22,208 genes expressed at least 1 transcript in at least 50% of HPV16-positive patients or HPV-negative normal individuals. These genes comprised 16,708 protein coding and 5,070 lncRNA genes (**Figure 1A**). For subsequent analysis, genes that were differentially expressed in CaCx patients as compared to HPV-negative healthy individuals, were identified based on |log2(FC)| ≥ 2, with FDR corrected *p* < 0.05. We identified 1486 DEcGs and 775 DElncGs, which are marked in red (upregulated genes) and blue (downregulated genes) on **Figure 1A** and the details are provided in **Supplementary Table S3**. Unsupervised hierarchical clustering using the Ward.D2 method and the Euclidean distance was then performed, which revealed contrasting expression profiles of both coding (**Figure 1B**) and noncoding genes (**Figure 1C**) in CaCx patients, as compared to normal individuals.

**Figure 1.**
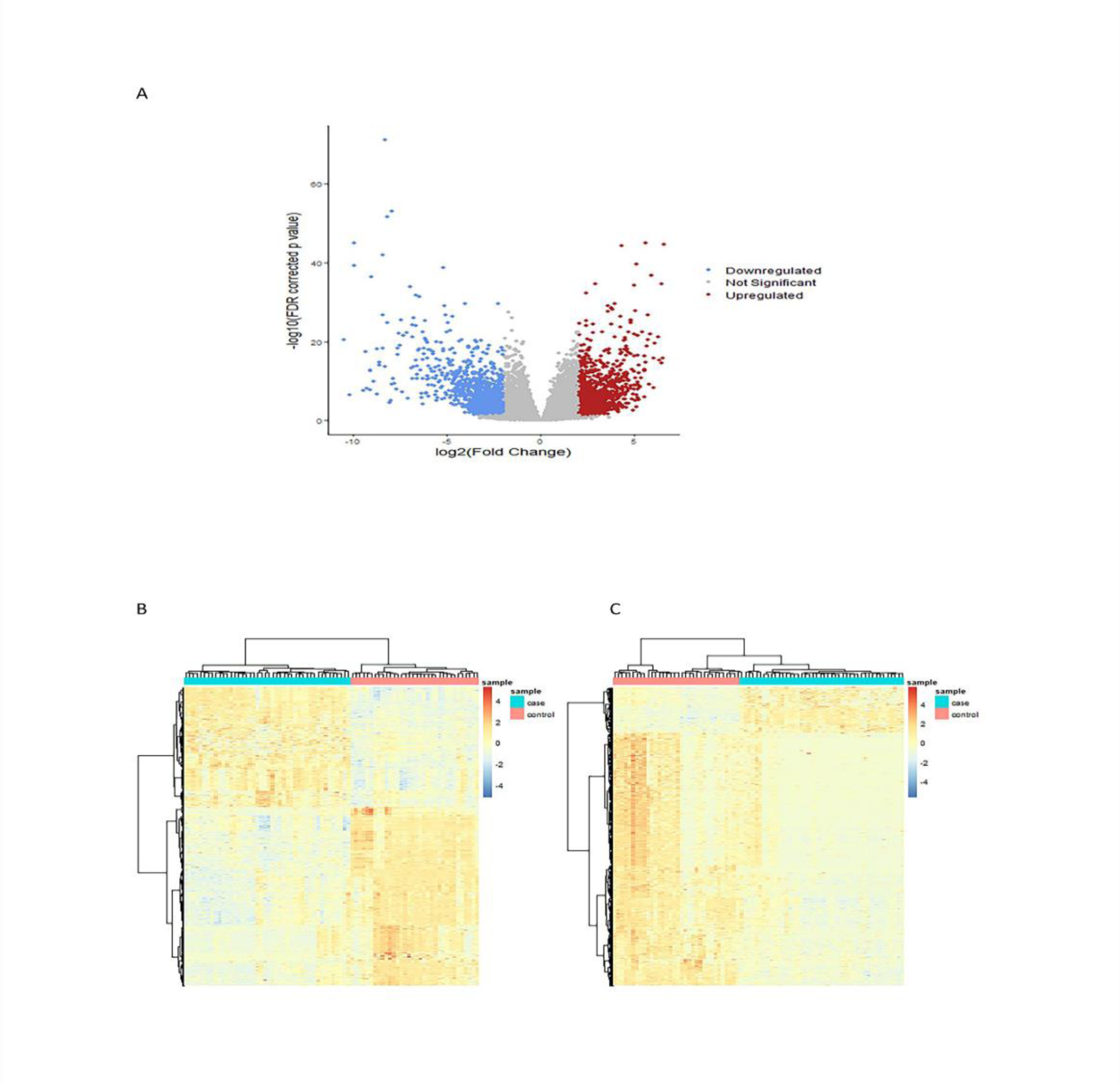
Visualisation of DEGs. (A)Volcano plot depicting the Differentially Expressed Genes (DEGs) in HPV16-positive CaCx patients as compared to HPV-negative normal individuals. The X-axis represents log2 (Fold Change) and Y-axis the -log10 (FDR corrected p value). Red dots represent the differentially expressed upregulated genes [FDR corrected *p* < 0.05 and log2(Fold Change) ≥ 2]. Blue dots represent the differentially expressed downregulated genes (FDR corrected *p* < 0.05 and log2(Fold Change) ≤ -2). Grey dots represent genes that were not differentially expressed. **(B-C) Unsupervised hierarchical clustering demarcating the gene expression profiles of CaCx patients from the normal individuals.** The Heatmap represents normalized counts, which were log2 transformed after adding constant 1 to all values of **(B)** the differentially expressed coding genes (DEcGs) and **(C)** the differentially expressed lncRNA genes (DElncGs). The rows represent the gene names, and the columns represent the samples (blue bar represents the CaCx patient samples and pink bar represents the samples from normal individuals).

### 3.2 Validation of the expression of some DEGs in HPV16-positive CaCx patients compared to HPV-negative normal individuals using qRT-PCR

We validated the expression levels of some of the DEGs (both coding and noncoding) using qRT-PCR. For validation, we selected the upregulated DEcG and DElncG (FOXD3 and RP4-792G4.2) out of the 3 pairs that were associated with patient-survival. Upon analysis significant upregulation of both FOXD3 and RP4-792G4.2 were observed among HPV16 positive CaCx patients compared to normal individuals, validating our finding from RNA-seq data. The fold change of the DEGs in patients and normal individuals are depicted in **Supplementary Figure S1**.

### 3.3 Pathways enriched and depleted in HPV16-positive CaCx patients in comparison with HPV-negative normal individuals

We then performed GSEA from MSigdb (msigdb.v7.4.symbols.gmt) considering only the GO (Gene Ontology) Biological Processes. In comparison to healthy individuals, genes involved in both humoral and cell mediated immune processes (Immunoglobulin family members, VTCN1, CD70, FOXJ1, LAMP3, ICAM4 etc) and cell cycle processes like cell migration, chromosome segregation (CDKN2A, SMC1B, MCM2, STAG3, E2F1) appeared to be significantly (FDR corrected *p* < 0.05) enriched in CaCx patients (**Supplementary Figure S2**). Similarly, the processes that were found to be significantly depleted in the CaCx patients included keratinization (KRT family genes), cornification (LCE family genes) and epithelial cell development and differentiation **(Supplementary Figure S3).** The details of all the processes are shown in **Supplementary Table S4.** This clearly reflected the functional distinctiveness of the depleted processes, as compared to the enriched processes.

Next, we explored the PPI network among the DEcGs by using the STRING database. We then determined the GO Biological processes corresponding to the upregulated and downregulated DEcGs from STRING, the details of which are provided in **Supplementary Table S5 and S6,** respectively. The findings replicated the processes identified through GSEA. In addition, the upregulated genes were mostly involved in chromatin assembly, cellular differentiation, cytokine mediated signalling pathways, DNA replication, G1-S transition, Extracellular Matrix Remodelling and Organization. The downregulated genes, on the other hand, were associated with cell development and differentiation, cellular signalling, cell-cell adhesion, cell death, lipoxygenase activity, etc.

### 3.4 Significantly correlated co-expression of DElncGs (ncNATs and sense intronic) with their corresponding sense DEcGs at the same location

We have previously identified global transcriptional reprogramming mediated by HPV16 encoded oncoprotein E7 through expression modulation and functional inactivation of lncRNA, HOTAIR (4). Therefore, in this study we focussed on DElncGs to determine their crosstalk with DEcGs in CaCx pathogenesis. Of the 775 DElncGs among the CaCx patients, 274 encoded ncNATs, 45 were sense intronic, while the remaining were other types of lncRNA genes. For further analysis, we considered only the ncNATs and sense intronic genes, because they are known to regulate their protein coding gene counterparts at the same locus (11), thus bearing biological relevance. We identified coding genes in the neighbourhood of these 319 noncoding genes (274 antisense and 45 sense intronic), through Ensembl database, followed by identification of 83 noncoding: coding gene pairs and a single sense intronic: ncNAT gene pair, for which both members of the pair were significantly and differentially expressed (|log2(FC)| ≥ 2 and FDR corrected *p* < 0.05).

Using the TPM values, we further determined that of the 83 ncNATs/sense intronic: coding gene pairs, 79 pairs were significantly and positively correlated (FDR corrected *p* <0.05), i.e., co-expressed, in both the CaCx patients and normal individuals, one pair (RP11-256L6.3: HAL) showed significant positive correlation among patients only, while two pairs (NOVA1-AS1:NOVA1 and RP1-40E16.11: TUBB2A) showed significant positive correlation only in the normal individuals. The remaining pair RP11-96C23.10: FAM25A failed to show significant correlation among both the patients and normal individuals. Both members of all these gene pairs showed expression dysregulation in the same direction, excepting for eight pairs. The details of the findings are provided in **Supplementary Table S7.** As functions of most of the noncoding partners remain unknown, correlation of these ncNATs/sense intronic genes with their neighbouring DEcGs, reveal their association in the regulation of their respective coding gene expression.

Of the 79 significantly correlated gene pairs in both patients and normal individuals, we identified 24 pairs that showed significant difference in the correlation coefficient between the patients and normal individuals. This gene set included 22 pairs of ncNATs: coding genes and 2 pairs of sense intronic: coding genes (RP11-687M24.7: PKNOX2 and RP11-483C6.1: NOVA1). The correlations were significantly higher in the patients than in the normal individuals for 17 pairs, while for the remaining the pattern was the opposite (**Supplementary Table S7)**. The significant alteration in the strength of correlation between such gene pairs in tumorigenesis, calls for functional validation of the finding. The remaining 55 correlated pairs, which did not show any difference in correlative strength between the normal and patient samples, may jointly play a role in cervical tissue specific functions.

### 3.5 Identification of putative actionable target gene pairs of DElncGs (ncNATs and sense intronic) and DEcGs associated with tumorigenesis

To decipher the functional relevance of the ncNAT/sense intronic: coding gene pairs in CaCx, we considered the 24 gene pairs that showed significant alteration in strength of correlation between the patients and the normal individuals (**Supplementary Table S7**), with 70% showing increased correlated coefficients as opposed to 30% with reduced correlated correlations. We further considered these for determining their potential as actionable targets of HPV16 related CaCx.

To accomplish this, we attempted to identify if any of the coding gene counterparts of these 24 gene pairs were part of functionally relevant gene clusters. Thus, we performed PPI network analysis of 1486 DEcGs recorded in our study, using STRING database (interaction score > 0.7). This revealed 1324 nodes (genes) and 3445 edges (interactions), as shown in **Supplementary Figure S4.** The genes clustered into 16 modules identified through MCODE plug-in (Degree cut-off ≥2, node score cut-off 0.2, K –core 2 and MCODE score≥5) of Cytoscape. Thereby we determined the GO processes corresponding to these clusters employing DAVID (FDR corrected *p*<0.05), revealing their association with cancer related functions. Of the 24 tumor specific ncNAT/sense intronic: coding gene pairs, 6 coding genes appeared to be a part of some of these gene clusters as depicted in **Figure 2 (A-E)** and the detailed analysis of these 6 clusters is summarised in **Table 1**. The details of the remaining clusters, which did not incorporate any of the coding genes of the 24-tumor specific ncNAT/sense intronic: coding gene pairs, are provided in **Supplementary Table S8.** Two of these 6 gene pairs had concordant upregulated expression (CTD-3214H19.6: PCP2 and RP11-845M18.6: KRT86), 3 pairs had concordant downregulated expression (AC019349.5: KRT13; FLG-AS1: FLG2 and AC053503.6: DES), and one pair (RP11-74H8.1: CACNG4) portrayed discordant expression with upregulation of the coding gene and downregulation of the noncoding partner, with respect to the healthy individuals. These gene clusters were associated with biological processes such as, signalling cascades related to cellular processes, Epidermal cell development and differentiation, ECM organisation, Voltage-gated metabolic processes and Muscle contraction and cytoskeleton reorganisation as indicated in **Table 1**. These findings indicate that perturbation of the correlated co-expression of these six noncoding: coding gene pairs is likely to impact the expression of other genes of the clusters, thereby, affecting key biological processes related to cancer development and progression.

**Figure 2:**
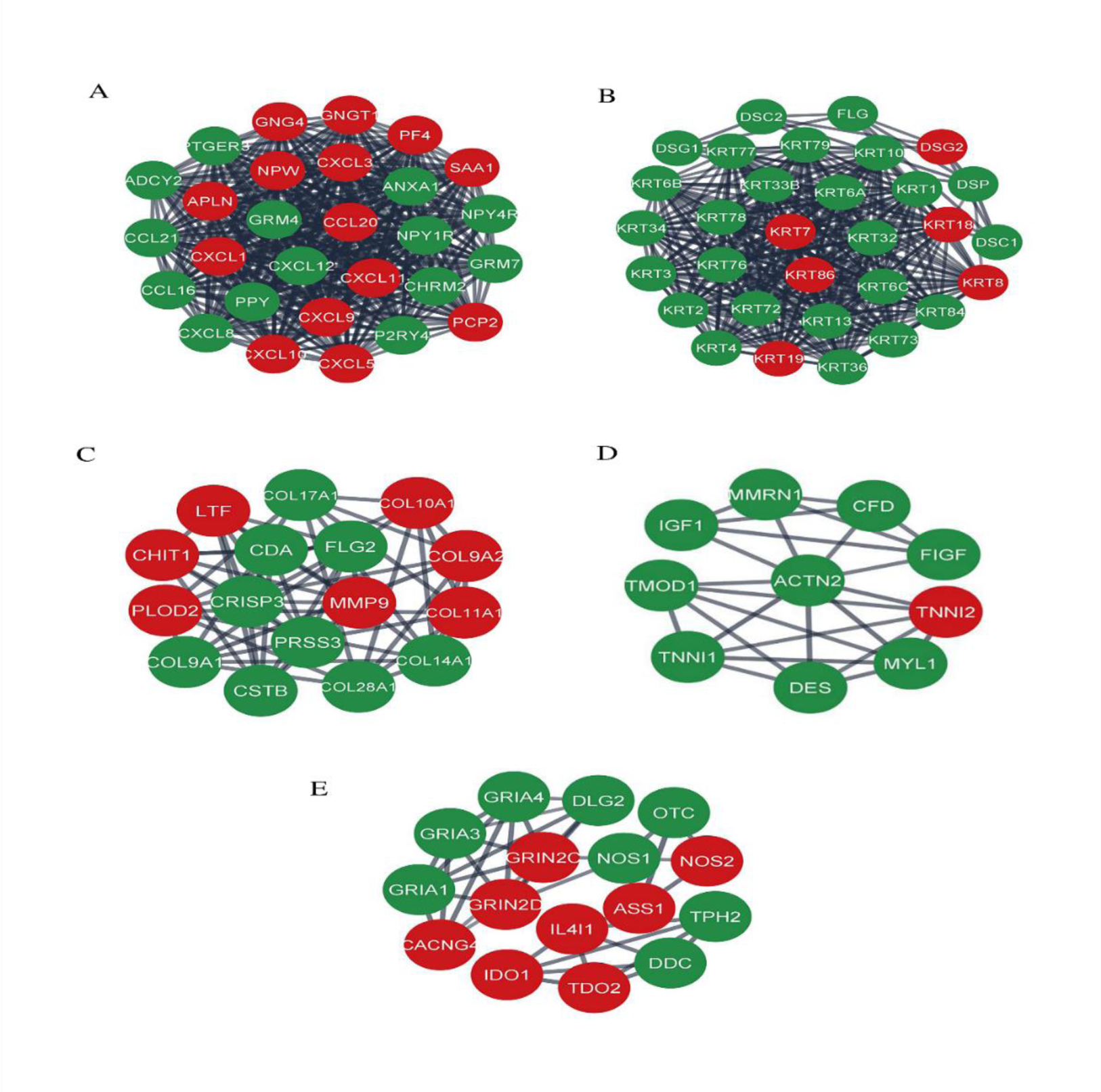
Visualisation of the gene clusters formed by considering the coding gene counterparts of the gene pairs DElncG (ncNAT/sense intronic):DEcG that revealed significantly altered correlation among HPV16-positive CaCx patients in comparison with HPV-negative normal individuals. The gene clusters correspond to **(A)** PCP2 (CTD-3214H19.6: PCP2), **(B)** KRT13 (KRT13:AC019349.5) and KRT86 (KRT86:RP11-845M18.6), **(C)** FLG2 (FLG2: FLG-AS1), **(D)** DES (DES: AC053503.6), **(E)** CACNG4 (CACNG4:RP11-74H8.1). The green nodes in the clusters represent genes with downregulated expression, whereas red nodes represent the genes with upregulated expression in CaCx patients as compared to normal individuals.

**Table 1:**
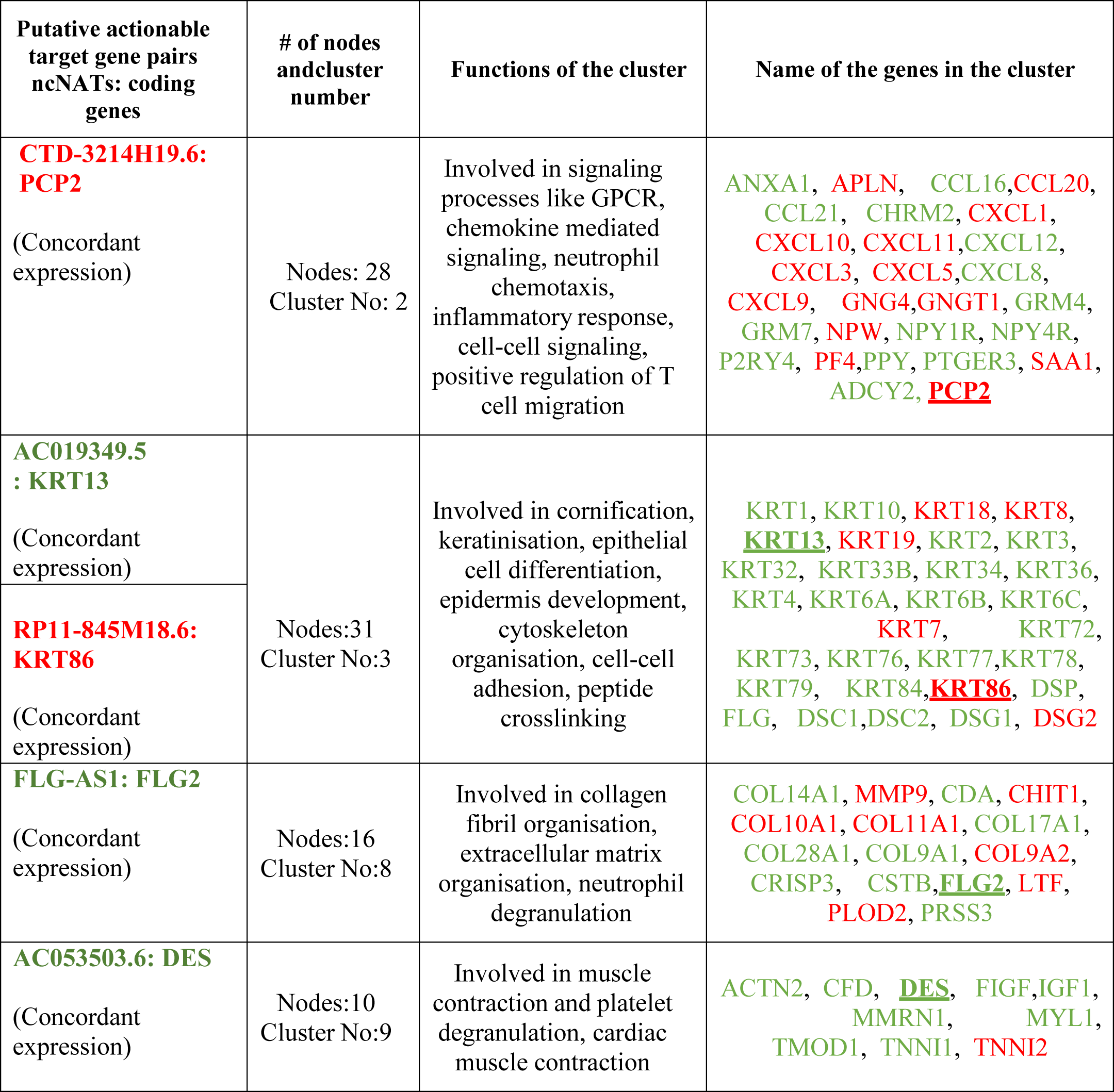
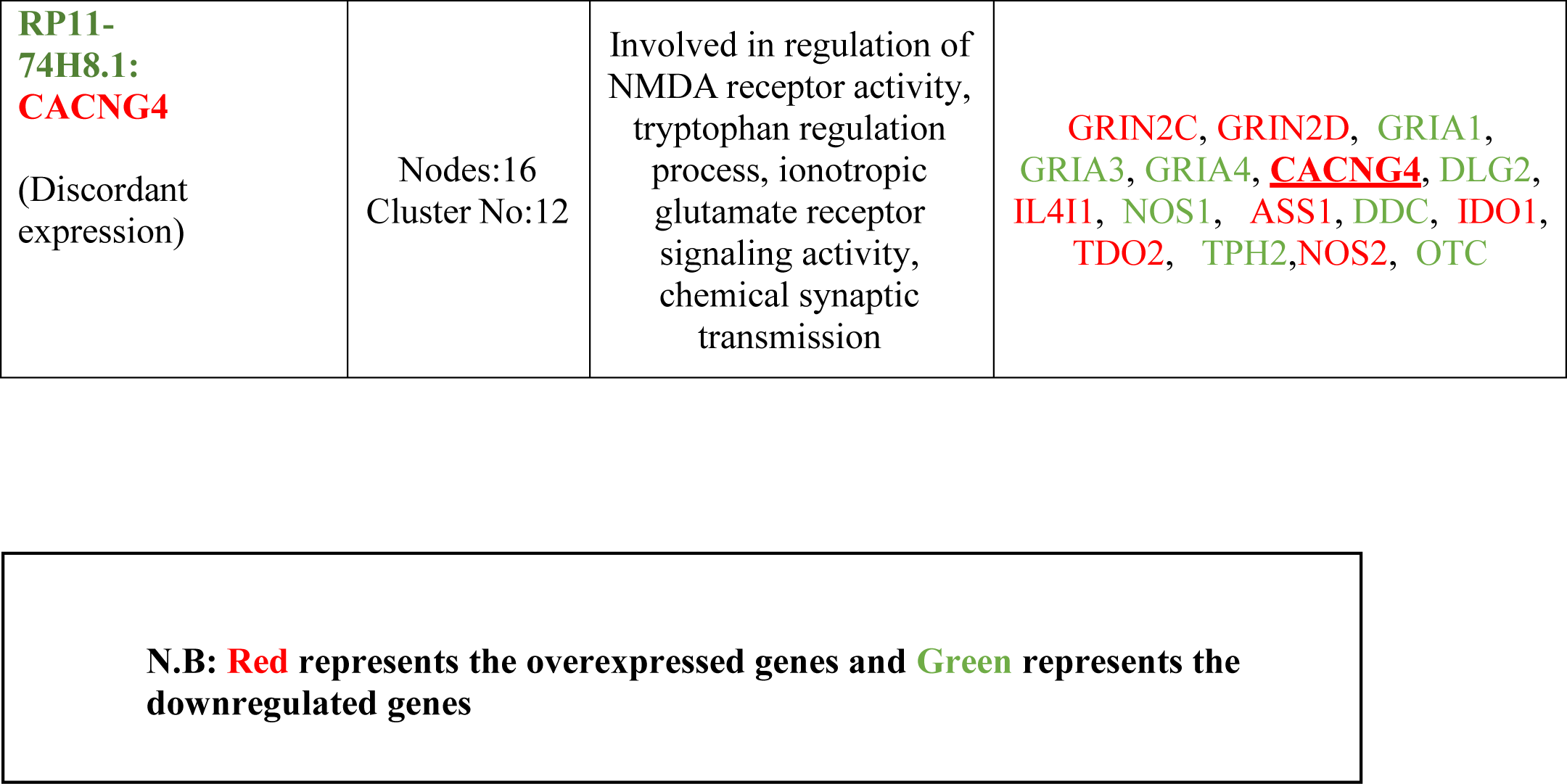
Clusters of coding genes associated with specific cancer related functions and incorporating the coding genes of the ncNATs:coding gene pairs portraying significant altered correlated co-expression in CaCx patients as compared to normal individuals.

| Putative actionable target gene pairs<br>ncNATs: coding genes | # of nodes and cluster number | Functions of the cluster | Name of the genes in the cluster |
| --- | --- | --- | --- |
| **CTD-3214H19.6: PCP2**<br>(Concordant expression) | Nodes: 28<br>Cluster No: 2 | Involved in signaling processes like GPCR, chemokine mediated signaling, neutrophil chemotaxis, inflammatory response, cell-cell signaling, positive regulation of T cell migration | ANXA1, APLN, CCL16, CCL20, CCL21, CHRM2, CXCL1, CXCL10, CXCL11, CXCL12, CXCL3, CXCL5, CXCL8, CXCL9, GNG4, GNGT1, GRM4, GRM7, NPW, NPY1R, NPY4R, P2RY4, PF4, PPY, PTGER3, SAA1, ADCY2, **PCP2** |
| **AC019349.5 : KRT13**<br>(Concordant expression) | Nodes: 31<br>Cluster No: 3 | Involved in cornification, keratinisation, epithelial cell differentiation, epidermis development, cytoskeleton organisation, cell-cell adhesion, peptide crosslinking | KRT1, KRT10, KRT18, KRT8, **KRT13**, KRT19, KRT2, KRT3, KRT32, KRT33B, KRT34, KRT36, KRT4, KRT6A, KRT6B, KRT6C, KRT7, KRT72, KRT73, KRT76, KRT77, KRT78, KRT79, KRT84, **KRT86**, DSP, FLG, DSC1, DSC2, DSG1, DSG2 |
| **RP11-845M18.6: KRT86**<br>(Concordant expression) |  |  |  |
| **FLG-AS1: FLG2**<br>(Concordant expression) | Nodes: 16<br>Cluster No: 8 | Involved in collagen fibril organisation, extracellular matrix organisation, neutrophil degranulation | COL14A1, MMP9, CDA, CHIT1, COL10A1, COL11A1, COL17A1, COL28A1, COL9A1, COL9A2, CRISP3, CSTB, **FLG2**, LTF, PLOD2, PRSS3 |
| **AC053503.6: DES**<br>(Concordant expression) | Nodes: 10<br>Cluster No: 9 | Involved in muscle contraction and platelet degranulation, cardiac muscle contraction | ACTN2, CFD, **DES**, FIGF, IGF1, MMRN1, MYL1, TMOD1, TNNI1, **TNNI2** |
| **RP11-74H8.1:**<br>**CACNG4**<br>(Discordant expression) | Nodes:16<br>Cluster No:12 | Involved in regulation of NMDA receptor activity, tryptophan regulation process, ionotropic glutamate receptor signaling activity, chemical synaptic transmission | **GRIN2C, GRIN2D, GRIA1, GRIA3, GRIA4, <u>CACNG4</u>, DLG2, IL4I1, NOS1, ASS1, DDC, IDO1, TDO2, TPH2, NOS2, OTC** |

**N.B: Red** represents the overexpressed genes and **Green** represents the downregulated genes

### 3.6 Deciphering the prognostic implication of tumor-specific significantly correlated DElncGs (ncNATs and sense intronic):DEcGs pairs in CaCx patients

To estimate the prognostic value of the 24 co-expressed and significantly correlated DElncG: DEcG pairs in CaCx patients, we determined the impact of the expression status of these coding and noncoding genes on patient overall survival (∼ 4 years) in our dataset, employing KM Plotter with default settings. Thereby, we identified 2 gene pairs with concordant downregulated expression (FLG-AS1:FLG2 and RP11-687M24.7:PKNOX2) and 1 pair (MNX1-AS1:MNX1) with concordant upregulated expression among patients, as survival associated genes. Both members of the 3 gene pairs at high expression levels, were significantly (p<0.05) associated with increased patient overall survival compared to those patients with low expression levels, as indicated in **Figure 3 (A-F)**.

**Figure 3:**
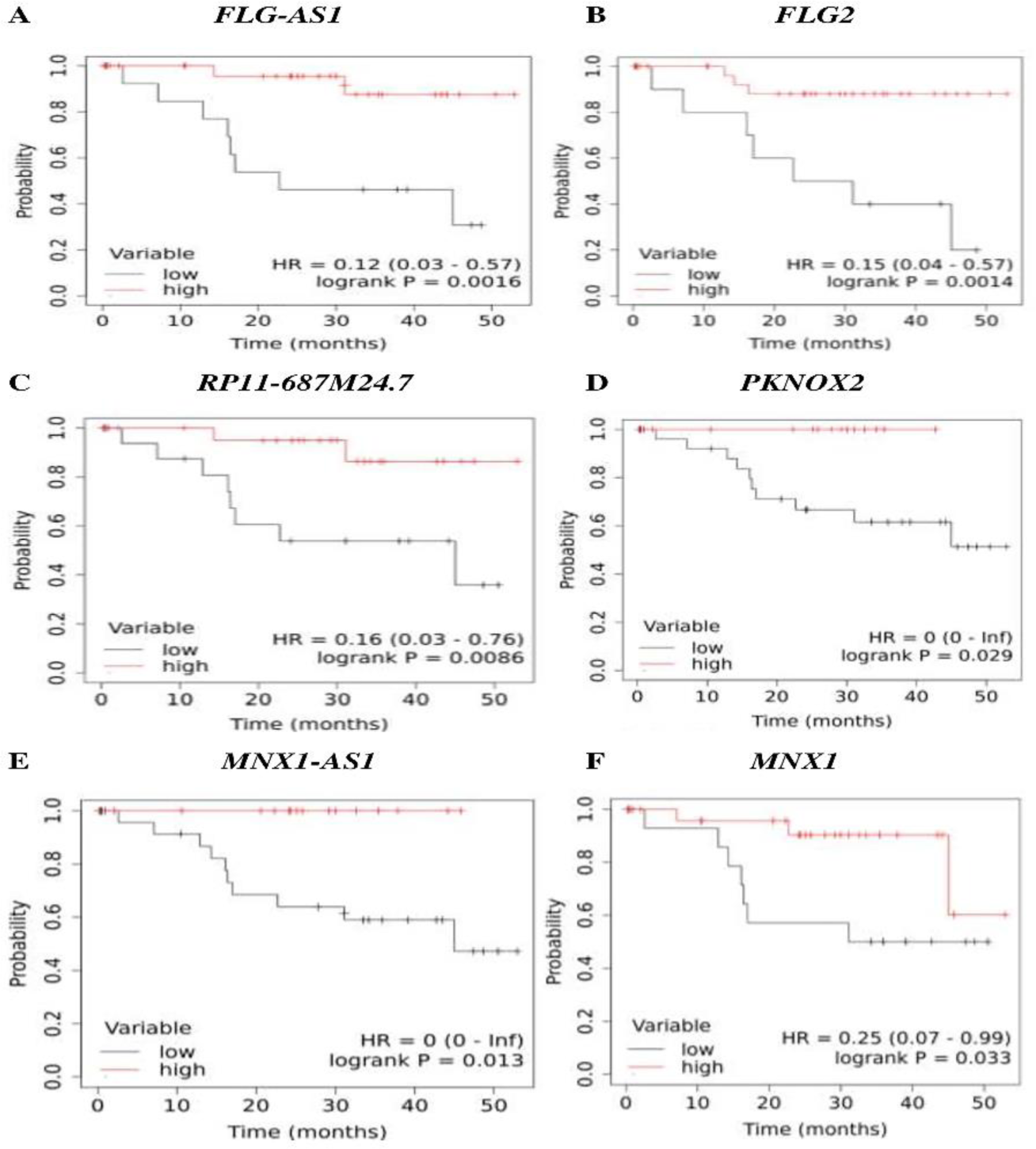
Impact of the expression status on patient overall survival of coding and noncoding counterparts of the DElncG (ncNAT/sense intronic):DEcG gene pairs that revealed significantly altered correlation among HPV16-positive CaCx patients in comparison with HPV-negative normal individuals. **(A-F)** Kaplan-Meier overall survival curve depicting CaCx patients with high expression and low expression of **(A)** FLG-AS1 (DElncG), **(B)** FLG2 (DEcG), **(C)** RP11-687M24.7 (DElncG), **(D)** PKNOX2 (DEcG), **(E)** MNX1-AS1 (DElncG) and **(F)** MNX1 (DEcG), employing KM-Plotter.

Furthermore, to understand the prognostic relevance of the 5 gene clusters, we identified the hub genes of each of these 5 clusters based on the higher degree of connectivity. The hub genes of each cluster and the corresponding degree of connectivity are represented in **Supplementary Table S9**. The prognostic value of each of these hub genes was explored by Kaplan-Meier Plotter. Among these clusters, all genes belonging to Cluster 2 were densely interconnected with each other, based on which, these were considered as the hub genes. Most of the genes (PPY, CXCL1, GRM4, GRM7, GNG4 and NPW) belonging to this cluster showed significant association with patient survival. Association of higher expression with better patient survival was recorded for PPY, GRM4, GRM7, GNG4 and NPW, while CXCL1 was the only hub gene that was associated with better patient survival at lower expression. Therefore, this cluster of genes appears to be significantly associated with patient survival. The survival curves are depicted in **Figure 4 (A-F)**. Moreover, this cluster incorporated the hub gene PCP2, which was significantly correlated with its ncNAT partner, CTD-3214H19.6 and both genes failed to reveal any association with patient survival. On the other hand, the survival associated gene pair, FLG-AS1:FLG2, belonged to cluster 8, where none of the coding hub genes revealed any association with patient survival.

**Figure 4:**
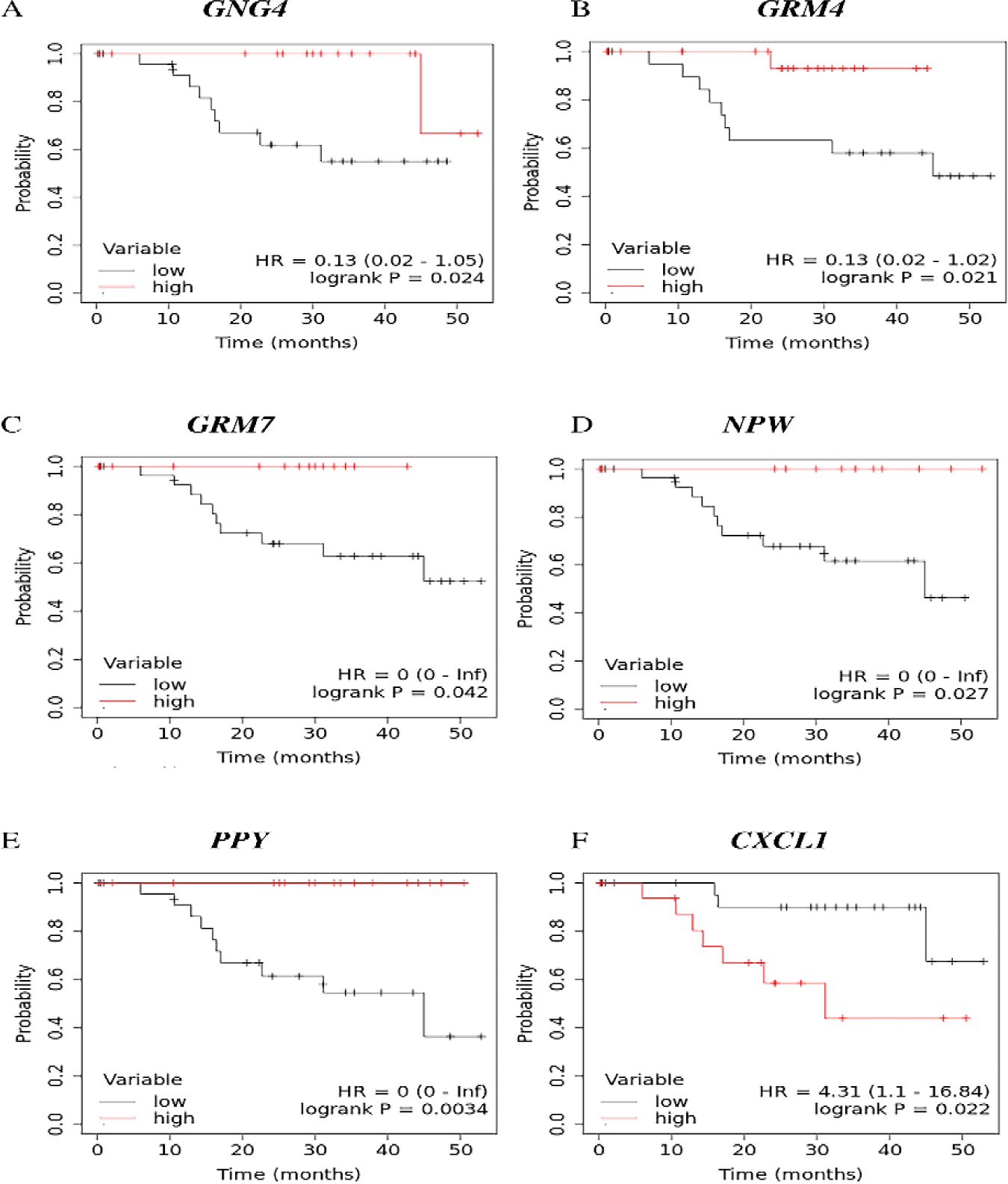
Impact of the expression status on patient overall survival of coding hub genes belonging to the cluster 2 incorporating the coding gene of the ncNATs:coding gene pairs portraying significant altered correlated co-expression in CaCx patients as compared to normal individuals. **(A-F)** Kaplan-Meier overall survival curve depicting CaCx patients with high expression and low expression of **(A)** GNG4, **(B)** GRM4, **(C)** GRM7, **(D)** NPW, **(E)** PPY and **(F)** CXCL1, employing KM-Plotter.

Thus, a substantial proportion of the coding genes and/or their noncoding (antisense/sense intronic) counterparts of such gene pairs with altered correlative strength in patients, appear to be of prognostic relevance in our patient cohort. The observation thereby points towards the biological relevance of such correlations in HPV16-positive CaCx patients.

## 4 Discussion

To our knowledge, this is the first study to report the global transcriptome profile (including both coding and noncoding) of HPV16 related CaCx in Indian population employing patient and normal individuals with the help of ssRNA-seq. The functions of a large majority of identified DElncGs remain to be characterized and reported in CaCx. Therefore, we aimed to identify crosstalk, if any, between the DElncGs (focussing on only the ncNATs and sense intronic genes) and their corresponding DEcG counterparts, to draw insights related to their functions based on those of the coding gene partners. Further, we determined the biological relevance of such interactions pertaining to identification of putative actionable targets and prognostic markers of HPV16-positive CaCx.

The ncNATs and sense intronic lncRNAs are known to affect expression of their proximal coding genes through multiple ways (23) and their dysregulated expression have been recorded among various cancer types (24,25) including CaCx (26,27). These genes, therefore, play an important role in cancer development, progression (23,28,29), therapeutics (11) and could also serve as biomarkers of cancer pathogenesis (30). We observed that several differentially expressed ncNATs and sense intronic genes in the CaCx patients showed significant correlated co-expression with the corresponding proximal DEcGs, concomitant with altered correlative relationships (both stronger and weaker), as compared to the normal individuals. Our study, therefore, is the first of its kind to reveal the significant joint role of correlated ncNATs/sense intronic genes and their coding gene counterparts at the expression level in HPV16 related CaCx pathogenesis. This underscores the existence of regulatory circuits between such noncoding and coding gene pairs. Such findings are similar to recent studies on other cancers (9,31), which recorded that a large majority of the ncNATs revealed positive and significant correlated co-expression with their corresponding sense genes, with stronger correlations among the tumors than in the normal samples. We also identified a large majority of such correlations, which failed to differ in correlative strength between cancer and normal individuals, justifying their roles in cervical tissue house-keeping functions. Taken together, our analysis strongly supports the biological relevance of correlated co-expression of ncNATs and sense intronic genes with their sense coding genes, both in cervical carcinogenesis as well as in cervical tissue specific functions.

In addition, we identified some functionally relevant gene clusters through PPI analysis of the DEcGs. These clusters were associated with the cancer related processes like keratinocyte and epidermal cell differentiation, cellular signalling cascades for proliferation, migration, cell cycle regulation, post translational modifications, interferon and neutrophils mediated immune pathways, extracellular matrix organisation etc. Thus, these gene clusters could potentially be highlighted as therapeutic targets, subjected to identification of ways and means to target them. We addressed this issue by examining the crosstalk between DElncG (ncNATs and sense intronic):DEcG pairs where the DEcG counterparts belong to such clusters. Our analysis reflected that the coding genes of 6 such gene pairs, were part of 5 large coding gene clusters, comprising of functionally related genes of cancer-associated pathways. Based on this observation, we propose that perturbations of the differential correlative relationships of the noncoding:coding gene pairs, where the coding gene counterpart appears to be a part of a functionally relevant gene cluster associated with CaCx pathogenesis, could serve as an option of disrupting the entire cluster (**Table 1**). This could thereby open up an interesting avenue of tackling such cancers.

In recent times, targeting ncNATs, to perturb the correlated ncNAT: coding gene pairs by use of antisense oligos (ASOs), is under exploration and some ASOs have been approved by FDA for the treatment of various diseases including cancers (32). Apart from this, some other possibilities based on advanced genome modification techniques (33) could also be employed for this. Our study therefore lays the foundation for exploring such possibilities. Also, the findings strongly confirm the participation of the DElncGs of these 6 gene pairs in cancer-associated mechanisms, pertaining to HPV16 positive CaCx cases.

Besides revealing the merit of altered significant correlated co-expressions of the ncNAT/sense intronic and coding gene pairs in CaCx patients as therapeutically relevant, our study also depicted the prognostic relevance of such pairs, involving both members (FLG-AS1:FLG2, MNX1-AS1:MNX1 and RP11-687M24.7:PKNOX2). As these genes show association with cancer development and are correlated at the expression level, therefore they may also seem to be of importance as prognostic targets, identified for the first time in case of HPV16 related CaCx.

Recent Pan-cancer studies (34) have provided convincing data that underscores the identification of pathway based biomarkers, which happen to be more proficient than single gene biomarkers. With respect to this, we moved a step further to show that of the 5 functionally relevant gene clusters in CaCx that incorporated 6 significantly correlated noncoding:coding gene pairs, cluster number 2 associated with various signaling processes and incorporating the CTD-3214H19.6: PCP2, also could be characterized as a patient survival-associated gene cluster. In fact, all genes of this cluster including PCP2, were hub genes and 6 such hub genes revealed association with patient survival. While both members of the gene pair CTD-3214H19.6: PCP2 were not associated with survival of patients, however, disruption of their correlated co-expression, could affect gene cluster 2, and hence patient survival. Therefore, this gene pair could potentially be considered as a prognostic biomarker of HPV16 related CaCx.

## 5. Conclusion

Through GSEA and PPI analysis of DEcGs, we identified enrichment of the processes crucial for abortive virus life cycle, cancer progression, immunity, and depletion of epithelial cell differentiation. The dysregulated DEcGs also formed several clusters, with characteristic cancer-related functions. In addition, we unfurled significant correlated co-expression of DElncGs (sense intronic and ncNATs) with DEcGs at proximal genomic loci within our sample cohort. A subset of these gene pairs portrayed significant alterations of correlation coefficients among patients than normal samples. The DEcGs of some such gene pairs showing altered correlative strengths, were part of a few gene clusters implicated in CaCx, including a survival associated gene cluster. Thus, such gene pairs portray the potential for serving as both therapeutic and prognostic targets. Three such pairs were also identified, where expression of both the members of the pairs were significantly associated with overall patient survival. Our study strongly highlights the cooperative role of DElncGs and DEcGs in cervical carcinogenesis, with the potential of translation.

## Authors’ contributions

AG^1*^ AS^1*^ and SSG^1^ conceived the study; AG^1*^, AS^1*^, SR^1^, SM^1^ and VK^1^ performed the experiments, AG^1*^,AS^1*^, NKB^1^ and SSG^1^ wrote the manuscript; NKB^1^ and AG^1^^ developed the RNA-seq data analysis pipeline and was involved in raw data processing, AG^1*^, AS^1*^ and SSG^1^ performed downstream data analysis, SM^2^, JB^2^, AM^2^, SS^2^, AC^2^ coordinated patient recruitment, clinical assessments and collection of data on patients and provided patient materials; AM^1^ coordinated massively parallel sequencing experiments; PPM^1^ provided guidance of the study design and analysis and SSG^1^ provided overall guidance of the study; all authors contributed towards reviewing the manuscript.

## Abbreviations

DEG: Differentially Expressed Gene
DEcG: Differentially Expressed coding Gene
DElncG: Differentially Expressed long noncoding RNA Gene
FDR: False Discovery Rate
FIGO: International Federation of Gynecology and Obstetrics
GSEA: Gene Set Enrichment Analysis
HPV: Human Papillomavirus
LncRNA: Long noncoding RNA
ncNAT: Noncoding Natural Antisense Transcript
NGS: Next Generation Sequencing
PPI: Protein-Protein Interaction
RIN: RNA Integrity Number
STRING: Search Tool for the Retrieval of Interacting Genes/Proteins
TPM: Transcripts Per Million

## Acknowledgements

We are grateful to the Government of India, Department of Biotechnology, Ministry of Science and Technology, for financial support to this study, through the Systems Medicine Cluster (SyMeC) project (BT/Med-II/NIBMG/SyMeC/2014/Vol. II). We are also grateful to all participating members of SyMeC for their inputs and contributions to this study. We acknowledge Tata Memorial Centre, Kolkata, and College of Medicine and J.N.M. Hospital, Kalyani, for providing us with the clinical samples for the study; all the members of the Core Technology Research Initiative (CoTeRI) of the National Institute of Biomedical Genomics, Kalyani, India for their technical support during the work; and special thanks also to Council of Scientific and Industrial Research (CSIR), India, for providing research fellowships to Abhisikta Ghosh (JRF and SRF), Department of Science and Technology (INSPIRE), Govt. of India for providing research fellowship to Abarna Sinha (JRF and SRF), Department of Biotechnology, Government of India, for funding Sumana Mallick (Project RA), and Department of Science and Technology, Govt. of India for funding Vinoth Kumar (National Post-doctoral Fellowship), to work on this project. We also acknowledge the support of Prof. Indranil Mukhopadhyay (Indian Statistical Institute, Kolkata), Dr. Samsiddhi Bhattacharjee and Ms. Sahana Ghosh (National Institute of Biomedical Genomics, Kalyani) and Dr. Shreoshi Sengupta (Indian Institute of Science, Bangalore) in data analyses.

## Data availability Statement

Raw RNA seq BAM files of individual patients can be accessed from European Nucleotide Archive (ENA) via accession number: PRJEB40877 and secondary accession number: ERP124576. Raw data will be made available upon request.

## Ethics Statement

All samples, from both healthy individuals and cancer patients, were collected with written informed consent. The study was approved by the Institutional Ethics Committees of all collaborating institutions (Tata Medical Centre, Kolkata and College of Medicine and Jawaharlal Nehru Memorial Hospital, Kalyani) and National Institute of Biomedical Genomics, Kalyani.

## Supplementary Figure Legends

**Supplementary Figure S1: Validation of the expression of DEGs in HPV16-positive CaCx patients compared to HPV-negative normal individuals using quantitative Real time PCR (qRT-PCR).** Bar diagram representing the fold change of the mRNA expression levels of the DEGs-**(A)** FOXD3 and **(B)** RP4-792G4.2 in HPV16-positve CaCx patients compared to HPV-negative normal individuals.

**Supplementary Figure S2**: **Gene Set Enrichment Analysis (GSEA) considering DEcGs representing significantly enriched GO Biological processes in HPV16-positive CaCx patients.** Graphical representation depicting the enrichment plots of the top enriched GO processes: **(A)** Adaptive Immune Response **(B)** Phagocytosis **(C)** B cell mediated immunity **(D)** Immune response regulating signalling pathway **(E)** Lymphocyte mediated immunity **(F)** Complement activation **(G)** Regulation of Complement Activation **(H)** Fc Receptor Signalling Pathway and **(I)** Immunoglobulin Production. The pathways were considered significant for FDR corrected p < 0.05.

**Supplementary Figure S3: Gene Set Enrichment Analysis (GSEA) considering DEcGs representing significantly depleted GO Biological processes in HPV16-positive CaCx patients.** Graphical representation of the enrichment plots of the depleted GO processes: **(A)** Skin development **(B)** Keratinocyte Differentiation **(C)** Keratinization **(D)** Epidermal cell differentiation **(E)** Epidermis development **(F)** Epithelial cell differentiation **(G)** Cornification **(H)** Epithelial development and **(I)** Peptide crosslinking. The pathways were considered significant for FDR corrected p < 0.05.

**Supplementary Figure S4: Protein-Protein Interaction (PPI) network.** PPI network (interaction score > 0.7, high confidence) was constructed with the DEcGs identified among the HPV16 positive CaCx patients as compared to normal individuals considered in this study, using STRING database, and visualised through Cytoscape.

## Supplementary Table Headings

**Supplementary Table S1**: **Details of the sequencing quality and coverage for all sequenced samples (44 HPV16 positive CaCx patients and 34 HPV-negative normal individuals).**

**Supplementary Table S2: List of primers used for the estimation of RNA expression of the selected DEGs by qRT-PCR**

**Supplementary Table S3**: **Details of all differentially expressed genes (DEcGs and DElncGs) in HPV16-positive CaCx patients as compared to HPV-negative normal individuals portraying expression difference of fold change (FC) cut-off |log2(FC)| ≥ 2 with False Discovery Rate (FDR) corrected p < 0.05.**

**Supplementary Table S4**: **Details of significantly (FDR corrected p <0.05) enriched and depleted GO Biological processes in HPV16-positive CaCx patients as compared to HPV-negative normal individuals obtained from Gene Set Enrichment Analysis (GSEA).**

**Supplementary Table S5**: **Details of the GO Biological processes corresponding to the upregulated DEcGs from STRING database.**

**Supplementary Table S6**: **Details of the GO Biological processes corresponding to the downregulated DEcGs from STRING database.**

**Supplementary Table S7: Differential correlation of the noncoding:coding gene pairs among the HPV16 positive CaCx patients and HPV negative normal individuals.**

**Supplementary Table S8**: **Details of the gene clusters, that excluded DEcGs of significantly correlated ncNAT/sense intronic: coding gene pairs, identified through PPI network analysis of all DEcGs employing the MCODE plug-in of Cytoscape, along with their corresponding functions determined through DAVID.**

**Supplementary Table S9: The details of the hub coding genes of the clusters incorporating the coding genes of the ncNATs:coding gene pairs portraying significant altered correlated co-expression in CaCx patients and their association with overall patient survival.**

