## Supplementary Materials for "Prognostic relevance of correlated co-expression of coding and noncoding RNAs in cervical cancers"

**Supplementary Figure S1**

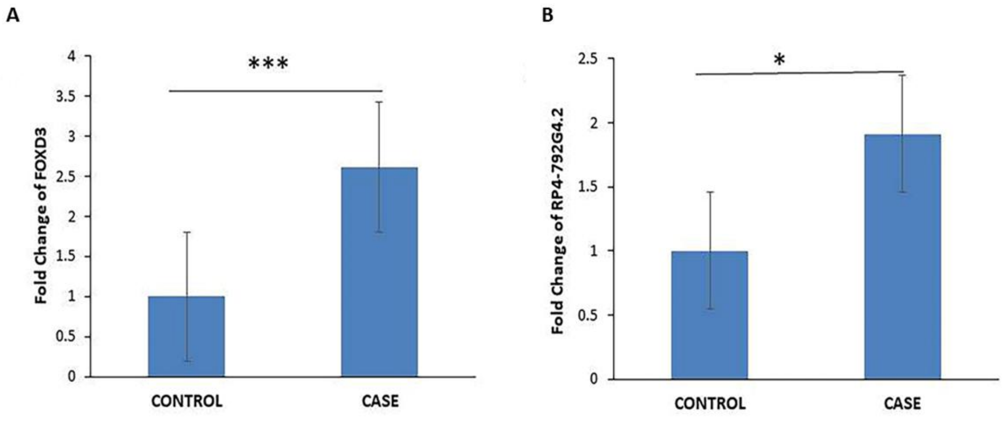


**Supplementary Figure S1: Validation of the expression of DEGs in HPV16-positive CaCx patients compared to HPV-negative normal individuals using quantitative Real time PCR (qRT-PCR).** Bar diagram representing the fold change of the mRNA expression levels of the DEGs- **(A)** FOXD3 and **(B)** RP4-792G4.2 in HPV16-positve CaCx patients compared to HPV-negative normal individuals.

**Supplementary Figure S2**

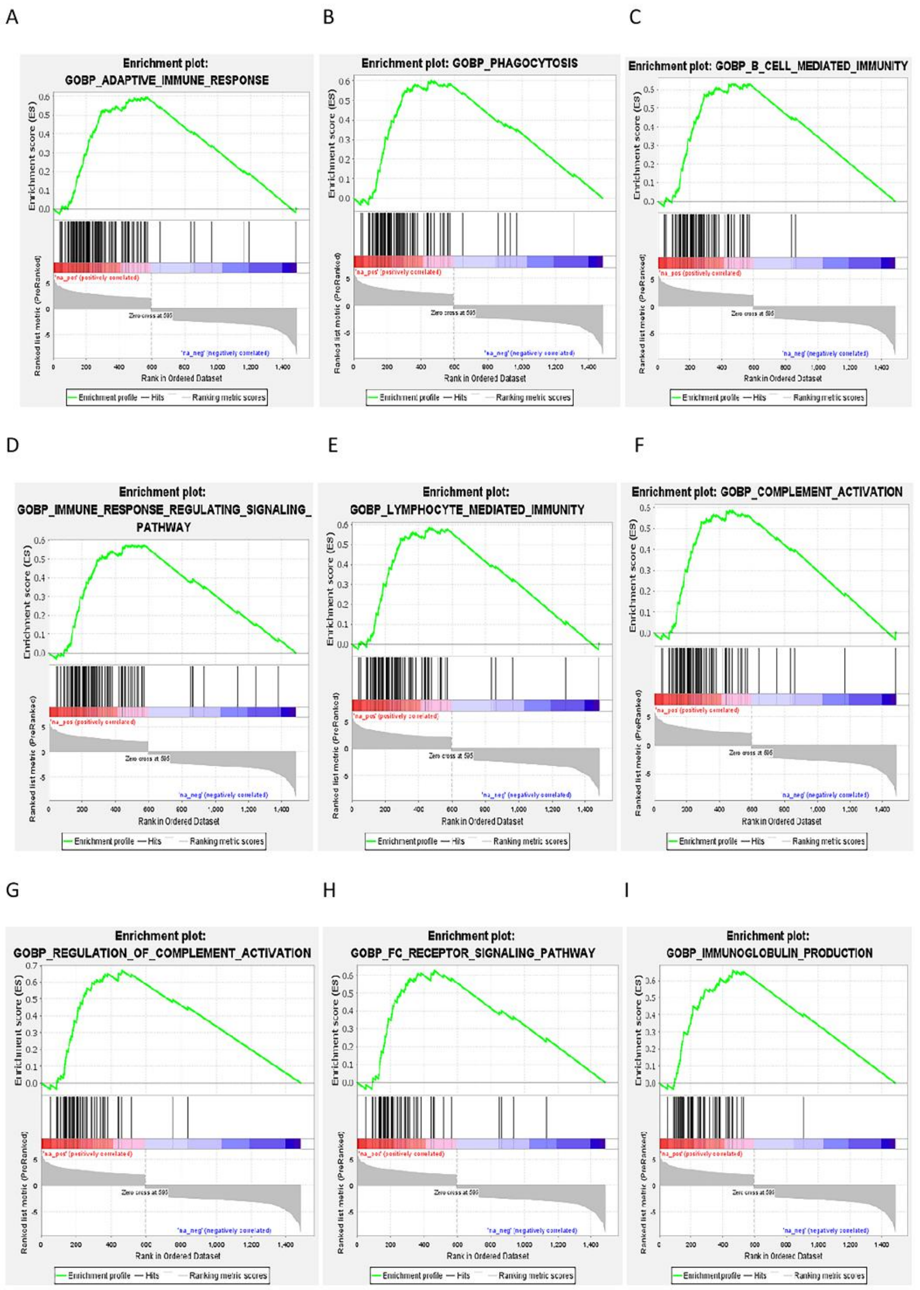


**Supplementary Figure S2**: **Gene Set Enrichment Analysis (GSEA) considering DEcGs representing significantly enriched GO Biological processes in HPV16-positive CaCx patients.** Graphical representation depicting the enrichment plots of the top enriched GO processes: **(A)** Adaptive Immune Response **(B)** Phagocytosis **(C)** B cell mediated immunity

**(D)** Immune response regulating signalling pathway **(E)** Lymphocyte mediated immunity **(F)** Complement activation **(G)** Regulation of Complement Activation **(H)** Fc Receptor Signalling Pathway and **(I)** Immunoglobulin Production. The pathways were considered significant for FDR corrected p < 0.05.

**Supplementary Figure S3**

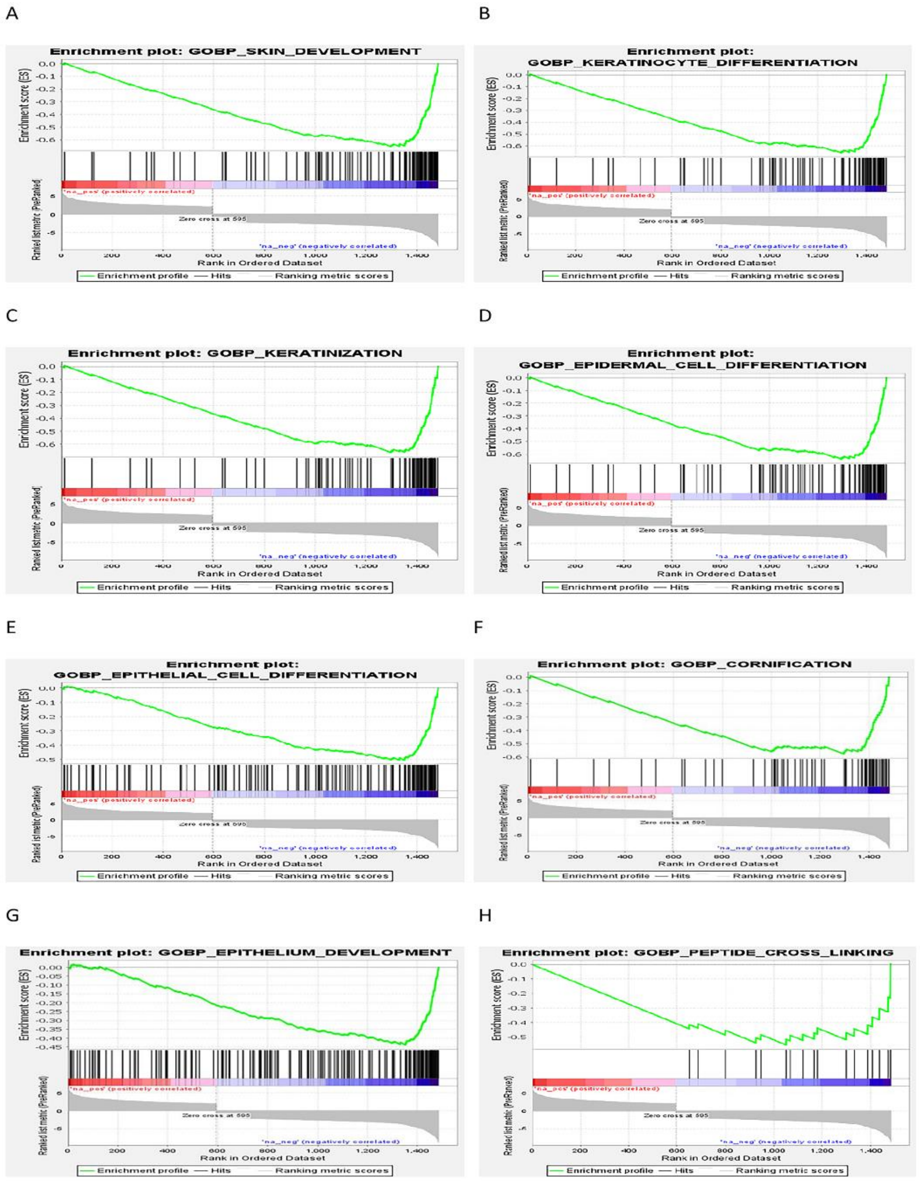


**Supplementary Figure S3: Gene Set Enrichment Analysis (GSEA) considering DEcGs representing significantly depleted GO Biological processes in HPV16-positive CaCx patients.** Graphical representation of the enrichment plots of the depleted GO processes: **(A)**

Skin development **(B)** Keratinocyte Differentiation **(C)** Keratinization **(D)** Epidermal cell differentiation **(E)** Epidermis development **(F)** Epithelial cell differentiation **(G)** Cornification **(H)** Epithelial development and **(I)** Peptide crosslinking. The pathways were considered significant for FDR corrected $p < 0.05$.

**Supplementary Figure S4**

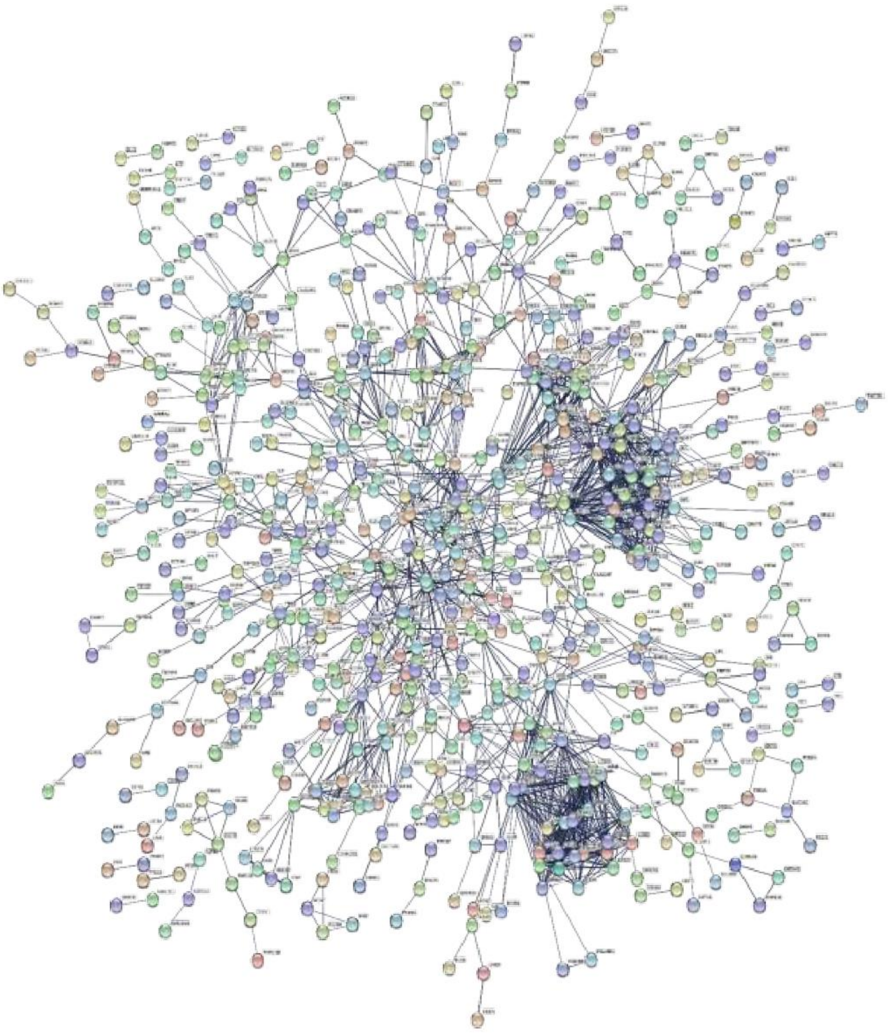

**Supplementary Figure S4: Protein-Protein Interaction (PPI) network.** PPI network (interaction score > 0.7, high confidence) was constructed with the DEcGs identified among the HPV16 positive CaCx patients as compared to normal individuals considered in this study, using STRING database, and visualised through Cytoscape.

**Supplementary Table S1: Details of the sequencing quality and coverage for all sequenced samples (44 HPV16 positive CaCx patients and 34 HPV-negative normal individuals)**

| Sample ID | Annotation | Total number of sequenced reads | Total number of uniquely mapped reads (GRCh37) | RNA Integrity Number (RIN) | Ratio of exon-mapped reads to total uniquely mapped reads (Expression Profile Efficiency) (GRCh37, Ensemble_v87) | Total number of detected transcripts [genes] with reads ≥ 1 [total number of genes (ensembl.v87) = 55638] |
|---|---|---|---|---|---|---|
| 106 | Patient | 80912360 | 70338538 | 8 | 0.793203521 | 35901 |
| 107 | Patient | 80206786 | 40821876 | 7.1 | 0.547513128 | 34937 |
| 108 | Patient | 65224463 | 57259111 | 8 | 0.810313646 | 32059 |
| 109 | Patient | 65287385 | 58200441 | 7.3 | 0.619372025 | 41567 |
| 112 | Patient | 63213895 | 55851994 | 8.4 | 0.788233308 | 35398 |
| 114 | Patient | 54865942 | 48538702 | 8.2 | 0.793219955 | 32206 |
| 115 | Patient | 49491526 | 44100890 | 8.4 | 0.811881574 | 32703 |
| 117 | Patient | 164666734 | 148654476 | 7.5 | 0.782062001 | 35266 |
| 118 | Patient | 197199661 | 177503079 | 7 | 0.850935065 | 34247 |
| 121 | Patient | 145021711 | 131267799 | 7.4 | 0.804786328 | 35511 |
| 131 | Patient | 174661264 | 159544695 | 6.5 | 0.614608201 | 40465 |
| 132 | Patient | 64963905 | 57193150 | 8.3 | 0.845369 | 30547 |
| 135 | Patient | 177125851 | 160360533 | 8.1 | 0.810837016 | 38903 |
| 136 | Patient | 67526479 | 58418582 | 8.4 | 0.734015249 | 41307 |
| 140 | Patient | 61494207 | 54032575 | 7 | 0.578111667 | 42177 |
| 142 | Patient | 150981348 | 135115291 | 8.4 | 0.699890577 | 35732 |
| 143 | Patient | 54021319 | 47441633 | 8.1 | 0.727481514 | 31876 |
| 144 | Patient | 60865383 | 53554893 | 8.1 | 0.83915128 | 31749 |
| 147 | Patient | 128067988 | 116647473 | 6.9 | 0.702048599 | 43431 |
| 148 | Patient | 178370875 | 159750526 | 7.4 | 0.737924738 | 39998 |
| 17-373 | Patient | 83604824 | 55638445 | 8.3 | 0.75456744 | 29387 |
| 17-387 | Patient | 73592808 | 45434265 | 8 | 0.751651534 | 29177 |
| 17-397 | Patient | 84573947 | 69727734 | 7.4 | 0.785158471 | 32113 |
| 17-417 | Patient | 72037683 | 46277527 | 7 | 0.783032723 | 28862 |
| 17-491 | Patient | 47852894 | 17100794 | 6.7 | 0.416782168 | 24941 |
| 17-499 | Patient | 65100117 | 33949555 | 7.5 | 0.632697925 | 29231 |
| 17-580 | Patient | 84469811 | 67456478 | 8 | 0.752516059 | 31992 |
| 17-590 | Patient | 93777517 | 78957240 | 8.5 | 0.833736362 | 31181 |
| 17-599 | Patient | 96189519 | 42146322 | 8.3 | 0.573260058 | 28657 |

| | | | | | | |
|---|---|---|---|---|---|---|
| 17-633 | Patient | 49929082 | 29165923 | 8.6 | 0.731296246 | 26597 |
| 17-641 | Patient | 47761144 | 24813303 | 7.8 | 0.661969186 | 25143 |
| 17-652 | Patient | 88187381 | 46484328 | 7.6 | 0.613460519 | 27983 |
| 17-736 | Patient | 49153118 | 17884269 | 8.1 | 0.446879489 | 23676 |
| 17-782 | Patient | 83153927 | 57124294 | 8.7 | 0.817590726 | 29136 |
| 17-857 | Patient | 57360379 | 46789723 | 7.5 | 0.770248437 | 31464 |
| 18-1062 | Patient | 103973584 | 53345812 | 8.5 | 0.688232471 | 28552 |
| 18-335 | Patient | 58305378 | 36864999 | 8.4 | 0.748615455 | 28362 |
| 18-373 | Patient | 121202000 | 97402532 | 7.4 | 0.741195044 | 33773 |
| 18-636 | Patient | 82551531 | 56350662 | 8.4 | 0.80203977 | 29032 |
| 18-815 | Patient | 104862467 | 60473302 | 7.8 | 0.666334658 | 31115 |
| 96 | Patient | 71235281 | 59232908 | 8.3 | 0.77517901 | 38202 |
| 97 | Patient | 87177957 | 71302363 | 8.1 | 0.762419108 | 33407 |
| 99 | Patient | 56999332 | 49407295 | 7 | 0.575368921 | 43068 |
| C108 | Patient | 58697777 | 47654138 | 6.6 | 0.788127675 | 34000 |
| C104 | Normal | 57234507 | 50835625 | 7 | 0.672165475 | 41254 |
| C116 | Normal | 52539680 | 46029522 | 6.9 | 0.5162527 | 46890 |
| C139 | Normal | 58706163 | 50860273 | 7.3 | 0.630034467 | 45279 |
| C144 | Normal | 61002456 | 52887028 | 7.4 | 0.49545189 | 48670 |
| C153 | Normal | 77194541 | 70910540 | 6.8 | 0.600652061 | 42406 |
| C162 | Normal | 50302216 | 44486299 | 6.8 | 0.578220836 | 43448 |
| C169 | Normal | 55493859 | 48109350 | 7.3 | 0.54218072 | 47853 |
| C173 | Normal | 51564715 | 45801472 | 7.1 | 0.55459866 | 45812 |
| C184 | Normal | 53353731 | 46418688 | 6.1 | 0.357629884 | 48726 |
| C203 | Normal | 60372072 | 53857692 | 6 | 0.558519663 | 41298 |
| C215 | Normal | 57982053 | 50515864 | 6.9 | 0.744556205 | 39712 |
| C216 | Normal | 54358767 | 46466799 | 6.1 | 0.653477357 | 34555 |
| C218 | Normal | 56848203 | 49478924 | 6.3 | 0.616283107 | 40067 |
| C236 | Normal | 76036933 | 66409598 | 5.8 | 0.602501343 | 47007 |
| C237 | Normal | 77911585 | 62524438 | 6 | 0.653319059 | 35996 |
| C239 | Normal | 63329025 | 56413056 | 7.6 | 0.662171484 | 35585 |
| C241 | Normal | 51200580 | 45659041 | 5.8 | 0.613866419 | 42464 |
| C399 | Normal | 91146090 | 35576953 | 8.4 | 0.496135827 | 26844 |
| C401 | Normal | 66304436 | 29047874 | 7.3 | 0.558320137 | 27401 |
| C402 | Normal | 79270927 | 25719050 | 7.2 | 0.378597227 | 25835 |
| C407 | Normal | 85334623 | 32126639 | 7.3 | 0.34492746 | 34453 |
| C409 | Normal | 71321519 | 37909140 | 6.8 | 0.52530767 | 37464 |
| C85 | Normal | 98322944 | 32767586 | 7 | 0.359957398 | 27732 |
| CMCT-107 | Normal | 68022926 | 50423000 | 6.7 | 0.790867818 | 34607 |
| CMCT-121 | Normal | 39607536 | 31672147 | 7.9 | 0.871281192 | 25302 |
| CMCT136 | Normal | 68467490 | 46801455 | 7.5 | 0.741682305 | 37721 |
| CMCT-137 | Normal | 58251639 | 47833407 | 7.4 | 0.868333569 | 31458 |
| CMCT-138 | Normal | 47622185 | 41751280 | 7.3 | 0.741126931 | 41271 |

| | | | | | | |
|---|---|---|---|---|---|---|
| CMCT-140 | Normal | 99321138 | 60490325 | 7.8 | 0.735441164 | 31173 |
| CMCT-141 | Normal | 70699401 | 52649330 | 7.6 | 0.824234667 | 29377 |
| CMCT-142 | Normal | 98008778 | 52266552 | 6.3 | 0.580762454 | 43366 |
| CMCT-181 | Normal | 37698952 | 32292270 | 5.7 | 0.572454646 | 43938 |
| JT-109 | Normal | 69219444 | 58934075 | 6.4 | 0.79163618 | 41073 |
| JT-92 | Normal | 47073207 | 34606004 | 6.8 | 0.8137329 | 28979 |

**N.B**: Patient in annotation column represents the HPV16 positive CaCx patients
Normal represents the histopathologically normal HPV negative healthy individuals

| **Supplementary Table S2: List of primers used for the estimation of RNA expression of the selected DEGs by qRT-PCR** | | |
|---|---|---|
| **Target Name** | **primer** | **Primer sequence (5'-3')** |
| FOXD3 | Forward Primer | GAC GAC GGG CTG GAA GAG AA |
| | Reverse Primer | GCC TCC TTG GGC AAT GTC A |
| FOXD3-AS1 | Forward Primer | GGT GGA GGA GGC GAG GAT G |
| | Reverse Primer | AGC GGA CAG ACA GGG ATT GG |
| GAPDH | Forward Primer | CAG CCT CAA GAT CAT CAG CA |
| | Reverse Primer | TGT GGT CAT GAG TCC TTC CA |

**<u>Supplementary Table S3</u>: Details of all differentially expressed genes (DEcGs and DElncGs) in HPV16-positive CaCx patients as compared to HPV-negative normal individuals portraying expression difference of fold change (FC) cut-off |log2(FC)| ≥ 2 with False Discovery Rate (FDR) corrected p < 0.05**

| Gene Name | log2(FC) | Regulation Status | FDR corrected p value | gene type (GENCODE annotation) |
|---|---|---|---|---|
| YBX2 | 3.282 | Upregulation | 6.8608E-43 | protein coding |
| CCL18 | 2.584 | Upregulation | 3.8457E-09 | protein coding |
| TMEM132A | 2.222 | Upregulation | 1.4173E-31 | protein coding |
| PRSS21 | 3.072 | Upregulation | 4.4054E-13 | protein coding |
| PROM1 | 2.700 | Upregulation | 4.0105E-05 | protein coding |
| NOS2 | 2.195 | Upregulation | 8.012E-06 | protein coding |
| PAX6 | 2.754 | Upregulation | 1.9537E-11 | protein coding |
| FMO3 | 4.435 | Upregulation | 3.027E-20 | protein coding |
| CELSR3 | 3.579 | Upregulation | 2.227E-60 | protein coding |
| IL32 | 2.566 | Upregulation | 4.7083E-28 | protein coding |
| PAX7 | 3.363 | Upregulation | 1.1917E-08 | protein coding |
| LTF | 3.678 | Upregulation | 1.4799E-08 | protein coding |
| CYP24A1 | 2.487 | Upregulation | 1.2215E-07 | protein coding |
| HMGB3 | 2.349 | Upregulation | 3.3294E-35 | protein coding |
| DNAH5 | 2.529 | Upregulation | 2.0856E-13 | protein coding |
| SOX30 | 3.796 | Upregulation | 1.0232E-22 | protein coding |
| ZIC2 | 5.390 | Upregulation | 6.4558E-49 | protein coding |
| ADRB1 | 2.635 | Upregulation | 1.3173E-18 | protein coding |
| DSG2 | 2.457 | Upregulation | 2.8386E-34 | protein coding |
| CP | 3.458 | Upregulation | 8.2897E-11 | protein coding |
| ROS1 | 2.851 | Upregulation | 8.5991E-10 | protein coding |
| TNFRSF17 | 2.271 | Upregulation | 1.1525E-08 | protein coding |
| COL9A2 | 3.756 | Upregulation | 5.8312E-35 | protein coding |
| TNIP3 | 2.664 | Upregulation | 1.3397E-11 | protein coding |
| LAMC2 | 2.515 | Upregulation | 3.752E-18 | protein coding |
| COL11A1 | 3.623 | Upregulation | 9.9271E-14 | protein coding |
| DMRT3 | 2.212 | Upregulation | 3.8127E-05 | protein coding |
| STAG3 | 4.418 | Upregulation | 1.3472E-36 | protein coding |

| | | | | |
|---|---|---|---|---|
| TFRC | 2.076 | Upregulation | 8.3608E-27 | protein coding |
| MCM2 | 2.921 | Upregulation | 3.5134E-77 | protein coding |
| CACNG4 | 2.429 | Upregulation | 1.2846E-07 | protein coding |
| MOCOS | 2.601 | Upregulation | 1.6227E-23 | protein coding |
| UBE2T | 2.343 | Upregulation | 4.1132E-47 | protein coding |
| SMC1B | 5.094 | Upregulation | 2.8357E-65 | protein coding |
| LAMP3 | 2.126 | Upregulation | 3.3166E-21 | protein coding |
| PTPRH | 3.036 | Upregulation | 3.6001E-23 | protein coding |
| SLC4A4 | 2.191 | Upregulation | 8.193E-06 | protein coding |
| EPYC | 3.125 | Upregulation | 1.0077E-09 | protein coding |
| ORC1 | 2.078 | Upregulation | 5.8693E-31 | protein coding |
| HSD17B2 | 2.424 | Upregulation | 3.4803E-09 | protein coding |
| AURKA | 2.043 | Upregulation | 3.7747E-37 | protein coding |
| ARHGAP4 | 2.194 | Upregulation | 8.1174E-31 | protein coding |
| KIF4A | 2.278 | Upregulation | 9.5945E-39 | protein coding |
| DLL3 | 2.423 | Upregulation | 2.6469E-10 | protein coding |
| SEL1L3 | 2.262 | Upregulation | 6.5661E-18 | protein coding |
| CLSPN | 2.081 | Upregulation | 5.2017E-33 | protein coding |
| CDC6 | 2.141 | Upregulation | 8.8334E-39 | protein coding |
| MYO3A | 2.349 | Upregulation | 4.8248E-12 | protein coding |
| TREM2 | 3.100 | Upregulation | 7.9071E-30 | protein coding |
| TSPAN15 | 2.343 | Upregulation | 6.2264E-41 | protein coding |
| IGFALS | 2.193 | Upregulation | 1.435E-05 | protein coding |
| MISP | 2.995 | Upregulation | 3.7091E-13 | protein coding |
| MMP11 | 3.835 | Upregulation | 1.2684E-32 | protein coding |
| DERL3 | 3.105 | Upregulation | 9.4864E-30 | protein coding |
| OSM | 2.291 | Upregulation | 3.9459E-11 | protein coding |
| SLC16A8 | 2.592 | Upregulation | 1.1194E-23 | protein coding |
| CENPM | 2.015 | Upregulation | 1.3317E-25 | protein coding |
| SEPTIN3 | 2.942 | Upregulation | 8.7022E-37 | protein coding |
| NEFH | 3.372 | Upregulation | 3.2511E-16 | protein coding |
| APOL1 | 3.139 | Upregulation | 6.978E-29 | protein coding |
| SIX4 | 3.456 | Upregulation | 6.0986E-46 | protein coding |
| MMP9 | 2.847 | Upregulation | 6.4521E-15 | protein coding |
| GINS1 | 2.007 | Upregulation | 3.2748E-39 | protein coding |
| MYBL2 | 2.826 | Upregulation | 3.9618E-48 | protein coding |

| | | | | |
|---|---|---|---|---|
| SALL4 | 3.408 | Upregulation | 2.032E-28 | protein coding |
| E2F1 | 2.525 | Upregulation | 3.6585E-45 | protein coding |
| WFDC2 | 3.077 | Upregulation | 5.6752E-09 | protein coding |
| VGLL1 | 2.025 | Upregulation | 2.9159E-05 | protein coding |
| TAF7L | 3.770 | Upregulation | 2.1856E-31 | protein coding |
| MSLN | 4.268 | Upregulation | 1.2744E-20 | protein coding |
| TMC5 | 2.294 | Upregulation | 8.4594E-09 | protein coding |
| OIP5 | 2.258 | Upregulation | 7.2918E-34 | protein coding |
| RNASEH2A | 2.130 | Upregulation | 3.9772E-51 | protein coding |
| AMH | 2.718 | Upregulation | 4.0587E-14 | protein coding |
| IL4I1 | 2.750 | Upregulation | 7.2288E-33 | protein coding |
| ASF1B | 2.728 | Upregulation | 5.5877E-51 | protein coding |
| CNTD2 | 2.427 | Upregulation | 2.3344E-11 | protein coding |
| ZFR2 | 4.385 | Upregulation | 2.8201E-51 | protein coding |
| CD79A | 2.143 | Upregulation | 1.0884E-09 | protein coding |
| ICAM4 | 2.758 | Upregulation | 5.7614E-17 | protein coding |
| GRIN2D | 3.932 | Upregulation | 2.0333E-33 | protein coding |
| SLC5A5 | 2.789 | Upregulation | 2.6487E-07 | protein coding |
| MEST | 2.174 | Upregulation | 7.5852E-21 | protein coding |
| AGR2 | 3.838 | Upregulation | 1.3299E-12 | protein coding |
| LHX2 | 4.303 | Upregulation | 1.1178E-29 | protein coding |
| ELAVL2 | 2.708 | Upregulation | 1.1032E-12 | protein coding |
| CA9 | 5.028 | Upregulation | 7.2234E-29 | protein coding |
| CSF3 | 2.171 | Upregulation | 0.00012256 | protein coding |
| HOXB6 | 2.434 | Upregulation | 4.6877E-12 | protein coding |
| MDK | 2.423 | Upregulation | 2.9866E-32 | protein coding |
| KRT18 | 2.086 | Upregulation | 6.9992E-24 | protein coding |
| PPM1H | 2.087 | Upregulation | 7.5923E-11 | protein coding |
| RAD51AP1 | 2.247 | Upregulation | 4.1434E-38 | protein coding |
| OAS2 | 2.007 | Upregulation | 6.5476E-17 | protein coding |
| PRDM13 | 3.420 | Upregulation | 1.029E-10 | protein coding |
| GMNN | 2.009 | Upregulation | 2.1033E-32 | protein coding |
| KIAA1244 | 2.084 | Upregulation | 5.4716E-15 | protein coding |
| STC2 | 2.252 | Upregulation | 2.0601E-11 | protein coding |
| RBP1 | 2.416 | Upregulation | 3.2208E-17 | protein coding |
| LRRC31 | 2.295 | Upregulation | 0.0003553 | protein coding |

| | | | | |
|---|---|---|---|---|
| ECT2 | 2.351 | Upregulation | 4.9596E-67 | protein coding |
| PODXL2 | 3.445 | Upregulation | 1.1487E-67 | protein coding |
| CSPG5 | 2.115 | Upregulation | 4.5518E-16 | protein coding |
| CCL20 | 4.305 | Upregulation | 4.0539E-19 | protein coding |
| CENPA | 2.065 | Upregulation | 3.3218E-32 | protein coding |
| TLX2 | 3.558 | Upregulation | 1.9082E-31 | protein coding |
| OTX1 | 3.120 | Upregulation | 1.8167E-34 | protein coding |
| VAX2 | 2.135 | Upregulation | 5.2084E-14 | protein coding |
| KIAA1324 | 3.303 | Upregulation | 1.0596E-09 | protein coding |
| AGMAT | 2.326 | Upregulation | 9.0033E-20 | protein coding |
| BMP8B | 2.348 | Upregulation | 1.1024E-33 | protein coding |
| CDC20 | 2.037 | Upregulation | 3.9952E-23 | protein coding |
| ARTN | 2.053 | Upregulation | 1.0932E-11 | protein coding |
| TSPAN1 | 3.238 | Upregulation | 1.7992E-20 | protein coding |
| NEK2 | 2.543 | Upregulation | 5.428E-44 | protein coding |
| CHRNB4 | 2.131 | Upregulation | 8.1656E-12 | protein coding |
| ZNF541 | 4.478 | Upregulation | 3.7446E-55 | protein coding |
| SLC8A2 | 2.333 | Upregulation | 2.3789E-09 | protein coding |
| KIF14 | 2.073 | Upregulation | 1.8083E-29 | protein coding |
| SPP1 | 5.707 | Upregulation | 8.4184E-49 | protein coding |
| RARRES1 | 3.875 | Upregulation | 3.8886E-16 | protein coding |
| ONECUT2 | 3.179 | Upregulation | 1.3631E-16 | protein coding |
| EPCAM | 3.469 | Upregulation | 4.6649E-38 | protein coding |
| ELOVL3 | 2.980 | Upregulation | 5.5853E-16 | protein coding |
| IFIT3 | 2.135 | Upregulation | 1.7108E-16 | protein coding |
| NKX2-3 | 3.250 | Upregulation | 1.0719E-09 | protein coding |
| RDH10 | 2.655 | Upregulation | 3.8038E-23 | protein coding |
| PLAU | 2.012 | Upregulation | 2.2113E-20 | protein coding |
| CDKN2C | 2.075 | Upregulation | 2.1049E-33 | protein coding |
| HOXC13 | 2.237 | Upregulation | 1.3036E-08 | protein coding |
| HOXC11 | 3.357 | Upregulation | 1.1337E-15 | protein coding |
| STIL | 2.251 | Upregulation | 4.7331E-50 | protein coding |
| COL10A1 | 5.140 | Upregulation | 1.5327E-36 | protein coding |
| G0S2 | 2.407 | Upregulation | 1.1986E-14 | protein coding |
| CTCFL | 4.910 | Upregulation | 1.4819E-17 | protein coding |
| KCNS1 | 4.264 | Upregulation | 4.9759E-54 | protein coding |

| | | | | |
|---|---|---|---|---|
| ZBP1 | 2.093 | Upregulation | 7.3368E-12 | protein coding |
| IL17C | 2.035 | Upregulation | 1.4455E-05 | protein coding |
| HIST1H2BJ | 2.051 | Upregulation | 1.913E-30 | protein coding |
| OR2B6 | 3.495 | Upregulation | 4.7988E-18 | protein coding |
| SPDEF | 3.292 | Upregulation | 4.5303E-09 | protein coding |
| TCP11 | 3.533 | Upregulation | 2.1451E-35 | protein coding |
| MYRF | 2.995 | Upregulation | 4.0637E-12 | protein coding |
| SRMS | 2.050 | Upregulation | 2.8084E-08 | protein coding |
| CD70 | 2.688 | Upregulation | 6.0408E-14 | protein coding |
| NKX2-4 | 3.255 | Upregulation | 3.6748E-05 | protein coding |
| TCF15 | 3.281 | Upregulation | 4.7346E-17 | protein coding |
| BPIFB1 | 2.485 | Upregulation | 0.00050858 | protein coding |
| IFI6 | 2.844 | Upregulation | 6.0315E-22 | protein coding |
| SIX1 | 3.740 | Upregulation | 4.1566E-25 | protein coding |
| TSPAN8 | 2.103 | Upregulation | 4.3298E-05 | protein coding |
| AUNIP | 2.216 | Upregulation | 1.0899E-39 | protein coding |
| SYNGR3 | 4.645 | Upregulation | 1.6316E-96 | protein coding |
| PKMYT1 | 2.149 | Upregulation | 1.1997E-26 | protein coding |
| GNGT1 | 3.634 | Upregulation | 6.845E-13 | protein coding |
| RIBC2 | 2.817 | Upregulation | 3.4002E-43 | protein coding |
| HOXD1 | 2.968 | Upregulation | 4.4345E-26 | protein coding |
| FAM64A | 2.130 | Upregulation | 1.8669E-26 | protein coding |
| FOXJ1 | 4.290 | Upregulation | 1.5489E-18 | protein coding |
| APOC1 | 3.860 | Upregulation | 2.364E-40 | protein coding |
| BST2 | 2.029 | Upregulation | 2.5334E-25 | protein coding |
| KLHDC7B | 5.551 | Upregulation | 9.8917E-48 | protein coding |
| GDF15 | 4.659 | Upregulation | 2.6935E-40 | protein coding |
| TNNI2 | 2.061 | Upregulation | 2.0889E-06 | protein coding |
| MNX1 | 3.234 | Upregulation | 2.4059E-16 | protein coding |
| ASS1 | 2.227 | Upregulation | 1.6362E-16 | protein coding |
| SMPDL3B | 3.265 | Upregulation | 2.0097E-38 | protein coding |
| SLC6A8 | 2.054 | Upregulation | 2.6114E-16 | protein coding |
| PNCK | 3.502 | Upregulation | 1.1314E-19 | protein coding |
| DUSP9 | 3.216 | Upregulation | 6.9299E-18 | protein coding |
| C1QL1 | 3.193 | Upregulation | 1.1633E-17 | protein coding |
| GINS2 | 2.345 | Upregulation | 1.0769E-45 | protein coding |

| | | | | |
|---|---|---|---|---|
| IDO1 | 5.533 | Upregulation | 3.7285E-53 | protein coding |
| BARX1 | 4.506 | Upregulation | 5.9051E-11 | protein coding |
| RHPN2 | 2.417 | Upregulation | 3.7273E-20 | protein coding |
| NUP210 | 2.299 | Upregulation | 1.2973E-53 | protein coding |
| ACY3 | 2.063 | Upregulation | 1.9631E-06 | protein coding |
| RBM38 | 2.015 | Upregulation | 1.7678E-41 | protein coding |
| VSTM2L | 2.453 | Upregulation | 3.3087E-07 | protein coding |
| CHIT1 | 2.328 | Upregulation | 1.3379E-10 | protein coding |
| EPHB2 | 2.256 | Upregulation | 1.5896E-19 | protein coding |
| NTS | 5.782 | Upregulation | 2.5693E-21 | protein coding |
| GSC | 2.137 | Upregulation | 2.3701E-08 | protein coding |
| ADAMDEC1 | 2.523 | Upregulation | 3.5145E-11 | protein coding |
| VTCN1 | 3.735 | Upregulation | 6.4465E-16 | protein coding |
| RSAD2 | 2.789 | Upregulation | 6.6734E-20 | protein coding |
| CMPK2 | 2.600 | Upregulation | 2.5439E-24 | protein coding |
| ERN2 | 2.856 | Upregulation | 4.0203E-09 | protein coding |
| GRP | 2.483 | Upregulation | 7.0746E-08 | protein coding |
| IL2RA | 2.049 | Upregulation | 7.4477E-17 | protein coding |
| CLDN10 | 2.912 | Upregulation | 2.456E-05 | protein coding |
| GOLM1 | 2.003 | Upregulation | 1.8163E-15 | protein coding |
| SDS | 2.753 | Upregulation | 5.4035E-17 | protein coding |
| RNFT2 | 2.025 | Upregulation | 1.734E-23 | protein coding |
| KRT7 | 5.238 | Upregulation | 6.2632E-41 | protein coding |
| IGF2BP3 | 2.268 | Upregulation | 3.6287E-09 | protein coding |
| ALPK3 | 2.027 | Upregulation | 4.8592E-11 | protein coding |
| IGFBPL1 | 4.063 | Upregulation | 2.1778E-22 | protein coding |
| HIST1H2AB | 2.135 | Upregulation | 4.2438E-20 | protein coding |
| TMPRSS4 | 2.042 | Upregulation | 6.5967E-17 | protein coding |
| MMP7 | 2.221 | Upregulation | 4.2651E-05 | protein coding |
| MMP13 | 5.571 | Upregulation | 3.5085E-26 | protein coding |
| NUSAP1 | 2.127 | Upregulation | 8.7891E-34 | protein coding |
| ITPKA | 2.143 | Upregulation | 8.8225E-16 | protein coding |
| GCHFR | 2.393 | Upregulation | 8.135E-36 | protein coding |
| IFI44L | 2.279 | Upregulation | 3.7256E-12 | protein coding |
| TRIM54 | 2.167 | Upregulation | 5.2025E-05 | protein coding |
| MSMB | 3.192 | Upregulation | 4.3433E-11 | protein coding |

| | | | | |
|---|---|---|---|---|
| CXCL9 | 2.734 | Upregulation | 5.2721E-15 | protein coding |
| LGR5 | 2.390 | Upregulation | 2.9509E-06 | protein coding |
| ZIC5 | 5.839 | Upregulation | 6.9231E-50 | protein coding |
| WARS | 2.221 | Upregulation | 4.3144E-16 | protein coding |
| MARVELD3 | 2.017 | Upregulation | 2.514E-48 | protein coding |
| CHST4 | 2.559 | Upregulation | 1.0182E-06 | protein coding |
| SLC16A3 | 2.380 | Upregulation | 2.7499E-29 | protein coding |
| ATHL1 | 2.276 | Upregulation | 8.8468E-16 | protein coding |
| PADI3 | 3.586 | Upregulation | 1.0027E-15 | protein coding |
| FHAD1 | 2.024 | Upregulation | 1.7667E-11 | protein coding |
| DMRTA2 | 6.225 | Upregulation | 1.0858E-54 | protein coding |
| KIF2C | 2.433 | Upregulation | 3.0562E-47 | protein coding |
| NUF2 | 2.111 | Upregulation | 4.6649E-38 | protein coding |
| DTL | 2.440 | Upregulation | 3.2939E-45 | protein coding |
| HES6 | 3.674 | Upregulation | 7.5382E-35 | protein coding |
| ALDH1L1 | 2.006 | Upregulation | 1.1689E-06 | protein coding |
| HAPLN1 | 4.537 | Upregulation | 7.0472E-33 | protein coding |
| PLA2G7 | 2.459 | Upregulation | 7.4201E-23 | protein coding |
| PNLDC1 | 3.245 | Upregulation | 3.6007E-09 | protein coding |
| RAB19 | 2.123 | Upregulation | 1.7579E-17 | protein coding |
| GINS4 | 2.076 | Upregulation | 7.9848E-38 | protein coding |
| CDKN2A | 5.426 | Upregulation | 4.734E-124 | protein coding |
| SCGB1A1 | 3.677 | Upregulation | 7.3172E-11 | protein coding |
| SYT8 | 2.792 | Upregulation | 4.9875E-11 | protein coding |
| KLHL35 | 4.130 | Upregulation | 7.8572E-78 | protein coding |
| MMP3 | 3.917 | Upregulation | 2.0233E-15 | protein coding |
| FCGR1A | 2.032 | Upregulation | 5.3105E-15 | protein coding |
| LYPD1 | 3.000 | Upregulation | 3.3732E-22 | protein coding |
| C11orf53 | 4.376 | Upregulation | 3.2829E-24 | protein coding |
| GPR158 | 2.187 | Upregulation | 1.4336E-05 | protein coding |
| ADAM8 | 2.267 | Upregulation | 4.1972E-25 | protein coding |
| TDO2 | 3.310 | Upregulation | 1.3704E-16 | protein coding |
| POU4F1 | 2.484 | Upregulation | 6.0526E-08 | protein coding |
| CCNO | 2.328 | Upregulation | 5.3509E-18 | protein coding |
| BMP3 | 2.487 | Upregulation | 7.3097E-06 | protein coding |
| PLOD2 | 2.222 | Upregulation | 4.1449E-30 | protein coding |

| | | | | |
|---|---|---|---|---|
| CLGN | 3.676 | Upregulation | 2.6562E-16 | protein coding |
| GAL3ST2 | 2.642 | Upregulation | 4.5569E-10 | protein coding |
| C4orf19 | 2.285 | Upregulation | 1.3121E-10 | protein coding |
| GBP5 | 3.383 | Upregulation | 3.1242E-20 | protein coding |
| SLFN13 | 2.546 | Upregulation | 2.3984E-33 | protein coding |
| SKA1 | 2.017 | Upregulation | 2.4765E-26 | protein coding |
| ALX3 | 3.874 | Upregulation | 1.1692E-09 | protein coding |
| TDRD9 | 3.068 | Upregulation | 1.2896E-09 | protein coding |
| GLYATL2 | 3.843 | Upregulation | 2.1764E-12 | protein coding |
| ATAD2 | 2.022 | Upregulation | 6.5179E-58 | protein coding |
| B3GNT7 | 2.203 | Upregulation | 7.0273E-20 | protein coding |
| SHCBP1L | 3.769 | Upregulation | 8.0304E-15 | protein coding |
| TMEM171 | 2.749 | Upregulation | 1.5903E-25 | protein coding |
| SLC34A2 | 2.518 | Upregulation | 0.00024059 | protein coding |
| TMSB15A | 2.065 | Upregulation | 6.3203E-10 | protein coding |
| CDC25C | 2.189 | Upregulation | 2.5723E-38 | protein coding |
| IGF2BP1 | 2.613 | Upregulation | 1.1627E-06 | protein coding |
| SIM2 | 3.187 | Upregulation | 6.6372E-19 | protein coding |
| M1AP | 2.676 | Upregulation | 1.8605E-24 | protein coding |
| TFF1 | 2.061 | Upregulation | 0.0015358 | protein coding |
| TMPRSS3 | 3.308 | Upregulation | 1.678E-14 | protein coding |
| CBS | 2.300 | Upregulation | 9.5748E-17 | protein coding |
| COX6B2 | 2.245 | Upregulation | 5.8386E-11 | protein coding |
| LY6K | 2.300 | Upregulation | 4.4118E-21 | protein coding |
| RECQL4 | 2.306 | Upregulation | 3.2751E-33 | protein coding |
| GRIN2C | 2.449 | Upregulation | 2.7073E-14 | protein coding |
| SYCE2 | 3.949 | Upregulation | 9.9374E-44 | protein coding |
| ALOX15 | 2.584 | Upregulation | 6.0112E-10 | protein coding |
| MEIOB | 3.696 | Upregulation | 1.0594E-13 | protein coding |
| C16orf59 | 2.539 | Upregulation | 2.5107E-44 | protein coding |
| PAQR4 | 2.492 | Upregulation | 3.0842E-43 | protein coding |
| ASRGL1 | 2.933 | Upregulation | 8.2739E-12 | protein coding |
| HENMT1 | 2.985 | Upregulation | 2.1956E-97 | protein coding |
| FCGR3B | 2.198 | Upregulation | 1.3756E-08 | protein coding |
| TDRD5 | 2.679 | Upregulation | 1.3969E-08 | protein coding |
| IL24 | 3.190 | Upregulation | 3.5192E-13 | protein coding |

| | | | | |
|---|---|---|---|---|
| PIGR | 3.115 | Upregulation | 9.4254E-06 | protein coding |
| TRIM17 | 2.247 | Upregulation | 2.578E-10 | protein coding |
| CAPN13 | 2.016 | Upregulation | 0.00011779 | protein coding |
| TDRD10 | 2.126 | Upregulation | 1.7681E-08 | protein coding |
| ALPPL2 | 3.891 | Upregulation | 5.9038E-07 | protein coding |
| AIM2 | 3.731 | Upregulation | 6.6434E-22 | protein coding |
| CXCL3 | 2.813 | Upregulation | 1.3039E-13 | protein coding |
| CXCL5 | 3.606 | Upregulation | 2.8879E-11 | protein coding |
| PF4 | 2.048 | Upregulation | 6.7292E-05 | protein coding |
| CXCL1 | 2.442 | Upregulation | 6.2826E-08 | protein coding |
| CAMK2N2 | 4.101 | Upregulation | 1.0663E-26 | protein coding |
| RFC4 | 2.499 | Upregulation | 3.6901E-73 | protein coding |
| RPL39L | 2.580 | Upregulation | 3.5281E-42 | protein coding |
| SLC51A | 3.377 | Upregulation | 1.7994E-26 | protein coding |
| MFI2 | 2.467 | Upregulation | 3.4787E-31 | protein coding |
| S100P | 2.167 | Upregulation | 3.8063E-06 | protein coding |
| ESM1 | 3.673 | Upregulation | 8.4696E-33 | protein coding |
| GJB7 | 2.735 | Upregulation | 3.7909E-07 | protein coding |
| TLX3 | 5.244 | Upregulation | 1.5355E-23 | protein coding |
| HNF4G | 2.342 | Upregulation | 1.1195E-07 | protein coding |
| GBX1 | 2.795 | Upregulation | 4.3042E-06 | protein coding |
| RNF183 | 2.861 | Upregulation | 1.6443E-15 | protein coding |
| CLDN3 | 5.320 | Upregulation | 5.4477E-22 | protein coding |
| NGB | 3.864 | Upregulation | 9.792E-12 | protein coding |
| E2F7 | 2.143 | Upregulation | 8.9234E-31 | protein coding |
| MOGAT2 | 2.209 | Upregulation | 1.4144E-05 | protein coding |
| WDR72 | 4.374 | Upregulation | 1.2804E-19 | protein coding |
| MMP10 | 2.762 | Upregulation | 4.1497E-07 | protein coding |
| MEI1 | 4.209 | Upregulation | 2.3705E-49 | protein coding |
| B4GALNT2 | 2.172 | Upregulation | 6.3958E-05 | protein coding |
| JSRP1 | 2.340 | Upregulation | 1.1245E-17 | protein coding |
| CDT1 | 2.252 | Upregulation | 4.2257E-35 | protein coding |
| NLRP7 | 4.199 | Upregulation | 8.0238E-16 | protein coding |
| TK1 | 2.315 | Upregulation | 1.4173E-31 | protein coding |
| RAB26 | 2.231 | Upregulation | 1.949E-15 | protein coding |
| BATF2 | 2.679 | Upregulation | 1.437E-21 | protein coding |

| | | | | |
|---|---|---|---|---|
| C11orf85 | 2.514 | Upregulation | 7.6796E-16 | protein coding |
| GNG4 | 2.016 | Upregulation | 2.0753E-07 | protein coding |
| TAP1 | 2.047 | Upregulation | 1.3141E-20 | protein coding |
| GBX2 | 2.117 | Upregulation | 2.5046E-06 | protein coding |
| PCSK9 | 2.132 | Upregulation | 3.0093E-11 | protein coding |
| CXCL10 | 3.178 | Upregulation | 7.3188E-18 | protein coding |
| CXCL11 | 3.549 | Upregulation | 1.2338E-16 | protein coding |
| GPRIN1 | 2.284 | Upregulation | 7.8929E-20 | protein coding |
| IL8 | 2.705 | Upregulation | 5.5545E-11 | protein coding |
| ALCAM | 2.150 | Upregulation | 3.6579E-24 | protein coding |
| CDK1 | 2.011 | Upregulation | 1.2649E-34 | protein coding |
| KRT8 | 3.310 | Upregulation | 4.2512E-41 | protein coding |
| METTL7B | 2.154 | Upregulation | 3.7306E-08 | protein coding |
| KRT86 | 2.192 | Upregulation | 3.4018E-21 | protein coding |
| MZB1 | 2.744 | Upregulation | 4.6727E-13 | protein coding |
| FOXA3 | 3.519 | Upregulation | 2.3802E-12 | protein coding |
| HOXB9 | 2.673 | Upregulation | 1.3762E-11 | protein coding |
| CEL | 5.444 | Upregulation | 1.6027E-39 | protein coding |
| PLAC1 | 3.100 | Upregulation | 9.6487E-12 | protein coding |
| HS6ST2 | 3.840 | Upregulation | 1.0022E-27 | protein coding |
| KCNG3 | 2.253 | Upregulation | 5.6722E-18 | protein coding |
| KRT19 | 2.683 | Upregulation | 1.8117E-22 | protein coding |
| APLN | 2.790 | Upregulation | 7.0881E-14 | protein coding |
| CYP4F11 | 2.065 | Upregulation | 6.9389E-07 | protein coding |
| CCL11 | 2.223 | Upregulation | 4.2142E-07 | protein coding |
| HSPA6 | 2.118 | Upregulation | 2.9229E-15 | protein coding |
| DMRT2 | 4.099 | Upregulation | 1.0684E-19 | protein coding |
| OLR1 | 2.528 | Upregulation | 1.419E-15 | protein coding |
| SAA1 | 3.175 | Upregulation | 1.8369E-10 | protein coding |
| AGR3 | 2.760 | Upregulation | 2.807E-05 | protein coding |
| GPR160 | 2.574 | Upregulation | 8.5052E-21 | protein coding |
| MFSD4 | 2.165 | Upregulation | 2.6039E-10 | protein coding |
| GOLT1A | 3.444 | Upregulation | 3.3188E-18 | protein coding |
| UGT8 | 2.285 | Upregulation | 3.9801E-09 | protein coding |
| PCP2 | 2.069 | Upregulation | 1.084E-19 | protein coding |
| SEZ6L2 | 2.635 | Upregulation | 4.3638E-16 | protein coding |

| | | | | |
|---|---|---|---|---|
| UBE2C | 2.291 | Upregulation | 2.6026E-40 | protein coding |
| SPSB4 | 2.419 | Upregulation | 1.5612E-06 | protein coding |
| MARCKSL1 | 2.020 | Upregulation | 1.3948E-30 | protein coding |
| NRIP3 | 3.082 | Upregulation | 7.525E-45 | protein coding |
| RMI2 | 2.370 | Upregulation | 1.5327E-48 | protein coding |
| ACBD7 | 2.717 | Upregulation | 1.2274E-20 | protein coding |
| PRR15 | 2.973 | Upregulation | 2.525E-09 | protein coding |
| CNTD1 | 2.384 | Upregulation | 1.2325E-08 | protein coding |
| FOXL1 | 2.307 | Upregulation | 6.7424E-11 | protein coding |
| FOXC2 | 2.184 | Upregulation | 1.333E-05 | protein coding |
| EFCAB4A | 2.459 | Upregulation | 1.5142E-17 | protein coding |
| MLF1 | 2.476 | Upregulation | 1.5601E-24 | protein coding |
| RNF212 | 2.780 | Upregulation | 6.7476E-37 | protein coding |
| FAM132B | 3.260 | Upregulation | 2.2684E-14 | protein coding |
| LBX2 | 2.304 | Upregulation | 4.2548E-27 | protein coding |
| APOBEC3B | 2.941 | Upregulation | 5.3686E-27 | protein coding |
| ABCA13 | 3.564 | Upregulation | 2.3426E-23 | protein coding |
| C1orf194 | 2.265 | Upregulation | 0.00340059 | protein coding |
| C17orf104 | 2.813 | Upregulation | 2.2537E-12 | protein coding |
| SHISA2 | 3.036 | Upregulation | 4.5906E-16 | protein coding |
| HOXC9 | 3.642 | Upregulation | 4.0579E-22 | protein coding |
| HOXC10 | 4.333 | Upregulation | 9.3135E-25 | protein coding |
| C12orf36 | 2.340 | Upregulation | 2.1389E-07 | protein coding |
| NQO1 | 2.283 | Upregulation | 2.8596E-14 | protein coding |
| MUC16 | 2.266 | Upregulation | 5.2769E-06 | protein coding |
| HIST3H2A | 2.108 | Upregulation | 3.7949E-17 | protein coding |
| SYNE4 | 2.142 | Upregulation | 9.983E-18 | protein coding |
| C6orf223 | 4.370 | Upregulation | 3.8379E-43 | protein coding |
| FDCSP | 3.111 | Upregulation | 1.7464E-05 | protein coding |
| HIST2H4B | 2.103 | Upregulation | 1.1736E-18 | protein coding |
| NXPH4 | 2.181 | Upregulation | 2.4862E-13 | protein coding |
| VMO1 | 2.436 | Upregulation | 1.1825E-15 | protein coding |
| RIPPLY3 | 3.415 | Upregulation | 2.2118E-26 | protein coding |
| GPR19 | 2.357 | Upregulation | 1.3447E-16 | protein coding |
| C2CD4C | 2.438 | Upregulation | 3.951E-13 | protein coding |
| CHST6 | 3.004 | Upregulation | 2.0401E-12 | protein coding |

| | | | | |
|---|---|---|---|---|
| CCDC60 | 2.095 | Upregulation | 5.7506E-09 | protein coding |
| HIST2H2AA3 | 2.009 | Upregulation | 2.7776E-13 | protein coding |
| ASCL2 | 2.053 | Upregulation | 7.5552E-18 | protein coding |
| IQGAP3 | 2.357 | Upregulation | 1.3472E-36 | protein coding |
| HIST2H4A | 2.204 | Upregulation | 6.6927E-17 | protein coding |
| NPW | 2.891 | Upregulation | 6.2508E-12 | protein coding |
| MAP7D2 | 3.118 | Upregulation | 2.2772E-15 | protein coding |
| NCMAP | 2.578 | Upregulation | 9.1927E-08 | protein coding |
| PTP4A3 | 2.011 | Upregulation | 3.1891E-23 | protein coding |
| LRRC26 | 2.214 | Upregulation | 3.4976E-05 | protein coding |
| USP18 | 2.360 | Upregulation | 3.7211E-29 | protein coding |
| NUPR1L | 2.175 | Upregulation | 3.0899E-08 | protein coding |
| MUC1 | 2.383 | Upregulation | 6.5518E-12 | protein coding |
| IRF7 | 2.034 | Upregulation | 2.1849E-26 | protein coding |
| PRAME | 2.486 | Upregulation | 1.4089E-05 | protein coding |
| C1orf110 | 2.482 | Upregulation | 1.9707E-06 | protein coding |
| LEMD1 | 2.438 | Upregulation | 5.5449E-11 | protein coding |
| ZPBP2 | 3.180 | Upregulation | 1.4644E-07 | protein coding |
| LILRB4 | 2.220 | Upregulation | 5.1566E-18 | protein coding |
| TNFRSF18 | 2.482 | Upregulation | 2.0372E-19 | protein coding |
| C17orf82 | 2.007 | Upregulation | 1.0991E-14 | protein coding |
| CCK | 2.244 | Upregulation | 9.1786E-06 | protein coding |
| FOXD3 | 2.287 | Upregulation | 1.7565E-09 | protein coding |
| HIST1H1T | 2.072 | Upregulation | 5.4908E-10 | protein coding |
| PLEKHG7 | 2.297 | Upregulation | 3.6167E-09 | protein coding |
| ISG15 | 3.408 | Upregulation | 1.5969E-30 | protein coding |
| AMTN | 4.567 | Upregulation | 1.7844E-17 | protein coding |
| KBTBD12 | 4.369 | Upregulation | 3.6746E-35 | protein coding |
| DNAH17 | 2.229 | Upregulation | 1.8791E-07 | protein coding |
| NWD1 | 2.800 | Upregulation | 4.6931E-16 | protein coding |
| HEPACAM2 | 2.376 | Upregulation | 7.2645E-05 | protein coding |
| SBK1 | 2.412 | Upregulation | 2.1541E-21 | protein coding |
| IER5L | 2.022 | Upregulation | 1.901E-21 | protein coding |
| HMX2 | 4.373 | Upregulation | 7.7211E-21 | protein coding |
| FAM26F | 2.180 | Upregulation | 2.4028E-21 | protein coding |
| ENTPD8 | 2.456 | Upregulation | 8.3246E-07 | protein coding |

| | | | | |
|---|---|---|---|---|
| FAM111B | 2.384 | Upregulation | 2.6127E-40 | protein coding |
| ANKRD34B | 2.869 | Upregulation | 1.2825E-14 | protein coding |
| ALG1L | 4.790 | Upregulation | 2.5635E-63 | protein coding |
| SYCP2 | 4.386 | Upregulation | 5.105E-132 | protein coding |
| PLEKHG4 | 2.185 | Upregulation | 3.189E-59 | protein coding |
| HIST1H2BO | 2.020 | Upregulation | 2.204E-25 | protein coding |
| NOXO1 | 2.137 | Upregulation | 2.5607E-10 | protein coding |
| C1orf186 | 2.007 | Upregulation | 2.2105E-06 | protein coding |
| MMP1 | 4.481 | Upregulation | 1.36E-25 | protein coding |
| HSH2D | 2.038 | Upregulation | 2.2042E-13 | protein coding |
| SERPINA1 | 2.102 | Upregulation | 3.7021E-11 | protein coding |
| ZNF695 | 2.291 | Upregulation | 3.6998E-12 | protein coding |
| DMBX1 | 2.772 | Upregulation | 4.912E-10 | protein coding |
| NMB | 2.810 | Upregulation | 4.3325E-31 | protein coding |
| HOXC6 | 3.203 | Upregulation | 5.0405E-21 | protein coding |
| ELOVL2 | 2.190 | Upregulation | 4.1338E-07 | protein coding |
| HIST1H4I | 2.337 | Upregulation | 7.9914E-24 | protein coding |
| HOXC4 | 2.484 | Upregulation | 3.2279E-08 | protein coding |
| KRTAP4-1 | 2.421 | Upregulation | 1.8739E-05 | protein coding |
| C2CD4A | 3.657 | Upregulation | 2.7465E-23 | protein coding |
| EGFL6 | 2.623 | Upregulation | 7.4991E-20 | protein coding |
| SOWAHA | 2.045 | Upregulation | 1.714E-09 | protein coding |
| FCGR3A | 2.578 | Upregulation | 6.778E-24 | protein coding |
| CENPW | 2.431 | Upregulation | 2.0865E-37 | protein coding |
| HIST2H3C | 2.790 | Upregulation | 1.1876E-20 | protein coding |
| HIST2H2AA4 | 2.293 | Upregulation | 8.4176E-19 | protein coding |
| FAM72C | 2.008 | Upregulation | 1.8276E-10 | protein coding |
| HIST2H3A | 2.513 | Upregulation | 2.0404E-16 | protein coding |
| ZYG11A | 2.475 | Upregulation | 6.3197E-32 | protein coding |
| SLC44A4 | 3.071 | Upregulation | 1.1773E-12 | protein coding |
| TRIM15 | 2.518 | Upregulation | 9.5234E-07 | protein coding |
| TRIM31 | 3.737 | Upregulation | 1.3469E-13 | protein coding |
| PTCHD2 | 2.598 | Upregulation | 3.792E-13 | protein coding |
| ZFP57 | 2.092 | Upregulation | 9.6837E-05 | protein coding |
| AC011484.1 | 2.254 | Upregulation | 2.2356E-15 | protein coding |
| CCNI2 | 2.973 | Upregulation | 1.1912E-23 | protein coding |

| | | | | |
|---|---|---|---|---|
| EXOC3L4 | 2.627 | Upregulation | 5.4746E-19 | protein coding |
| C2CD4B | 2.049 | Upregulation | 3.2863E-13 | protein coding |
| IGKV4-1 | 2.893 | Upregulation | 2.9035E-13 | IG_V gene |
| IGKV5-2 | 2.426 | Upregulation | 1.0947E-08 | IG_V gene |
| IGKV6-21 | 3.186 | Upregulation | 1.7492E-09 | IG_V gene |
| IGKV2-40 | 2.710 | Upregulation | 5.6895E-05 | IG_V gene |
| IGKV3D-20 | 2.777 | Upregulation | 1.0662E-15 | IG_V gene |
| IGKV1D-13 | 2.207 | Upregulation | 3.4674E-05 | IG_V gene |
| IGKV3D-11 | 3.007 | Upregulation | 1.6715E-12 | IG_V gene |
| IGLV4-69 | 3.419 | Upregulation | 2.1155E-15 | IG_V gene |
| IGLV8-61 | 4.057 | Upregulation | 5.0669E-18 | IG_V gene |
| IGLV4-60 | 3.514 | Upregulation | 2.428E-12 | IG_V gene |
| IGLV6-57 | 2.347 | Upregulation | 2.2069E-09 | IG_V gene |
| IGLV10-54 | 3.352 | Upregulation | 2.2089E-12 | IG_V gene |
| IGLV1-51 | 2.617 | Upregulation | 8.4159E-14 | IG_V gene |
| IGLV5-48 | 3.300 | Upregulation | 4.1773E-09 | IG_V gene |
| IGLV1-47 | 3.394 | Upregulation | 4.1436E-16 | IG_V gene |
| IGLV7-46 | 2.973 | Upregulation | 1.1308E-09 | IG_V gene |
| IGLV1-44 | 3.249 | Upregulation | 4.8555E-17 | IG_V gene |
| IGLV7-43 | 2.110 | Upregulation | 4.1749E-06 | IG_V gene |
| IGLV1-40 | 3.197 | Upregulation | 1.0275E-18 | IG_V gene |
| IGLV1-36 | 2.147 | Upregulation | 1.9669E-05 | IG_V gene |
| IGLV3-27 | 3.553 | Upregulation | 1.1386E-16 | IG_V gene |
| IGLV3-25 | 3.497 | Upregulation | 1.5634E-17 | IG_V gene |
| IGLV2-23 | 3.248 | Upregulation | 1.6159E-14 | IG_V gene |
| IGLV3-19 | 2.401 | Upregulation | 9.2274E-09 | IG_V gene |
| IGLV2-18 | 2.255 | Upregulation | 1.1682E-06 | IG_V gene |
| IGLV3-16 | 2.658 | Upregulation | 1.6775E-10 | IG_V gene |
| IGLV2-14 | 3.046 | Upregulation | 8.5494E-17 | IG_V gene |
| IGLV2-11 | 3.027 | Upregulation | 4.1974E-14 | IG_V gene |
| IGLV3-9 | 2.356 | Upregulation | 9.2519E-08 | IG_V gene |
| IGLV2-8 | 3.267 | Upregulation | 2.2893E-12 | IG_V gene |
| IGLV3-1 | 3.162 | Upregulation | 3.0689E-13 | IG_V gene |
| IGLC2 | 2.922 | Upregulation | 1.1847E-12 | IG_C gene |
| IGLC3 | 3.004 | Upregulation | 6.9647E-14 | IG_C gene |
| IGHA2 | 2.247 | Upregulation | 3.1465E-07 | IG_C gene |

| | | | | |
|---|---|---|---|---|
| IGHG2 | 2.828 | Upregulation | 3.4801E-11 | IG_C gene |
| IGHA1 | 2.276 | Upregulation | 3.0502E-08 | IG_C gene |
| IGHG1 | 3.093 | Upregulation | 5.652E-16 | IG_C gene |
| IGHG3 | 2.843 | Upregulation | 2.2168E-11 | IG_C gene |
| IGHD | 2.691 | Upregulation | 1.6159E-08 | IG_C gene |
| IGHM | 4.170 | Upregulation | 4.3229E-20 | IG_C gene |
| IGHJ6 | 2.020 | Upregulation | 8.1682E-06 | IG_J gene |
| IGHV6-1 | 3.146 | Upregulation | 5.999E-15 | IG_V gene |
| IGHV1-2 | 3.126 | Upregulation | 8.7053E-15 | IG_V gene |
| IGHV1-3 | 2.237 | Upregulation | 2.3624E-06 | IG_V gene |
| IGHV4-4 | 3.386 | Upregulation | 4.9104E-17 | IG_V gene |
| IGHV2-5 | 3.114 | Upregulation | 1.9407E-11 | IG_V gene |
| IGHV3-7 | 3.121 | Upregulation | 1.4031E-16 | IG_V gene |
| IGHV1-8 | 2.849 | Upregulation | 7.9219E-09 | IG_V gene |
| IGHV3-9 | 3.454 | Upregulation | 1.1931E-12 | IG_V gene |
| IGHV3-11 | 2.882 | Upregulation | 5.6443E-12 | IG_V gene |
| IGHV3-13 | 2.618 | Upregulation | 1.6295E-09 | IG_V gene |
| IGHV3-15 | 2.851 | Upregulation | 1.4391E-12 | IG_V gene |
| IGHV1-18 | 2.783 | Upregulation | 5.2893E-13 | IG_V gene |
| IGHV3-20 | 2.037 | Upregulation | 9.3683E-05 | IG_V gene |
| IGHV3-21 | 3.343 | Upregulation | 4.5943E-15 | IG_V gene |
| IGHV3-23 | 3.551 | Upregulation | 7.4653E-17 | IG_V gene |
| IGHV1-24 | 2.510 | Upregulation | 3.9034E-08 | IG_V gene |
| IGHV2-26 | 2.673 | Upregulation | 3.0569E-09 | IG_V gene |
| IGHV4-28 | 2.428 | Upregulation | 1.709E-07 | IG_V gene |
| IGHV3-30 | 2.765 | Upregulation | 2.0879E-13 | IG_V gene |
| IGHV3-33 | 2.891 | Upregulation | 4.0537E-11 | IG_V gene |
| IGHV4-34 | 3.142 | Upregulation | 1.5521E-15 | IG_V gene |
| IGHV4-39 | 3.922 | Upregulation | 4.0623E-21 | IG_V gene |
| IGHV1-45 | 2.623 | Upregulation | 0.0002073 | IG_V gene |
| IGHV1-46 | 3.178 | Upregulation | 1.0224E-13 | IG_V gene |
| IGHV3-48 | 3.207 | Upregulation | 1.2823E-14 | IG_V gene |
| IGHV3-49 | 3.110 | Upregulation | 3.089E-14 | IG_V gene |
| IGHV5-51 | 2.663 | Upregulation | 1.1705E-11 | IG_V gene |
| IGHV3-53 | 2.237 | Upregulation | 7.2089E-08 | IG_V gene |
| IGHV1-58 | 3.009 | Upregulation | 6.3133E-09 | IG_V gene |

| | | | | |
|---|---|---|---|---|
| IGHV4-61 | 2.684 | Upregulation | 6.9218E-10 | IG_V gene |
| IGHV1-69 | 2.545 | Upregulation | 1.777E-09 | IG_V gene |
| IGHV3-73 | 3.214 | Upregulation | 8.5416E-14 | IG_V gene |
| IGHV7-81 | 2.021 | Upregulation | 0.00086584 | IG_V gene |
| UBD | 2.120 | Upregulation | 4.4093E-08 | protein coding |
| CLDN9 | 2.249 | Upregulation | 5.6943E-08 | protein coding |
| C12orf74 | 2.690 | Upregulation | 1.8098E-12 | protein coding |
| MUC5AC | 2.136 | Upregulation | 0.00193632 | protein coding |
| SP9 | 4.126 | Upregulation | 2.7265E-16 | protein coding |
| HMSD | 2.892 | Upregulation | 6.8681E-17 | protein coding |
| IGLV9-49 | 2.877 | Upregulation | 3.6019E-08 | IG_V gene |
| IGHV3-64 | 2.831 | Upregulation | 5.8382E-10 | IG_V gene |
| IGKV3D-15 | 2.935 | Upregulation | 4.8604E-16 | IG_V gene |
| IGHV4-59 | 3.055 | Upregulation | 3.9597E-14 | IG_V gene |
| IGHV3-74 | 3.008 | Upregulation | 2.6744E-16 | IG_V gene |
| IGHV3-72 | 2.077 | Upregulation | 3.6366E-07 | IG_V gene |
| IGHV4-31 | 2.173 | Upregulation | 3.9729E-06 | IG_V gene |
| IGHV3-43 | 2.731 | Upregulation | 2.3843E-10 | IG_V gene |
| IGKV3OR2-268 | 2.762 | Upregulation | 1.7471E-07 | IG_V gene |
| MCIDAS | 3.490 | Upregulation | 6.4869E-29 | protein coding |
| ANKRD65 | 2.756 | Upregulation | 2.004E-20 | protein coding |
| ARHGEF38 | 2.178 | Upregulation | 9.2008E-05 | protein coding |
| IGKV2D-30 | 2.200 | Upregulation | 1.3633E-06 | IG_V gene |
| IGKV3-20 | 3.249 | Upregulation | 2.3979E-17 | IG_V gene |
| IGKV1D-33 | 2.211 | Upregulation | 1.2996E-07 | IG_V gene |
| IGHJ4 | 2.149 | Upregulation | 2.7795E-06 | IG_J gene |
| IGKV1-17 | 2.752 | Upregulation | 5.0114E-11 | IG_V gene |
| IGKV1-8 | 2.356 | Upregulation | 6.9087E-07 | IG_V gene |
| IGKV1D-12 | 3.235 | Upregulation | 6.0441E-13 | IG_V gene |
| IGKV1-16 | 3.233 | Upregulation | 2.4307E-13 | IG_V gene |
| IGKV1D-16 | 2.404 | Upregulation | 5.554E-09 | IG_V gene |
| IGKV2-24 | 2.672 | Upregulation | 3.1219E-09 | IG_V gene |
| IGKV3-11 | 2.756 | Upregulation | 9.7451E-13 | IG_V gene |
| PDXP | 2.086 | Upregulation | 1.6944E-09 | protein coding |
| IGKV2D-24 | 2.878 | Upregulation | 6.1175E-09 | IG_V gene |
| IGKV1-9 | 2.925 | Upregulation | 1.3238E-11 | IG_V gene |

| | | | | |
|---|---|---|---|---|
| IGKV1-33 | 2.074 | Upregulation | 6.1516E-07 | IG_V gene |
| PEG10 | 2.343 | Upregulation | 1.9598E-06 | protein coding |
| IGKV1-39 | 2.419 | Upregulation | 3.684E-10 | IG_V gene |
| IGHJ5 | 2.279 | Upregulation | 2.2628E-07 | IG_J gene |
| IGKV2D-28 | 2.444 | Upregulation | 5.4757E-09 | IG_V gene |
| IGKV1D-17 | 2.275 | Upregulation | 8.5418E-05 | IG_V gene |
| IGHJ3 | 2.139 | Upregulation | 1.8809E-05 | IG_J gene |
| IGKV2-30 | 2.874 | Upregulation | 3.9982E-11 | IG_V gene |
| IGKV1-12 | 2.242 | Upregulation | 1.1864E-08 | IG_V gene |
| IGKV1-5 | 2.635 | Upregulation | 9.7831E-12 | IG_V gene |
| CFB | 2.790 | Upregulation | 2.2912E-13 | protein coding |
| IGKV2-28 | 2.240 | Upregulation | 9.3106E-09 | IG_V gene |
| IGKV3-15 | 2.722 | Upregulation | 9.6487E-12 | IG_V gene |
| IGKV1-27 | 3.351 | Upregulation | 6.7999E-14 | IG_V gene |
| IGKV2D-40 | 3.742 | Upregulation | 1.8564E-13 | IG_V gene |
| FOXD1 | 2.320 | Upregulation | 1.434E-07 | protein coding |
| IGKV1D-39 | 2.513 | Upregulation | 3.9774E-10 | IG_V gene |
| IGLL5 | 2.070 | Upregulation | 1.35E-07 | protein coding |
| CCDC177 | 3.585 | Upregulation | 3.5386E-18 | protein coding |
| HIST1H3G | 2.028 | Upregulation | 2.7964E-15 | protein coding |
| HIST1H3F | 2.148 | Upregulation | 3.7455E-21 | protein coding |
| CTD-2021H9.3 | 2.043 | Upregulation | 0.00739015 | protein coding |
| IGHV4OR15-8 | 3.586 | Upregulation | 1.1564E-12 | IG_V gene |
| TCF24 | 2.689 | Upregulation | 1.1944E-09 | protein coding |
| SMIM22 | 4.287 | Upregulation | 2.0097E-38 | protein coding |
| CCDC177 | 3.038 | Upregulation | 3.6049E-13 | protein coding |
| AC019171.1 | 4.458 | Upregulation | 1.6188E-40 | protein coding |
| IGHV3OR16-13 | 2.000 | Upregulation | 1.5477E-06 | IG_V gene |
| WNT16 | -3.926 | Downregulation | 5.4337E-20 | protein coding |
| HSPB6 | -3.370 | Downregulation | 1.6511E-23 | protein coding |
| ABCB5 | -2.038 | Downregulation | 7.237E-07 | protein coding |
| ABCC8 | -3.279 | Downregulation | 3.9929E-15 | protein coding |
| CACNA1G | -2.083 | Downregulation | 5.9082E-12 | protein coding |
| GAS7 | -2.231 | Downregulation | 6.8781E-30 | protein coding |
| NOX1 | -2.147 | Downregulation | 9.3544E-08 | protein coding |
| TENM1 | -2.418 | Downregulation | 3.4511E-10 | protein coding |

| | | | | |
|---|---|---|---|---|
| HHATL | -2.609 | Downregulation | 7.9466E-09 | protein coding |
| PRSS3 | -5.281 | Downregulation | 3.4126E-37 | protein coding |
| DCN | -2.107 | Downregulation | 5.1414E-12 | protein coding |
| STMN4 | -2.505 | Downregulation | 7.2864E-11 | protein coding |
| ISL1 | -2.281 | Downregulation | 3.3234E-08 | protein coding |
| IGF1 | -2.672 | Downregulation | 5.6596E-13 | protein coding |
| FHL1 | -2.632 | Downregulation | 9.5153E-14 | protein coding |
| OTC | -2.986 | Downregulation | 7.3259E-08 | protein coding |
| USP2 | -2.738 | Downregulation | 1.3855E-16 | protein coding |
| MYO16 | -2.247 | Downregulation | 2.8996E-12 | protein coding |
| BARX2 | -2.555 | Downregulation | 9.403E-18 | protein coding |
| PTGER3 | -3.505 | Downregulation | 3.0421E-28 | protein coding |
| NNAT | -2.763 | Downregulation | 7.0439E-18 | protein coding |
| CHRDL2 | -2.927 | Downregulation | 9.7266E-11 | protein coding |
| F7 | -2.541 | Downregulation | 1.6353E-17 | protein coding |
| DGAT2 | -2.407 | Downregulation | 5.5816E-22 | protein coding |
| DLX3 | -2.846 | Downregulation | 1.2435E-12 | protein coding |
| WISP2 | -3.258 | Downregulation | 2.3449E-13 | protein coding |
| GLP2R | -2.871 | Downregulation | 5.3251E-14 | protein coding |
| ROPN1 | -2.372 | Downregulation | 4.153E-08 | protein coding |
| SNAP91 | -2.497 | Downregulation | 4.8206E-12 | protein coding |
| COL17A1 | -2.690 | Downregulation | 5.2089E-15 | protein coding |
| CTNNA2 | -2.715 | Downregulation | 1.8612E-09 | protein coding |
| ACSM2B | -3.198 | Downregulation | 1.968E-10 | protein coding |
| NAV3 | -2.653 | Downregulation | 1.1876E-20 | protein coding |
| MAOB | -2.544 | Downregulation | 1.5035E-20 | protein coding |
| TGFBR3 | -2.397 | Downregulation | 9.3547E-17 | protein coding |
| GUCY2C | -3.409 | Downregulation | 1.1432E-13 | protein coding |
| FGF10 | -3.211 | Downregulation | 2.0583E-16 | protein coding |
| FGF22 | -2.682 | Downregulation | 1.8975E-14 | protein coding |
| FRMPD1 | -5.366 | Downregulation | 2.1847E-30 | protein coding |
| CDH19 | -3.240 | Downregulation | 4.874E-13 | protein coding |
| SPP2 | -2.894 | Downregulation | 1.5641E-07 | protein coding |
| RPS6KA6 | -2.382 | Downregulation | 4.3396E-10 | protein coding |
| TRHDE | -3.188 | Downregulation | 2.2539E-16 | protein coding |
| RGS11 | -2.268 | Downregulation | 7.6407E-13 | protein coding |

| | | | | |
|---|---|---|---|---|
| CTTNBP2 | -2.256 | Downregulation | 8.2672E-15 | protein coding |
| ACTL6B | -3.224 | Downregulation | 2.535E-10 | protein coding |
| ACTN2 | -2.035 | Downregulation | 9.4847E-08 | protein coding |
| NAALAD2 | -2.012 | Downregulation | 1.3284E-07 | protein coding |
| ADCY2 | -2.074 | Downregulation | 4.3698E-07 | protein coding |
| RBFOX1 | -3.020 | Downregulation | 3.9263E-18 | protein coding |
| RUNX1T1 | -2.030 | Downregulation | 7.5058E-14 | protein coding |
| DUSP13 | -2.467 | Downregulation | 8.0293E-23 | protein coding |
| EPHA6 | -2.522 | Downregulation | 1.1204E-09 | protein coding |
| AFP | -2.556 | Downregulation | 1.8808E-09 | protein coding |
| PCDHGA2 | -2.155 | Downregulation | 1.285E-17 | protein coding |
| PGR | -2.104 | Downregulation | 1.2284E-05 | protein coding |
| TRPM3 | -2.494 | Downregulation | 8.4073E-09 | protein coding |
| HAL | -4.788 | Downregulation | 8.0626E-29 | protein coding |
| TMPRSS11E | -2.853 | Downregulation | 7.6536E-11 | protein coding |
| PHACTR3 | -2.357 | Downregulation | 7.5018E-10 | protein coding |
| SULT2B1 | -2.793 | Downregulation | 5.6618E-26 | protein coding |
| GP6 | -2.038 | Downregulation | 4.4182E-14 | protein coding |
| SLC15A1 | -2.207 | Downregulation | 1.0347E-07 | protein coding |
| ARHGAP28 | -2.563 | Downregulation | 4.3115E-17 | protein coding |
| RPH3A | -2.556 | Downregulation | 9.9941E-10 | protein coding |
| TBX5 | -2.492 | Downregulation | 2.8533E-09 | protein coding |
| NOS1 | -2.986 | Downregulation | 5.5007E-10 | protein coding |
| FETUB | -5.261 | Downregulation | 1.3893E-31 | protein coding |
| LAMB4 | -3.339 | Downregulation | 2.0824E-25 | protein coding |
| ZFHX4 | -3.028 | Downregulation | 9.1521E-21 | protein coding |
| ESR1 | -2.416 | Downregulation | 3.1686E-13 | protein coding |
| CMA1 | -2.152 | Downregulation | 3.4084E-06 | protein coding |
| TGM1 | -2.750 | Downregulation | 4.6109E-14 | protein coding |
| SEMA6A | -2.112 | Downregulation | 3.0419E-11 | protein coding |
| FMO2 | -3.022 | Downregulation | 1.1732E-08 | protein coding |
| NXPE1 | -3.133 | Downregulation | 3.4438E-13 | protein coding |
| CRISP3 | -5.244 | Downregulation | 1.868E-22 | protein coding |
| PGC | -2.387 | Downregulation | 0.00140488 | protein coding |
| DSP | -2.267 | Downregulation | 2.775E-25 | protein coding |
| PLA2G3 | -2.813 | Downregulation | 1.2159E-13 | protein coding |

| | | | | |
|---|---|---|---|---|
| SOX10 | -2.899 | Downregulation | 5.4061E-11 | protein coding |
| ACR | -2.886 | Downregulation | 2.9802E-20 | protein coding |
| MLC1 | -2.319 | Downregulation | 1.275E-11 | protein coding |
| C14orf166B | -2.540 | Downregulation | 8.8857E-07 | protein coding |
| CPNE6 | -2.224 | Downregulation | 7.0264E-08 | protein coding |
| RIMS4 | -3.529 | Downregulation | 9.5056E-13 | protein coding |
| SEL1L2 | -2.883 | Downregulation | 4.5733E-13 | protein coding |
| EPPIN | -2.349 | Downregulation | 1.4624E-05 | protein coding |
| CHRDL1 | -2.973 | Downregulation | 7.8146E-08 | protein coding |
| PAGE4 | -2.478 | Downregulation | 2.8062E-07 | protein coding |
| ZC3H12B | -2.217 | Downregulation | 8.3305E-15 | protein coding |
| KLHL4 | -2.220 | Downregulation | 2.2897E-09 | protein coding |
| PCDH11X | -3.304 | Downregulation | 1.428E-19 | protein coding |
| NALCN | -2.020 | Downregulation | 1.5955E-07 | protein coding |
| MEDAG | -2.104 | Downregulation | 2.2629E-14 | protein coding |
| MT4 | -7.591 | Downregulation | 3.7123E-31 | protein coding |
| NDRG4 | -3.201 | Downregulation | 2.0339E-25 | protein coding |
| SALL1 | -2.030 | Downregulation | 1.625E-08 | protein coding |
| TGM5 | -2.996 | Downregulation | 1.7248E-15 | protein coding |
| EYA1 | -3.361 | Downregulation | 1.2387E-13 | protein coding |
| SFRP1 | -2.228 | Downregulation | 5.7807E-11 | protein coding |
| TRPS1 | -2.035 | Downregulation | 6.0432E-27 | protein coding |
| GML | -2.695 | Downregulation | 2.2757E-08 | protein coding |
| TULP2 | -2.309 | Downregulation | 9.2585E-12 | protein coding |
| LGALS13 | -2.947 | Downregulation | 5.7922E-07 | protein coding |
| CRX | -2.782 | Downregulation | 2.8986E-07 | protein coding |
| CNFN | -4.660 | Downregulation | 5.3268E-35 | protein coding |
| CABP5 | -2.895 | Downregulation | 7.5393E-09 | protein coding |
| UPK1A | -7.539 | Downregulation | 9.358E-102 | protein coding |
| ATP4A | -2.802 | Downregulation | 2.3017E-08 | protein coding |
| ANKRD7 | -2.011 | Downregulation | 5.7027E-10 | protein coding |
| EVX1 | -2.012 | Downregulation | 4.7915E-05 | protein coding |
| CRHR2 | -2.562 | Downregulation | 3.5932E-08 | protein coding |
| GHRHR | -2.892 | Downregulation | 3.9408E-11 | protein coding |
| HYAL4 | -2.680 | Downregulation | 1.66E-08 | protein coding |
| MEOX2 | -2.896 | Downregulation | 3.3007E-18 | protein coding |

| | | | | |
|---|---|---|---|---|
| POU6F2 | -2.273 | Downregulation | 1.4114E-08 | protein coding |
| OGN | -3.408 | Downregulation | 1.4983E-09 | protein coding |
| PTGDS | -2.692 | Downregulation | 1.854E-12 | protein coding |
| CXCL12 | -2.477 | Downregulation | 4.633E-15 | protein coding |
| GDF10 | -2.104 | Downregulation | 1.5996E-09 | protein coding |
| SORCS1 | -3.063 | Downregulation | 4.4475E-19 | protein coding |
| CYP2C18 | -2.132 | Downregulation | 7.8769E-08 | protein coding |
| ASPA | -2.403 | Downregulation | 2.6929E-11 | protein coding |
| SLC6A4 | -2.983 | Downregulation | 2.8328E-09 | protein coding |
| KRT32 | -4.429 | Downregulation | 3.3601E-34 | protein coding |
| SGCA | -2.134 | Downregulation | 2.6455E-08 | protein coding |
| ALOX12 | -6.458 | Downregulation | 8.6744E-66 | protein coding |
| PPY | -2.526 | Downregulation | 5.5331E-06 | protein coding |
| SLC16A6 | -2.654 | Downregulation | 7.7693E-35 | protein coding |
| EFNB3 | -2.693 | Downregulation | 3.1412E-28 | protein coding |
| GABRA4 | -3.501 | Downregulation | 4.1749E-18 | protein coding |
| GNRHR | -3.050 | Downregulation | 2.2799E-15 | protein coding |
| CWH43 | -4.043 | Downregulation | 2.7298E-18 | protein coding |
| CRYAB | -2.515 | Downregulation | 2.3191E-13 | protein coding |
| ZBTB16 | -3.274 | Downregulation | 6.7803E-22 | protein coding |
| CCND1 | -2.255 | Downregulation | 6.2757E-20 | protein coding |
| ELMOD1 | -5.640 | Downregulation | 1.9082E-31 | protein coding |
| PTPN5 | -2.842 | Downregulation | 8.1477E-16 | protein coding |
| DAO | -3.176 | Downregulation | 4.6163E-13 | protein coding |
| SYT10 | -2.196 | Downregulation | 1.4238E-05 | protein coding |
| ACSS3 | -2.129 | Downregulation | 1.8341E-11 | protein coding |
| GLI1 | -2.119 | Downregulation | 6.2139E-09 | protein coding |
| FGF6 | -3.503 | Downregulation | 2.0695E-10 | protein coding |
| ART4 | -2.289 | Downregulation | 2.0644E-13 | protein coding |
| RERGL | -2.209 | Downregulation | 2.005E-06 | protein coding |
| ENDOU | -6.104 | Downregulation | 9.6869E-59 | protein coding |
| GPR133 | -2.886 | Downregulation | 5.9892E-15 | protein coding |
| GYS2 | -6.081 | Downregulation | 1.5949E-42 | protein coding |
| AICDA | -2.445 | Downregulation | 1.9768E-07 | protein coding |
| RFX4 | -2.955 | Downregulation | 1.347E-09 | protein coding |
| RBM24 | -2.096 | Downregulation | 7.7582E-08 | protein coding |

| | | | | |
|---|---|---|---|---|
| GPR63 | -2.065 | Downregulation | 1.4066E-19 | protein coding |
| COL9A1 | -5.224 | Downregulation | 5.7054E-21 | protein coding |
| GPLD1 | -2.382 | Downregulation | 1.421E-13 | protein coding |
| UNC93A | -5.134 | Downregulation | 1.3635E-26 | protein coding |
| SLC22A2 | -2.658 | Downregulation | 2.5761E-09 | protein coding |
| WISP3 | -2.350 | Downregulation | 6.5945E-07 | protein coding |
| C7 | -2.324 | Downregulation | 2.6654E-07 | protein coding |
| SLC27A6 | -4.397 | Downregulation | 1.5833E-24 | protein coding |
| AGXT2 | -2.887 | Downregulation | 2.0945E-09 | protein coding |
| GNAT1 | -2.491 | Downregulation | 1.0038E-10 | protein coding |
| PEX5L | -2.263 | Downregulation | 2.2721E-07 | protein coding |
| AADAC | -2.456 | Downregulation | 2.8405E-07 | protein coding |
| PDE1A | -2.092 | Downregulation | 8.5535E-13 | protein coding |
| TACR1 | -2.491 | Downregulation | 2.3453E-09 | protein coding |
| IGFBP5 | -2.409 | Downregulation | 5.1319E-10 | protein coding |
| PLCD4 | -3.180 | Downregulation | 8.1198E-35 | protein coding |
| IL1R2 | -2.694 | Downregulation | 1.1506E-18 | protein coding |
| CD207 | -2.098 | Downregulation | 1.4711E-09 | protein coding |
| PAPPA2 | -3.047 | Downregulation | 5.6756E-14 | protein coding |
| ANGPTL1 | -3.131 | Downregulation | 8.1747E-13 | protein coding |
| SLC5A9 | -2.672 | Downregulation | 6.7785E-16 | protein coding |
| ELOVL4 | -3.286 | Downregulation | 1.2214E-19 | protein coding |
| ARG1 | -7.910 | Downregulation | 1.3163E-51 | protein coding |
| TCF21 | -2.187 | Downregulation | 1.2283E-06 | protein coding |
| PPL | -2.688 | Downregulation | 2.3376E-33 | protein coding |
| C2orf40 | -2.393 | Downregulation | 6.2353E-08 | protein coding |
| SLC46A2 | -2.394 | Downregulation | 2.1222E-08 | protein coding |
| NR4A3 | -2.914 | Downregulation | 6.6447E-22 | protein coding |
| LGALSL | -3.059 | Downregulation | 2.6565E-46 | protein coding |
| PPP1R3C | -4.876 | Downregulation | 1.7741E-40 | protein coding |
| CRHR1 | -2.245 | Downregulation | 4.6794E-09 | protein coding |
| DUSP1 | -2.275 | Downregulation | 5.9269E-15 | protein coding |
| TP53AIP1 | -2.641 | Downregulation | 2.445E-12 | protein coding |
| ARR3 | -3.208 | Downregulation | 2.7871E-11 | protein coding |
| MYOT | -2.689 | Downregulation | 1.0478E-08 | protein coding |
| EGR1 | -2.525 | Downregulation | 5.039E-17 | protein coding |

| | | | | |
|---|---|---|---|---|
| CHRNA2 | -2.782 | Downregulation | 3.4542E-11 | protein coding |
| ADRA1A | -3.407 | Downregulation | 1.9009E-11 | protein coding |
| PDLIM2 | -2.101 | Downregulation | 5.1906E-25 | protein coding |
| RGSL1 | -2.071 | Downregulation | 3.3802E-07 | protein coding |
| LHX4 | -2.953 | Downregulation | 2.1093E-19 | protein coding |
| CSTA | -2.890 | Downregulation | 5.3739E-16 | protein coding |
| GJB6 | -3.063 | Downregulation | 5.271E-17 | protein coding |
| XPNPEP2 | -2.422 | Downregulation | 1.242E-10 | protein coding |
| PTGFR | -2.260 | Downregulation | 1.0159E-08 | protein coding |
| KIAA1045 | -2.171 | Downregulation | 1.7208E-16 | protein coding |
| CNTFR | -2.205 | Downregulation | 1.0041E-07 | protein coding |
| NECAB1 | -2.410 | Downregulation | 5.3947E-08 | protein coding |
| NR4A1 | -2.136 | Downregulation | 3.9006E-07 | protein coding |
| PLP1 | -3.402 | Downregulation | 7.2071E-14 | protein coding |
| PI3 | -2.486 | Downregulation | 3.0839E-07 | protein coding |
| EDN3 | -5.911 | Downregulation | 7.7679E-25 | protein coding |
| PTGIS | -2.742 | Downregulation | 5.2128E-11 | protein coding |
| XG | -2.463 | Downregulation | 3.1347E-15 | protein coding |
| HIF3A | -2.774 | Downregulation | 8.0902E-17 | protein coding |
| LYPD3 | -2.121 | Downregulation | 8.6662E-14 | protein coding |
| CRISP2 | -2.842 | Downregulation | 3.7273E-07 | protein coding |
| GRM4 | -2.092 | Downregulation | 7.3803E-10 | protein coding |
| EREG | -5.099 | Downregulation | 1.3765E-36 | protein coding |
| MYH2 | -2.497 | Downregulation | 0.00028015 | protein coding |
| GRIA3 | -2.715 | Downregulation | 2.1036E-21 | protein coding |
| FOSB | -3.013 | Downregulation | 1.2653E-12 | protein coding |
| TGM3 | -5.635 | Downregulation | 5.5724E-27 | protein coding |
| PCSK2 | -4.448 | Downregulation | 1.1563E-27 | protein coding |
| F10 | -3.094 | Downregulation | 3.8031E-19 | protein coding |
| SLURP1 | -5.472 | Downregulation | 1.1698E-34 | protein coding |
| KRT36 | -3.108 | Downregulation | 3.2623E-10 | protein coding |
| FLRT1 | -3.005 | Downregulation | 4.0448E-12 | protein coding |
| STATH | -2.097 | Downregulation | 0.00043742 | protein coding |
| PRDM7 | -2.764 | Downregulation | 7.2247E-09 | protein coding |
| AIF1L | -2.078 | Downregulation | 1.6444E-16 | protein coding |
| TMEM35 | -2.307 | Downregulation | 7.1914E-11 | protein coding |

| | | | | |
|---|---|---|---|---|
| ZC4H2 | -2.089 | Downregulation | 5.1304E-20 | protein coding |
| OMD | -2.834 | Downregulation | 1.3869E-12 | protein coding |
| MASP1 | -3.002 | Downregulation | 1.0884E-20 | protein coding |
| BEST3 | -2.574 | Downregulation | 2.1313E-08 | protein coding |
| SPINK2 | -3.436 | Downregulation | 1.5353E-21 | protein coding |
| RFPL1 | -3.097 | Downregulation | 2.7115E-08 | protein coding |
| CPA4 | -2.260 | Downregulation | 6.3649E-09 | protein coding |
| FOXP2 | -2.065 | Downregulation | 4.416E-10 | protein coding |
| CGNL1 | -2.406 | Downregulation | 7.5628E-13 | protein coding |
| BBOX1 | -3.603 | Downregulation | 2.5436E-32 | protein coding |
| KCNC1 | -2.659 | Downregulation | 6.0131E-07 | protein coding |
| KLK14 | -5.629 | Downregulation | 3.6136E-35 | protein coding |
| KLK10 | -2.906 | Downregulation | 4.4104E-14 | protein coding |
| KLK8 | -2.742 | Downregulation | 1.1641E-09 | protein coding |
| PRRG3 | -2.313 | Downregulation | 3.0229E-10 | protein coding |
| CNN1 | -2.924 | Downregulation | 6.9936E-13 | protein coding |
| MAS1 | -3.021 | Downregulation | 9.4397E-09 | protein coding |
| ACSBG2 | -2.753 | Downregulation | 1.0534E-10 | protein coding |
| EPO | -3.146 | Downregulation | 1.1748E-22 | protein coding |
| KIAA1683 | -2.059 | Downregulation | 1.9E-17 | protein coding |
| SAG | -3.008 | Downregulation | 9.0388E-11 | protein coding |
| EDA2R | -2.137 | Downregulation | 1.0671E-15 | protein coding |
| GFAP | -3.386 | Downregulation | 4.9353E-12 | protein coding |
| TEX101 | -3.721 | Downregulation | 3.0733E-21 | protein coding |
| CA6 | -3.061 | Downregulation | 1.245E-09 | protein coding |
| CKMT2 | -2.063 | Downregulation | 7.5099E-11 | protein coding |
| KRT34 | -2.062 | Downregulation | 3.8952E-07 | protein coding |
| KRT33B | -3.432 | Downregulation | 8.0083E-11 | protein coding |
| RAI2 | -2.018 | Downregulation | 1.1577E-10 | protein coding |
| ABHD12B | -2.608 | Downregulation | 4.6682E-06 | protein coding |
| DDC | -2.422 | Downregulation | 9.6548E-09 | protein coding |
| RGS22 | -2.107 | Downregulation | 5.7962E-09 | protein coding |
| RHBG | -2.913 | Downregulation | 4.9875E-11 | protein coding |
| BCAN | -2.115 | Downregulation | 4.9964E-09 | protein coding |
| CRP | -2.376 | Downregulation | 2.8968E-07 | protein coding |
| LGR6 | -2.076 | Downregulation | 5.2428E-09 | protein coding |

| | | | | |
|---|---|---|---|---|
| DCLK1 | -2.645 | Downregulation | 4.6462E-15 | protein coding |
| MYH11 | -2.588 | Downregulation | 2.0972E-09 | protein coding |
| SPINK5 | -4.519 | Downregulation | 9.2653E-28 | protein coding |
| LYVE1 | -2.697 | Downregulation | 2.3705E-12 | protein coding |
| TRPM1 | -3.005 | Downregulation | 6.3463E-14 | protein coding |
| GSTM5 | -3.153 | Downregulation | 3.8437E-19 | protein coding |
| EMP1 | -2.941 | Downregulation | 6.3154E-25 | protein coding |
| DSC2 | -2.166 | Downregulation | 6.3472E-16 | protein coding |
| DSG1 | -6.776 | Downregulation | 4.4858E-41 | protein coding |
| DSC1 | -6.953 | Downregulation | 2.2241E-45 | protein coding |
| KLB | -2.678 | Downregulation | 2.7758E-24 | protein coding |
| ANXA1 | -2.065 | Downregulation | 1.7808E-16 | protein coding |
| OCM2 | -3.398 | Downregulation | 2.5565E-10 | protein coding |
| BAI3 | -3.036 | Downregulation | 2.7194E-18 | protein coding |
| EPHA7 | -2.333 | Downregulation | 3.1638E-08 | protein coding |
| FAIM2 | -2.047 | Downregulation | 3.9416E-09 | protein coding |
| STAB2 | -3.191 | Downregulation | 1.0055E-13 | protein coding |
| SCEL | -3.290 | Downregulation | 5.2727E-21 | protein coding |
| DGKB | -2.603 | Downregulation | 4.7949E-09 | protein coding |
| SCN7A | -3.138 | Downregulation | 1.632E-11 | protein coding |
| IL36A | -2.468 | Downregulation | 0.00011063 | protein coding |
| IL36RN | -2.783 | Downregulation | 7.9244E-09 | protein coding |
| IL36B | -2.651 | Downregulation | 1.7541E-07 | protein coding |
| IL1F10 | -3.453 | Downregulation | 4.2813E-13 | protein coding |
| KLF4 | -2.565 | Downregulation | 7.8666E-27 | protein coding |
| TMOD1 | -2.032 | Downregulation | 2.1437E-10 | protein coding |
| CCL21 | -2.531 | Downregulation | 1.6032E-09 | protein coding |
| TUBB2A | -2.131 | Downregulation | 3.0143E-17 | protein coding |
| CPEB2 | -2.251 | Downregulation | 6.9932E-49 | protein coding |
| PI15 | -2.963 | Downregulation | 4.529E-14 | protein coding |
| BTG4 | -2.067 | Downregulation | 5.9658E-07 | protein coding |
| ARHGAP20 | -2.371 | Downregulation | 4.3771E-22 | protein coding |
| UNC13C | -3.498 | Downregulation | 2.6708E-25 | protein coding |
| PAQR5 | -2.769 | Downregulation | 3.9702E-24 | protein coding |
| SEMA6D | -2.782 | Downregulation | 3.0228E-20 | protein coding |
| GCOM1 | -3.871 | Downregulation | 3.0516E-39 | protein coding |

| | | | | |
|---|---|---|---|---|
| LHCGR | -2.801 | Downregulation | 4.9668E-09 | protein coding |
| SULT6B1 | -2.331 | Downregulation | 5.4369E-06 | protein coding |
| AOX1 | -2.346 | Downregulation | 2.9273E-15 | protein coding |
| PCDH10 | -2.538 | Downregulation | 1.0113E-06 | protein coding |
| AGPAT9 | -2.039 | Downregulation | 8.2816E-19 | protein coding |
| FGF2 | -2.123 | Downregulation | 3.9995E-12 | protein coding |
| MMRN1 | -2.910 | Downregulation | 6.2569E-18 | protein coding |
| PLCZ1 | -2.391 | Downregulation | 1.149E-06 | protein coding |
| PIANP | -2.047 | Downregulation | 2.9884E-09 | protein coding |
| PPFIA2 | -2.188 | Downregulation | 3.8807E-14 | protein coding |
| TPH2 | -2.624 | Downregulation | 5.7527E-09 | protein coding |
| GLTP | -2.145 | Downregulation | 1.0162E-28 | protein coding |
| SRRM4 | -3.016 | Downregulation | 4.1969E-11 | protein coding |
| NOVA1 | -2.831 | Downregulation | 2.8522E-13 | protein coding |
| RDH12 | -5.816 | Downregulation | 1.5555E-57 | protein coding |
| FGF7 | -2.284 | Downregulation | 2.7019E-08 | protein coding |
| CYP11A1 | -2.200 | Downregulation | 1.657E-06 | protein coding |
| CYP1A1 | -4.099 | Downregulation | 2.32E-13 | protein coding |
| RHCG | -3.092 | Downregulation | 1.2321E-15 | protein coding |
| MYOCD | -2.665 | Downregulation | 2.1836E-10 | protein coding |
| SPATA22 | -2.847 | Downregulation | 6.8917E-12 | protein coding |
| SPACA3 | -2.600 | Downregulation | 6.6308E-08 | protein coding |
| ABCA8 | -2.363 | Downregulation | 8.1086E-10 | protein coding |
| ASXL3 | -2.337 | Downregulation | 5.3929E-11 | protein coding |
| ADCYAP1 | -3.759 | Downregulation | 1.0991E-15 | protein coding |
| GREB1L | -3.423 | Downregulation | 5.4102E-19 | protein coding |
| ARL5C | -2.576 | Downregulation | 1.6621E-08 | protein coding |
| DMRTC2 | -2.826 | Downregulation | 7.8524E-07 | protein coding |
| SLC6A3 | -2.391 | Downregulation | 2.0245E-11 | protein coding |
| KLK3 | -2.197 | Downregulation | 7.8163E-05 | protein coding |
| IL22RA1 | -2.023 | Downregulation | 1.8393E-15 | protein coding |
| CYR61 | -2.105 | Downregulation | 3.9319E-12 | protein coding |
| PROK1 | -2.443 | Downregulation | 1.239E-07 | protein coding |
| RXRG | -2.656 | Downregulation | 1.2961E-09 | protein coding |
| DPT | -2.946 | Downregulation | 6.6939E-14 | protein coding |
| CRABP2 | -2.187 | Downregulation | 1.0273E-11 | protein coding |

| | | | | |
|---|---|---|---|---|
| HMCN1 | -2.047 | Downregulation | 4.1435E-17 | protein coding |
| ANXA9 | -2.615 | Downregulation | 1.3352E-12 | protein coding |
| FLG2 | -7.826 | Downregulation | 1.5378E-30 | protein coding |
| CRNN | -6.588 | Downregulation | 3.9711E-47 | protein coding |
| S100A7 | -2.199 | Downregulation | 7.7341E-05 | protein coding |
| FLG | -7.247 | Downregulation | 2.8776E-39 | protein coding |
| GDF7 | -2.510 | Downregulation | 1.0449E-15 | protein coding |
| ATP6V1C2 | -2.585 | Downregulation | 3.5773E-08 | protein coding |
| ST6GAL2 | -2.023 | Downregulation | 6.2208E-11 | protein coding |
| THSD7B | -3.030 | Downregulation | 9.4401E-20 | protein coding |
| GALNT13 | -2.111 | Downregulation | 1.9457E-08 | protein coding |
| CPNE9 | -2.105 | Downregulation | 7.3098E-12 | protein coding |
| SLC22A14 | -2.176 | Downregulation | 2.9502E-18 | protein coding |
| BOC | -2.531 | Downregulation | 2.1224E-18 | protein coding |
| EPHA5 | -4.042 | Downregulation | 9.8843E-21 | protein coding |
| CORIN | -2.239 | Downregulation | 7.51E-12 | protein coding |
| ANK2 | -2.642 | Downregulation | 5.2081E-14 | protein coding |
| CDH18 | -2.824 | Downregulation | 7.6975E-15 | protein coding |
| ANKRD31 | -4.845 | Downregulation | 4.803E-71 | protein coding |
| CRHBP | -2.289 | Downregulation | 2.8546E-13 | protein coding |
| ADAMTS19 | -2.480 | Downregulation | 1.1132E-05 | protein coding |
| CXCL14 | -2.050 | Downregulation | 4.0476E-10 | protein coding |
| SPINK7 | -6.989 | Downregulation | 2.8203E-43 | protein coding |
| TENM2 | -3.257 | Downregulation | 9.8112E-15 | protein coding |
| LRRTM2 | -2.451 | Downregulation | 1.0982E-17 | protein coding |
| HMGCLL1 | -2.278 | Downregulation | 9.8898E-07 | protein coding |
| VIP | -2.419 | Downregulation | 3.8012E-11 | protein coding |
| SPIN2A | -2.244 | Downregulation | 2.7173E-11 | protein coding |
| RAB41 | -2.901 | Downregulation | 5.0718E-09 | protein coding |
| AWAT2 | -2.068 | Downregulation | 1.2767E-05 | protein coding |
| STAR | -2.262 | Downregulation | 6.3294E-15 | protein coding |
| SNTG1 | -3.213 | Downregulation | 4.6357E-10 | protein coding |
| PMP2 | -4.371 | Downregulation | 7.4216E-19 | protein coding |
| RSPO2 | -3.003 | Downregulation | 9.6303E-13 | protein coding |
| FAM135B | -2.285 | Downregulation | 6.1412E-08 | protein coding |
| FAM13C | -2.493 | Downregulation | 5.2586E-24 | protein coding |

| | | | | |
|---|---|---|---|---|
| CDHR1 | -2.405 | Downregulation | 1.3861E-08 | protein coding |
| NKX6-2 | -5.080 | Downregulation | 1.2761E-10 | protein coding |
| LRRC4C | -2.255 | Downregulation | 1.2015E-08 | protein coding |
| NCAM1 | -3.811 | Downregulation | 4.9012E-28 | protein coding |
| HTR3B | -4.788 | Downregulation | 4.2923E-33 | protein coding |
| ADAM33 | -2.178 | Downregulation | 7.6935E-12 | protein coding |
| SCN2B | -3.826 | Downregulation | 1.2411E-26 | protein coding |
| TM7SF2 | -2.045 | Downregulation | 4.039E-18 | protein coding |
| PCDH15 | -2.722 | Downregulation | 7.8129E-12 | protein coding |
| GPM6A | -3.133 | Downregulation | 2.4307E-13 | protein coding |
| DLG2 | -2.639 | Downregulation | 4.5374E-27 | protein coding |
| IL18 | -2.577 | Downregulation | 5.5372E-33 | protein coding |
| ZNF385D | -2.698 | Downregulation | 7.4383E-17 | protein coding |
| SLC35F4 | -2.192 | Downregulation | 1.6377E-07 | protein coding |
| GABRA2 | -3.025 | Downregulation | 5.5532E-15 | protein coding |
| GFRA1 | -3.248 | Downregulation | 8.1294E-18 | protein coding |
| GRID2 | -3.357 | Downregulation | 3.219E-21 | protein coding |
| RIT2 | -2.677 | Downregulation | 1.884E-08 | protein coding |
| OLAH | -2.651 | Downregulation | 8.8147E-12 | protein coding |
| GRIA4 | -2.306 | Downregulation | 1.9187E-05 | protein coding |
| SPARCL1 | -2.222 | Downregulation | 1.2925E-17 | protein coding |
| GJA1 | -2.261 | Downregulation | 3.3435E-22 | protein coding |
| DDX4 | -2.952 | Downregulation | 6.258E-13 | protein coding |
| CLEC4F | -2.057 | Downregulation | 8.4327E-13 | protein coding |
| CNTNAP4 | -2.433 | Downregulation | 6.1201E-10 | protein coding |
| RAB3C | -2.330 | Downregulation | 6.4839E-11 | protein coding |
| SCN3A | -2.024 | Downregulation | 1.0148E-12 | protein coding |
| PTPRD | -2.220 | Downregulation | 5.9705E-10 | protein coding |
| JAZF1 | -2.114 | Downregulation | 1.4649E-16 | protein coding |
| SLC5A10 | -2.833 | Downregulation | 2.7106E-22 | protein coding |
| ABI3BP | -3.149 | Downregulation | 4.3475E-18 | protein coding |
| CERS3 | -2.047 | Downregulation | 2.941E-08 | protein coding |
| L3MBTL4 | -2.076 | Downregulation | 1.2852E-15 | protein coding |
| JAM2 | -2.034 | Downregulation | 6.5835E-15 | protein coding |
| ADAMTS5 | -2.161 | Downregulation | 5.4154E-12 | protein coding |
| CNTNAP5 | -3.186 | Downregulation | 6.8498E-11 | protein coding |

| | | | | |
|---|---|---|---|---|
| GRIA1 | -3.034 | Downregulation | 5.4527E-17 | protein coding |
| WIF1 | -4.175 | Downregulation | 6.0787E-15 | protein coding |
| PPEF2 | -2.765 | Downregulation | 1.6651E-13 | protein coding |
| CLDN17 | -7.004 | Downregulation | 1.131E-36 | protein coding |
| CLDN8 | -3.114 | Downregulation | 4.8194E-10 | protein coding |
| TIMP4 | -2.038 | Downregulation | 1.5093E-08 | protein coding |
| TRIM63 | -2.540 | Downregulation | 4.7747E-11 | protein coding |
| FAM46B | -2.412 | Downregulation | 2.4028E-20 | protein coding |
| NRG2 | -2.158 | Downregulation | 6.3139E-12 | protein coding |
| HMHB1 | -3.040 | Downregulation | 3.1701E-08 | protein coding |
| CDA | -2.917 | Downregulation | 8.3867E-19 | protein coding |
| APOA2 | -2.439 | Downregulation | 3.8129E-05 | protein coding |
| TNNI1 | -2.229 | Downregulation | 5.0955E-07 | protein coding |
| PRAC1 | -4.655 | Downregulation | 1.5224E-16 | protein coding |
| HOXB13 | -3.338 | Downregulation | 4.0712E-12 | protein coding |
| ACTC1 | -2.298 | Downregulation | 6.6869E-07 | protein coding |
| PLA2G4D | -5.965 | Downregulation | 1.3836E-56 | protein coding |
| LCE2B | -7.914 | Downregulation | 1.3614E-20 | protein coding |
| SPRR2G | -6.544 | Downregulation | 5.4477E-20 | protein coding |
| PGLYRP3 | -2.647 | Downregulation | 4.6409E-13 | protein coding |
| TPPP3 | -2.393 | Downregulation | 9.3047E-09 | protein coding |
| CSTB | -2.318 | Downregulation | 9.498E-20 | protein coding |
| CYP3A4 | -3.608 | Downregulation | 1.0657E-25 | protein coding |
| DMKN | -4.220 | Downregulation | 3.7551E-47 | protein coding |
| NPHS1 | -2.280 | Downregulation | 3.2046E-07 | protein coding |
| CCL16 | -3.471 | Downregulation | 9.7941E-18 | protein coding |
| CD300LG | -2.474 | Downregulation | 6.9218E-10 | protein coding |
| IZUMO2 | -2.208 | Downregulation | 3.0375E-05 | protein coding |
| KRT84 | -3.454 | Downregulation | 3.7991E-25 | protein coding |
| C1orf177 | -3.157 | Downregulation | 3.2088E-23 | protein coding |
| DIRAS3 | -2.458 | Downregulation | 1.1877E-17 | protein coding |
| AKNAD1 | -3.304 | Downregulation | 5.7963E-43 | protein coding |
| BRINP3 | -2.767 | Downregulation | 1.0658E-10 | protein coding |
| CADM3 | -3.644 | Downregulation | 1.1301E-18 | protein coding |
| FAM71A | -2.825 | Downregulation | 8.7849E-13 | protein coding |
| KLHDC8A | -2.821 | Downregulation | 6.434E-17 | protein coding |

| | | | | |
|---|---|---|---|---|
| KCNJ3 | -4.627 | Downregulation | 3.1286E-21 | protein coding |
| BNIPL | -2.660 | Downregulation | 1.6519E-19 | protein coding |
| C1QTNF7 | -3.402 | Downregulation | 1.6813E-16 | protein coding |
| LCE3D | -5.018 | Downregulation | 2.0891E-14 | protein coding |
| IVL | -3.015 | Downregulation | 7.9367E-13 | protein coding |
| SPRR3 | -4.461 | Downregulation | 3.2192E-26 | protein coding |
| SPRR2D | -2.223 | Downregulation | 2.2533E-05 | protein coding |
| PGLYRP4 | -2.222 | Downregulation | 1.3734E-10 | protein coding |
| S100A12 | -2.430 | Downregulation | 6.8718E-12 | protein coding |
| NIPAL1 | -2.022 | Downregulation | 1.3815E-21 | protein coding |
| DAPL1 | -2.959 | Downregulation | 4.6972E-15 | protein coding |
| LMOD1 | -2.043 | Downregulation | 3.5777E-09 | protein coding |
| SPTA1 | -2.083 | Downregulation | 2.7966E-08 | protein coding |
| SYNPR | -3.349 | Downregulation | 1.4581E-11 | protein coding |
| TCF23 | -3.680 | Downregulation | 2.5382E-09 | protein coding |
| TGM4 | -2.538 | Downregulation | 4.8654E-13 | protein coding |
| CLEC3B | -2.223 | Downregulation | 7.5692E-13 | protein coding |
| LRRC2 | -2.537 | Downregulation | 4.4298E-18 | protein coding |
| ETNPPL | -2.421 | Downregulation | 4.8253E-08 | protein coding |
| NDST3 | -2.581 | Downregulation | 1.7369E-11 | protein coding |
| HAND2 | -2.324 | Downregulation | 2.842E-12 | protein coding |
| HPGD | -2.935 | Downregulation | 6.1628E-20 | protein coding |
| ASB5 | -2.794 | Downregulation | 1.0789E-11 | protein coding |
| NPY1R | -2.353 | Downregulation | 6.2233E-10 | protein coding |
| RNF180 | -2.077 | Downregulation | 2.0262E-14 | protein coding |
| SCGB3A2 | -3.000 | Downregulation | 5.2923E-13 | protein coding |
| GPR111 | -2.964 | Downregulation | 3.2521E-31 | protein coding |
| RAET1E | -3.351 | Downregulation | 9.441E-27 | protein coding |
| PI16 | -4.597 | Downregulation | 2.367E-22 | protein coding |
| BMPER | -2.225 | Downregulation | 2.2811E-10 | protein coding |
| SLC13A4 | -3.236 | Downregulation | 4.5256E-24 | protein coding |
| CSMD3 | -3.978 | Downregulation | 9.1068E-32 | protein coding |
| FREM1 | -2.431 | Downregulation | 1.7125E-15 | protein coding |
| MAMDC2 | -2.750 | Downregulation | 3.8847E-18 | protein coding |
| SVEP1 | -2.008 | Downregulation | 1.1524E-14 | protein coding |
| PTCHD1 | -2.443 | Downregulation | 1.3315E-07 | protein coding |

| | | | | |
|---|---|---|---|---|
| FIGF | -2.092 | Downregulation | 7.6062E-07 | protein coding |
| FAT3 | -2.159 | Downregulation | 2.4864E-08 | protein coding |
| LRFN5 | -2.385 | Downregulation | 3.9915E-10 | protein coding |
| SLC16A9 | -2.055 | Downregulation | 1.3917E-12 | protein coding |
| GJB2 | -3.370 | Downregulation | 4.0444E-27 | protein coding |
| PKNOX2 | -2.842 | Downregulation | 1.925E-20 | protein coding |
| SLC18A2 | -2.929 | Downregulation | 4.8884E-22 | protein coding |
| SLC39A2 | -2.411 | Downregulation | 5.0732E-12 | protein coding |
| C12orf50 | -2.473 | Downregulation | 4.5433E-08 | protein coding |
| FAM194B | -3.180 | Downregulation | 9.4568E-12 | protein coding |
| OTOGL | -2.006 | Downregulation | 5.3857E-11 | protein coding |
| SERPINA12 | -7.654 | Downregulation | 7.91E-48 | protein coding |
| PDZRN4 | -2.940 | Downregulation | 1.2651E-11 | protein coding |
| NELL1 | -2.824 | Downregulation | 9.5445E-18 | protein coding |
| CACNB2 | -2.268 | Downregulation | 3.7211E-11 | protein coding |
| TMCO5A | -2.336 | Downregulation | 2.4736E-06 | protein coding |
| CMTM5 | -2.014 | Downregulation | 2.0679E-08 | protein coding |
| GLB1L3 | -3.754 | Downregulation | 1.0962E-28 | protein coding |
| SVOP | -2.214 | Downregulation | 2.8024E-05 | protein coding |
| SPATA19 | -3.492 | Downregulation | 5.0751E-17 | protein coding |
| TPTE | -2.875 | Downregulation | 1.1029E-09 | protein coding |
| TBATA | -3.256 | Downregulation | 1.0377E-09 | protein coding |
| SYNPO2L | -4.382 | Downregulation | 1.3085E-32 | protein coding |
| A2ML1 | -2.388 | Downregulation | 1.7616E-09 | protein coding |
| SERPINB12 | -3.028 | Downregulation | 5.9288E-08 | protein coding |
| GREM1 | -2.035 | Downregulation | 3.0809E-08 | protein coding |
| NYAP1 | -2.252 | Downregulation | 8.299E-19 | protein coding |
| CCDC178 | -3.162 | Downregulation | 1.3406E-22 | protein coding |
| RP11-93B14.6 | -2.126 | Downregulation | 5.7608E-14 | protein coding |
| SUN5 | -2.721 | Downregulation | 5.5685E-06 | protein coding |
| LOXHD1 | -2.164 | Downregulation | 1.2302E-09 | protein coding |
| CCL23 | -2.278 | Downregulation | 7.2375E-13 | protein coding |
| RBFOX3 | -2.510 | Downregulation | 3.7646E-06 | protein coding |
| ZNF229 | -2.107 | Downregulation | 7.6163E-14 | protein coding |
| ATCAY | -3.818 | Downregulation | 1.6888E-21 | protein coding |
| KLK2 | -2.392 | Downregulation | 6.484E-08 | protein coding |

| | | | | |
|---|---|---|---|---|
| KLK5 | -2.939 | Downregulation | 1.4289E-07 | protein coding |
| KLK6 | -3.259 | Downregulation | 2.6284E-12 | protein coding |
| KLK11 | -3.288 | Downregulation | 6.5523E-25 | protein coding |
| KLK13 | -4.479 | Downregulation | 2.5771E-26 | protein coding |
| KRT1 | -8.258 | Downregulation | 1.8725E-75 | protein coding |
| ACER1 | -7.167 | Downregulation | 3.4605E-84 | protein coding |
| IGFBP6 | -2.485 | Downregulation | 7.0326E-14 | protein coding |
| SOAT2 | -2.713 | Downregulation | 7.8148E-26 | protein coding |
| GSDMA | -4.812 | Downregulation | 8.0112E-32 | protein coding |
| SCARA5 | -2.741 | Downregulation | 4.2666E-08 | protein coding |
| KIF5C | -2.271 | Downregulation | 4.3152E-33 | protein coding |
| ACOX2 | -2.829 | Downregulation | 6.4165E-28 | protein coding |
| FAM107A | -2.185 | Downregulation | 2.7401E-09 | protein coding |
| MOBP | -2.962 | Downregulation | 1.8039E-11 | protein coding |
| TNXB | -3.008 | Downregulation | 5.0104E-24 | protein coding |
| PHYHIP | -3.642 | Downregulation | 3.4877E-36 | protein coding |
| SDPR | -2.408 | Downregulation | 1.1415E-19 | protein coding |
| MYL1 | -2.279 | Downregulation | 0.01971028 | protein coding |
| LRP1B | -4.053 | Downregulation | 2.2497E-31 | protein coding |
| WFDC12 | -5.141 | Downregulation | 7.6163E-14 | protein coding |
| NSG1 | -2.999 | Downregulation | 8.5298E-20 | protein coding |
| PLA2G4F | -3.177 | Downregulation | 3.4393E-27 | protein coding |
| KLK7 | -3.225 | Downregulation | 5.3971E-12 | protein coding |
| AR | -2.803 | Downregulation | 1.328E-17 | protein coding |
| GSG1L | -2.649 | Downregulation | 7.5637E-10 | protein coding |
| RSPO1 | -2.970 | Downregulation | 4.5325E-19 | protein coding |
| KCNAB1 | -2.258 | Downregulation | 5.1936E-11 | protein coding |
| NR0B1 | -2.278 | Downregulation | 5.7904E-05 | protein coding |
| GP2 | -3.500 | Downregulation | 1.7166E-12 | protein coding |
| ELSPBP1 | -3.424 | Downregulation | 9.1041E-12 | protein coding |
| SPRR1B | -3.417 | Downregulation | 3.748E-18 | protein coding |
| SPRR1A | -3.638 | Downregulation | 9.1904E-17 | protein coding |
| CRCT1 | -5.035 | Downregulation | 2.5578E-21 | protein coding |
| MUC15 | -3.196 | Downregulation | 1.053E-19 | protein coding |
| CLIC3 | -2.678 | Downregulation | 2.4343E-16 | protein coding |
| CTNND2 | -2.346 | Downregulation | 4.4886E-10 | protein coding |

| | | | | |
|---|---|---|---|---|
| NSG2 | -2.785 | Downregulation | 1.5723E-10 | protein coding |
| GYPA | -2.253 | Downregulation | 6.4951E-07 | protein coding |
| FOS | -2.386 | Downregulation | 1.9543E-12 | protein coding |
| SEMA3E | -2.857 | Downregulation | 3.9774E-10 | protein coding |
| KRT78 | -5.153 | Downregulation | 1.6136E-40 | protein coding |
| SDR9C7 | -4.063 | Downregulation | 1.1531E-21 | protein coding |
| KRT6C | -4.331 | Downregulation | 3.1169E-20 | protein coding |
| KRT4 | -4.220 | Downregulation | 8.1224E-19 | protein coding |
| KRT72 | -2.723 | Downregulation | 1.2755E-09 | protein coding |
| HSD17B13 | -3.763 | Downregulation | 2.4206E-18 | protein coding |
| DLGAP1 | -3.091 | Downregulation | 9.3602E-19 | protein coding |
| SYT9 | -3.285 | Downregulation | 3.7167E-19 | protein coding |
| GSTA4 | -2.085 | Downregulation | 1.0109E-13 | protein coding |
| PATE1 | -3.096 | Downregulation | 1.1968E-08 | protein coding |
| C8orf74 | -2.533 | Downregulation | 1.0168E-09 | protein coding |
| GRIK1 | -4.477 | Downregulation | 6.515E-30 | protein coding |
| SOSTDC1 | -4.005 | Downregulation | 3.2236E-17 | protein coding |
| KRT13 | -3.983 | Downregulation | 3.5619E-25 | protein coding |
| HOPX | -4.456 | Downregulation | 7.9797E-56 | protein coding |
| PAH | -2.082 | Downregulation | 0.00017057 | protein coding |
| ANGPTL7 | -3.485 | Downregulation | 1.6782E-08 | protein coding |
| CYP4F22 | -6.163 | Downregulation | 3.3577E-72 | protein coding |
| MAL | -6.813 | Downregulation | 1.2822E-63 | protein coding |
| LCE1D | -8.880 | Downregulation | 1.5727E-16 | protein coding |
| GPR22 | -3.491 | Downregulation | 2.0856E-11 | protein coding |
| CSDC2 | -2.132 | Downregulation | 1.7979E-07 | protein coding |
| ABCG4 | -4.439 | Downregulation | 8.0203E-44 | protein coding |
| PDZD3 | -2.267 | Downregulation | 2.6504E-08 | protein coding |
| PRSS27 | -2.608 | Downregulation | 1.0352E-14 | protein coding |
| SYNPO2 | -2.027 | Downregulation | 3.3198E-12 | protein coding |
| IL17D | -2.300 | Downregulation | 5.2762E-11 | protein coding |
| C2orf54 | -2.694 | Downregulation | 1.3647E-21 | protein coding |
| NIPAL4 | -3.022 | Downregulation | 1.124E-16 | protein coding |
| SNTG2 | -2.778 | Downregulation | 4.9619E-15 | protein coding |
| LRRC20 | -2.194 | Downregulation | 9.6374E-21 | protein coding |
| FADS6 | -3.481 | Downregulation | 5.7726E-13 | protein coding |

| | | | | |
|---|---|---|---|---|
| KRT2 | -8.444 | Downregulation | 9.0546E-71 | protein coding |
| AQPEP | -2.133 | Downregulation | 2.2134E-11 | protein coding |
| HPSE2 | -2.961 | Downregulation | 2.749E-12 | protein coding |
| ARPP21 | -3.093 | Downregulation | 8.0694E-10 | protein coding |
| BNC2 | -2.092 | Downregulation | 3.8479E-14 | protein coding |
| MAB21L3 | -2.465 | Downregulation | 2.1529E-14 | protein coding |
| LIPM | -7.045 | Downregulation | 4.0626E-33 | protein coding |
| KCNK7 | -2.832 | Downregulation | 1.9428E-13 | protein coding |
| SULT1B1 | -2.312 | Downregulation | 4.7557E-13 | protein coding |
| SPATA3 | -3.853 | Downregulation | 1.8674E-11 | protein coding |
| WFIKKN2 | -3.341 | Downregulation | 1.0187E-11 | protein coding |
| EVX2 | -3.524 | Downregulation | 6.2574E-21 | protein coding |
| CHRNA9 | -4.254 | Downregulation | 2.6882E-15 | protein coding |
| GOLGA6L2 | -3.295 | Downregulation | 5.7823E-11 | protein coding |
| ZCCHC12 | -2.738 | Downregulation | 3.0417E-08 | protein coding |
| GALNTL6 | -2.600 | Downregulation | 4.0585E-11 | protein coding |
| IGDCC3 | -2.587 | Downregulation | 6.898E-12 | protein coding |
| MYO1H | -2.155 | Downregulation | 4.0546E-16 | protein coding |
| KLK15 | -3.045 | Downregulation | 2.0644E-10 | protein coding |
| KY | -2.878 | Downregulation | 8.0814E-19 | protein coding |
| CATSPERD | -2.018 | Downregulation | 7.485E-07 | protein coding |
| CD164L2 | -2.276 | Downregulation | 1.3467E-21 | protein coding |
| ZIC4 | -2.035 | Downregulation | 0.00022712 | protein coding |
| DES | -2.349 | Downregulation | 1.2154E-05 | protein coding |
| WFDC5 | -2.587 | Downregulation | 3.5643E-07 | protein coding |
| CADM2 | -3.804 | Downregulation | 8.376E-24 | protein coding |
| SCUBE2 | -2.021 | Downregulation | 8.4441E-14 | protein coding |
| FOSL1 | -2.028 | Downregulation | 1.5597E-08 | protein coding |
| EIF4E1B | -3.073 | Downregulation | 1.2857E-09 | protein coding |
| CREG2 | -3.095 | Downregulation | 1.8345E-14 | protein coding |
| CIDEA | -6.788 | Downregulation | 5.6537E-53 | protein coding |
| LRRTM4 | -2.770 | Downregulation | 3.3112E-13 | protein coding |
| ZNF135 | -2.212 | Downregulation | 1.5201E-14 | protein coding |
| OR4N2 | -3.966 | Downregulation | 4.6301E-14 | protein coding |
| TAC4 | -3.401 | Downregulation | 4.9441E-09 | protein coding |
| GCNT4 | -2.199 | Downregulation | 1.8195E-28 | protein coding |

| | | | | |
|---|---|---|---|---|
| FIBIN | -2.309 | Downregulation | 2.4882E-09 | protein coding |
| DEFB104B | -3.321 | Downregulation | 5.4665E-06 | protein coding |
| FAM9B | -2.886 | Downregulation | 8.5991E-10 | protein coding |
| CLVS1 | -4.622 | Downregulation | 2.7804E-54 | protein coding |
| SPACA4 | -3.104 | Downregulation | 5.8372E-17 | protein coding |
| ZBED2 | -3.638 | Downregulation | 7.9686E-36 | protein coding |
| MYOZ1 | -2.248 | Downregulation | 9.8322E-09 | protein coding |
| SPINK6 | -4.816 | Downregulation | 3.5895E-14 | protein coding |
| TNP2 | -3.043 | Downregulation | 4.982E-08 | protein coding |
| DYNAP | -2.392 | Downregulation | 3.7697E-05 | protein coding |
| C5orf64 | -2.242 | Downregulation | 7.7019E-10 | protein coding |
| C5orf46 | -3.711 | Downregulation | 3.3251E-11 | protein coding |
| GDPD4 | -3.213 | Downregulation | 4.0723E-10 | protein coding |
| IGSF22 | -2.044 | Downregulation | 9.0685E-13 | protein coding |
| ALOXE3 | -4.012 | Downregulation | 4.6931E-24 | protein coding |
| SIGLECL1 | -2.622 | Downregulation | 2.6974E-07 | protein coding |
| EGR3 | -2.833 | Downregulation | 1.4272E-17 | protein coding |
| ALOX12B | -4.908 | Downregulation | 1.9082E-31 | protein coding |
| ALOX15B | -2.568 | Downregulation | 1.1133E-09 | protein coding |
| FCER1A | -2.350 | Downregulation | 2.2912E-18 | protein coding |
| NKPD1 | -2.206 | Downregulation | 3.8273E-12 | protein coding |
| NRXN1 | -3.033 | Downregulation | 1.0387E-16 | protein coding |
| C12orf40 | -3.427 | Downregulation | 1.8047E-17 | protein coding |
| F2 | -2.205 | Downregulation | 3.4157E-07 | protein coding |
| PNPLA1 | -3.331 | Downregulation | 8.8489E-15 | protein coding |
| KCTD4 | -7.019 | Downregulation | 1.1456E-21 | protein coding |
| YOD1 | -2.473 | Downregulation | 1.4488E-26 | protein coding |
| C1orf105 | -2.683 | Downregulation | 2.7148E-10 | protein coding |
| CHRM2 | -4.413 | Downregulation | 6.6089E-20 | protein coding |
| TMEM132C | -3.635 | Downregulation | 1.6538E-16 | protein coding |
| NLRP10 | -2.973 | Downregulation | 1.8517E-06 | protein coding |
| LIPF | -3.523 | Downregulation | 6.6781E-18 | protein coding |
| XKRX | -2.511 | Downregulation | 2.7078E-17 | protein coding |
| EPGN | -4.377 | Downregulation | 3.7778E-26 | protein coding |
| PLCXD3 | -3.121 | Downregulation | 1.6622E-11 | protein coding |
| GPIHBP1 | -2.418 | Downregulation | 4.3051E-16 | protein coding |

| | | | | |
|---|---|---|---|---|
| TCEAL7 | -2.159 | Downregulation | 5.5174E-12 | protein coding |
| OTOP3 | -3.085 | Downregulation | 1.0268E-08 | protein coding |
| ZNF662 | -2.115 | Downregulation | 4.3179E-18 | protein coding |
| SPNS2 | -2.391 | Downregulation | 1.8317E-14 | protein coding |
| PCP4 | -2.413 | Downregulation | 4.3025E-05 | protein coding |
| CSMD1 | -2.444 | Downregulation | 5.1433E-09 | protein coding |
| CALN1 | -3.278 | Downregulation | 1.6925E-11 | protein coding |
| CTNNA3 | -2.316 | Downregulation | 5.095E-10 | protein coding |
| SPDYE4 | -3.094 | Downregulation | 1.7621E-10 | protein coding |
| C10orf107 | -2.012 | Downregulation | 2.9017E-08 | protein coding |
| GBP6 | -2.290 | Downregulation | 2.5147E-10 | protein coding |
| TREX2 | -2.325 | Downregulation | 6.4138E-11 | protein coding |
| TNFAIP8L3 | -2.556 | Downregulation | 1.2129E-22 | protein coding |
| GPR1 | -4.147 | Downregulation | 3.8292E-46 | protein coding |
| NOG | -2.618 | Downregulation | 2.887E-12 | protein coding |
| OPCML | -2.802 | Downregulation | 8.2899E-16 | protein coding |
| ACSM2A | -3.157 | Downregulation | 5.0009E-09 | protein coding |
| PAPL | -3.383 | Downregulation | 8.0643E-13 | protein coding |
| ZNF730 | -2.005 | Downregulation | 1.0611E-11 | protein coding |
| CNTN2 | -2.111 | Downregulation | 4.2003E-11 | protein coding |
| SPRR4 | -3.912 | Downregulation | 4.3388E-11 | protein coding |
| PCDH9 | -2.796 | Downregulation | 4.9212E-14 | protein coding |
| CDR1 | -3.391 | Downregulation | 2.5415E-14 | protein coding |
| SLIT3 | -2.778 | Downregulation | 1.0734E-20 | protein coding |
| BPIFC | -5.819 | Downregulation | 1.5074E-22 | protein coding |
| SEPTIN5 | -2.077 | Downregulation | 2.402E-15 | protein coding |
| TCEAL2 | -2.033 | Downregulation | 3.0039E-06 | protein coding |
| SLC24A3 | -2.125 | Downregulation | 7.4339E-20 | protein coding |
| KRT76 | -5.939 | Downregulation | 2.7711E-30 | protein coding |
| KLHL33 | -2.337 | Downregulation | 2.6151E-13 | protein coding |
| WBSCR17 | -2.083 | Downregulation | 1.646E-09 | protein coding |
| KRT6B | -3.075 | Downregulation | 3.6817E-12 | protein coding |
| LSAMP | -2.243 | Downregulation | 5.0732E-15 | protein coding |
| DBX2 | -3.164 | Downregulation | 5.5388E-09 | protein coding |
| KRT79 | -2.676 | Downregulation | 1.225E-15 | protein coding |
| NTF3 | -2.107 | Downregulation | 1.3485E-09 | protein coding |

| | | | | |
|---|---|---|---|---|
| SMIM23 | -3.434 | Downregulation | 1.808E-09 | protein coding |
| KCNIP4 | -2.876 | Downregulation | 5.1377E-07 | protein coding |
| TMPRSS11B | -4.671 | Downregulation | 5.9399E-19 | protein coding |
| LCE3A | -5.618 | Downregulation | 5.9025E-12 | protein coding |
| LCE3E | -5.708 | Downregulation | 1.9235E-21 | protein coding |
| PLK5 | -2.162 | Downregulation | 1.5746E-06 | protein coding |
| KRT73 | -2.667 | Downregulation | 5.3611E-10 | protein coding |
| CYP4F2 | -2.459 | Downregulation | 1.8711E-18 | protein coding |
| TEX38 | -2.060 | Downregulation | 6.3173E-08 | protein coding |
| LCE5A | -4.950 | Downregulation | 1.1743E-19 | protein coding |
| LCE1E | -6.120 | Downregulation | 1.0824E-19 | protein coding |
| NAP1L3 | -2.066 | Downregulation | 7.3008E-10 | protein coding |
| KRT10 | -4.844 | Downregulation | 2.0605E-24 | protein coding |
| TRDN | -2.764 | Downregulation | 6.0849E-08 | protein coding |
| KRT3 | -4.695 | Downregulation | 2.5365E-26 | protein coding |
| NAP1L2 | -2.602 | Downregulation | 4.6259E-17 | protein coding |
| KLK12 | -5.027 | Downregulation | 4.1956E-27 | protein coding |
| MYT1L | -3.177 | Downregulation | 7.3378E-12 | protein coding |
| PDE2A | -2.240 | Downregulation | 1.5141E-21 | protein coding |
| MPPED1 | -2.846 | Downregulation | 1.1944E-09 | protein coding |
| VSIG10L | -3.446 | Downregulation | 4.3026E-36 | protein coding |
| LCE1A | -7.167 | Downregulation | 1.2858E-16 | protein coding |
| C1QL4 | -2.644 | Downregulation | 3.1166E-09 | protein coding |
| P2RY4 | -3.206 | Downregulation | 4.5868E-26 | protein coding |
| C14orf23 | -3.189 | Downregulation | 7.7776E-09 | protein coding |
| TMPRSS11A | -2.077 | Downregulation | 1.5586E-06 | protein coding |
| VSTM2B | -4.885 | Downregulation | 1.4446E-21 | protein coding |
| LCE2A | -6.252 | Downregulation | 2.9437E-13 | protein coding |
| LCE2C | -6.447 | Downregulation | 2.061E-14 | protein coding |
| LCE2D | -6.594 | Downregulation | 3.7594E-16 | protein coding |
| CIDEC | -2.443 | Downregulation | 8.6068E-06 | protein coding |
| LUZP2 | -2.878 | Downregulation | 1.0586E-18 | protein coding |
| LHFPL3 | -4.417 | Downregulation | 1.5039E-19 | protein coding |
| LCN10 | -2.746 | Downregulation | 6.2301E-15 | protein coding |
| OVCH1 | -2.251 | Downregulation | 2.0103E-11 | protein coding |
| COL14A1 | -3.311 | Downregulation | 1.9537E-28 | protein coding |

| | | | | |
|---|---|---|---|---|
| TPRG1 | -3.633 | Downregulation | 5.1991E-48 | protein coding |
| SCGB1C1 | -3.069 | Downregulation | 9.036E-10 | protein coding |
| FAM25A | -4.007 | Downregulation | 6.8631E-14 | protein coding |
| ZNF626 | -2.185 | Downregulation | 5.8142E-14 | protein coding |
| C10orf99 | -2.675 | Downregulation | 1.8013E-13 | protein coding |
| NCCRP1 | -2.752 | Downregulation | 2.1551E-16 | protein coding |
| KRTDAP | -7.901 | Downregulation | 4.9596E-67 | protein coding |
| C2orf80 | -2.920 | Downregulation | 3.2764E-08 | protein coding |
| PRELP | -2.076 | Downregulation | 6.6711E-12 | protein coding |
| SBSN | -5.466 | Downregulation | 2.9208E-34 | protein coding |
| RNF222 | -4.094 | Downregulation | 1.1643E-20 | protein coding |
| APOD | -2.783 | Downregulation | 2.5312E-12 | protein coding |
| PLAC9 | -3.015 | Downregulation | 8.9765E-17 | protein coding |
| KRT77 | -2.653 | Downregulation | 6.7434E-11 | protein coding |
| FAM180A | -2.116 | Downregulation | 1.7764E-13 | protein coding |
| FAT4 | -2.012 | Downregulation | 2.4643E-18 | protein coding |
| ZNF681 | -2.307 | Downregulation | 1.4878E-14 | protein coding |
| GREB1 | -2.465 | Downregulation | 1.1817E-08 | protein coding |
| SFTA2 | -3.846 | Downregulation | 6.1351E-19 | protein coding |
| GRM7 | -2.403 | Downregulation | 5.6804E-13 | protein coding |
| THEM5 | -5.626 | Downregulation | 5.6307E-39 | protein coding |
| ADH1B | -4.151 | Downregulation | 9.8483E-21 | protein coding |
| ZNF418 | -2.178 | Downregulation | 3.5224E-19 | protein coding |
| LCE1B | -6.600 | Downregulation | 1.5567E-18 | protein coding |
| SPRR2B | -2.838 | Downregulation | 9.5709E-05 | protein coding |
| LCE1C | -6.704 | Downregulation | 3.3093E-19 | protein coding |
| C9orf169 | -3.077 | Downregulation | 2.1092E-16 | protein coding |
| DIO3 | -2.280 | Downregulation | 6.3974E-09 | protein coding |
| FABP12 | -2.401 | Downregulation | 1.8024E-07 | protein coding |
| CYP2F1 | -3.159 | Downregulation | 2.4175E-12 | protein coding |
| BCO2 | -2.646 | Downregulation | 9.6161E-36 | protein coding |
| CFD | -2.054 | Downregulation | 1.0454E-09 | protein coding |
| FAM25G | -4.229 | Downregulation | 1.665E-12 | protein coding |
| HRNR | -4.834 | Downregulation | 3.0357E-16 | protein coding |
| AADACL2 | -6.208 | Downregulation | 1.6067E-31 | protein coding |
| DNM3 | -2.281 | Downregulation | 1.34E-48 | protein coding |

| | | | | |
|---|---|---|---|---|
| PCDH20 | -2.054 | Downregulation | 0.00016367 | protein coding |
| AKR1B10 | -2.369 | Downregulation | 9.1184E-13 | protein coding |
| CYP2A7 | -2.692 | Downregulation | 4.313E-09 | protein coding |
| ADH4 | -2.930 | Downregulation | 1.6278E-12 | protein coding |
| PEG3 | -2.666 | Downregulation | 8.9217E-15 | protein coding |
| ANKRD35 | -2.125 | Downregulation | 3.3963E-14 | protein coding |
| PLN | -2.041 | Downregulation | 1.9425E-05 | protein coding |
| ITGBL1 | -2.312 | Downregulation | 3.07E-12 | protein coding |
| SLC34A3 | -3.837 | Downregulation | 3.9711E-27 | protein coding |
| ARC | -3.346 | Downregulation | 2.1483E-18 | protein coding |
| MT-ND6 | -2.263 | Downregulation | 6.375E-11 | protein coding |
| GPX6 | -2.644 | Downregulation | 1.0142E-06 | protein coding |
| RYR3 | -2.154 | Downregulation | 5.8907E-09 | protein coding |
| C1orf68 | -4.824 | Downregulation | 1.5176E-13 | protein coding |
| SAMD5 | -2.645 | Downregulation | 4.1557E-21 | protein coding |
| LOR | -8.608 | Downregulation | 1.3906E-40 | protein coding |
| SPRR2E | -2.777 | Downregulation | 2.3976E-06 | protein coding |
| KPRP | -6.251 | Downregulation | 1.3639E-29 | protein coding |
| PNLIPRP3 | -4.926 | Downregulation | 4.8791E-15 | protein coding |
| RBM20 | -3.008 | Downregulation | 3.6295E-30 | protein coding |
| SMIM9 | -2.227 | Downregulation | 5.32E-07 | protein coding |
| RP11-451M19.3 | -2.106 | Downregulation | 9.9776E-15 | protein coding |
| LINC00632 | -2.203 | Downregulation | 2.9807E-12 | protein coding |
| LIPN | -4.906 | Downregulation | 1.6381E-32 | protein coding |
| LIPK | -5.071 | Downregulation | 1.4148E-30 | protein coding |
| TCEAL5 | -2.836 | Downregulation | 7.6158E-21 | protein coding |
| NPY4R | -2.622 | Downregulation | 7.9727E-16 | protein coding |
| C6orf25 | -2.069 | Downregulation | 1.3316E-12 | protein coding |
| LY6G6C | -5.731 | Downregulation | 9.6135E-48 | protein coding |
| CDSN | -5.434 | Downregulation | 2.8156E-17 | protein coding |
| C6orf15 | -4.292 | Downregulation | 4.0371E-11 | protein coding |
| MUC21 | -4.379 | Downregulation | 1.3191E-23 | protein coding |
| FAM153C | -2.415 | Downregulation | 3.2208E-17 | protein coding |
| MALRD1 | -2.307 | Downregulation | 5.6505E-09 | protein coding |
| SPINK9 | -7.358 | Downregulation | 1.6506E-31 | protein coding |
| C19orf69 | -3.248 | Downregulation | 5.8454E-13 | protein coding |

| | | | | |
|---|---|---|---|---|
| VIT | -2.608 | Downregulation | 1.9146E-10 | protein coding |
| MT1A | -2.759 | Downregulation | 9.2712E-16 | protein coding |
| C15orf59 | -4.331 | Downregulation | 1.1777E-38 | protein coding |
| KRT6A | -2.602 | Downregulation | 1.3688E-13 | protein coding |
| IZUMO3 | -2.474 | Downregulation | 1.562E-07 | protein coding |
| TP53TG3D | -2.914 | Downregulation | 2.396E-12 | protein coding |
| C9orf92 | -2.401 | Downregulation | 8.5089E-05 | protein coding |
| AREGB | -2.631 | Downregulation | 4.1572E-07 | protein coding |
| XKR4 | -3.477 | Downregulation | 1.7568E-19 | protein coding |
| GPX3 | -3.088 | Downregulation | 1.0127E-28 | protein coding |
| DIO1 | -2.020 | Downregulation | 2.2312E-07 | protein coding |
| IGLJ4 | -2.114 | Downregulation | 0.02720374 | IG_J gene |
| IGLJ5 | -2.063 | Downregulation | 0.02454491 | IG_J gene |
| IGLJ7 | -2.483 | Downregulation | 0.00058798 | IG_J gene |
| KRTAP3-2 | -2.926 | Downregulation | 1.3456E-05 | protein coding |
| KRT222 | -3.066 | Downregulation | 7.1883E-12 | protein coding |
| CCL14 | -3.387 | Downregulation | 2.9315E-17 | protein coding |
| ZNF726 | -2.304 | Downregulation | 8.1462E-18 | protein coding |
| SPINK13 | -2.891 | Downregulation | 6.7659E-08 | protein coding |
| CAPN14 | -2.282 | Downregulation | 9.2095E-07 | protein coding |
| DCDC2C | -2.686 | Downregulation | 2.8443E-11 | protein coding |
| COL28A1 | -2.247 | Downregulation | 1.0509E-10 | protein coding |
| C14orf164 | -2.251 | Downregulation | 1.2991E-06 | protein coding |
| DEFB115 | -2.606 | Downregulation | 0.00022528 | protein coding |
| AL138815.1 | -2.636 | Downregulation | 1.3862E-05 | protein coding |
| KIR2DL2 | -2.118 | Downregulation | 4.7059E-06 | protein coding |
| RPTN | -6.224 | Downregulation | 1.2415E-30 | protein coding |
| AC091801.1 | -2.938 | Downregulation | 2.261E-17 | protein coding |
| AC110619.2 | -2.566 | Downregulation | 3.0888E-11 | protein coding |
| KRTAP1-5 | -2.579 | Downregulation | 1.201E-08 | protein coding |
| AC007557.1 | -2.766 | Downregulation | 1.9573E-07 | protein coding |
| RP11-433C9.2 | -2.221 | Downregulation | 1.014E-06 | protein coding |
| AC022431.2 | -3.132 | Downregulation | 1.8792E-18 | protein coding |
| AC018692.2 | -2.894 | Downregulation | 9.2623E-07 | protein coding |
| IQCF3 | -3.556 | Downregulation | 5.3977E-10 | protein coding |
| AC008271.1 | -3.771 | Downregulation | 2.0428E-14 | protein coding |

| | | | | |
|---|---|---|---|---|
| RNF224 | -3.180 | Downregulation | 5.5348E-21 | protein coding |
| AC109829.1 | -3.272 | Downregulation | 6.4785E-13 | protein coding |
| LCE6A | -6.880 | Downregulation | 1.2566E-14 | protein coding |
| RP11-147C23.1 | -2.688 | Downregulation | 3.699E-07 | protein coding |
| TEX35 | -2.322 | Downregulation | 3.9315E-08 | protein coding |
| LCE1F | -6.971 | Downregulation | 3.7728E-18 | protein coding |
| PNMA2 | -2.267 | Downregulation | 4.5881E-11 | protein coding |
| SPRR2A | -2.680 | Downregulation | 5.9595E-08 | protein coding |
| PSG11 | -3.626 | Downregulation | 3.9272E-18 | protein coding |
| VSIG8 | -4.505 | Downregulation | 1.2579E-39 | protein coding |
| SPRR2F | -2.904 | Downregulation | 4.5684E-07 | protein coding |
| ASPRV1 | -4.993 | Downregulation | 2.4051E-16 | protein coding |
| GYPB | -2.989 | Downregulation | 1.027E-08 | protein coding |
| PCDHGB6 | -2.052 | Downregulation | 6.7373E-24 | protein coding |
| RP11-1102P16.1 | -2.136 | Downregulation | 6.5389E-06 | protein coding |
| RP11-998D10.1 | -3.268 | Downregulation | 1.6792E-09 | protein coding |
| CARD18 | -4.664 | Downregulation | 4.436E-23 | protein coding |
| AL050302.1 | -3.686 | Downregulation | 5.2798E-11 | protein coding |
| RP11-324D17.1 | -3.740 | Downregulation | 5.7464E-14 | protein coding |
| BLID | -2.766 | Downregulation | 1.2021E-14 | protein coding |
| MUC22 | -4.468 | Downregulation | 3.1957E-17 | protein coding |
| CCER2 | -2.110 | Downregulation | 6.675E-08 | protein coding |
| FXYD1 | -2.927 | Downregulation | 1.7823E-19 | protein coding |
| AC037199.1 | -3.068 | Downregulation | 1.0363E-05 | protein coding |
| AC005549.3 | -2.404 | Downregulation | 4.0495E-05 | protein coding |
| AP001094.1 | -2.839 | Downregulation | 3.0752E-06 | protein coding |
| AC091948.1 | -3.544 | Downregulation | 2.8398E-15 | protein coding |
| AC061975.10 | -3.588 | Downregulation | 5.9404E-10 | protein coding |
| AC019206.1 | -2.931 | Downregulation | 0.00021044 | protein coding |
| AC003043.1 | -2.342 | Downregulation | 0.00216782 | protein coding |
| AC026407.1 | -2.203 | Downregulation | 0.00454645 | protein coding |
| BAGE5 | -3.418 | Downregulation | 1.7405E-12 | protein coding |
| AC026369.1 | -2.707 | Downregulation | 3.4735E-05 | protein coding |
| AC009802.1 | -2.167 | Downregulation | 0.00261458 | protein coding |
| AC068039.1 | -2.407 | Downregulation | 0.01015602 | protein coding |
| AC011308.1 | -2.145 | Downregulation | 0.00060493 | protein coding |

| | | | | |
|---|---|---|---|---|
| AC006014.1 | -2.148 | Downregulation | 0.00027353 | protein coding |
| AL645922.1 | -2.846 | Downregulation | 0.0026799 | protein coding |
| AC108868.1 | -2.693 | Downregulation | 0.00033471 | protein coding |
| AP001925.1 | -2.681 | Downregulation | 0.00157995 | protein coding |
| AL160286.1 | -2.563 | Downregulation | 3.6451E-06 | protein coding |
| ZNF728 | -2.089 | Downregulation | 4.6132E-07 | protein coding |
| AL035681.1 | -2.681 | Downregulation | 2.737E-05 | protein coding |
| TRABD2B | -2.177 | Downregulation | 1.3461E-15 | protein coding |
| AC004528.1 | -2.822 | Downregulation | 1.8542E-05 | protein coding |
| AL645608.2 | -2.356 | Downregulation | 0.00029738 | protein coding |
| AC020629.1 | -2.971 | Downregulation | 1.0317E-05 | protein coding |
| AL359693.1 | -2.010 | Downregulation | 0.00301687 | protein coding |
| AL137026.1 | -2.844 | Downregulation | 0.00037664 | protein coding |
| AL590560.1 | -2.251 | Downregulation | 2.0794E-09 | protein coding |
| AL109659.1 | -2.769 | Downregulation | 3.5809E-07 | protein coding |
| AL162424.1 | -2.107 | Downregulation | 0.01659099 | protein coding |
| AC015660.1 | -2.150 | Downregulation | 1.9725E-05 | protein coding |
| AC012313.1 | -2.927 | Downregulation | 8.9058E-15 | protein coding |
| TPTEP1 | -3.309 | Downregulation | 5.5575E-21 | lincRNA |
| MIR202HG | -2.784 | Downregulation | 1.5962E-07 | antisense |
| LINC00917 | -3.484 | Downregulation | 3.4117E-12 | lincRNA |
| GRIK1-AS1 | -2.000 | Downregulation | 2.0674E-08 | antisense |
| LINC00302 | -6.346 | Downregulation | 2.8426E-15 | lincRNA |
| LINC00207 | -3.369 | Downregulation | 2.0668E-07 | lincRNA |
| AC107218.3 | -3.212 | Downregulation | 5.8528E-16 | antisense |
| RP11-351J23.1 | -7.072 | Downregulation | 1.1392E-71 | lincRNA |
| HCG9 | -2.010 | Downregulation | 3.0067E-08 | lincRNA |
| AC104135.3 | -2.741 | Downregulation | 2.8602E-13 | lincRNA |
| AC007277.3 | -2.422 | Downregulation | 3.5016E-11 | antisense |
| MEG3 | -2.064 | Downregulation | 7.8888E-10 | lincRNA |
| LINC01020 | -2.966 | Downregulation | 5.4291E-08 | lincRNA |
| AC004041.2 | -2.185 | Downregulation | 4.0555E-05 | antisense |
| RP11-332E3.2 | -2.379 | Downregulation | 5.9704E-05 | sense intronic |
| AC003090.1 | -2.348 | Downregulation | 2.0304E-09 | lincRNA |
| RP11-543B16.3 | -2.856 | Downregulation | 1.1157E-05 | lincRNA |
| RP11-311H10.4 | -3.140 | Downregulation | 1.4555E-08 | antisense |

| | | | | |
|---|---|---|---|---|
| RP11-113O24.3 | -3.620 | Downregulation | 2.3208E-10 | lincRNA |
| RP11-57C13.4 | -2.922 | Downregulation | 6.0718E-05 | lincRNA |
| AC008074.4 | -5.638 | Downregulation | 3.5674E-31 | lincRNA |
| AC079630.2 | -3.715 | Downregulation | 1.62E-13 | lincRNA |
| AC007381.2 | -2.293 | Downregulation | 6.9284E-09 | lincRNA |
| RP11-475D12.1 | -3.793 | Downregulation | 4.2712E-12 | lincRNA |
| ELMO1-AS1 | -2.869 | Downregulation | 1.7471E-07 | antisense |
| POU6F2-AS1 | -2.018 | Downregulation | 0.00095655 | antisense |
| RP3-369A17.4 | -2.150 | Downregulation | 1.306E-09 | lincRNA |
| LINC00570 | -2.733 | Downregulation | 1.0528E-06 | lincRNA |
| RP4-604K5.2 | -2.249 | Downregulation | 0.00028249 | antisense |
| RP11-359D14.3 | -2.454 | Downregulation | 6.5082E-05 | antisense |
| AC010880.1 | -3.871 | Downregulation | 5.4823E-09 | lincRNA |
| RP11-48F14.1 | -2.445 | Downregulation | 1.8911E-05 | lincRNA |
| RP3-470L22.1 | -3.347 | Downregulation | 7.0734E-11 | lincRNA |
| AC096669.3 | -5.098 | Downregulation | 6.9344E-28 | lincRNA |
| AC007182.6 | -4.001 | Downregulation | 1.0313E-25 | antisense |
| AC016735.1 | -2.725 | Downregulation | 3.5857E-05 | lincRNA |
| TEX26-AS1 | -2.606 | Downregulation | 5.5018E-10 | antisense |
| RP11-54O15.3 | -2.626 | Downregulation | 1.3606E-10 | antisense |
| AC002115.5 | -3.608 | Downregulation | 1.1275E-11 | lincRNA |
| PGM5-AS1 | -3.984 | Downregulation | 4.7567E-22 | antisense |
| RP11-114L10.2 | -2.806 | Downregulation | 1.9674E-08 | antisense |
| AC010082.2 | -3.167 | Downregulation | 3.4227E-10 | antisense |
| RP4-660H19.1 | -2.051 | Downregulation | 0.00388533 | lincRNA |
| AC018643.4 | -3.229 | Downregulation | 5.609E-08 | antisense |
| LINC00457 | -2.503 | Downregulation | 6.4857E-06 | lincRNA |
| TBC1D4-AS1 | -2.235 | Downregulation | 5.7738E-12 | antisense |
| AC009478.1 | -3.480 | Downregulation | 1.1734E-11 | lincRNA |
| RP11-782C8.5 | -2.157 | Downregulation | 4.8102E-08 | lincRNA |
| AC019100.3 | -2.351 | Downregulation | 1.7728E-06 | antisense |
| RP11-764K9.1 | -2.845 | Downregulation | 1.5114E-15 | lincRNA |
| RP5-1178H5.2 | -2.636 | Downregulation | 2.0066E-05 | antisense |
| RP11-736E3.1 | -2.617 | Downregulation | 1.09E-06 | antisense |
| AC096669.1 | -2.725 | Downregulation | 1.0153E-06 | lincRNA |
| CTA-134P22.2 | -3.064 | Downregulation | 1.3219E-18 | antisense |

| | | | | |
|---|---|---|---|---|
| RP11-75C9.1 | -2.595 | Downregulation | 3.2998E-10 | lincRNA |
| RP11-3P22.2 | -3.420 | Downregulation | 3.5376E-10 | lincRNA |
| AL132709.5 | -2.124 | Downregulation | 5.6419E-10 | lincRNA |
| RP11-417O11.5 | -2.254 | Downregulation | 1.3741E-15 | sense intronic |
| FGF13-AS1 | -2.533 | Downregulation | 7.3172E-11 | antisense |
| RP1-300G12.2 | -2.048 | Downregulation | 1.3864E-11 | sense intronic |
| GRM7-AS3 | -2.967 | Downregulation | 6.2404E-08 | lincRNA |
| AC093382.1 | -2.261 | Downregulation | 5.2398E-19 | lincRNA |
| RP11-735G4.1 | -2.257 | Downregulation | 3.459E-08 | antisense |
| AC104655.3 | -2.214 | Downregulation | 1.1319E-10 | lincRNA |
| RP11-340I6.8 | -2.672 | Downregulation | 1.125E-15 | lincRNA |
| RP11-550C4.6 | -2.838 | Downregulation | 0.00192284 | lincRNA |
| RP11-239E10.3 | -2.155 | Downregulation | 1.1762E-05 | lincRNA |
| AC092155.4 | -3.353 | Downregulation | 1.8345E-14 | antisense |
| RP11-292F22.5 | -2.100 | Downregulation | 7.1568E-05 | lincRNA |
| DAB1-AS1 | -2.891 | Downregulation | 7.6603E-07 | antisense |
| RP11-122M14.3 | -3.605 | Downregulation | 5.3343E-09 | lincRNA |
| RP11-435B5.6 | -2.450 | Downregulation | 3.8244E-05 | lincRNA |
| RP11-302I18.3 | -2.470 | Downregulation | 0.00268369 | lincRNA |
| RP11-543F8.2 | -3.452 | Downregulation | 1.3094E-12 | lincRNA |
| AC012593.1 | -2.563 | Downregulation | 3.8103E-07 | lincRNA |
| LINC00658 | -2.426 | Downregulation | 1.1251E-06 | lincRNA |
| SLC8A1-AS1 | -3.081 | Downregulation | 1.1507E-11 | antisense |
| RP11-267C16.1 | -3.006 | Downregulation | 2.6701E-08 | lincRNA |
| LINC00595 | -2.246 | Downregulation | 3.3083E-07 | lincRNA |
| IL21-AS1 | -3.367 | Downregulation | 3.5709E-24 | antisense |
| RP11-214D15.2 | -2.212 | Downregulation | 2.2774E-05 | antisense |
| FAM155A-IT1 | -2.333 | Downregulation | 4.2181E-10 | sense intronic |
| AC116035.1 | -3.930 | Downregulation | 4.0542E-20 | lincRNA |
| LINC00866 | -2.386 | Downregulation | 5.1743E-08 | lincRNA |
| EIF2B5-IT1 | -2.211 | Downregulation | 6.0869E-08 | sense intronic |
| AC003051.1 | -2.691 | Downregulation | 7.4901E-14 | lincRNA |
| AC018647.3 | -2.310 | Downregulation | 1.0838E-12 | lincRNA |
| CERS6-AS1 | -2.326 | Downregulation | 5.5784E-09 | antisense |
| RP11-307P5.1 | -3.171 | Downregulation | 1.4614E-17 | lincRNA |
| SNAP25-AS1 | -2.156 | Downregulation | 1.6184E-08 | antisense |

| | | | | |
|---|---|---|---|---|
| RP3-323P13.2 | -2.554 | Downregulation | 6.0381E-09 | antisense |
| RP4-781K5.4 | -3.115 | Downregulation | 1.7804E-16 | lincRNA |
| AC012368.1 | -2.168 | Downregulation | 2.6869E-14 | lincRNA |
| LINC00568 | -3.189 | Downregulation | 7.8399E-33 | lincRNA |
| RP1-40E16.11 | -3.066 | Downregulation | 3.784E-15 | antisense |
| RP11-127L20.3 | -2.823 | Downregulation | 1.7396E-19 | lincRNA |
| RP11-132M7.3 | -3.776 | Downregulation | 7.1291E-11 | antisense |
| RP11-186N15.3 | -3.160 | Downregulation | 2.9027E-13 | antisense |
| GS1-204I12.1 | -2.577 | Downregulation | 6.918E-05 | lincRNA |
| BX322559.3 | -3.211 | Downregulation | 5.385E-07 | antisense |
| RP11-15B24.5 | -2.194 | Downregulation | 5.8378E-08 | lincRNA |
| AC093159.1 | -3.631 | Downregulation | 1.437E-21 | lincRNA |
| RP5-1051H14.2 | -3.102 | Downregulation | 5.0146E-06 | lincRNA |
| AC004692.4 | -3.087 | Downregulation | 1.8923E-07 | antisense |
| RP11-138M12.1 | -2.541 | Downregulation | 1.3679E-08 | lincRNA |
| RP11-129J12.1 | -2.812 | Downregulation | 1.5984E-05 | lincRNA |
| HCG22 | -6.232 | Downregulation | 1.4724E-46 | lincRNA |
| RP1-223B1.1 | -2.171 | Downregulation | 1.2675E-05 | lincRNA |
| RP11-473E2.4 | -2.621 | Downregulation | 3.1702E-09 | lincRNA |
| HCG23 | -2.567 | Downregulation | 7.0534E-14 | antisense |
| RP11-286B14.1 | -2.529 | Downregulation | 6.2862E-09 | lincRNA |
| RP11-88I18.3 | -2.968 | Downregulation | 5.7953E-10 | sense intronic |
| RP11-80I15.4 | -2.356 | Downregulation | 1.3205E-15 | antisense |
| RP11-20J15.3 | -2.336 | Downregulation | 2.3421E-09 | antisense |
| AC004870.3 | -2.068 | Downregulation | 6.1152E-05 | lincRNA |
| RP11-397O4.1 | -2.901 | Downregulation | 1.7188E-16 | sense intronic |
| AC022201.4 | -2.827 | Downregulation | 1.0827E-07 | antisense |
| LINC00446 | -2.492 | Downregulation | 5.8374E-06 | lincRNA |
| RP4-580N22.2 | -2.436 | Downregulation | 5.5788E-10 | lincRNA |
| AC007563.3 | -3.291 | Downregulation | 6.0958E-15 | lincRNA |
| RP11-342D14.1 | -2.644 | Downregulation | 3.6207E-07 | antisense |
| RP11-44N12.2 | -2.104 | Downregulation | 4.5699E-09 | lincRNA |
| RP11-305L7.6 | -2.464 | Downregulation | 3.3675E-18 | lincRNA |
| XX-CR54.1 | -2.982 | Downregulation | 3.567E-08 | lincRNA |
| RP11-557H15.5 | -2.705 | Downregulation | 5.9115E-10 | lincRNA |
| AC019349.5 | -2.792 | Downregulation | 5.3209E-13 | antisense |

| | | | | |
|---|---|---|---|---|
| AC118754.4 | -2.412 | Downregulation | 5.3048E-22 | antisense |
| RP11-366F6.2 | -2.477 | Downregulation | 2.5154E-06 | antisense |
| RP11-96C23.10 | -4.002 | Downregulation | 2.8449E-07 | antisense |
| RP11-215N21.1 | -3.250 | Downregulation | 6.7004E-11 | lincRNA |
| RP11-95P13.1 | -2.328 | Downregulation | 1.0362E-10 | lincRNA |
| RP11-550H2.2 | -2.461 | Downregulation | 5.2975E-06 | lincRNA |
| RP11-505C13.1 | -3.579 | Downregulation | 8.3952E-19 | lincRNA |
| CTA-929C8.8 | -3.234 | Downregulation | 3.7506E-09 | lincRNA |
| AC108025.2 | -2.263 | Downregulation | 0.0003553 | antisense |
| RP3-390M24.1 | -2.947 | Downregulation | 5.0612E-07 | lincRNA |
| LINC00856 | -2.127 | Downregulation | 9.8322E-09 | lincRNA |
| RP13-329D4.3 | -3.051 | Downregulation | 2.5333E-07 | lincRNA |
| AC007403.2 | -2.674 | Downregulation | 5.2214E-11 | lincRNA |
| AC093609.1 | -2.988 | Downregulation | 1.481E-28 | lincRNA |
| RP3-331H24.4 | -3.288 | Downregulation | 4.7404E-10 | lincRNA |
| AC097532.2 | -2.685 | Downregulation | 3.6352E-06 | lincRNA |
| AC104135.4 | -2.559 | Downregulation | 1.0803E-09 | lincRNA |
| RP5-968J1.1 | -2.313 | Downregulation | 3.1914E-21 | lincRNA |
| AC023115.1 | -2.712 | Downregulation | 9.9781E-09 | lincRNA |
| RP3-525N10.2 | -2.978 | Downregulation | 3.9323E-12 | antisense |
| AC008063.2 | -3.174 | Downregulation | 1.3309E-14 | antisense |
| HAO2-IT1 | -2.714 | Downregulation | 1.3483E-05 | sense intronic |
| RP11-394O9.1 | -3.033 | Downregulation | 1.6724E-08 | lincRNA |
| LINC00160 | -3.603 | Downregulation | 3.156E-22 | lincRNA |
| RP11-428F8.2 | -3.730 | Downregulation | 2.9765E-12 | antisense |
| AC011747.6 | -2.528 | Downregulation | 1.1942E-07 | lincRNA |
| RP5-1007G16.1 | -2.696 | Downregulation | 2.6162E-05 | antisense |
| AC116609.3 | -2.705 | Downregulation | 5.6186E-05 | lincRNA |
| FARP1-AS1 | -2.293 | Downregulation | 2.3633E-12 | antisense |
| AC011752.1 | -2.519 | Downregulation | 1.3406E-08 | lincRNA |
| RP5-965F6.2 | -2.766 | Downregulation | 3.9644E-12 | lincRNA |
| RP4-799P18.4 | -2.378 | Downregulation | 1.8833E-08 | sense intronic |
| RP1-225E12.2 | -2.675 | Downregulation | 2.7215E-11 | antisense |
| AC016995.3 | -2.178 | Downregulation | 8.5651E-13 | lincRNA |
| FGF12-AS1 | -2.939 | Downregulation | 1.0362E-06 | antisense |
| CTC-490G23.2 | -3.739 | Downregulation | 3.9747E-19 | lincRNA |

| | | | | |
|---|---|---|---|---|
| RP1-251M9.2 | -3.534 | Downregulation | 2.8873E-15 | lincRNA |
| RP11-82L18.4 | -2.236 | Downregulation | 8.7999E-15 | lincRNA |
| RP11-157N3.1 | -2.765 | Downregulation | 4.5428E-07 | lincRNA |
| RP11-339N8.1 | -2.061 | Downregulation | 7.1552E-06 | lincRNA |
| RP11-573D15.3 | -2.375 | Downregulation | 6.2702E-05 | antisense |
| RP11-189B4.6 | -2.727 | Downregulation | 7.9216E-10 | lincRNA |
| AC009158.1 | -2.816 | Downregulation | 3.9996E-06 | lincRNA |
| AC012594.1 | -3.217 | Downregulation | 3.6884E-09 | antisense |
| CTA-125H2.2 | -3.060 | Downregulation | 1.1066E-08 | antisense |
| RP4-610C12.1 | -2.885 | Downregulation | 1.1505E-08 | antisense |
| RP11-395L14.4 | -3.264 | Downregulation | 6.0055E-16 | lincRNA |
| RP11-557H15.3 | -3.347 | Downregulation | 2.9747E-28 | lincRNA |
| RP11-82L20.1 | -3.232 | Downregulation | 1E-08 | antisense |
| RP11-53B5.1 | -2.780 | Downregulation | 5.4185E-06 | lincRNA |
| AL132709.8 | -2.103 | Downregulation | 2.2838E-08 | lincRNA |
| AC092168.2 | -2.126 | Downregulation | 1.1084E-14 | sense intronic |
| LINC00867 | -3.732 | Downregulation | 8.1002E-16 | lincRNA |
| RP11-370K11.1 | -2.068 | Downregulation | 2.746E-12 | sense intronic |
| AC008440.5 | -2.782 | Downregulation | 1.4203E-06 | antisense |
| RP4-715N11.2 | -2.083 | Downregulation | 8.4806E-05 | lincRNA |
| AC011286.1 | -3.354 | Downregulation | 2.8349E-16 | lincRNA |
| VIPR1-AS1 | -2.790 | Downregulation | 2.2026E-11 | antisense |
| AC016768.1 | -2.775 | Downregulation | 2.1387E-07 | lincRNA |
| RP11-313D6.3 | -2.210 | Downregulation | 6.0221E-07 | lincRNA |
| OSBPL10-AS1 | -3.029 | Downregulation | 5.7276E-08 | antisense |
| RP1-37J18.2 | -2.907 | Downregulation | 5.411E-06 | lincRNA |
| AC010090.1 | -2.630 | Downregulation | 3.8355E-08 | lincRNA |
| AC004862.6 | -3.358 | Downregulation | 1.7267E-12 | antisense |
| LINC01022 | -3.108 | Downregulation | 4.0347E-06 | antisense |
| LMLN-AS1 | -2.284 | Downregulation | 6.9063E-06 | antisense |
| RP11-429H9.4 | -3.312 | Downregulation | 6.1479E-09 | antisense |
| AC067959.1 | -2.788 | Downregulation | 7.8991E-12 | lincRNA |
| RP11-24P14.1 | -2.937 | Downregulation | 5.822E-06 | lincRNA |
| AC006372.6 | -2.039 | Downregulation | 0.00028066 | antisense |
| STXBP5-AS1 | -2.982 | Downregulation | 7.9377E-35 | antisense |
| AC005162.5 | -2.022 | Downregulation | 1.5085E-11 | antisense |

| | | | | |
|---|---|---|---|---|
| RP11-177F11.1 | -2.550 | Downregulation | 7.6615E-05 | antisense |
| RP5-1172A22.1 | -2.655 | Downregulation | 5.5515E-08 | lincRNA |
| RP4-799D16.1 | -2.413 | Downregulation | 1.6426E-07 | lincRNA |
| AC098617.1 | -2.431 | Downregulation | 6.6939E-14 | antisense |
| RP3-348I23.2 | -3.033 | Downregulation | 1.9274E-08 | antisense |
| AC103563.8 | -3.869 | Downregulation | 3.6102E-17 | antisense |
| AC007131.2 | -2.760 | Downregulation | 9.0594E-09 | lincRNA |
| RP11-310H4.2 | -2.384 | Downregulation | 1.6069E-07 | lincRNA |
| PABPC5-AS1 | -2.387 | Downregulation | 2.0685E-10 | antisense |
| AC009505.4 | -3.735 | Downregulation | 3.1879E-10 | lincRNA |
| RP11-433J22.3 | -2.685 | Downregulation | 6.5161E-07 | lincRNA |
| RP11-799O21.2 | -2.059 | Downregulation | 3.5412E-05 | lincRNA |
| AC012370.3 | -3.119 | Downregulation | 1.5346E-07 | lincRNA |
| RP11-308N19.1 | -2.037 | Downregulation | 5.4544E-07 | lincRNA |
| JAZF1-AS1 | -2.874 | Downregulation | 3.3111E-14 | antisense |
| RNF219-AS1 | -2.236 | Downregulation | 5.1329E-06 | antisense |
| AC007278.3 | -2.214 | Downregulation | 6.4254E-11 | sense intronic |
| RP11-560I19.1 | -2.977 | Downregulation | 5.2737E-08 | lincRNA |
| AC068489.1 | -3.400 | Downregulation | 1.434E-10 | lincRNA |
| RP4-784A16.1 | -2.209 | Downregulation | 2.9406E-06 | antisense |
| AC009305.1 | -2.927 | Downregulation | 7.9828E-08 | antisense |
| RP1-149A16.12 | -2.412 | Downregulation | 8.126E-14 | antisense |
| AC053503.6 | -3.064 | Downregulation | 7.2903E-08 | antisense |
| LINC00444 | -2.290 | Downregulation | 1.2205E-07 | lincRNA |
| RP11-120C12.3 | -2.103 | Downregulation | 2.1659E-06 | lincRNA |
| AC109828.1 | -2.678 | Downregulation | 1.7683E-09 | antisense |
| MLIP-AS1 | -2.844 | Downregulation | 5.2276E-07 | antisense |
| AC091153.4 | -2.716 | Downregulation | 8.6904E-07 | antisense |
| LINC00330 | -3.221 | Downregulation | 1.1163E-18 | lincRNA |
| AC068286.1 | -2.843 | Downregulation | 9.1345E-09 | lincRNA |
| RP1-60O19.1 | -2.039 | Downregulation | 2.2065E-12 | lincRNA |
| RP1-159G19.1 | -4.131 | Downregulation | 1.2346E-10 | lincRNA |
| RP11-532N4.2 | -3.125 | Downregulation | 1.4142E-11 | antisense |
| AC009411.2 | -3.385 | Downregulation | 6.4549E-09 | lincRNA |
| LINC00484 | -2.466 | Downregulation | 8.024E-14 | lincRNA |
| MAGI2-IT1 | -2.925 | Downregulation | 3.1106E-14 | sense intronic |

| | | | | |
|---|---|---|---|---|
| AC079779.5 | -2.809 | Downregulation | 5.0879E-07 | lincRNA |
| AL109767.1 | -3.001 | Downregulation | 5.1656E-10 | antisense |
| LINC01073 | -2.245 | Downregulation | 2.4301E-06 | lincRNA |
| AC012363.13 | -3.934 | Downregulation | 7.1791E-20 | lincRNA |
| AC023115.2 | -2.132 | Downregulation | 1.881E-10 | lincRNA |
| RP11-217L21.1 | -3.152 | Downregulation | 6.5921E-06 | antisense |
| RP3-523C21.2 | -2.230 | Downregulation | 5.0946E-08 | lincRNA |
| hsa-mir-7515 | -3.080 | Downregulation | 4.4255E-09 | lincRNA |
| RP1-287H17.1 | -2.487 | Downregulation | 1.4816E-14 | lincRNA |
| RP11-107M16.2 | -2.202 | Downregulation | 2.31E-09 | lincRNA |
| CTA-796E4.3 | -2.462 | Downregulation | 0.00078229 | lincRNA |
| RP11-443A13.5 | -3.319 | Downregulation | 1.4136E-09 | antisense |
| AC011284.3 | -3.247 | Downregulation | 7.127E-10 | antisense |
| AL035610.2 | -2.502 | Downregulation | 9.1481E-19 | lincRNA |
| SMAD9-AS1 | -2.086 | Downregulation | 8.5292E-08 | sense intronic |
| AL121656.5 | -2.790 | Downregulation | 1.5894E-06 | lincRNA |
| RP11-44H4.1 | -2.168 | Downregulation | 0.00027439 | lincRNA |
| AC007563.5 | -2.122 | Downregulation | 1.6568E-08 | antisense |
| AC092661.1 | -3.265 | Downregulation | 1.7773E-10 | lincRNA |
| RP11-154H17.1 | -2.090 | Downregulation | 0.00132815 | lincRNA |
| LINC01141 | -2.572 | Downregulation | 1.982E-09 | lincRNA |
| WASF3-AS1 | -3.263 | Downregulation | 8.3507E-12 | antisense |
| AC008067.2 | -2.927 | Downregulation | 1.0765E-06 | antisense |
| RP11-365O16.6 | -3.266 | Downregulation | 1.4355E-23 | antisense |
| HAND2-AS1 | -2.440 | Downregulation | 1.584E-11 | antisense |
| RP11-415D17.3 | -3.421 | Downregulation | 4.6515E-10 | antisense |
| RP11-85G21.2 | -2.691 | Downregulation | 6.3512E-06 | lincRNA |
| AC007099.2 | -2.587 | Downregulation | 6.8906E-06 | antisense |
| AC019064.1 | -4.336 | Downregulation | 4.8083E-15 | lincRNA |
| RP11-145M4.3 | -2.524 | Downregulation | 1.0296E-13 | lincRNA |
| CAMTA1-IT1 | -2.149 | Downregulation | 1.525E-05 | sense intronic |
| RP11-356I2.4 | -2.263 | Downregulation | 1.1308E-22 | antisense |
| AC012668.2 | -2.172 | Downregulation | 8.5369E-05 | lincRNA |
| LINC00407 | -2.246 | Downregulation | 2.6154E-06 | lincRNA |
| AP002856.7 | -3.694 | Downregulation | 1.0657E-12 | lincRNA |
| RP11-462G2.1 | -2.639 | Downregulation | 2.3992E-11 | lincRNA |

| | | | | |
|---|---|---|---|---|
| ERICH1-AS1 | -2.373 | Downregulation | 5.1732E-10 | antisense |
| RP5-940F7.2 | -3.230 | Downregulation | 4.7459E-15 | lincRNA |
| LINC00844 | -3.485 | Downregulation | 1.0654E-09 | lincRNA |
| FLG-AS1 | -2.795 | Downregulation | 4.2266E-25 | antisense |
| RP11-667F9.2 | -2.960 | Downregulation | 0.00194089 | lincRNA |
| AC009312.1 | -2.708 | Downregulation | 2.1257E-10 | antisense |
| ST6GAL2-IT1 | -2.861 | Downregulation | 1.1708E-11 | sense intronic |
| RP11-435B5.5 | -2.142 | Downregulation | 8.1341E-12 | lincRNA |
| RP11-384F7.2 | -3.328 | Downregulation | 1.5271E-10 | lincRNA |
| PDZRN3-AS1 | -2.288 | Downregulation | 2.8049E-08 | antisense |
| RP11-246A10.1 | -2.161 | Downregulation | 3.7894E-08 | lincRNA |
| RP13-635I23.3 | -2.914 | Downregulation | 1.4478E-23 | sense intronic |
| RP11-167H9.5 | -4.508 | Downregulation | 2.1333E-19 | lincRNA |
| RP11-344B5.3 | -2.587 | Downregulation | 5.4472E-05 | lincRNA |
| ADAMTS9-AS1 | -2.227 | Downregulation | 3.7935E-05 | antisense |
| RP11-768G7.2 | -3.612 | Downregulation | 1.4279E-11 | lincRNA |
| LINC00635 | -2.720 | Downregulation | 4.51E-08 | lincRNA |
| RP11-641D5.2 | -3.147 | Downregulation | 2.4832E-11 | antisense |
| RP11-444P10.1 | -3.463 | Downregulation | 5.5421E-09 | lincRNA |
| ADAMTS9-AS2 | -2.640 | Downregulation | 2.1899E-13 | antisense |
| CTC-340A15.2 | -3.326 | Downregulation | 3.0561E-11 | antisense |
| RP11-457K10.1 | -2.561 | Downregulation | 5.5792E-06 | lincRNA |
| CTD-2377D24.4 | -2.048 | Downregulation | 5.8103E-06 | lincRNA |
| RP11-501O2.5 | -2.471 | Downregulation | 3.8923E-05 | lincRNA |
| RP11-416O18.1 | -3.394 | Downregulation | 7.3696E-20 | lincRNA |
| RP11-435B5.3 | -2.136 | Downregulation | 4.2396E-06 | lincRNA |
| RP11-190P13.2 | -2.001 | Downregulation | 3.0352E-08 | lincRNA |
| RP11-166N6.2 | -2.142 | Downregulation | 2.2121E-06 | antisense |
| LINC00359 | -2.346 | Downregulation | 1.1191E-06 | lincRNA |
| FGF14-IT1 | -3.425 | Downregulation | 4.4106E-24 | lincRNA |
| RP11-475O23.2 | -2.919 | Downregulation | 1.576E-09 | antisense |
| RP11-521D12.5 | -3.699 | Downregulation | 1.1308E-22 | lincRNA |
| RP11-483E7.1 | -6.566 | Downregulation | 1.5813E-40 | lincRNA |
| RP11-889D3.2 | -3.012 | Downregulation | 3.0713E-09 | lincRNA |
| RP11-846F4.12 | -3.690 | Downregulation | 2.4883E-15 | antisense |
| IL12A-AS1 | -2.949 | Downregulation | 2.0573E-24 | antisense |

| | | | | |
|---|---|---|---|---|
| RP11-85M11.2 | -2.225 | Downregulation | 2.5116E-06 | lincRNA |
| ALKBH3-AS1 | -3.242 | Downregulation | 2.2925E-10 | antisense |
| LINC00968 | -2.208 | Downregulation | 4.5546E-19 | lincRNA |
| RP11-68L18.1 | -2.036 | Downregulation | 4.6237E-17 | antisense |
| RP11-44F21.2 | -2.532 | Downregulation | 4.1459E-07 | antisense |
| RP11-148L24.1 | -2.479 | Downregulation | 2.5834E-11 | lincRNA |
| CTB-174D11.1 | -3.308 | Downregulation | 8.1693E-11 | antisense |
| RP11-556I14.2 | -2.374 | Downregulation | 2.4508E-18 | lincRNA |
| MEF2C-AS1 | -2.024 | Downregulation | 4.3797E-14 | antisense |
| RP11-236J17.5 | -2.774 | Downregulation | 3.6877E-09 | sense intronic |
| LINC00504 | -3.279 | Downregulation | 7.1082E-28 | lincRNA |
| CTD-2010I22.2 | -2.763 | Downregulation | 1.231E-06 | lincRNA |
| RP11-321E2.3 | -2.058 | Downregulation | 0.0001998 | lincRNA |
| FGF10-AS1 | -3.557 | Downregulation | 9.4925E-18 | antisense |
| RP11-807H7.2 | -3.495 | Downregulation | 1.6677E-09 | lincRNA |
| RP11-125O18.1 | -2.046 | Downregulation | 9.9403E-07 | lincRNA |
| RP11-386B13.4 | -3.189 | Downregulation | 2.5294E-06 | lincRNA |
| RP11-2O17.2 | -3.402 | Downregulation | 2.2297E-13 | antisense |
| OTX2-AS1 | -3.014 | Downregulation | 2.5526E-08 | antisense |
| GDNF-AS1 | -2.377 | Downregulation | 4.0023E-11 | lincRNA |
| RP11-5N11.2 | -3.005 | Downregulation | 4.5607E-12 | sense intronic |
| CTD-2215E18.2 | -2.647 | Downregulation | 0.00023093 | antisense |
| CTD-2275D24.4 | -3.084 | Downregulation | 3.8283E-09 | lincRNA |
| CTD-2251F13.1 | -2.927 | Downregulation | 1.1069E-06 | lincRNA |
| RP11-73G16.2 | -2.724 | Downregulation | 2.3719E-08 | lincRNA |
| LINC01033 | -2.156 | Downregulation | 7.9095E-07 | lincRNA |
| RP11-241G9.3 | -2.672 | Downregulation | 5.9601E-06 | antisense |
| AC010468.2 | -3.026 | Downregulation | 4.7855E-08 | antisense |
| LINC01016 | -4.078 | Downregulation | 2.9047E-20 | lincRNA |
| RP11-434D9.1 | -2.006 | Downregulation | 3.0617E-06 | lincRNA |
| CASC11 | -2.265 | Downregulation | 1.4423E-11 | lincRNA |
| IL20RB-AS1 | -2.071 | Downregulation | 1.3759E-05 | antisense |
| ADAMTS19-AS1 | -2.455 | Downregulation | 1.7187E-05 | antisense |
| RP11-1E22.1 | -2.879 | Downregulation | 2.2249E-08 | lincRNA |
| AC091969.1 | -3.058 | Downregulation | 2.5329E-09 | lincRNA |
| RP11-438N16.1 | -2.603 | Downregulation | 6.4081E-09 | lincRNA |

| | | | | |
|---|---|---|---|---|
| RP11-619J20.1 | -3.032 | Downregulation | 1.3149E-07 | antisense |
| RP11-259O2.3 | -2.187 | Downregulation | 5.3377E-06 | lincRNA |
| RP11-60A8.1 | -4.422 | Downregulation | 2.682E-36 | lincRNA |
| CTD-2143L24.1 | -2.764 | Downregulation | 2.8936E-10 | lincRNA |
| CTD-3179P9.1 | -3.269 | Downregulation | 2.5662E-15 | lincRNA |
| RP11-314N14.1 | -3.656 | Downregulation | 6.1548E-10 | lincRNA |
| CACNA1G-AS1 | -2.242 | Downregulation | 2.7185E-09 | antisense |
| CTC-321K16.1 | -2.799 | Downregulation | 2.5101E-21 | antisense |
| RP11-305O6.3 | -2.045 | Downregulation | 1.4277E-14 | sense intronic |
| CTD-2089N3.1 | -3.835 | Downregulation | 1.4665E-17 | lincRNA |
| RP11-119J18.1 | -2.701 | Downregulation | 2.5485E-15 | lincRNA |
| RP11-257I8.2 | -2.698 | Downregulation | 6.776E-07 | lincRNA |
| RP11-83M16.6 | -2.189 | Downregulation | 7.748E-05 | lincRNA |
| RP13-577H12.2 | -2.098 | Downregulation | 0.00024754 | antisense |
| RP11-6C14.1 | -3.460 | Downregulation | 2.8663E-09 | lincRNA |
| RP11-302F12.10 | -3.347 | Downregulation | 6.3875E-11 | antisense |
| CTC-467M3.2 | -2.461 | Downregulation | 0.00033466 | antisense |
| RP11-725M22.1 | -3.030 | Downregulation | 7.4728E-07 | lincRNA |
| RP11-230G5.2 | -3.230 | Downregulation | 5.2459E-11 | lincRNA |
| RP11-55L3.1 | -3.519 | Downregulation | 2.3577E-12 | lincRNA |
| RP11-807H7.1 | -3.152 | Downregulation | 1.2633E-11 | lincRNA |
| CTD-2281M20.1 | -2.685 | Downregulation | 1.3329E-09 | lincRNA |
| AC097467.2 | -3.230 | Downregulation | 4.9109E-11 | antisense |
| KB-1639H6.4 | -2.340 | Downregulation | 4.6415E-05 | antisense |
| LNX1-AS1 | -3.162 | Downregulation | 8.9137E-11 | antisense |
| RP11-94C24.11 | -2.485 | Downregulation | 0.00055134 | antisense |
| CTD-2533K21.4 | -2.541 | Downregulation | 9.4204E-07 | lincRNA |
| F11-AS1 | -2.499 | Downregulation | 5.025E-07 | antisense |
| RP11-506N2.1 | -3.736 | Downregulation | 1.8815E-12 | lincRNA |
| RP11-427M20.1 | -2.350 | Downregulation | 1.0858E-07 | lincRNA |
| RP11-234K19.1 | -2.024 | Downregulation | 9.2998E-07 | lincRNA |
| RP11-420A23.1 | -2.192 | Downregulation | 6.0927E-27 | lincRNA |
| LINC01099 | -3.509 | Downregulation | 4.4182E-14 | lincRNA |
| RP11-381N20.1 | -2.644 | Downregulation | 1.6782E-08 | lincRNA |
| RP11-166A12.1 | -2.716 | Downregulation | 2.4407E-09 | lincRNA |
| RP11-745L13.2 | -2.189 | Downregulation | 1.1846E-05 | lincRNA |

| | | | | |
|---|---|---|---|---|
| RP11-6N13.1 | -2.522 | Downregulation | 0.0001472 | lincRNA |
| RP11-614F17.2 | -2.557 | Downregulation | 1.0502E-05 | lincRNA |
| CTC-575N7.1 | -3.331 | Downregulation | 1.9306E-13 | antisense |
| KB-1448A5.1 | -3.056 | Downregulation | 1.2127E-16 | lincRNA |
| RP1-84O15.2 | -3.380 | Downregulation | 6.4081E-09 | lincRNA |
| CTD-3023L14.2 | -2.877 | Downregulation | 4.7118E-07 | lincRNA |
| RP11-563N12.2 | -2.711 | Downregulation | 1.3048E-06 | lincRNA |
| RP11-100L22.1 | -3.006 | Downregulation | 2.49E-11 | lincRNA |
| CTD-3046C4.1 | -2.951 | Downregulation | 2.4135E-07 | lincRNA |
| LINC00599 | -3.638 | Downregulation | 9.0516E-11 | lincRNA |
| CTD-2313P7.1 | -3.762 | Downregulation | 1.0616E-09 | lincRNA |
| RP11-108E14.1 | -2.667 | Downregulation | 7.5401E-06 | lincRNA |
| RP11-513O17.2 | -2.398 | Downregulation | 1.5514E-05 | lincRNA |
| RP1-170O19.17 | -2.430 | Downregulation | 9.7942E-10 | lincRNA |
| CTA-392C11.2 | -2.873 | Downregulation | 1.9577E-06 | lincRNA |
| RP11-21C17.1 | -2.863 | Downregulation | 5.0339E-09 | lincRNA |
| EVX1-AS | -2.395 | Downregulation | 7.3244E-11 | antisense |
| RP11-114O8.1 | -2.877 | Downregulation | 2.4633E-06 | lincRNA |
| RP11-653B10.1 | -3.403 | Downregulation | 6.7926E-11 | lincRNA |
| CTD-3080F16.3 | -2.871 | Downregulation | 4.8526E-12 | sense intronic |
| RP11-586K2.1 | -2.766 | Downregulation | 5.2202E-14 | antisense |
| RP11-32K4.1 | -2.097 | Downregulation | 0.00057237 | antisense |
| RP11-351C8.1 | -2.853 | Downregulation | 5.8335E-06 | lincRNA |
| RP11-513H8.1 | -2.056 | Downregulation | 3.9561E-06 | lincRNA |
| RP11-600K15.1 | -2.258 | Downregulation | 7.1373E-09 | lincRNA |
| ZFHX4-AS1 | -3.550 | Downregulation | 2.0064E-23 | antisense |
| CTB-33O18.1 | -2.920 | Downregulation | 8.7656E-08 | lincRNA |
| KB-173C10.1 | -2.702 | Downregulation | 2.4864E-08 | lincRNA |
| RP11-909N17.2 | -3.104 | Downregulation | 4.5914E-10 | antisense |
| CTB-32H22.1 | -2.102 | Downregulation | 3.9147E-06 | lincRNA |
| NRG1-IT1 | -2.807 | Downregulation | 6.4746E-09 | sense intronic |
| RP11-326E22.1 | -2.571 | Downregulation | 1.9574E-08 | antisense |
| RP11-421P23.2 | -2.643 | Downregulation | 1.3721E-07 | lincRNA |
| RP11-730G20.1 | -6.852 | Downregulation | 1.8876E-30 | antisense |
| RP11-30J20.1 | -2.272 | Downregulation | 7.2527E-07 | lincRNA |
| RP11-386D6.1 | -3.000 | Downregulation | 3.9609E-08 | antisense |

| | | | | |
|---|---|---|---|---|
| RP11-67M9.1 | -2.964 | Downregulation | 2.6204E-09 | lincRNA |
| RP1-273G13.3 | -2.581 | Downregulation | 2.7733E-05 | lincRNA |
| AC008691.1 | -3.577 | Downregulation | 3.5546E-11 | lincRNA |
| RP11-622O11.4 | -2.752 | Downregulation | 9.4626E-07 | lincRNA |
| RP11-115J16.1 | -2.709 | Downregulation | 5.2759E-07 | lincRNA |
| RP11-466I1.1 | -2.681 | Downregulation | 3.0449E-08 | lincRNA |
| RP11-115E19.1 | -2.817 | Downregulation | 2.1164E-06 | lincRNA |
| CTD-2530H12.2 | -2.052 | Downregulation | 3.0302E-08 | sense intronic |
| RP1-17K7.3 | -3.225 | Downregulation | 2.033E-06 | antisense |
| RP11-831A10.2 | -2.399 | Downregulation | 1.1198E-05 | lincRNA |
| RP11-152H18.4 | -3.571 | Downregulation | 3.4916E-09 | antisense |
| CTD-2562J17.9 | -2.346 | Downregulation | 0.00035583 | antisense |
| RP11-1081L13.4 | -2.241 | Downregulation | 1.9417E-12 | antisense |
| RP11-680F20.6 | -5.421 | Downregulation | 3.9779E-46 | antisense |
| RP11-65M17.3 | -2.270 | Downregulation | 1.1066E-08 | lincRNA |
| RP11-563P16.1 | -2.152 | Downregulation | 6.8186E-08 | lincRNA |
| RP11-680F20.9 | -3.771 | Downregulation | 1.2574E-17 | antisense |
| CTD-2572N17.1 | -3.239 | Downregulation | 1.0408E-09 | lincRNA |
| OVOL1-AS1 | -2.586 | Downregulation | 3.1478E-09 | antisense |
| RP11-646J21.2 | -2.541 | Downregulation | 2.2986E-06 | lincRNA |
| RP4-791M13.4 | -2.886 | Downregulation | 4.4E-06 | antisense |
| RP11-318C2.1 | -3.783 | Downregulation | 7.1792E-12 | antisense |
| RP4-541C22.5 | -2.058 | Downregulation | 3.1554E-14 | antisense |
| CTD-2210P24.3 | -2.167 | Downregulation | 3.9E-05 | antisense |
| RP11-166D19.1 | -2.220 | Downregulation | 7.5937E-14 | sense overlapping |
| RP5-916O11.2 | -2.172 | Downregulation | 2.4839E-12 | sense intronic |
| TMPRSS4-AS1 | -2.264 | Downregulation | 3.8262E-06 | antisense |
| RP11-160H12.2 | -3.449 | Downregulation | 3.7144E-12 | antisense |
| TBX5-AS1 | -2.758 | Downregulation | 1.187E-14 | antisense |
| RP11-483L5.1 | -2.166 | Downregulation | 3.705E-06 | antisense |
| RP11-736K20.5 | -2.052 | Downregulation | 1.7841E-13 | antisense |
| RP11-283I3.1 | -3.483 | Downregulation | 8.5893E-11 | antisense |
| AP000439.3 | -3.141 | Downregulation | 8.3845E-14 | lincRNA |
| RP11-711K1.7 | -2.401 | Downregulation | 6.0554E-06 | sense intronic |
| CCND2-AS2 | -2.093 | Downregulation | 2.7245E-08 | antisense |
| RP11-357K6.3 | -3.392 | Downregulation | 3.7705E-08 | lincRNA |

| | | | | |
|---|---|---|---|---|
| RP11-243M5.1 | -2.946 | Downregulation | 1.2961E-09 | lincRNA |
| RP11-434C1.2 | -2.165 | Downregulation | 0.00077345 | lincRNA |
| RP11-221N13.3 | -2.255 | Downregulation | 1.2526E-07 | lincRNA |
| RP11-277P12.10 | -3.509 | Downregulation | 7.2441E-11 | lincRNA |
| RP11-662M24.1 | -3.032 | Downregulation | 2.8603E-09 | lincRNA |
| RP11-554A11.6 | -2.001 | Downregulation | 2.6773E-07 | antisense |
| RP11-76I14.1 | -3.393 | Downregulation | 3.4698E-10 | lincRNA |
| RP11-349K16.1 | -2.784 | Downregulation | 5.7512E-07 | lincRNA |
| RP11-64D24.2 | -3.132 | Downregulation | 8.4882E-09 | antisense |
| RP11-551L14.6 | -2.123 | Downregulation | 4.9949E-07 | lincRNA |
| RP11-685B13.2 | -3.132 | Downregulation | 1.0507E-08 | sense intronic |
| RP11-266O8.1 | -2.176 | Downregulation | 1.6226E-06 | lincRNA |
| C12orf80 | -2.370 | Downregulation | 4.4086E-11 | lincRNA |
| RP11-778J16.3 | -3.090 | Downregulation | 2.4199E-09 | lincRNA |
| RP11-1028N23.4 | -4.007 | Downregulation | 3.2254E-21 | lincRNA |
| RP11-121G22.3 | -3.151 | Downregulation | 2.2685E-12 | lincRNA |
| RP11-850F7.7 | -3.268 | Downregulation | 6.6745E-09 | lincRNA |
| RP11-70F11.8 | -2.901 | Downregulation | 5.6663E-11 | lincRNA |
| RP11-630C16.1 | -3.645 | Downregulation | 2.102E-09 | lincRNA |
| RP11-596D21.1 | -3.119 | Downregulation | 9.1499E-18 | antisense |
| RP1-97G4.1 | -2.230 | Downregulation | 4.4866E-08 | lincRNA |
| NOVA1-AS1 | -2.834 | Downregulation | 2.7859E-07 | antisense |
| RP11-616L12.1 | -2.741 | Downregulation | 1.5349E-05 | lincRNA |
| RP11-256L6.3 | -2.339 | Downregulation | 3.4036E-09 | antisense |
| RP11-100F15.2 | -3.252 | Downregulation | 6.2054E-09 | lincRNA |
| RP11-148B3.1 | -3.156 | Downregulation | 8.6199E-12 | lincRNA |
| RP11-781A6.1 | -2.770 | Downregulation | 1.4136E-09 | lincRNA |
| RP11-809C9.2 | -2.493 | Downregulation | 5.5357E-11 | lincRNA |
| RP11-1022B3.1 | -2.758 | Downregulation | 6.0795E-07 | lincRNA |
| RP11-887P2.5 | -4.049 | Downregulation | 1.2913E-16 | antisense |
| RP11-386G11.3 | -2.951 | Downregulation | 1.2824E-05 | antisense |
| RP11-148B3.2 | -2.991 | Downregulation | 1.3856E-06 | lincRNA |
| CTD-2307P3.1 | -3.901 | Downregulation | 1.4446E-15 | sense overlapping |
| FRMD6-AS2 | -3.256 | Downregulation | 1.4312E-06 | lincRNA |
| CTD-2298J14.2 | -2.265 | Downregulation | 3.0038E-08 | lincRNA |
| RP11-509A17.3 | -3.376 | Downregulation | 1.0228E-09 | lincRNA |

| | | | | |
|---|---|---|---|---|
| RP11-588P7.2 | -3.057 | Downregulation | 5.2734E-06 | antisense |
| LINC00871 | -2.037 | Downregulation | 7.1921E-05 | lincRNA |
| RP11-794A8.1 | -3.324 | Downregulation | 1.2741E-08 | lincRNA |
| CTD-3049M7.1 | -3.116 | Downregulation | 1.5144E-23 | lincRNA |
| RP11-404P21.3 | -2.525 | Downregulation | 4.8932E-11 | antisense |
| RP11-1152H15.1 | -2.952 | Downregulation | 1.6342E-07 | lincRNA |
| RP11-300J18.1 | -2.742 | Downregulation | 6.1641E-08 | lincRNA |
| RP11-30G8.1 | -3.091 | Downregulation | 7.5387E-12 | lincRNA |
| RP11-463J10.4 | -3.086 | Downregulation | 4.4771E-08 | antisense |
| RP11-219E7.4 | -2.591 | Downregulation | 3.9E-10 | lincRNA |
| RP11-507K2.2 | -2.083 | Downregulation | 1.8498E-05 | antisense |
| RP11-26L16.1 | -2.610 | Downregulation | 3.2912E-05 | lincRNA |
| CTD-2058B24.2 | -2.584 | Downregulation | 9.2327E-10 | antisense |
| RP11-187O7.3 | -2.529 | Downregulation | 2.2842E-13 | lincRNA |
| LINC00648 | -3.216 | Downregulation | 4.3423E-09 | lincRNA |
| LINC00929 | -3.022 | Downregulation | 1.7578E-08 | lincRNA |
| RP6-65G23.3 | -2.110 | Downregulation | 2.1556E-15 | lincRNA |
| RP11-69H14.6 | -2.577 | Downregulation | 1.2864E-07 | sense overlapping |
| CTD-2647E9.3 | -3.035 | Downregulation | 4.6771E-09 | antisense |
| RP11-17L5.4 | -3.281 | Downregulation | 6.0729E-08 | lincRNA |
| RP11-687M24.7 | -2.781 | Downregulation | 2.5912E-07 | sense intronic |
| RP11-489D6.2 | -2.617 | Downregulation | 1.1674E-06 | antisense |
| RP11-352D13.5 | -2.508 | Downregulation | 1.4297E-08 | lincRNA |
| RP11-253M7.6 | -3.504 | Downregulation | 2.3667E-16 | sense intronic |
| CTD-2314G24.2 | -2.355 | Downregulation | 1.5705E-10 | antisense |
| CTD-2007H18.1 | -4.162 | Downregulation | 6.5637E-18 | lincRNA |
| RP11-475A13.1 | -2.016 | Downregulation | 0.00027363 | lincRNA |
| RP11-999E24.3 | -2.366 | Downregulation | 3.8115E-22 | sense overlapping |
| RP11-753D20.4 | -3.149 | Downregulation | 1.4556E-08 | lincRNA |
| LA16c-306A4.1 | -2.660 | Downregulation | 1.8798E-05 | antisense |
| RP11-616M22.5 | -2.209 | Downregulation | 4.438E-08 | sense intronic |
| RP11-481J2.2 | -3.101 | Downregulation | 1.1336E-26 | lincRNA |
| RP11-401P9.5 | -2.118 | Downregulation | 1.9173E-13 | antisense |
| RP11-101E7.2 | -2.923 | Downregulation | 2.3006E-10 | antisense |
| RP11-264B17.4 | -2.365 | Downregulation | 1.9869E-05 | antisense |
| RP11-521I2.3 | -2.637 | Downregulation | 2.8675E-25 | sense overlapping |

| | | | | |
|---|---|---|---|---|
| AC012065.7 | -2.727 | Downregulation | 1.2043E-16 | lincRNA |
| RP11-56L13.1 | -5.354 | Downregulation | 7.8882E-23 | lincRNA |
| RP11-8P11.3 | -4.710 | Downregulation | 9.1826E-23 | lincRNA |
| RP11-592N21.2 | -2.076 | Downregulation | 5.8396E-10 | sense intronic |
| RP11-24D15.1 | -3.122 | Downregulation | 1.3394E-07 | antisense |
| RP4-529N6.2 | -3.741 | Downregulation | 1.8869E-10 | lincRNA |
| RP11-18F14.1 | -2.778 | Downregulation | 8.8821E-10 | lincRNA |
| RP11-254F19.2 | -2.961 | Downregulation | 4.2637E-10 | sense intronic |
| RP11-22H5.2 | -2.866 | Downregulation | 1.1184E-22 | sense intronic |
| RP11-345M22.1 | -2.265 | Downregulation | 4.743E-16 | lincRNA |
| CTD-2012K14.4 | -2.481 | Downregulation | 5.4355E-09 | antisense |
| CTD-3247F14.2 | -4.752 | Downregulation | 6.3971E-37 | sense overlapping |
| CTD-2588J6.2 | -2.061 | Downregulation | 3.5257E-05 | lincRNA |
| RP11-7K24.3 | -2.570 | Downregulation | 9.3135E-25 | lincRNA |
| RP11-345M22.3 | -3.738 | Downregulation | 1.2265E-11 | lincRNA |
| LINC02167 | -2.455 | Downregulation | 1.0448E-05 | lincRNA |
| RP11-215E13.1 | -2.757 | Downregulation | 1.5097E-07 | lincRNA |
| RP3-507I15.2 | -2.169 | Downregulation | 3.1377E-06 | sense overlapping |
| AC140912.1 | -2.876 | Downregulation | 2.869E-09 | lincRNA |
| RP11-297L17.4 | -2.844 | Downregulation | 1.1399E-06 | lincRNA |
| RP11-308D13.3 | -2.622 | Downregulation | 9.5945E-11 | lincRNA |
| RP11-389G6.3 | -2.999 | Downregulation | 2.4121E-12 | sense overlapping |
| RP13-514E23.1 | -2.095 | Downregulation | 4.3279E-13 | sense overlapping |
| RP11-408H20.1 | -2.816 | Downregulation | 3.1822E-15 | lincRNA |
| RP11-575H3.1 | -2.761 | Downregulation | 1.491E-12 | lincRNA |
| RP11-554A11.4 | -2.085 | Downregulation | 7.066E-13 | sense overlapping |
| RP11-390D11.1 | -2.335 | Downregulation | 7.7835E-09 | antisense |
| RP11-679B19.2 | -2.571 | Downregulation | 3.0419E-09 | lincRNA |
| AC144831.1 | -2.143 | Downregulation | 2.777E-20 | lincRNA |
| RP11-483C6.1 | -2.872 | Downregulation | 7.9504E-13 | sense intronic |
| RP11-55L4.1 | -2.242 | Downregulation | 0.00083872 | antisense |
| RP11-141J13.3 | -3.600 | Downregulation | 1.0766E-17 | lincRNA |
| RP11-667K14.3 | -2.281 | Downregulation | 9.7575E-10 | lincRNA |
| RP11-333E1.2 | -3.263 | Downregulation | 2.212E-10 | lincRNA |
| RP11-204E4.3 | -4.127 | Downregulation | 1.485E-16 | lincRNA |
| LINC00675 | -2.302 | Downregulation | 4.9406E-07 | lincRNA |

| | | | | |
|---|---|---|---|---|
| CTC-304I17.4 | -2.546 | Downregulation | 0.0001269 | sense intronic |
| RP11-963H4.3 | -2.648 | Downregulation | 1.4076E-12 | lincRNA |
| RP11-269G24.3 | -2.638 | Downregulation | 2.4481E-09 | antisense |
| RP11-746B8.1 | -3.617 | Downregulation | 7.2343E-13 | antisense |
| RP11-344B2.3 | -2.173 | Downregulation | 1.8842E-11 | sense intronic |
| RP11-728E14.2 | -2.374 | Downregulation | 3.161E-06 | lincRNA |
| MIR451B | -2.309 | Downregulation | 0.00053112 | lincRNA |
| RP11-357H3.1 | -3.039 | Downregulation | 9.6551E-06 | lincRNA |
| RP11-680F20.12 | -5.011 | Downregulation | 4.7841E-25 | antisense |
| RP11-805F19.1 | -2.609 | Downregulation | 2.9379E-07 | lincRNA |
| RP11-387H17.4 | -2.137 | Downregulation | 2.0595E-10 | lincRNA |
| MIR1-2 | -2.122 | Downregulation | 2.5142E-06 | antisense |
| RP11-74H8.1 | -2.403 | Downregulation | 1.3049E-05 | antisense |
| RP11-849F2.5 | -3.457 | Downregulation | 3.7377E-23 | antisense |
| CYP4F35P | -2.241 | Downregulation | 3.0516E-08 | lincRNA |
| ESRG | -7.902 | Downregulation | 5.6184E-92 | sense intronic |
| RP11-172F10.1 | -3.677 | Downregulation | 5.347E-14 | lincRNA |
| RP11-101O21.1 | -3.226 | Downregulation | 1.5246E-11 | lincRNA |
| RP11-449D8.5 | -2.979 | Downregulation | 8.8522E-08 | lincRNA |
| CTD-2382H12.2 | -2.408 | Downregulation | 0.00069299 | lincRNA |
| RP1-56K13.5 | -2.853 | Downregulation | 5.2949E-09 | lincRNA |
| RP11-1058N17.1 | -2.044 | Downregulation | 0.00026947 | lincRNA |
| RP11-354P11.8 | -2.714 | Downregulation | 7.0746E-08 | antisense |
| CTD-2081K17.2 | -3.782 | Downregulation | 3.6142E-16 | lincRNA |
| RP11-92C4.3 | -2.789 | Downregulation | 3.8813E-09 | antisense |
| AC002398.13 | -2.548 | Downregulation | 2.1975E-06 | antisense |
| CTB-30L5.1 | -2.031 | Downregulation | 3.4363E-08 | lincRNA |
| CTB-91J4.1 | -2.018 | Downregulation | 2.5555E-05 | antisense |
| RP11-116O18.3 | -2.272 | Downregulation | 7.7722E-09 | antisense |
| AC005307.3 | -2.900 | Downregulation | 2.2738E-12 | lincRNA |
| RP11-712C7.1 | -3.600 | Downregulation | 1.7922E-12 | lincRNA |
| CTC-232P5.3 | -2.694 | Downregulation | 4.0723E-10 | antisense |
| AC005780.1 | -3.263 | Downregulation | 3.9226E-11 | lincRNA |
| KB-7G2.9 | -2.891 | Downregulation | 2.7683E-12 | lincRNA |
| LINC00906 | -4.699 | Downregulation | 2.1145E-32 | lincRNA |
| RP11-640A1.3 | -3.177 | Downregulation | 5.2737E-08 | antisense |

| | | | | |
|---|---|---|---|---|
| CTC-296K1.4 | -2.366 | Downregulation | 2.372E-08 | lincRNA |
| CTD-2189E23.2 | -3.221 | Downregulation | 2.845E-07 | lincRNA |
| RP11-120M18.5 | -2.919 | Downregulation | 1.2033E-08 | lincRNA |
| CTD-2537I9.12 | -2.432 | Downregulation | 5.0878E-08 | antisense |
| AC011524.1 | -5.580 | Downregulation | 2.4836E-34 | lincRNA |
| LINC00907 | -2.229 | Downregulation | 1.2888E-15 | lincRNA |
| CTD-2050I18.2 | -3.670 | Downregulation | 3.869E-11 | lincRNA |
| AC138472.6 | -3.085 | Downregulation | 1.9822E-07 | lincRNA |
| CTD-2553C6.1 | -2.671 | Downregulation | 3.103E-07 | antisense |
| RP11-136H19.1 | -2.260 | Downregulation | 5.9023E-05 | antisense |
| CTB-55O6.4 | -2.113 | Downregulation | 3.9527E-06 | antisense |
| AC008991.1 | -3.843 | Downregulation | 4.8797E-16 | lincRNA |
| CTB-151G24.1 | -2.869 | Downregulation | 2.4844E-07 | lincRNA |
| RP11-815J4.1 | -2.667 | Downregulation | 1.4228E-05 | lincRNA |
| CTD-2534I21.9 | -3.174 | Downregulation | 1.9464E-09 | lincRNA |
| CTD-3193O13.11 | -3.376 | Downregulation | 8.1442E-12 | lincRNA |
| CTD-2619J13.13 | -3.813 | Downregulation | 4.5422E-36 | lincRNA |
| RP11-678G14.3 | -2.423 | Downregulation | 3.4721E-07 | lincRNA |
| LINC00664 | -2.177 | Downregulation | 2.2209E-07 | lincRNA |
| CTC-518B2.12 | -3.191 | Downregulation | 3.9329E-12 | antisense |
| CTC-518B2.9 | -2.984 | Downregulation | 1.5733E-15 | antisense |
| CTC-218B8.3 | -2.553 | Downregulation | 0.00015198 | antisense |
| CTB-92J24.3 | -2.436 | Downregulation | 3.9416E-09 | antisense |
| AC092071.1 | -2.100 | Downregulation | 1.1058E-05 | sense intronic |
| CTB-92J24.2 | -2.248 | Downregulation | 1.5077E-13 | sense intronic |
| CTB-175P5.4 | -2.609 | Downregulation | 4.7653E-08 | lincRNA |
| CTD-2265M8.2 | -2.867 | Downregulation | 4.5432E-07 | antisense |
| AC004257.1 | -2.277 | Downregulation | 3.8858E-08 | lincRNA |
| AC006262.4 | -5.068 | Downregulation | 4.5062E-13 | lincRNA |
| AC006115.3 | -2.420 | Downregulation | 1.8394E-09 | antisense |
| CTB-180A7.3 | -2.383 | Downregulation | 1.2209E-09 | antisense |
| CTD-3032J10.4 | -2.167 | Downregulation | 0.00096571 | antisense |
| RP3-333A15.2 | -3.451 | Downregulation | 3.5636E-20 | sense intronic |
| RP11-635N19.3 | -2.948 | Downregulation | 5.7962E-19 | sense intronic |
| RP11-88H12.2 | -2.482 | Downregulation | 4.3174E-18 | sense intronic |
| RP11-327F22.6 | -2.197 | Downregulation | 6.0759E-17 | sense intronic |

| | | | | |
|---|---|---|---|---|
| RP11-118F19.1 | -2.525 | Downregulation | 7.0833E-10 | lincRNA |
| RP11-373D23.3 | -2.321 | Downregulation | 3.0224E-19 | lincRNA |
| RP11-2L8.2 | -2.472 | Downregulation | 0.00010969 | lincRNA |
| AC025811.3 | -2.990 | Downregulation | 1.1174E-07 | lincRNA |
| NAMA | -2.724 | Downregulation | 2.387E-11 | lincRNA |
| RP5-1182A14.5 | -2.423 | Downregulation | 3.1086E-05 | lincRNA |
| RP11-254F7.3 | -2.020 | Downregulation | 9.2584E-08 | antisense |
| CTD-2235C13.3 | -2.866 | Downregulation | 7.7397E-20 | lincRNA |
| RP11-464F9.21 | -2.470 | Downregulation | 4.2703E-10 | antisense |
| RP11-16D22.2 | -2.437 | Downregulation | 4.3316E-06 | lincRNA |
| CTD-2260A17.3 | -2.634 | Downregulation | 5.3268E-10 | antisense |
| LL22NC03-2H8.5 | -2.046 | Downregulation | 8.2256E-15 | lincRNA |
| RP5-855D21.1 | -2.470 | Downregulation | 1.5734E-07 | antisense |
| RP11-554D15.3 | -2.935 | Downregulation | 2.1252E-08 | lincRNA |
| LINC00551 | -3.809 | Downregulation | 2.3821E-22 | lincRNA |
| RP11-180N14.1 | -2.925 | Downregulation | 4.6736E-17 | lincRNA |
| GS1-166A23.1 | -2.495 | Downregulation | 3.4282E-12 | lincRNA |
| RP11-351J23.2 | -5.351 | Downregulation | 2.1407E-47 | lincRNA |
| RP11-459I19.1 | -2.008 | Downregulation | 1.9199E-08 | lincRNA |
| RP1-288H2.4 | -2.093 | Downregulation | 5.1844E-05 | sense intronic |
| RP11-572C15.6 | -2.198 | Downregulation | 1.2286E-11 | lincRNA |
| RP11-335L23.5 | -2.192 | Downregulation | 1.4311E-08 | lincRNA |
| RP11-295M18.6 | -2.127 | Downregulation | 6.8186E-08 | lincRNA |
| RP11-307C18.1 | -2.000 | Downregulation | 7.6359E-09 | antisense |
| RP11-314C9.2 | -2.907 | Downregulation | 1.9762E-10 | lincRNA |
| RP11-350J20.12 | -2.478 | Downregulation | 4.6388E-10 | antisense |
| RP11-398A8.4 | -3.544 | Downregulation | 6.8348E-11 | lincRNA |
| AC144450.2 | 2.311 | Upregulation | 5.8181E-05 | lincRNA |
| RP11-453F18__B.1 | 3.178 | Upregulation | 8.9725E-27 | lincRNA |
| AC106786.1 | 2.787 | Upregulation | 1.6887E-11 | antisense |
| AC009487.4 | 2.923 | Upregulation | 6.4172E-06 | antisense |
| HOXD-AS1 | 2.692 | Upregulation | 4.3832E-32 | antisense |
| AP000697.6 | 2.310 | Upregulation | 1.0824E-05 | antisense |
| RP11-191L9.4 | 4.414 | Upregulation | 2.714E-17 | lincRNA |
| RP11-190J1.3 | 2.481 | Upregulation | 0.00027205 | lincRNA |
| RP11-415J8.3 | 4.171 | Upregulation | 2.9505E-42 | antisense |

| | | | | |
|---|---|---|---|---|
| RFPL1S | 2.342 | Upregulation | 1.523E-11 | antisense |
| RP11-511I2.2 | 2.127 | Upregulation | 4.5139E-06 | antisense |
| AC009336.24 | 3.639 | Upregulation | 9.6577E-27 | lincRNA |
| RP11-137H2.4 | 2.296 | Upregulation | 6.6458E-16 | antisense |
| CTA-384D8.31 | 6.226 | Upregulation | 1.3943E-37 | antisense |
| AC005082.12 | 2.524 | Upregulation | 1.6704E-13 | lincRNA |
| RP3-340N1.2 | 3.150 | Upregulation | 1.4884E-10 | lincRNA |
| RP11-492E3.2 | 2.868 | Upregulation | 7.0071E-09 | antisense |
| LINC00392 | 2.748 | Upregulation | 0.00026285 | lincRNA |
| RP5-884M6.1 | 2.401 | Upregulation | 8.941E-07 | lincRNA |
| RP11-567G11.1 | 2.458 | Upregulation | 0.00030757 | lincRNA |
| RP11-73M7.1 | 2.396 | Upregulation | 2.3846E-11 | antisense |
| RP11-284F21.7 | 3.032 | Upregulation | 4.0052E-27 | antisense |
| RP11-135A1.3 | 3.437 | Upregulation | 2.2914E-07 | lincRNA |
| FEZF1-AS1 | 3.219 | Upregulation | 1.2697E-16 | antisense |
| RP13-455A7.1 | 2.182 | Upregulation | 0.00221595 | lincRNA |
| RP4-792G4.2 | 4.174 | Upregulation | 2.5447E-32 | antisense |
| AC093850.2 | 2.207 | Upregulation | 0.00026285 | lincRNA |
| AC018470.4 | 4.416 | Upregulation | 2.4606E-26 | lincRNA |
| DLG3-AS1 | 2.247 | Upregulation | 2.1726E-06 | antisense |
| PCAT7 | 3.147 | Upregulation | 6.031E-10 | antisense |
| AC073342.12 | 2.033 | Upregulation | 1.59E-14 | antisense |
| RP11-281A20.1 | 3.828 | Upregulation | 2.9986E-12 | lincRNA |
| AC104088.1 | 2.691 | Upregulation | 2.7619E-08 | lincRNA |
| LINC00884 | 2.688 | Upregulation | 7.0509E-18 | antisense |
| AP000695.4 | 2.518 | Upregulation | 1.0331E-19 | antisense |
| GS1-600G8.5 | 2.078 | Upregulation | 8.2937E-05 | lincRNA |
| MORC2-AS1 | 2.022 | Upregulation | 3.5055E-10 | antisense |
| RP11-323C15.2 | 4.848 | Upregulation | 2.4352E-28 | lincRNA |
| AC074389.9 | 2.634 | Upregulation | 1.8247E-07 | lincRNA |
| RP11-346D19.1 | 3.677 | Upregulation | 1.9524E-06 | lincRNA |
| AP001631.9 | 3.968 | Upregulation | 8.3842E-15 | antisense |
| FOXD2-AS1 | 2.294 | Upregulation | 9E-24 | antisense |
| AP000251.3 | 2.024 | Upregulation | 6.9552E-17 | antisense |
| RP5-1120P11.1 | 2.312 | Upregulation | 9.85E-17 | antisense |
| RP11-385J1.2 | 2.760 | Upregulation | 1.4745E-11 | antisense |

| | | | | |
|---|---|---|---|---|
| RP11-200A1.1 | 2.396 | Upregulation | 3.0689E-13 | lincRNA |
| CDKN2B-AS1 | 3.197 | Upregulation | 1.0602E-61 | antisense |
| MNX1-AS1 | 3.056 | Upregulation | 4.8858E-11 | antisense |
| RP11-475N22.4 | 2.066 | Upregulation | 2.2697E-20 | antisense |
| RP11-206M11.7 | 2.648 | Upregulation | 7.6509E-06 | antisense |
| AC108676.1 | 2.926 | Upregulation | 1.8738E-07 | sense overlapping |
| RP11-93K22.13 | 2.121 | Upregulation | 1.334E-12 | lincRNA |
| RP11-159F24.6 | 2.215 | Upregulation | 1.9287E-09 | antisense |
| RP11-294O2.2 | 2.108 | Upregulation | 0.00156618 | lincRNA |
| RP11-742B18.1 | 2.533 | Upregulation | 2.4081E-08 | antisense |
| CASC9 | 2.658 | Upregulation | 2.2851E-06 | lincRNA |
| RP11-98D18.3 | 2.306 | Upregulation | 2.1146E-14 | antisense |
| CTD-2116N20.1 | 2.260 | Upregulation | 8.4839E-10 | lincRNA |
| HOXC-AS2 | 2.821 | Upregulation | 1.0746E-10 | antisense |
| HOXC-AS1 | 3.021 | Upregulation | 7.0019E-08 | antisense |
| LINC00491 | 2.423 | Upregulation | 1.4359E-05 | lincRNA |
| AC084082.3 | 2.062 | Upregulation | 1.6547E-09 | lincRNA |
| RP11-434I12.3 | 3.769 | Upregulation | 1.4732E-08 | lincRNA |
| MIR146A | 2.139 | Upregulation | 6.8592E-10 | lincRNA |
| RP11-44K6.2 | 3.901 | Upregulation | 3.3396E-16 | sense intronic |
| RP11-156K13.1 | 2.165 | Upregulation | 7.0619E-14 | antisense |
| RP11-1L12.3 | 2.303 | Upregulation | 3.3682E-09 | antisense |
| RP11-326C3.2 | 3.278 | Upregulation | 9.8146E-23 | antisense |
| LINC00925 | 5.213 | Upregulation | 5.429E-112 | lincRNA |
| TMPO-AS1 | 2.143 | Upregulation | 4.1612E-42 | antisense |
| RP11-70F11.7 | 3.175 | Upregulation | 1.5962E-08 | lincRNA |
| RP11-469H8.6 | 2.366 | Upregulation | 3.4055E-06 | antisense |
| RP11-845M18.6 | 2.535 | Upregulation | 5.8111E-18 | antisense |
| RP11-903H12.3 | 3.449 | Upregulation | 6.8562E-10 | antisense |
| RP4-755D9.1 | 4.171 | Upregulation | 1.8781E-20 | antisense |
| RP11-1042B17.3 | 4.489 | Upregulation | 4.6197E-22 | antisense |
| CTD-3035D6.2 | 2.864 | Upregulation | 9.5786E-15 | lincRNA |
| RP11-356O9.2 | 2.101 | Upregulation | 8.6202E-06 | lincRNA |
| RP11-209K10.2 | 2.038 | Upregulation | 0.00214065 | lincRNA |
| RP11-20G13.1 | 2.231 | Upregulation | 1.751E-07 | lincRNA |
| RP11-519G16.5 | 2.728 | Upregulation | 2.4422E-06 | antisense |

| | | | | |
|---|---|---|---|---|
| RP11-30K9.5 | 3.583 | Upregulation | 8.7293E-15 | antisense |
| RP11-89K21.1 | 2.050 | Upregulation | 0.00020334 | lincRNA |
| RP11-69G7.1 | 3.034 | Upregulation | 1.4393E-05 | lincRNA |
| RP11-304L19.1 | 2.728 | Upregulation | 9.0414E-13 | sense overlapping |
| RP11-22P6.3 | 2.413 | Upregulation | 3.6713E-22 | antisense |
| RP11-49I11.1 | 2.097 | Upregulation | 1.0718E-13 | antisense |
| AC012531.25 | 3.279 | Upregulation | 8.1979E-17 | lincRNA |
| CTA-363E6.7 | 2.387 | Upregulation | 2.084E-07 | antisense |
| RP1-118J21.25 | 2.703 | Upregulation | 1.144E-15 | antisense |
| RP11-1260E13.1 | 2.273 | Upregulation | 0.00154863 | antisense |
| RP11-485G7.5 | 2.215 | Upregulation | 1.4279E-11 | antisense |
| AF186192.6 | 2.282 | Upregulation | 0.01090437 | lincRNA |
| SNORD3A | 2.624 | Upregulation | 1.3998E-12 | lincRNA |
| SNORD3C | 2.462 | Upregulation | 2.7197E-13 | lincRNA |
| AC145343.2 | 2.412 | Upregulation | 1.1559E-20 | lincRNA |
| RP11-149I2.4 | 3.268 | Upregulation | 7.2639E-30 | sense intronic |
| RP6-114E22.1 | 3.128 | Upregulation | 2.3204E-07 | lincRNA |
| RNF157-AS1 | 2.172 | Upregulation | 1.1971E-11 | antisense |
| MIR2117 | 2.502 | Upregulation | 8.0281E-09 | lincRNA |
| RP11-397A16.1 | 2.714 | Upregulation | 1.6276E-06 | lincRNA |
| CTB-186G2.1 | 2.733 | Upregulation | 1.4417E-06 | antisense |
| CTD-3214H19.6 | 2.171 | Upregulation | 1.2744E-20 | antisense |
| AC010524.2 | 3.004 | Upregulation | 3.2891E-11 | antisense |
| AC104534.2 | 2.916 | Upregulation | 1.0531E-06 | antisense |
| RP11-276H19.2 | 2.216 | Upregulation | 0.00049112 | lincRNA |
| TERC | 2.466 | Upregulation | 6.6801E-20 | lincRNA |
| LA16c-380H5.4 | 2.484 | Upregulation | 3.6325E-05 | lincRNA |
| RP11-109M17.2 | 3.881 | Upregulation | 3.4697E-06 | lincRNA |
| U47924.29 | 2.344 | Upregulation | 1.5499E-18 | antisense |
| RP11-573G6.8 | 2.374 | Upregulation | 4.8825E-06 | lincRNA |
| RP4-736L20.3 | 2.677 | Upregulation | 1.6521E-10 | antisense |
| RP11-337N6.1 | 2.093 | Upregulation | 0.00019238 | antisense |
| RP11-523H20.3 | 2.245 | Upregulation | 5.3944E-15 | antisense |
| RP11-284F21.10 | 2.816 | Upregulation | 9.266E-28 | antisense |
| RP1-86C11.7 | 2.893 | Upregulation | 3.8303E-23 | lincRNA |
| CTA-384D8.35 | 2.766 | Upregulation | 6.7928E-17 | lincRNA |

| | | | | |
|---|---|---|---|---|
| RP1-74M1.3 | 2.312 | Upregulation | 1.3018E-11 | lincRNA |

**N.B:** FDR corrected p value was calculated after multiple testing correction by Benjamini-Hochberg method

**<u>Supplementary Table S4</u>: Details of significantly (FDR corrected p <0.05) enriched and depleted GO Biological processes in HPV16-positive CaCx patients as compared to HPV- negative normal individuals obtained from Gene Set Enrichment Analysis (GSEA)**

| GO Biological Processes | Size of Dataset | Enrich-ment Score | Normalised Enrichment Score | Nominal p-value | FDR q-value | Rank at Max | Status |
|---|---|---|---|---|---|---|---|
| Adaptive Immune Response | 119 | 0.594 | 5.494 | <0.001 | <0.001 | 562 | Enriched |
| Phagocytosis | 88 | 0.600 | 5.190 | <0.001 | <0.001 | 462 | Enriched |
| B Cell Mediated Immunity | 84 | 0.632 | 5.174 | <0.001 | <0.001 | 462 | Enriched |
| Immune Response Regulating Signaling Pathway | 94 | 0.575 | 5.057 | <0.001 | <0.001 | 499 | Enriched |
| Lymphocyte Mediated Immunity | 89 | 0.585 | 5.042 | <0.001 | <0.001 | 462 | Enriched |
| Complement Activation | 85 | 0.587 | 4.959 | <0.001 | <0.001 | 462 | Enriched |
| Regulation of Complement Activation | 54 | 0.669 | 4.891 | <0.001 | <0.001 | 462 | Enriched |
| Fc Receptor Signaling Pathway | 58 | 0.625 | 4.779 | <0.001 | <0.001 | 462 | Enriched |
| Immunoglobulin Production | 57 | 0.658 | 4.752 | <0.001 | <0.001 | 462 | Enriched |
| Humoral Immune Response Mediated By Circulating Immunoglobulin | 81 | 0.647 | 5.460 | <0.001 | <0.001 | 462 | Enriched |
| Fc Receptor Mediated Stimulatory Signaling Pathway | 55 | 0.651 | 4.946 | <0.001 | <0.001 | 462 | Enriched |
| Adaptive Immune Response Based On Somatic Recombination of Immune Receptors Built From Immunoglobulin Superfamily Domains | 90 | 0.579 | 4.870 | <0.001 | <0.001 | 462 | Enriched |
| Activation of Immune Response | 101 | 0.529 | 4.723 | <0.001 | <0.001 | 537 | Enriched |
| Fc Epsilon Receptor Signaling Pathway | 51 | 0.643 | 4.657 | <0.001 | <0.001 | 462 | Enriched |
| B Cell Activation | 55 | 0.606 | 4.541 | <0.001 | <0.001 | 572 | Enriched |
| Regulation of B Cell Activation | 50 | 0.619 | 4.452 | <0.001 | <0.001 | 572 | Enriched |
| Positive Regulation of B Cell Activation | 46 | 0.634 | 4.434 | <0.001 | <0.001 | 572 | Enriched |
| Regulation of Humoral Immune Response | 60 | 0.584 | 4.408 | <0.001 | <0.001 | 462 | Enriched |

| | | | | | | | |
|---|---|---|---|---|---|---|---|
| B Cell Receptor Signaling Pathway | 46 | 0.634 | 4.406 | <0.001 | <0.001 | 572 | Enriched |
| Membrane Invagination | 49 | 0.609 | 4.389 | <0.001 | <0.001 | 572 | Enriched |
| Antigen Receptor Mediated Signaling Pathway | 48 | 0.635 | 4.319 | <0.001 | <0.001 | 572 | Enriched |
| Leukocyte Migration | 101 | 0.474 | 4.280 | <0.001 | <0.001 | 462 | Enriched |
| Positive Regulation of Immune Response | 116 | 0.462 | 4.249 | <0.001 | <0.001 | 572 | Enriched |
| Humoral Immune Response | 115 | 0.466 | 4.219 | <0.001 | <0.001 | 363 | Enriched |
| Phagocytosis Recognition | 46 | 0.620 | 4.179 | <0.001 | <0.001 | 443 | Enriched |
| Endocytosis | 119 | 0.448 | 4.008 | <0.001 | <0.001 | 462 | Enriched |
| Production of Molecular Mediator of Immune Response | 69 | 0.501 | 3.998 | <0.001 | <0.001 | 462 | Enriched |
| Regulation of Immune Effector Process | 76 | 0.473 | 3.895 | <0.001 | <0.001 | 464 | Enriched |
| Regulation of Immune Response | 136 | 0.398 | 3.809 | <0.001 | <0.001 | 463 | Enriched |
| Immune Effector Process | 155 | 0.382 | 3.767 | <0.001 | <0.001 | 564 | Enriched |
| Leukocyte Mediated Immunity | 123 | 0.398 | 3.691 | <0.001 | <0.001 | 537 | Enriched |
| Receptor Mediated Endocytosis | 76 | 0.449 | 3.674 | <0.001 | <0.001 | 462 | Enriched |
| Positive Regulation of Immune System Process | 148 | 0.372 | 3.610 | <0.001 | <0.001 | 464 | Enriched |
| Cell Recognition | 60 | 0.488 | 3.601 | <0.001 | <0.001 | 443 | Enriched |
| Regulation of Lymphocyte Activation | 74 | 0.417 | 3.362 | <0.001 | <0.001 | 453 | Enriched |
| Lymphocyte Activation | 80 | 0.400 | 3.327 | <0.001 | <0.001 | 572 | Enriched |
| Regulation of Immune System Process | 179 | 0.325 | 3.297 | <0.001 | <0.001 | 464 | Enriched |
| Positive Regulation of Cell Activation | 66 | 0.423 | 3.291 | <0.001 | <0.001 | 572 | Enriched |
| Regulation of Cell Activation | 81 | 0.362 | 3.022 | <0.001 | <0.001 | 453 | Enriched |
| Defense Response To Bacterium | 79 | 0.354 | 2.989 | <0.001 | <0.001 | 318 | Enriched |
| Innate Immune Response | 115 | 0.302 | 2.897 | <0.001 | <0.001 | 572 | Enriched |
| Embryonic Skeletal System Development | 15 | 0.621 | 2.702 | <0.001 | <0.001 | 345 | Enriched |
| Membrane Organization | 84 | 0.319 | 2.642 | <0.001 | <0.001 | 318 | Enriched |
| Response To Bacterium | 121 | 0.270 | 2.552 | <0.001 | 0.001 | 293 | Enriched |
| Cell Migration | 175 | 0.244 | 2.551 | <0.001 | 0.001 | 369 | Enriched |
| Chromosome Segregation | 25 | 0.478 | 2.550 | <0.001 | 0.001 | 577 | Enriched |

| | | | | | | | |
|---|---|---|---|---|---|---|---|
| Nuclear Chromosome Segregation | 22 | 0.474 | 2.401 | <0.001 | 0.003 | 560 | Enriched |
| Defense Response To Virus | 18 | 0.508 | 2.384 | <0.001 | 0.003 | 591 | Enriched |
| Sister Chromatid Segregation | 14 | 0.565 | 2.368 | 0.002 | 0.004 | 560 | Enriched |
| Embryonic Skeletal System Morphogenesis | 12 | 0.596 | 2.353 | <0.001 | 0.004 | 497 | Enriched |
| Chromosome Organization | 55 | 0.323 | 2.351 | <0.001 | 0.004 | 583 | Enriched |
| Defense Response | 195 | 0.215 | 2.301 | <0.001 | 0.005 | 564 | Enriched |
| Skeletal System Development | 50 | 0.328 | 2.284 | <0.001 | 0.006 | 354 | Enriched |
| Defense Response To Other Organism | 140 | 0.238 | 2.264 | <0.001 | 0.007 | 324 | Enriched |
| Neuron Fate Commitment | 11 | 0.584 | 2.239 | <0.001 | 0.008 | 326 | Enriched |
| Embryo Development Ending In Birth Or Egg Hatching | 48 | 0.324 | 2.225 | <0.001 | 0.008 | 615 | Enriched |
| Pattern Specification Process | 44 | 0.329 | 2.206 | <0.001 | 0.010 | 513 | Enriched |
| Mitotic Sister Chromatid Segregation | 12 | 0.550 | 2.167 | <0.001 | 0.012 | 560 | Enriched |
| Response To Biotic Stimulus | 181 | 0.216 | 2.166 | <0.001 | 0.012 | 327 | Enriched |
| Anterior Posterior Pattern Specification | 22 | 0.416 | 2.146 | 0.006 | 0.014 | 354 | Enriched |
| Regionalization | 36 | 0.329 | 2.106 | <0.001 | 0.017 | 513 | Enriched |
| Male Gamete Generation | 43 | 0.317 | 2.105 | 0.004 | 0.017 | 310 | Enriched |
| Biological Process Involved In Symbiotic Interaction | 41 | 0.310 | 2.084 | <0.001 | 0.019 | 591 | Enriched |
| Skeletal System Morphogenesis | 21 | 0.416 | 2.071 | 0.003 | 0.020 | 345 | Enriched |
| Cytokinesis | 11 | 0.544 | 2.065 | 0.002 | 0.021 | 555 | Enriched |
| Cell Activation | 126 | 0.223 | 2.065 | 0.000 | 0.021 | 353 | Enriched |
| DNA Dependent DNA Replication | 12 | 0.516 | 2.027 | 0.005 | 0.025 | 590 | Enriched |
| Cell Cycle Process | 82 | 0.242 | 2.017 | 0.004 | 0.026 | 590 | Enriched |
| Regulation of Viral Life Cycle | 11 | 0.523 | 1.984 | 0.013 | 0.031 | 591 | Enriched |
| Locomotion | 211 | 0.182 | 1.972 | <0.001 | 0.033 | 356 | Enriched |
| DNA Replication | 22 | 0.375 | 1.935 | 0.003 | 0.040 | 590 | Enriched |
| Organelle Fission | 40 | 0.297 | 1.919 | 0.007 | 0.043 | 583 | Enriched |
| DNA Recombination | 14 | 0.442 | 1.906 | 0.003 | 0.046 | 532 | Enriched |
| Embryonic Organ Development | 34 | 0.307 | 1.890 | 0.006 | 0.050 | 513 | Enriched |

| | | | | | | | |
|---|---|---|---|---|---|---|---|
| Skin Development | 101 | -0.652 | -4.601 | <0.001 | <0.001 | 188 | Depleted |
| Keratinocyte Differentiation | 89 | -0.658 | -4.487 | <0.001 | <0.001 | 188 | Depleted |
| Keratinization | 81 | -0.663 | -4.473 | <0.001 | <0.001 | 188 | Depleted |
| Epidermal Cell Differentiation | 93 | -0.638 | -4.428 | <0.001 | <0.001 | 188 | Depleted |
| Epidermis Development | 109 | -0.603 | -4.299 | <0.001 | <0.001 | 188 | Depleted |
| Epithelial Cell Differentiation | 131 | -0.504 | -3.816 | <0.001 | <0.001 | 155 | Depleted |
| Cornification | 61 | -0.577 | -3.541 | <0.001 | <0.001 | 188 | Depleted |
| Epithelium Development | 166 | -0.438 | -3.422 | <0.001 | <0.001 | 134 | Depleted |
| Peptide Cross Linking | 18 | -0.553 | -2.237 | <0.001 | 0.033 | 435 | Depleted |
| Water Homeostasis | 10 | -0.774 | -2.507 | <0.001 | 0.002 | 127 | Depleted |

**N.B**: Nominal p value and FDR q value are calculated by GSEA

<u>Supplementary Table S5</u>: **Details of the GO Biological processes corresponding to the upregulated DEcGs from STRING database**

| GO Term Description | Observed Gene Count | Background Gene Count | FDR corrected p | Matching proteins in network (labels) |
|---|---|---|---|---|
| cell cycle process | 57 | 976 | 7E-05 | CDC6, MISP, AURKA, MYBL2, GMNN, ERN2, PKMYT1, GINS1, CDKN2C, MCM2, NUF2, CCNO, M1AP, SYCE2, CDT1, C11orf85, CXCL8, CLSPN, CDC25C, E2F7, CENPA, E2F1, RBM38, UBE2C, SMC1B, SYCP2, IQGAP3, NEK2, KIF14, SHCBP1L, CENPW, ORC1, STIL, KIF2C, CDC20, KIF4A, RFC4, ECT2, MLF1, CDK1, SEPT3, MEI1, C17orf104, TDRD9, RNF212, HEPACAM2, MEIOB, CNTD2, PAX6, CDKN2A, AUNIP, NUSAP1, NUPR1L, CNTD1, STAG3, CCNI2, MUC1 |
| cytokine-mediated signaling pathway | 44 | 678 | 1E-04 | TNFRSF17, OSM, CSF3, IL17C, CD70, BST2, CXCL3, CXCL5, PF4, MMP3, CCL11, CXCL10, CXCL8, CXCL11, MMP1, NOS2, TNFRSF18, OAS2, IFI6, CCL20, CXCL9, C1orf186, AIM2, HIST2H3C, FCGR1A, IFIT3, MMP9, TRIM31, ISG15, IL2RA, RSAD2, KRT18, IL24, CXCL1, IRF7, SAA1, HIST2H3A, IL32, KRT8, ALOX15, CCL18, HIST1H3F, HIST1H3G, MUC1 |
| immune response | 76 | 1588 | 2E-04 | TNFRSF17, OSM, CD79A, CSF3, LTF, CD70, BST2, MNX1, BPIFB1, NTS, ADAMDEC1, CHRNB4, LAMP3, CXCL3, CXCL5, PF4, S100P, CCL11, CXCL10, ALCAM, CXCL8, CXCL11, OLR1, HSPA6, APOL1, FOXJ1, NOS2, APOBEC3B, CHST4, OAS2, IFI6, NKX2-3, TAP1, PIGR, CCL20, CXCL9, TRIM17, CHIT1, FCGR3A, AIM2, FCGR1A, VTCN1, HMGB3, GBP5, IFI44L, ZBP1, IFIT3, MMP9, ASS1, WFDC2, PLAU, TREM2, SMPDL3B, TRIM15, TRIM31, UBD, ISG15, IL2RA, RSAD2, LILRB4, CXCL1, IRF7, SAA1, MDK, APLN, ABCA13, SERPINA1, CFB, IGLL5, IL32, FCGR3B, ADAM8, HIST1H2BJ, IGHV3-11, IGHV3-15, CCL18 |
| pattern specification process | 32 | 432 | 4E-04 | DMRT3, DLL3, AURKA, VAX2, GSC, HOXC13, HOXC6, TCF15, SIX1, BARX1, DNAH5, HES6, OTX1, SIM2, GBX2, HOXC9, HOXC10, HOXB9, FOXJ1, FOXC2, RIPPLY3, ALX3, STIL, LHX2, PAX7, DMRTA2, HOXC4, PAX6, TDRD5, HOXB6, HOXC11, FOXD1 |
| nuclear division | 25 | 291 | 6E-04 | MISP, AURKA, MYBL2, NUF2, M1AP, SYCE2, CDT1, C11orf85, UBE2C, SYCP2, NEK2, KIF14, SHCBP1L, KIF2C, CDC20, KIF4A, CDK1, MEI1, C17orf104, TDRD9, RNF212, MEIOB, NUSAP1, CNTD1, STAG3 |
| DNA packaging | 21 | 215 | 6E-04 | CENPM, OIP5, ERN2, ASF1B, MCM2, M1AP, CENPA, HIST1H1T, CENPW, HIST2H3C, CDK1, HIST2H3A, CDKN2A, NUSAP1, HIST2H4B, HIST2H4A, HIST1H2BJ, HIST1H2BO, HIST1H4I, HIST1H3F, HIST1H3G |
| killing of cells of other organism | 14 | 91 | 6E-04 | LTF, NTS, CXCL3, PF4, CCL11, CXCL10, CXCL8, CXCL11, APOL1, CCL20, CXCL9, CXCL1, HIST1H2BJ, CCL18 |

| | | | | |
|---|---|---|---|---|
| developmental process | 197 | 5841 | 6E-04 | PRSS21, YBX2, TNFRSF17, CELSR3, DMRT3, HSD17B2, DLL3, RAB26, MMP11, OSM, SIX4, CYP24A1, AURKA, SALL4, SRMS, AMH, CD79A, MEST, CSF3, SLC44A4, GMNN, LTF, TLX2, VAX2, GSC, RDH10, HOXC13, HOXC6, TCF15, SIX1, SYNGR3, GNGT1, MNX1, C1QL1, BARX1, IGF2BP3, ALPK3, MMP7, MMP13, ITPKA, EPYC, DSG2, GINS1, CDKN2C, GRIN2D, ASF1B, EPCAM, LAMC2, SOX30, MCM2, STC2, DNAH5, MYO3A, LGR5, ZIC5, HES6, HAPLN1, GINS4, SCGB1A1, MYRF, OTX1, CCNO, BMP3, IGF2BP1, SIM2, M1AP, TFF1, GRIN2C, PTCHD2, TRIM54, RPL39L, TLX3, GBX1, BATF2, CCL11, GBX2, HOXC9, PCSK9, FOXA3, CXCL10, GPRIN1, ALCAM, CXCL8, HOXC10, TCP11, HOXB9, UGT8, NEFH, SHISA2, SYNE4, PRR15, NQO1, E2F7, FOXJ1, FOXL1, FOXC2, COL10A1, HOXD1, KRT7, RIPPLY3, ASCL2, ZPBP2, AMTN, HMX2, HIST1H1T, NKX2-3, NKX2-4, E2F1, WARS, RBM38, SYCP2, PLAC1, DMBX1, TFRC, KRT19, NEK2, C1orf186, KIF14, SHCBP1L, ROS1, PRDM13, ADRB1, ALX3, ARHGAP4, COL11A1, HMGB3, FOXD3, STIL, CEL, MMP9, ASS1, CDC20, WFDC2, COL9A2, BMP8B, TAF7L, TREM2, VSTM2L, LHX2, SPDEF, NCMAP, EPHB2, PAX7, ZIC2, TRIM15, UBD, POU4F1, CA9, EGFL6, ESM1, RSAD2, SLC34A2, DMRT2, MSLN, CSPG5, KRT18, ZNF541, PNLDC1, ECT2, MLF1, TSPAN8, SP9, SPP1, CLDN3, CDK1, CXCL1, CCK, WDR72, HNF4G, KRTAP4-1, DMRTA2, MEI1, MDK, C17orf104, TDRD9, ARTN, AGR2, APLN, HOXC4, PAX6, TDRD5, ZFP57, CDKN2A, PEG10, ONECUT2, HOXB6, MCIDAS, PROM1, KRT86, PRAME, HOXC11, KRT8, ADAM8, ALOX15, CNTD1, FOXD1, RECQL4 |
| response to cytokine | 57 | 1101 | 0.00064 | TNFRSF17, OSM, MYBL2, CSF3, IL17C, CD70, BST2, MCM2, LAMP3, SCGB1A1, CXCL3, CXCL5, PF4, MMP3, CCL11, CXCL10, CXCL8, CXCL11, NEFH, MMP1, NOS2, TNFRSF18, PTP4A3, OAS2, IFI6, CCL20, TFRC, CXCL9, C1orf186, AIM2, HIST2H3C, FCGR1A, GBP5, IFIT3, MMP9, ASS1, TRIM31, UBD, POU4F1, ISG15, IL2RA, RSAD2, KRT18, IL24, CXCL1, IRF7, SAA1, HIST2H3A, PAX6, IL32, KRT8, ALOX15, NLRP7, CCL18, HIST1H3F, HIST1H3G, MUC1 |

| | | | | |
|---|---|---|---|---|
| regulation of molecular function | 171 | 4913 | 0.00064 | DMRT3, CDC6, OSM, SIX4, AURKA, MYBL2, SALL4, AMH, CSF3, PPM1H, GMNN, LTF, TLX2, VAX2, SLC8A2, RARRES1, GSC, HOXC13, HOXC6, IL17C, CD70, TCF15, SIX1, SYNGR3, KIAA1244, BST2, GDF15, MNX1, BARX1, NTS, ERN2, GRP, GCHFR, CACNG4, PKMYT1, CDKN2C, SOX30, MCM2, STC2, LAMP3, LGR5, ZIC5, HES6, SCGB1A1, MYRF, OTX1, CCNO, BMP3, SIM2, TFF1, TMPRSS3, CXCL3, CXCL5, PF4, CAMK2N2, TLX3, GBX1, JSRP1, CDT1, BATF2, CCL11, GBX2, HOXC9, PCSK9, FOXA3, CXCL10, CXCL8, CXCL11, HOXC10, KCNS1, HOXB9, PCP2, CLSPN, NQO1, CDC25C, E2F7, FOXJ1, FOXL1, FOXC2, NOS2, HOXD1, ASCL2, HMX2, OAS2, IFI6, NKX2-3, PLEKHG7, NKX2-4, E2F1, DUSP9, WARS, UBE2C, CCL20, DMBX1, TFRC, PLEKHG4, IQGAP3, CXCL9, NEK2, KIF14, AIM2, ROS1, FCGR1A, ADRB1, ALX3, ARHGAP4, VGLL1, FOXD3, LRRC26, MMP9, CDC20, WFDC2, PLAU, BMP8B, TAF7L, TREM2, LHX2, SPDEF, EPHB2, PAX7, ZIC2, PTPRH, TRIM15, TRIM31, POU4F1, LBX2, DMRT2, CSPG5, ZNF541, IL24, RFC4, ECT2, NMB, SP9, SPP1, CLDN3, CDK1, CXCL1, CCK, HNF4G, NOXO1, LYPD1, IRF7, DMRTA2, SAA1, MDK, HMSD, ARTN, APLN, CNTD2, HOXC4, PAX6, CTCFL, SERPINA1, ARHGEF38, CDKN2A, ONECUT2, HOXB6, MCIDAS, NWD1, IL32, FAM132B, HOXC11, ADAM8, TCF24, APOC1, NLRP7, CCNI2, CCL18, FOXD1, MUC1 |
| cellular response to cytokine stimulus | 54 | 1013 | 0.00064 | TNFRSF17, OSM, MYBL2, CSF3, IL17C, CD70, BST2, MCM2, CXCL3, CXCL5, PF4, MMP3, CCL11, CXCL10, CXCL8, CXCL11, NEFH, MMP1, NOS2, TNFRSF18, PTP4A3, OAS2, IFI6, CCL20, TFRC, CXCL9, C1orf186, AIM2, HIST2H3C, FCGR1A, GBP5, IFIT3, MMP9, ASS1, TRIM31, POU4F1, ISG15, IL2RA, RSAD2, KRT18, IL24, CXCL1, IRF7, SAA1, HIST2H3A, PAX6, IL32, KRT8, ALOX15, NLRP7, CCL18, HIST1H3F, HIST1H3G, MUC1 |
| embryonic skeletal system development | 16 | 130 | 0.00068 | SIX4, GSC, RDH10, HOXC6, SIX1, HOXC9, HOXB9, FOXC2, HOXD1, ALX3, COL11A1, PAX7, DMRT2, HOXC4, HOXB6, HOXC11 |
| DNA conformation change | 26 | 328 | 0.00068 | CENPM, OIP5, GINS2, ERN2, ASF1B, MCM2, GINS4, M1AP, CENPA, HIST1H1T, CENPW, HIST2H3C, HMGB3, RFC4, CDK1, HIST2H3A, CDKN2A, NUSAP1, HIST2H4B, HIST2H4A, HIST1H2BJ, HIST1H2BO, HIST1H4I, RECQL4, HIST1H3F, HIST1H3G |
| cell cycle | 63 | 1313 | 0.0007 | CDC6, MISP, AURKA, MYBL2, OIP5, GMNN, FAM64A, ERN2, PKMYT1, GINS1, CDKN2C, MCM2, NUF2, CCNO, SKA1, M1AP, SYCE2, CDT1, C11orf85, CXCL8, CLSPN, CDC25C, E2F7, CENPA, E2F1, RBM38, UBE2C, SMC1B, SYCP2, IQGAP3, NEK2, KIF14, SHCBP1L, CENPW, ORC1, STIL, KIF2C, CDC20, KIF4A, KRT18, RFC4, ECT2, MLF1, CDK1, SEPT3, MEI1, C17orf104, TDRD9, RNF212, HEPACAM2, MEIOB, CNTD2, PAX6, CTCFL, CDKN2A, MCIDAS, AUNIP, NUSAP1, NUPR1L, CNTD1, STAG3, CCNI2, MUC1 |

| | | | | |
|---|---|---|---|---|
| immune system process | 100 | 2481 | 0.00071 | TNFRSF17, OSM, SIX4, CD79A, CSF3, LTF, CD70, SIX1, BST2, MNX1, BPIFB1, BARX1, NTS, ADAMDEC1, CHRNB4, EPCAM, LAMP3, SLFN13, CXCL3, CXCL5, PF4, S100P, BATF2, CCL11, CXCL10, ALCAM, CXCL8, CXCL11, OLR1, HSPA6, APOL1, SLC16A8, MMP1, FOXJ1, FOXL1, NOS2, APOBEC3B, CHST4, OAS2, IFI6, NKX2-3, PODXL2, TAP1, PIGR, CCL20, TFRC, CXCL9, TRIM17, C1orf186, CHIT1, FCGR3A, AIM2, FCGR1A, VTCN1, HMGB3, GBP5, IFI44L, ZBP1, IFIT3, KIF2C, MMP9, ASS1, WFDC2, PLAU, TREM2, SMPDL3B, KIF4A, TRIM15, TRIM31, UBD, ISG15, IL2RA, RSAD2, LILRB4, MLF1, CXCL1, IRF7, MUC16, SAA1, MDK, HMSD, ARTN, APLN, ABCA13, SERPINA1, CFB, HOXB6, IDO1, IGLL5, IL32, FCGR3B, ADAM8, SLC16A3, HIST1H2BJ, IGHV3-11, IGHV3-15, CCL18, HSH2D, MUC1, MUC5AC |
| regionalization | 26 | 332 | 0.00071 | DMRT3, DLL3, AURKA, VAX2, GSC, HOXC13, HOXC6, TCF15, BARX1, HES6, OTX1, GBX2, HOXC9, HOXC10, HOXB9, FOXJ1, FOXC2, LHX2, PAX7, DMRTA2, HOXC4, PAX6, TDRD5, HOXB6, HOXC11, FOXD1 |
| chromatin assembly or disassembly | 19 | 193 | 0.00071 | CENPM, OIP5, ASF1B, MCM2, M1AP, CENPA, HIST1H1T, HIST3H2A, CENPW, HIST2H3C, HIST2H3A, CDKN2A, HIST2H4B, HIST2H4A, HIST1H2BJ, HIST1H2BO, HIST1H4I, HIST1H3F, HIST1H3G |
| nucleosome assembly | 16 | 135 | 0.00071 | CENPM, OIP5, ASF1B, MCM2, CENPA, HIST1H1T, CENPW, HIST2H3C, HIST2H3A, HIST2H4B, HIST2H4A, HIST1H2BJ, HIST1H2BO, HIST1H4I, HIST1H3F, HIST1H3G |
| defense response | 62 | 1296 | 0.00071 | LTF, IL17C, BST2, BPIFB1, SLFN13, CXCL3, CXCL5, PF4, BATF2, CCL11, CXCL10, CXCL8, CXCL11, OLR1, APOL1, NOS2, APOBEC3B, CHST4, OAS2, IFI6, TAP1, C2CD4A, CCL20, CXCL9, TRIM17, AIM2, FCGR1A, HMGB3, GBP5, IFI44L, ZBP1, IFIT3, ASS1, WFDC2, TREM2, SMPDL3B, TRIM15, TRIM31, UBD, ISG15, IL2RA, C2CD4B, RSAD2, SPP1, CXCL1, LYPD1, IRF7, CYP4F11, SAA1, MDK, SERPINA1, CFB, TNIP3, IDO1, IGLL5, IL32, ADAM8, ALOX15, HIST1H2BJ, IGHV3-11, IGHV3-15, CCL18 |
| humoral immune response | 23 | 275 | 0.00071 | LTF, MNX1, BPIFB1, NTS, CXCL3, CXCL5, PF4, CCL11, CXCL10, CXCL8, CXCL11, FOXJ1, CCL20, CXCL9, WFDC2, TREM2, CXCL1, CFB, IGLL5, HIST1H2BJ, IGHV3-11, IGHV3-15, CCL18 |
| embryo development | 52 | 1002 | 0.00071 | HSD17B2, DLL3, SIX4, SALL4, SLC44A4, TLX2, VAX2, GSC, RDH10, HOXC6, TCF15, SIX1, GINS1, ASF1B, MYO3A, GINS4, OTX1, SIM2, GBX2, HOXC9, CXCL8, HOXC10, HOXB9, E2F7, FOXC2, HOXD1, RIPPLY3, ASCL2, HMX2, KRT19, NEK2, ALX3, COL11A1, FOXD3, STIL, MMP9, LHX2, EPHB2, PAX7, TRIM15, SLC34A2, DMRT2, PNLDC1, SP9, APLN, HOXC4, PAX6, TDRD5, ZFP57, HOXB6, HOXC11, KRT8 |
| neutrophil chemotaxis | 12 | 74 | 0.00071 | CXCL3, CXCL5, PF4, CCL11, CXCL10, CXCL8, CXCL11, CCL20, CXCL9, CXCL1, SAA1, CCL18 |
| chromatin assembly | 18 | 172 | 0.00071 | CENPM, OIP5, ASF1B, MCM2, M1AP, CENPA, HIST1H1T, CENPW, HIST2H3C, HIST2H3A, CDKN2A, HIST2H4B, HIST2H4A, HIST1H2BJ, HIST1H2BO, HIST1H4I, HIST1H3F, HIST1H3G |

| | | | | |
|---|---|---|---|---|
| multicellular organismal process | 223 | 6933 | 0.00071 | PRSS21, YBX2, TNFRSF17, CELSR3, DMRT3, MOGAT2, HSD17B2, DLL3, RAB26, MMP11, OSM, SIX4, CYP24A1, AURKA, SALL4, AMH, CD79A, CSF3, SLC44A4, GMNN, LTF, TLX2, VAX2, SLC8A2, GSC, RDH10, HOXC13, HOXC6, OR2B6, TCF15, SIX1, SYNGR3, GNGT1, GDF15, MNX1, SLC6A8, C1QL1, BARX1, IGF2BP3, ALPK3, MMP7, MMP13, ITPKA, EPYC, DSG2, CHRNB4, CACNG4, GINS1, CDKN2C, GRIN2D, ASF1B, EPCAM, LAMC2, SOX30, MCM2, STC2, DNAH5, MYO3A, LGR5, ZIC5, HES6, HAPLN1, PLA2G7, GINS4, SCGB1A1, MYRF, OTX1, BMP3, IGF2BP1, SIM2, M1AP, TFF1, TMPRSS3, GRIN2C, PTCHD2, PF4, TRIM54, RPL39L, S100P, TLX3, GBX1, JSRP1, BATF2, CCL11, GBX2, HOXC9, PCSK9, FOXA3, CXCL10, GPRIN1, ALCAM, CXCL8, HOXC10, TCP11, OLR1, HOXB9, UGT8, NEFH, SHISA2, PRR15, E2F7, FOXJ1, FOXL1, FOXC2, NOS2, COL10A1, HOXD1, KRT7, RIPPLY3, PTP4A3, ASCL2, ZPBP2, AMTN, HMX2, HIST1H1T, NKX2-3, NKX2-4, E2F1, WARS, C2CD4A, PIGR, SYCP2, PLAC1, DMBX1, TFRC, KRT19, NEK2, C1orf186, CHIT1, KIF14, SHCBP1L, AIM2, ROS1, HIST2H3C, PRDM13, ADRB1,<br><br>VTCN1, ALX3, ARHGAP4, COL11A1, HMGB3, GBP5, FOXD3, STIL, CEL, MMP9, ASS1, CDC20, WFDC2, COL9A2, PLAU, BMP8B, TAF7L, TREM2, VSTM2L, LHX2, SPDEF, NCMAP, EPHB2, PAX7, ZIC2, TRIM15, UBD, POU4F1, ISG15, C2CD4B, EGFL6, ESM1, TNNI2, RSAD2, SLC34A2, DMRT2, MSLN, CSPG5, KRT18, ZNF541, PNLDC1, ECT2, MLF1, TSPAN8, SP9, SPP1, CLDN3, CDK1, CXCL1, CCK, WDR72, LYPD1, IRF7, KRTAP4-1, DMRTA2, MEI1, CYP4F11, SAA1, MDK, HIST2H3A, C17orf104, TDRD9, ARTN, MEIOB, AGR2, APLN, HOXC4, PAX6, TDRD5, CTCFL, SERPINA1, ZFP57, ONECUT2, HOXB6, PROM1, IDO1, KRT86, HOXC11, KRT8, ADAM8, ALOX15, CNTD1, APOC1, FOXD1, RECQL4, HIST1H3F, HIST1H3G |
| leukocyte migration | 25 | 316 | 0.00071 | EPCAM, CXCL3, CXCL5, PF4, CCL11, CXCL10, CXCL8, CXCL11, OLR1, SLC16A8, MMP1, FOXJ1, NKX2-3, PODXL2, CCL20, CXCL9, CXCL1, SAA1, MDK, ARTN, IGLL5, ADAM8, SLC16A3, IGHV3-11, CCL18 |
| chromosome organization | 54 | 1066 | 0.00071 | CENPM, AURKA, OIP5, GINS2, ERN2, ASF1B, MCM2, NUF2, GINS4, ATAD2, M1AP, SYCE2, CDT1, C11orf85, FOXA3, RMI2, CENPA, HIST1H1T, SMC1B, SYCP2, HIST3H2A, NEK2, KIF14, CENPW, HIST2H3C, HIST2H2AA, PRDM13, HMGB3, KIF2C, CDC20, KIF4A, PAX7, PADI3, RFC4, CDK1, MEI1, HIST2H3A, C17orf104, RNF212, MEIOB, CTCFL, CDKN2A, NUSAP1, HIST2H4B, HIST2H4A, HIST2H2AA3, HIST1H2BJ, HIST1H2BO, STAG3, HIST1H4I, RECQL4, HIST1H2AB, HIST1H3F, HIST1H3G |
| antimicrobial humoral immune response mediated by antimicrobial peptide | 14 | 113 | 0.0014 | LTF, NTS, CXCL3, CXCL5, PF4, CCL11, CXCL10, CXCL8, CXCL11, CCL20, CXCL9, CXCL1, HIST1H2BJ, CCL18 |

| | | | | |
|---|---|---|---|---|
| anatomical structure development | 180 | 5402 | 0.0015 | YBX2, TNFRSF17, CELSR3, DMRT3, HSD17B2, DLL3, RAB26, MMP11, OSM, SIX4, AURKA, SALL4, AMH, CD79A, MEST, CSF3, SLC44A4, GMNN, LTF, TLX2, VAX2, GSC, RDH10, HOXC13, HOXC6, TCF15, SIX1, SYNGR3, GNGT1, MNX1, C1QL1, BARX1, IGF2BP3, ALPK3, MMP13, ITPKA, EPYC, DSG2, GINS1, CDKN2C, GRIN2D, ASF1B, EPCAM, LAMC2, SOX30, MCM2, STC2, DNAH5, MYO3A, LGR5, ZIC5, HES6, HAPLN1, GINS4, SCGB1A1, MYRF, OTX1, CCNO, BMP3, IGF2BP1, SIM2, GRIN2C, PTCHD2, TRIM54, TLX3, GBX1, BATF2, CCL11, GBX2, HOXC9, PCSK9, FOXA3, CXCL10, GPRIN1, ALCAM, CXCL8, HOXC10, TCP11, HOXB9, UGT8, NEFH, SHISA2, SYNE4, PRR15, E2F7, FOXJ1, FOXL1, FOXC2, COL10A1, HOXD1, KRT7, RIPPLY3, ASCL2, ZPBP2, AMTN, HMX2, HIST1H1T, NKX2-3, NKX2-4, E2F1, WARS, SYCP2, PLAC1, DMBX1, TFRC, KRT19, NEK2, C1orf186, KIF14, ROS1, PRDM13, ALX3, ARHGAP4, COL11A1, HMGB3, FOXD3, STIL, CEL, MMP9, ASS1, CDC20, COL9A2, BMP8B, TAF7L, TREM2, VSTM2L, LHX2, SPDEF, NCMAP, EPHB2, PAX7, ZIC2, TRIM15, UBD, POU4F1, CA9, EGFL6, ESM1, RSAD2, SLC34A2, DMRT2, MSLN, CSPG5, KRT18, ZNF541, PNLDC1, ECT2, MLF1, SP9, SPP1, CLDN3, CDK1, CXCL1, CCK, WDR72, HNF4G, KRTAP4-1, DMRTA2, MEI1, MDK, C17orf104, TDRD9, ARTN, AGR2, APLN, HOXC4, PAX6, TDRD5, ZFP57, ONECUT2, HOXB6, MCIDAS, PROM1, KRT86, HOXC11, KRT8, ADAM8, ALOX15, FOXD1, RECQL4 |
| inflammatory response | 32 | 515 | 0.0019 | IL17C, CXCL3, CXCL5, PF4, CCL11, CXCL10, CXCL8, CXCL11, OLR1, NOS2, CHST4, C2CD4A, CCL20, CXCL9, AIM2, GBP5, ASS1, TREM2, SMPDL3B, IL2RA, C2CD4B, SPP1, CXCL1, CYP4F11, SAA1, MDK, SERPINA1, TNIP3, IDO1, ADAM8, ALOX15, CCL18 |

| | | | | |
|---|---|---|---|---|
| multicellular organism development | 169 | 5023 | 0.0019 | TNFRSF17, CELSR3, DMRT3, HSD17B2, DLL3, RAB26, MMP11, OSM, SIX4, AURKA, SALL4, AMH, CD79A, CSF3, SLC44A4, GMNN, LTF, TLX2, VAX2, GSC, RDH10, HOXC13, HOXC6, TCF15, SIX1, SYNGR3, GNGT1, MNX1, C1QL1, BARX1, IGF2BP3, ALPK3, MMP13, ITPKA, EPYC, DSG2, GINS1, CDKN2C, GRIN2D, ASF1B, EPCAM, LAMC2, MCM2, STC2, DNAH5, MYO3A, LGR5, ZIC5, HES6, HAPLN1, GINS4, SCGB1A1, MYRF, OTX1, BMP3, IGF2BP1, SIM2, GRIN2C, PTCHD2, TRIM54, TLX3, GBX1, BATF2, CCL11, GBX2, HOXC9, PCSK9, FOXA3, CXCL10, GPRIN1, ALCAM, CXCL8, HOXC10, TCP11, HOXB9, UGT8, NEFH, SHISA2, PRR15, E2F7, FOXJ1, FOXL1, FOXC2, COL10A1, HOXD1, KRT7, RIPPLY3, ASCL2, AMTN, HMX2, HIST1H1T, NKX2-3, NKX2-4, E2F1, WARS, SYCP2, PLAC1, DMBX1, TFRC, KRT19, NEK2, C1orf186, KIF14, ROS1, PRDM13, ALX3, ARHGAP4, COL11A1, HMGB3, FOXD3, STIL, CEL, MMP9, ASS1, CDC20, COL9A2, BMP8B, TAF7L, TREM2, VSTM2L, LHX2, SPDEF, NCMAP, EPHB2, PAX7, ZIC2, TRIM15, UBD, POU4F1, EGFL6, ESM1, RSAD2, SLC34A2, DMRT2, MSLN, CSPG5, KRT18, ZNF541, PNLDC1, ECT2, MLF1, SP9, SPP1, CLDN3, CDK1, CXCL1, CCK, WDR72, KRTAP4-1, DMRTA2, MDK, TDRD9, ARTN, AGR2, APLN, HOXC4, PAX6, TDRD5, ZFP57, ONECUT2, HOXB6, PROM1, KRT86, HOXC11, KRT8, ADAM8, ALOX15, FOXD1, RECQL4 |
| protein-DNA complex assembly | 19 | 213 | 0.0019 | CENPM, OIP5, GMNN, ASF1B, MCM2, CDT1, CENPA, HIST1H1T, CENPW, HIST2H3C, TAF7L, HIST2H3A, HIST2H4B, HIST2H4A, HIST1H2BJ, HIST1H2BO, HIST1H4I, HIST1H3F, HIST1H3G |
| nucleosome organization | 17 | 176 | 0.0022 | CENPM, OIP5, ASF1B, MCM2, CENPA, HIST1H1T, HIST3H2A, CENPW, HIST2H3C, HIST2H3A, HIST2H4B, HIST2H4A, HIST1H2BJ, HIST1H2BO, HIST1H4I, HIST1H3F, HIST1H3G |
| neuron fate specification | 8 | 33 | 0.0024 | DMRT3, SIX1, MNX1, TLX3, HOXC10, POU4F1, DMRTA2, PAX6 |
| antimicrobial humoral response | 16 | 160 | 0.0026 | LTF, BPIFB1, NTS, CXCL3, CXCL5, PF4, CCL11, CXCL10, CXCL8, CXCL11, CCL20, CXCL9, WFDC2, CXCL1, HIST1H2BJ, CCL18 |

| | | | | |
|---|---|---|---|---|
| response to stress | 125 | 3485 | 0.0031 | AURKA, RNASEH2A, RAD51AP1, LTF, SLC8A2, IL17C, GNGT1, BST2, BPIFB1, GINS2, ERN2, MCM2, STC2, GINS4, SCGB1A1, PLOD2, SLFN13, IGF2BP1, TFF1, GRIN2C, CXCL3, CXCL5, PF4, MMP3, MARVELD3, BATF2, CCL11, PCSK9, FOXA3, CXCL10, CXCL8, CXCL11, OLR1, HSPA6, RMI2, CLSPN, APOL1, NQO1, CDC25C, E2F7, CLGN, NOS2, APOBEC3B, ASCL2, CHST4, OAS2, IFI6, E2F1, DUSP9, TAP1, C2CD4A, RBM38, CCL20, CXCL9, HIST3H2A, TRIM17, DTL, UBE2T, AIM2, HIST2H3C, FCGR1A, ADRB1, KIAA1324, HMGB3, GBP5, IFI44L, ZBP1, IFIT3, MMP9, ASS1, WFDC2, PLAU, TREM2, SMPDL3B, TRIM15, TRIM31, UBD, CA9, ISG15, IL2RA, C2CD4B, RSAD2, IL24, RFC4, ECT2, SPP1, CLDN3, CDK1, CXCL1, LYPD1, IRF7, CBSL, CYP4F11, DERL3, SAA1, MDK, HIST2H3A, C17orf104, MEIOB, PAX6, SERPINA1, CFB, CDKN2A, RNF183, TNIP3, IDO1, IGLL5, IL32, AUNIP, KRT8, ADAM8, NUPR1L, ALOX15, HIST2H4B, HIST2H4A, HIST1H2BJ, TMPRSS4, IGHV3-11, IGHV3-15, CCL18, HIST1H4I, RECQL4, HIST1H3F, HIST1H3G, MUC1 |
| chromosome segregation | 21 | 268 | 0.0032 | OIP5, NUF2, SKA1, M1AP, SYCE2, CDT1, C11orf85, SMC1B, SYCP2, NEK2, KIF14, CENPW, KIF2C, CDC20, KIF4A, MEI1, C17orf104, RNF212, MEIOB, NUSAP1, STAG3 |
| chordate embryonic development | 36 | 640 | 0.0032 | HSD17B2, DLL3, SIX4, SALL4, GSC, RDH10, HOXC6, TCF15, SIX1, GINS1, ASF1B, GINS4, GBX2, HOXC9, HOXB9, E2F7, FOXC2, HOXD1, RIPPLY3, ASCL2, KRT19, NEK2, ALX3, COL11A1, FOXD3, STIL, LHX2, PAX7, SLC34A2, DMRT2, PNLDC1, HOXC4, PAX6, HOXB6, HOXC11, KRT8 |
| mitotic cell cycle process | 35 | 616 | 0.0034 | CDC6, MISP, AURKA, MYBL2, GMNN, PKMYT1, GINS1, CDKN2C, MCM2, NUF2, CCNO, CDT1, CLSPN, CDC25C, E2F7, CENPA, E2F1, RBM38, UBE2C, IQGAP3, NEK2, KIF14, ORC1, STIL, KIF2C, CDC20, KIF4A, ECT2, CDK1, CNTD2, PAX6, CDKN2A, NUSAP1, CCNI2, MUC1 |
| nuclear chromosome segregation | 18 | 209 | 0.004 | NUF2, M1AP, SYCE2, CDT1, C11orf85, SMC1B, SYCP2, NEK2, KIF14, KIF2C, CDC20, KIF4A, MEI1, C17orf104, RNF212, MEIOB, NUSAP1, STAG3 |
| response to biotic stimulus | 58 | 1289 | 0.0042 | CSF3, LTF, BST2, BPIFB1, NTS, CMPK2, GRP, SCGB1A1, SLFN13, CXCL3, CXCL5, PF4, BATF2, CCL11, CXCL10, CXCL8, CXCL11, APOL1, NOS2, APOBEC3B, OAS2, IFI6, CCL20, CXCL9, TRIM17, CHIT1, AIM2, FCGR1A, VTCN1, HMGB3, GBP5, IFI44L, ZBP1, IFIT3, ASS1, WFDC2, TREM2, SMPDL3B, TRIM15, TRIM31, UBD, IGFBPL1, ISG15, RSAD2, IL24, CXCL1, IRF7, SAA1, CFB, TNIP3, IDO1, IGLL5, KRT8, NLRP7, HIST1H2BJ, IGHV3-11, IGHV3-15, CCL18 |

| | | | | |
|---|---|---|---|---|
| response to external biotic stimulus | 57 | 1258 | 0.0042 | CSF3, LTF, BST2, BPIFB1, NTS, CMPK2, GRP, SCGB1A1, SLFN13, CXCL3, CXCL5, PF4, BATF2, CCL11, CXCL10, CXCL8, CXCL11, APOL1, NOS2, APOBEC3B, OAS2, IFI6, CCL20, CXCL9, TRIM17, CHIT1, AIM2, FCGR1A, VTCN1, HMGB3, GBP5, IFI44L, ZBP1, IFIT3, ASS1, WFDC2, TREM2, SMPDL3B, TRIM15, TRIM31, UBD, ISG15, RSAD2, IL24, CXCL1, IRF7, SAA1, CFB, TNIP3, IDO1, IGLL5, KRT8, NLRP7, HIST1H2BJ, IGHV3-11, IGHV3-15, CCL18 |
| chemokine-mediated signaling pathway | 11 | 80 | 0.0042 | CXCL3, CXCL5, PF4, CCL11, CXCL10, CXCL8, CXCL11, CCL20, CXCL9, CXCL1, CCL18 |
| embryonic skeletal system morphogenesis | 12 | 97 | 0.0043 | SIX4, GSC, RDH10, SIX1, HOXC9, HOXB9, FOXC2, ALX3, COL11A1, HOXC4, HOXB6, HOXC11 |
| protein-DNA complex subunit organization | 20 | 255 | 0.0043 | CENPM, OIP5, GMNN, ASF1B, MCM2, CDT1, CENPA, HIST1H1T, HIST3H2A, CENPW, HIST2H3C, TAF7L, HIST2H3A, HIST2H4B, HIST2H4A, HIST1H2BJ, HIST1H2BO, HIST1H4I, HIST1H3F, HIST1H3G |
| anterior/posterior pattern specification | 18 | 214 | 0.0047 | DLL3, AURKA, HOXC13, HOXC6, TCF15, BARX1, HES6, OTX1, GBX2, HOXC9, HOXC10, HOXB9, FOXC2, HOXC4, PAX6, TDRD5, HOXB6, HOXC11 |
| extracellular matrix disassembly | 10 | 66 | 0.0047 | MMP11, MMP7, MMP13, MMP10, MMP3, MMP1, MMP9, WDR72, NOXO1, ADAM8 |
| neuron fate commitment | 10 | 66 | 0.0047 | DMRT3, SIX1, MNX1, TLX3, GBX1, HOXC10, PAX7, POU4F1, DMRTA2, PAX6 |
| mitotic cell cycle | 37 | 695 | 0.0058 | CDC6, MISP, AURKA, MYBL2, GMNN, PKMYT1, GINS1, CDKN2C, MCM2, NUF2, CCNO, SKA1, CDT1, CLSPN, CDC25C, E2F7, CENPA, E2F1, RBM38, UBE2C, IQGAP3, NEK2, KIF14, CENPW, ORC1, STIL, KIF2C, CDC20, KIF4A, ECT2, CDK1, CNTD2, PAX6, CDKN2A, NUSAP1, CCNI2, MUC1 |

| | | | | |
|---|---|---|---|---|
| negative regulation of biological process | 175 | 5389 | 0.006 | DLL3, CDC6, MMP11, OSM, SIX4, AURKA, SALL4, SRMS, AMH, CSF3, GMNN, LTF, TLX2, VAX2, RARRES1, GSC, HOXC6, SIX1, KIAA1244, BST2, GDF15, BPIFB1, BARX1, ADAMDEC1, ERN2, IGF2BP3, GCHFR, PKMYT1, CDKN2C, EPCAM, SOX30, MCM2, STC2, LAMP3, HES6, SCGB1A1, ATAD2, TMSNB, IGF2BP1, SIM2, TFF1, GRIN2C, PTCHD2, PF4, TRIM54, CAMK2N2, MFI2, TLX3, MMP3, MARVELD3, B4GALNT2, CDT1, CCL11, PCSK9, CXCL10, CXCL8, HOXC10, AGR3, RMI2, CLSPN, SHISA2, NQO1, CDC25C, E2F7, FOXJ1, FOXC2, NOS2, APOBEC3B, TNFRSF18, RIPPLY3, ASCL2, ZPBP2, HIST1H1T, OAS2, IFI6, E2F1, DUSP9, WARS, RBM38, SYCP2, DMBX1, TFRC, IQGAP3, HIST3H2A, DTL, NEK2, KIF14, AIM2, ROS1, HIST2H3C, HIST2H2AA, PRDM13, ADRB1, VTCN1, ARHGAP4, HENMT1, HMGB3, FOXD3, ORC1, IFIT3, STIL, MMP9, ASS1, CDC20, WFDC2, PLAU, TREM2, TSPAN15, VSTM2L, LHX2, SMPDL3B, SPDEF, EPHB2, PAX7, ZIC2, PTPRH, TRIM15, TRIM31, POU4F1, ISG15, IL2RA, RSAD2, KRT18, LILRB4, GNG4, IL24, PNLDC1, MLF1, TSPAN8, NMB, SPP1, CLDN3, CDK1, CXCL1, CCK, LYPD1, IRF7, DERL3, SAA1, MDK, HIST2H3A, HMSD, C17orf104, TDRD9, AGR2, APLN, PAX6, SERPINA1, CDKN2A, PEG10, ONECUT2, TNIP3, NWD1, IDO1, IL32, FAM132B, AUNIP, PRAME, ADAM8, NUPR1L, ALOX15, HIST2H4B, HIST2H4A, APOC1, NLRP7, HIST2H2AA3, HIST1H2BJ, TMPRSS4, HIST1H4I, FOXD1, HSH2D, HIST1H2AB, HIST1H3F, HIST1H3G, MUC1 |
| response to other organism | 56 | 1256 | 0.0062 | CSF3, LTF, BST2, BPIFB1, NTS, CMPK2, SCGB1A1, SLFN13, CXCL3, CXCL5, PF4, BATF2, CCL11, CXCL10, CXCL8, CXCL11, APOL1, NOS2, APOBEC3B, OAS2, IFI6, CCL20, CXCL9, TRIM17, CHIT1, AIM2, FCGR1A, VTCN1, HMGB3, GBP5, IFI44L, ZBP1, IFIT3, ASS1, WFDC2, TREM2, SMPDL3B, TRIM15, TRIM31, UBD, ISG15, RSAD2, IL24, CXCL1, IRF7, SAA1, CFB, TNIP3, IDO1, IGLL5, KRT8, NLRP7, HIST1H2BJ, IGHV3-11, IGHV3-15, CCL18 |
| negative regulation of gene expression, epigenetic | 12 | 103 | 0.0064 | HIST3H2A, HIST2H3C, HIST2H2AA, LHX2, HIST2H3A, HIST2H4B, HIST2H4A, HIST2H2AA3, HIST1H4I, HIST1H2AB, HIST1H3F, HIST1H3G |
| leukocyte chemotaxis | 14 | 142 | 0.0075 | CXCL3, CXCL5, PF4, CCL11, CXCL10, CXCL8, CXCL11, CCL20, CXCL9, CXCL1, SAA1, MDK, ADAM8, CCL18 |
| response to external stimulus | 88 | 2310 | 0.0086 | CELSR3, CYP24A1, CSF3, LTF, GNGT1, BST2, GDF15, BPIFB1, NTS, CMPK2, GRP, MMP7, GRIN2D, LAMC2, STC2, SCGB1A1, SLFN13, CXCL3, CXCL5, PF4, GBX1, BATF2, CCL11, GBX2, PCSK9, FOXA3, CXCL10, ALCAM, CXCL8, CXCL11, HOXB9, APOL1, NQO1, NOS2, APOBEC3B, OAS2, IFI6, CCL20, CXCL9, TRIM17, CHIT1, AIM2, FCGR1A, ADRB1, VTCN1, KIAA1324, COL11A1, HMGB3, GBP5, IFI44L, ZBP1, IFIT3, ASS1, WFDC2, PLAU, TREM2, VSTM2L, LHX2, SMPDL3B, EPHB2, TRIM15, TRIM31, UBD, ISG15, RSAD2, IL24, SPP1, CXCL1, CCK, IRF7, CBSL, SAA1, MDK, ARTN, PAX6, CFB, TNIP3, IDO1, IGLL5, KRT8, ADAM8, NUPR1L, NLRP7, HIST1H2BJ, IGHV3-11, IGHV3-15, CCL18, FOXD1 |

| | | | | |
|---|---|---|---|---|
| chromatin organization involved in negative regulation of transcription | 12 | 108 | 0.0091 | HIST3H2A, HIST2H3C, HIST2H2AA, HIST2H3A, CDKN2A, HIST2H4B, HIST2H4A, HIST2H2AA3, HIST1H4I, HIST1H2AB, HIST1H3F, HIST1H3G |
| regulation of metabolic process | 214 | 6948 | 0.0103 | YBX2, CELSR3, DMRT3, CDC6, RAB26, OSM, SIX4, AURKA, MYBL2, SALL4, AMH, CSF3, PPM1H, RAD51AP1, GMNN, LTF, TLX2, VAX2, SLC8A2, RARRES1, GSC, RDH10, HOXC13, HOXC6, TCF15, SIX1, KIAA1244, BST2, GDF15, MNX1, BARX1, NUP210, ERN2, IGF2BP3, GCHFR, PKMYT1, CDKN2C, EPCAM, SOX30, STC2, LAMP3, ZIC5, HES6, SCGB1A1, MYRF, OTX1, CCNO, BMP3, ATAD2, IGF2BP1, SIM2, GRIN2C, PTCHD2, PF4, TRIM54, CAMK2N2, MFI2, TLX3, GBX1, MMP3, MARVELD3, CDT1, BATF2, CCL11, GBX2, HOXC9, PCSK9, MZB1, FOXA3, CXCL10, HOXC10, HOXB9, RMI2, SPSB4, CLSPN, NQO1, CDC25C, E2F7, FOXJ1, FOXL1, FOXC2, NOS2, TNFRSF18, HOXD1, RIPPLY3, PTP4A3, ASCL2, HMX2, HIST1H1T, ZNF695, OAS2, IFI6, NKX2-3, NKX2-4, E2F1, DUSP9, WARS, RBM38, UBE2C, CCL20, DMBX1, TFRC, IQGAP3, HIST3H2A, TRIM17, DTL, NEK2, G0S2, KIF14, AIM2, ROS1, HIST2H3C, HIST2H2AA, FCGR1A, PRDM13, ADRB1,<br><br>VTCN1, ALX3, KIAA1324, ELOVL3, HENMT1, HMGB3, GBP5, VGLL1, FOXD3, ZBP1, ORC1, MMP9, ASS1, CDC20, WFDC2, BMP8B, TAF7L, TREM2, LHX2, SPDEF, EPHB2, PAX7, ZIC2, PTPRH, TRIM15, TRIM31, POU4F1, LBX2, CA9, ISG15, TNNI2, RSAD2, DMRT2, GOLM1, LILRB4, ZNF541, IL24, PNLDC1, RFC4, ECT2, MLF1, TSPAN8, SP9, SPP1, CLDN3, CDK1, CCK, HNF4G, NOXO1, ELAVL2, IRF7, DMRTA2, DERL3, SAA1, MDK, HIST2H3A, HMSD, C17orf104, TDRD9, AGR2, APLN, SLC4A4, CNTD2, HOXC4, PAX6, CTCFL, SERPINA1, ZFP57, CDKN2A, ONECUT2, HOXB6, MCIDAS, TNIP3, NWD1, IDO1, IL32, FAM132B, AUNIP, PRAME, HOXC11, ADAM8, NUPR1L, TCF24, ALOX15, HIST2H4B, HIST2H4A, APOC1, NLRP7, HIST2H2AA3, TMPRSS4, CCNI2, CCL18, HIST1H4I, FOXD1, HIST1H2AB, HIST1H3F, HIST1H3G, MUC1 |
| embryonic organ morphogenesis | 21 | 301 | 0.0103 | SIX4, SLC44A4, VAX2, GSC, RDH10, SIX1, MYO3A, OTX1, GBX2, HOXC9, HOXB9, FOXC2, HMX2, ALX3, COL11A1, STIL, EPHB2, HOXC4, PAX6, HOXB6, HOXC11 |

| | | | | |
|---|---|---|---|---|
| regulation of nitrogen compound metabolic process | 185 | 5836 | 0.0106 | YBX2, CELSR3, DMRT3, CDC6, RAB26, OSM, SIX4, AURKA, MYBL2, SALL4, AMH, CSF3, RAD51AP1, GMNN, LTF, TLX2, VAX2, SLC8A2, RARRES1, GSC, HOXC13, HOXC6, TCF15, SIX1, BST2, GDF15, MNX1, BARX1, NUP210, ERN2, IGF2BP3, PKMYT1, CDKN2C, EPCAM, SOX30, LAMP3, ZIC5, HES6, SCGB1A1, MYRF, OTX1, CCNO, BMP3, ATAD2, IGF2BP1, SIM2, GRIN2C, PF4, CAMK2N2, MFI2, TLX3, GBX1, MARVELD3, CDT1, BATF2, CCL11, GBX2, HOXC9, PCSK9, FOXA3, CXCL10, HOXC10, HOXB9, RMI2, SPSB4, CLSPN, NQO1, CDC25C, E2F7, FOXJ1, FOXL1, FOXC2, NOS2, TNFRSF18, HOXD1, RIPPLY3, PTP4A3, ASCL2, HMX2, HIST1H1T, ZNF695, OAS2, IFI6, NKX2-3, NKX2-4, E2F1, DUSP9, WARS, RBM38, UBE2C, CCL20, DMBX1, TFRC, IQGAP3, HIST3H2A, DTL, NEK2, KIF14, AIM2, ROS1, HIST2H3C, HIST2H2AA, FCGR1A, PRDM13, ALX3, HMGB3, VGLL1, FOXD3, ORC1, MMP9, ASS1, CDC20, WFDC2, BMP8B, TAF7L, TREM2, LHX2, SPDEF, EPHB2, PAX7, ZIC2, PTPRH, TRIM15, TRIM31, POU4F1, LBX2, CA9, ISG15, TNNI2, DMRT2, LILRB4, ZNF541, IL24, RFC4, ECT2, MLF1, SP9, SPP1, CLDN3, CDK1, CCK, HNF4G, ELAVL2, IRF7, DMRTA2, DERL3, SAA1, MDK, HIST2H3A, HMSD, C17orf104, APLN, SLC4A4, CNTD2, HOXC4, PAX6, CTCFL, SERPINA1, ZFP57, CDKN2A, ONECUT2, HOXB6, MCIDAS, TNIP3, NWD1, AUNIP, PRAME, HOXC11, ADAM8, NUPR1L, TCF24, ALOX15, HIST2H4B, HIST2H4A, APOC1, NLRP7, HIST2H2AA3, CCNI2, CCL18, HIST1H4I, FOXD1, HIST1H2AB, HIST1H3F, HIST1H3G, MUC1 |
| anatomical structure morphogenesis | 83 | 2165 | 0.011 | CELSR3, DLL3, SIX4, AURKA, SALL4, AMH, SLC44A4, GMNN, LTF, TLX2, VAX2, GSC, RDH10, HOXC13, TCF15, SIX1, GNGT1, MNX1, IGF2BP3, MMP13, LAMC2, MYO3A, LGR5, OTX1, GBX1, CCL11, GBX2, HOXC9, FOXA3, ALCAM, CXCL8, HOXC10, HOXB9, UGT8, NEFH, SYNE4, E2F7, FOXJ1, FOXL1, FOXC2, ZPBP2, AMTN, HMX2, NKX2-3, WARS, SYCP2, KRT19, KIF14, ALX3, COL11A1, FOXD3, STIL, MMP9, VSTM2L, LHX2, NCMAP, EPHB2, PAX7, TRIM15, POU4F1, CA9, ESM1, CSPG5, KRT18, PNLDC1, ECT2, SP9, CLDN3, CCK, WDR72, MDK, ARTN, AGR2, APLN, HOXC4, PAX6, ONECUT2, HOXB6, PROM1, HOXC11, KRT8, ADAM8, FOXD1 |

| | | | | |
|---|---|---|---|---|
| regulation of biological process | 325 | 11475 | 0.0112 | YBX2, TNFRSF17, CELSR3, DMRT3, DLL3, CDC6, RAB26, MISP, MMP11, OSM, USP18, PDXP, SIX4, CYP24A1, AURKA, MYBL2, SALL4, SRMS, AMH, CD79A, MEST, CSF3, PPM1H, RAD51AP1, SLC44A4, GMNN, LTF, TLX2, VAX2, SLC8A2, RARRES1, GSC, RDH10, HOXC13, HOXC6, IL17C, OR2B6, CD70, TCF15, SIX1, SYNGR3, GNGT1, KIAA1244, BST2, GDF15, MNX1, BPIFB1, BARX1, RHPN2, NUP210, NTS, ADAMDEC1, ERN2, GRP, IGF2BP3, ITPKA, GCHFR, DSG2, CHRNB4, CACNG4, PKMYT1, CDKN2C, GRIN2D, EPCAM, LAMC2, SOX30, MCM2, STC2, LAMP3, LGR5, ZIC5, HES6, PLA2G7, SCGB1A1, MYRF, OTX1, CCNO, BMP3, SKA1, ATAD2, TMSNB, IGF2BP1, SIM2, TFF1, GRIN2C, PTCHD2, CXCL3, CXCL5, PF4, TRIM54, CAMK2N2, MFI2, TLX3, GBX1, CLDN10, MMP3, MARVELD3, B4GALNT2, JSRP1, CDT1, BATF2, CCL11, GBX2, HOXC9, PCSK9, MZB1, FOXA3, KCNG3, CXCL10, ALCAM, CXCL8, CXCL11, HOXC10, KCNS1, AGR3, TCP11, HOXB9, RMI2, SPSB4, NEFH, CLSPN, SHISA2, NQO1, CDC25C, MMP1, E2F7, FOXJ1, FOXL1, FOXC2, CLGN, NOS2, APOBEC3B, TNFRSF18, HOXD1, NPW, RIPPLY3, PTP4A3, ASCL2, NXPH4, ZPBP2, AMTN, HMX2, HIST1H1T, ZNF695, ICAM4, OAS2, IFI6, NKX2-3, NKX2-4, E2F1, DUSP9, WARS, C2CD4A, GPR160, RBM38, UBE2C, PIGR, SYCP2, CCL20, DMBX1, TFRC, IQGAP3, CXCL9, C16orf59, KRT19, HIST3H2A, TRIM17, DTL, NEK2, G0S2, C1orf186, KIF14, SHCBP1L, FCGR3A, AIM2, ROS1, HIST2H3C, HIST2H2AA, FCGR1A, PRDM13, ADRB1, VTCN1, ALX3, KIAA1324, ELOVL3, ARHGAP4, HENMT1, HMGB3, GBP5, VGLL1, FOXD3, ZBP1, LRRC26, ORC1, IFIT3, STIL, CEL, KIF2C, MMP9, ASS1, CDC20, WFDC2, PLAU, BMP8B, TAF7L, TREM2, TSPAN15, VSTM2L, MARCKSL1, LHX2, SMPDL3B, SPDEF, NCMAP, EPHB2, PAX7, ZIC2, PTPRH, GPR158, TRIM15, TRIM31, UBD, POU4F1, LBX2, IGFBPL1, CA9, ISG15, IL2RA, C2CD4B, EGFL6, ESM1, TNNI2, SYT8, RSAD2, DMRT2, CSPG5, GOLM1, KRT18, LILRB4, GNG4, ZNF541, IL24, PNLDC1, RFC4, ECT2, MLF1, TSPAN8, NMB, SP9, SPP1, CLDN3, CDK1, CXCL1, CCK, HNF4G, NOXO1, ELAVL2, LYPD1, IRF7, MUC16, DMRTA2, DERL3, SAA1, MDK, HIST2H3A, HMSD, C17orf104, TDRD9, ARTN, AGR2, APLN, SLC4A4, CNTD2, HOXC4, PAX6, CTCFL, SERPINA1, CFB, IGFALS, ZFP57, CDKN2A, PEG10, ONECUT2, RNF183, HOXB6, MCIDAS, TNIP3, PROM1, NWD1, IDO1, IGLL5, IL32, FCGR3B, CRACR2B, RAB19, GPR19, FAM132B, AUNIP, PRAME, HOXC11, KRT8, ADAM8, NUSAP1, NUPR1L, TCF24, ALOX15, HIST2H4B, HIST2H4A, APOC1, NLRP7, HIST2H2AA3, HIST1H2BJ, TMPRSS4, CCNI2, IGHV3-11, IGHV3-15, CCL18, HIST1H4I, FOXD1, SMIM22, HSH2D, HIST1H2AB, HIST1H3F, HIST1H3G, MUC1, MUC5AC |
| meiosis I | 12 | 112 | 0.0114 | AURKA, M1AP, SYCE2, C11orf85, SYCP2, MEI1, C17orf104, TDRD9, RNF212, MEIOB, CNTD1, STAG3 |

| | | | | |
|---|---|---|---|---|
| meiotic nuclear division | 14 | 152 | 0.0125 | AURKA, NUF2, M1AP, SYCE2, C11orf85, SYCP2, SHCBP1L, MEI1, C17orf104, TDRD9, RNF212, MEIOB, CNTD1, STAG3 |
| cellular response to molecule of bacterial origin | 16 | 195 | 0.0131 | CSF3, CMPK2, CXCL3, CXCL5, PF4, CXCL10, CXCL8, CXCL11, NOS2, CXCL9, ASS1, TREM2, IL24, CXCL1, TNIP3, NLRP7 |
| reproductive process | 59 | 1400 | 0.0135 | PRSS21, YBX2, DMRT3, HSD17B2, SIX4, AURKA, AMH, RDH10, MMP7, EPYC, DSG2, ASF1B, SOX30, STC2, DNAH5, LGR5, NUF2, SCGB1A1, M1AP, LY6K, SYCE2, RPL39L, C11orf85, FOXA3, TCP11, E2F7, FOXJ1, CLGN, ASCL2, ZPBP2, HIST1H1T, E2F1, SMC1B, SYCP2, PLAC1, KRT19, NEK2, SHCBP1L, ROS1, MMP9, WFDC2, TAF7L, SYT8, ZNF541, TSPAN8, SPP1, CDK1, MEI1, MDK, C17orf104, TDRD9, RNF212, MEIOB, TDRD5, CTCFL, IDO1, KRT8, CNTD1, STAG3 |
| cellular response to biotic stimulus | 17 | 219 | 0.0142 | CSF3, CMPK2, CXCL3, CXCL5, PF4, CXCL10, CXCL8, CXCL11, NOS2, CXCL9, ASS1, TREM2, IGFBPL1, IL24, CXCL1, TNIP3, NLRP7 |
| regulation of primary metabolic process | 189 | 6032 | 0.0143 | YBX2, CELSR3, DMRT3, CDC6, RAB26, OSM, SIX4, AURKA, MYBL2, SALL4, AMH, CSF3, RAD51AP1, GMNN, LTF, TLX2, VAX2, SLC8A2, RARRES1, GSC, RDH10, HOXC13, HOXC6, TCF15, SIX1, BST2, GDF15, MNX1, BARX1, NUP210, ERN2, IGF2BP3, PKMYT1, CDKN2C, EPCAM, SOX30, LAMP3, ZIC5, HES6, SCGB1A1, MYRF, OTX1, CCNO, BMP3, ATAD2, IGF2BP1, SIM2, GRIN2C, PTCHD2, PF4, CAMK2N2, MFI2, TLX3, GBX1, MARVELD3, CDT1, BATF2, CCL11, GBX2, HOXC9, PCSK9, FOXA3, CXCL10, HOXC10, HOXB9, RMI2, SPSB4, CLSPN, NQO1, CDC25C, E2F7, FOXJ1, FOXL1, FOXC2, NOS2, TNFRSF18, HOXD1, RIPPLY3, PTP4A3, ASCL2, HMX2, HIST1H1T, ZNF695, OAS2, IFI6, NKX2-3, NKX2-4, E2F1, DUSP9, WARS, RBM38, UBE2C, CCL20, DMBX1, TFRC, IQGAP3, HIST3H2A, DTL, NEK2, G0S2, KIF14, AIM2, ROS1, HIST2H3C, HIST2H2AA, FCGR1A, PRDM13, ALX3, HMGB3, VGLL1, FOXD3, ORC1, MMP9, CDC20, WFDC2, BMP8B, TAF7L, TREM2, LHX2, SPDEF, EPHB2, PAX7, ZIC2, PTPRH, TRIM15, TRIM31, POU4F1, LBX2, CA9, ISG15, TNNI2, DMRT2, GOLM1, LILRB4, ZNF541, IL24, RFC4, ECT2, MLF1, SP9, SPP1, CLDN3, CDK1, CCK, HNF4G, ELAVL2, IRF7, DMRTA2, DERL3, SAA1, MDK, HIST2H3A, HMSD, C17orf104, APLN, SLC4A4, CNTD2, HOXC4, PAX6, CTCFL, SERPINA1, ZFP57, CDKN2A, ONECUT2, HOXB6, MCIDAS, TNIP3, NWD1, FAM132B, AUNIP, PRAME, HOXC11, ADAM8, NUPR1L, TCF24, ALOX15, HIST2H4B, HIST2H4A, APOC1, NLRP7, HIST2H2AA3, CCNI2, CCL18, HIST1H4I, FOXD1, HIST1H2AB, HIST1H3F, HIST1H3G, MUC1 |

| | | | | |
|---|---|---|---|---|
| rDNA heterochromatin assembly | 7 | 37 | 0.0177 | HIST2H3C, HIST2H3A, HIST2H4B, HIST2H4A, HIST1H4I, HIST1H3F, HIST1H3G |
| positive regulation of metabolic process | 131 | 3893 | 0.0177 | YBX2, CDC6, OSM, SIX4, AURKA, MYBL2, SALL4, AMH, CSF3, PPM1H, LTF, TLX2, SLC8A2, RDH10, HOXC13, TCF15, SIX1, GDF15, BARX1, ERN2, EPCAM, SOX30, LAMP3, MYRF, OTX1, BMP3, ATAD2, IGF2BP1, SIM2, PTCHD2, PF4, MFI2, TLX3, GBX1, CDT1, CCL11, GBX2, HOXC9, PCSK9, MZB1, FOXA3, CXCL10, HOXC10, HOXB9, SPSB4, CLSPN, E2F7, FOXJ1, FOXC2, NOS2, TNFRSF18, ASCL2, HMX2, HIST1H1T, NKX2-3, NKX2-4, E2F1, DUSP9, WARS, RBM38, UBE2C, CCL20, TFRC, IQGAP3, DTL, NEK2, G0S2, KIF14, AIM2, ROS1, FCGR1A, ADRB1, VTCN1, KIAA1324, ELOVL3, HMGB3, GBP5, VGLL1, FOXD3, ZBP1, MMP9, ASS1, CDC20, BMP8B, TAF7L, TREM2, LHX2, SPDEF, EPHB2, PAX7, ZIC2, TRIM15, POU4F1, TNNI2, RSAD2, DMRT2, IL24, RFC4, ECT2, SP9, SPP1, CLDN3, CDK1, CCK, HNF4G, IRF7, SAA1, MDK, HMSD, C17orf104, AGR2, APLN, SLC4A4, HOXC4, PAX6, CTCFL, CDKN2A, ONECUT2, MCIDAS, TNIP3, NWD1, IDO1, IL32, HOXC11, ADAM8, ALOX15, APOC1, NLRP7, CCL18, FOXD1, MUC1 |
| regulation of gene expression, epigenetic | 16 | 202 | 0.0177 | ATAD2, HIST1H1T, HIST3H2A, HIST2H3C, HIST2H2AA, LHX2, HIST2H3A, CTCFL, ZFP57, HIST2H4B, HIST2H4A, HIST2H2AA3, HIST1H4I, HIST1H2AB, HIST1H3F, HIST1H3G |
| regulation of cell population proliferation | 66 | 1642 | 0.0177 | CDC6, OSM, SIX4, SRMS, CSF3, LTF, RARRES1, CD70, SIX1, BST2, CDKN2C, EPCAM, LAMC2, LGR5, SCGB1A1, TFF1, PTCHD2, CXCL5, PF4, MARVELD3, CCL11, MZB1, CXCL10, CXCL8, CXCL11, HOXC10, E2F7, FOXJ1, NOS2, RIPPLY3, ASCL2, HMX2, NKX2-3, E2F1, WARS, RBM38, TFRC, IQGAP3, CXCL9, KIF14, VTCN1, IFIT3, MMP9, CDC20, PLAU, MARCKSL1, LHX2, PAX7, IL2RA, ESM1, IL24, NMB, CLDN3, CDK1, CXCL1, CCK, DMRTA2, MDK, APLN, PAX6, CDKN2A, IDO1, PRAME, NUPR1L, SMIM22, HIST1H2AB |
| skeletal system development | 28 | 499 | 0.0178 | DLL3, SIX4, LTF, GSC, RDH10, HOXC6, TCF15, SIX1, MMP13, EPYC, HAPLN1, BMP3, HOXC9, HOXC10, HOXB9, FOXC2, COL10A1, HOXD1, ALX3, COL11A1, MMP9, COL9A2, BMP8B, PAX7, DMRT2, HOXC4, HOXB6, HOXC11 |
| embryonic organ development | 26 | 448 | 0.0188 | SIX4, SLC44A4, VAX2, GSC, RDH10, SIX1, MYO3A, OTX1, GBX2, HOXC9, CXCL8, HOXB9, E2F7, FOXC2, ASCL2, HMX2, KRT19, ALX3, COL11A1, STIL, EPHB2, HOXC4, PAX6, HOXB6, HOXC11, KRT8 |

| | | | | |
|---|---|---|---|---|
| positive regulation of biological process | 190 | 6112 | 0.0197 | YBX2, CDC6, OSM, PDXP, SIX4, AURKA, MYBL2, SALL4, AMH, CD79A, CSF3, PPM1H, SLC44A4, LTF, TLX2, SLC8A2, RDH10, HOXC13, CD70, TCF15, SIX1, SYNGR3, BST2, GDF15, BARX1, ERN2, GRP, ITPKA, CHRNB4, CACNG4, GRIN2D, EPCAM, LAMC2, SOX30, LAMP3, LGR5, PLA2G7, MYRF, OTX1, BMP3, ATAD2, IGF2BP1, SIM2, GRIN2C, PTCHD2, CXCL5, PF4, MFI2, TLX3, GBX1, MMP3, CDT1, CCL11, GBX2, HOXC9, PCSK9, MZB1, FOXA3, CXCL10, CXCL8, CXCL11, HOXC10, HOXB9, SPSB4, CLSPN, NQO1, CDC25C, MMP1, E2F7, FOXJ1, FOXC2, NOS2, TNFRSF18, PTP4A3, ASCL2, AMTN, HMX2, HIST1H1T, NKX2-3, NKX2-4, E2F1, DUSP9, WARS, C2CD4A, RBM38, UBE2C, CCL20, TFRC, IQGAP3, CXCL9, C16orf59, DTL, NEK2, G0S2, C1orf186, KIF14, SHCBP1L, FCGR3A, AIM2, ROS1, FCGR1A, ADRB1, VTCN1, KIAA1324, ELOVL3, HMGB3, GBP5, VGLL1, FOXD3, ZBP1, LRRC26, MMP9, ASS1, CDC20, PLAU, BMP8B, TAF7L, TREM2, MARCKSL1, LHX2, SPDEF, NCMAP, EPHB2, PAX7, ZIC2, TRIM15, TRIM31, UBD, POU4F1, ISG15, IL2RA, C2CD4B, EGFL6, ESM1, TNNI2, RSAD2, DMRT2, CSPG5, LILRB4, IL24, RFC4, ECT2, NMB, SP9, SPP1, CLDN3, CDK1, CCK, HNF4G, IRF7, MUC16, DMRTA2, SAA1, MDK, HMSD, C17orf104, ARTN, AGR2, APLN, SLC4A4, HOXC4, PAX6, CTCFL, CFB, CDKN2A, ONECUT2, RNF183, MCIDAS, TNIP3, PROM1, NWD1, IDO1, IGLL5, IL32, FAM132B, PRAME, HOXC11, ADAM8, NUSAP1, NUPR1L, ALOX15, APOC1, NLRP7, IGHV3-11, IGHV3-15, CCL18, FOXD1, SMIM22, MUC1, MUC5AC |
| DNA replication-independent nucleosome assembly | 8 | 53 | 0.0208 | CENPM, OIP5, ASF1B, CENPA, CENPW, HIST2H4B, HIST2H4A, HIST1H4I |
| cellular response to lipopolysaccharide | 15 | 185 | 0.021 | CSF3, CMPK2, CXCL3, CXCL5, PF4, CXCL10, CXCL8, CXCL11, NOS2, CXCL9, ASS1, IL24, CXCL1, TNIP3, NLRP7 |

| | | | | |
|---|---|---|---|---|
| cell differentiation | 125 | 3702 | 0.0215 | YBX2, CELSR3, DMRT3, DLL3, SIX4, CYP24A1, AURKA, SRMS, AMH, CD79A, CSF3, SLC44A4, TLX2, VAX2, GSC, RDH10, TCF15, SIX1, GNGT1, MNX1, C1QL1, BARX1, ALPK3, ITPKA, DSG2, CDKN2C, ASF1B, EPCAM, LAMC2, SOX30, LGR5, ZIC5, HES6, MYRF, CCNO, BMP3, SIM2, M1AP, TFF1, PTCHD2, TRIM54, TLX3, GBX1, BATF2, CCL11, GBX2, PCSK9, FOXA3, GPRIN1, ALCAM, HOXC10, TCP11, UGT8, NEFH, SYNE4, E2F7, FOXJ1, FOXL1, FOXC2, HOXD1, KRT7, ASCL2, ZPBP2, HMX2, HIST1H1T, NKX2-3, NKX2-4, E2F1, RBM38, TFRC, KRT19, C1orf186, KIF14, SHCBP1L, ROS1, PRDM13, ADRB1, ARHGAP4, COL11A1, FOXD3, MMP9, CDC20, BMP8B, TAF7L, TREM2, VSTM2L, LHX2, SPDEF, NCMAP, EPHB2, PAX7, ZIC2, TRIM15, UBD, POU4F1, EGFL6, RSAD2, CSPG5, KRT18, ZNF541, ECT2, MLF1, SPP1, CLDN3, CDK1, CCK, HNF4G, KRTAP4-1, DMRTA2, MEI1, MDK, C17orf104, TDRD9, ARTN, AGR2, PAX6, TDRD5, PEG10, ONECUT2, MCIDAS, PROM1, KRT86, PRAME, KRT8, FOXD1 |
| meiotic chromosome segregation | 10 | 87 | 0.0221 | NUF2, M1AP, SYCE2, C11orf85, SYCP2, MEI1, C17orf104, RNF212, MEIOB, STAG3 |
| positive regulation of gene expression | 86 | 2337 | 0.0226 | OSM, SIX4, MYBL2, SALL4, AMH, CSF3, TLX2, HOXC13, TCF15, SIX1, BARX1, EPCAM, SOX30, LAMP3, MYRF, OTX1, ATAD2, IGF2BP1, SIM2, PF4, MFI2, TLX3, GBX1, GBX2, HOXC9, FOXA3, CXCL10, HOXC10, HOXB9, E2F7, FOXJ1, FOXC2, ASCL2, HMX2, HIST1H1T, NKX2-3, NKX2-4, E2F1, WARS, RBM38, TFRC, IQGAP3, AIM2, VTCN1, HMGB3, GBP5, FOXD3, ZBP1, TREM2, LHX2, SPDEF, EPHB2, PAX7, ZIC2, TRIM15, POU4F1, TNNI2, RSAD2, DMRT2, SP9, SPP1, CLDN3, CDK1, HNF4G, IRF7, SAA1, MDK, HMSD, C17orf104, AGR2, APLN, HOXC4, PAX6, CTCFL, CDKN2A, ONECUT2, MCIDAS, TNIP3, NWD1, IDO1, IL32, HOXC11, ADAM8, NLRP7, FOXD1, MUC1 |

| | | | | |
|---|---|---|---|---|
| regulation of macromolecule metabolic process | 197 | 6407 | 0.0226 | YBX2, CELSR3, DMRT3, CDC6, RAB26, OSM, SIX4, AURKA, MYBL2, SALL4, AMH, CSF3, RAD51AP1, GMNN, LTF, TLX2, VAX2, SLC8A2, RARRES1, GSC, HOXC13, HOXC6, TCF15, SIX1, BST2, GDF15, MNX1, BARX1, NUP210, ERN2, IGF2BP3, PKMYT1, CDKN2C, EPCAM, SOX30, STC2, LAMP3, ZIC5, HES6, SCGB1A1, MYRF, OTX1, CCNO, BMP3, ATAD2, IGF2BP1, SIM2, GRIN2C, PF4, TRIM54, CAMK2N2, MFI2, TLX3, GBX1, MARVELD3, CDT1, BATF2, CCL11, GBX2, HOXC9, PCSK9, MZB1, FOXA3, CXCL10, HOXC10, HOXB9, RMI2, SPSB4, CLSPN, CDC25C, E2F7, FOXJ1, FOXL1, FOXC2, NOS2, TNFRSF18, HOXD1, RIPPLY3, PTP4A3, ASCL2, HMX2, HIST1H1T, ZNF695, OAS2, IFI6, NKX2-3, NKX2-4, E2F1, DUSP9, WARS, RBM38, UBE2C, CCL20, DMBX1, TFRC, IQGAP3, HIST3H2A, TRIM17, DTL, NEK2, KIF14, AIM2, ROS1, HIST2H3C, HIST2H2AA, FCGR1A, PRDM13, VTCN1, ALX3, HENMT1, HMGB3, GBP5, VGLL1, FOXD3, ZBP1, ORC1, MMP9, CDC20, WFDC2, BMP8B, TAF7L, TREM2, LHX2, SPDEF, EPHB2, PAX7, ZIC2, PTPRH, TRIM15, TRIM31, POU4F1, LBX2, CA9, ISG15, TNNI2, RSAD2, DMRT2, LILRB4, ZNF541, IL24, PNLDC1, RFC4, ECT2, MLF1, TSPAN8, SP9, SPP1, CLDN3, CDK1, CCK, HNF4G, ELAVL2, IRF7, DMRTA2, DERL3, SAA1, MDK, HIST2H3A, HMSD, C17orf104, TDRD9, AGR2, APLN, CNTD2, HOXC4, PAX6, CTCFL, SERPINA1, ZFP57, CDKN2A, ONECUT2, HOXB6, MCIDAS, TNIP3, NWD1, IDO1, IL32, AUNIP, PRAME, HOXC11, ADAM8, NUPR1L, TCF24, ALOX15, HIST2H4B, HIST2H4A, NLRP7, HIST2H2AA3, TMPRSS4, CCNI2, CCL18, HIST1H4I, FOXD1, HIST1H2AB, HIST1H3F, HIST1H3G, MUC1 |
| mitotic cell cycle phase transition | 19 | 280 | 0.0227 | CDC6, AURKA, GMNN, PKMYT1, CDKN2C, MCM2, CCNO, CDT1, CDC25C, E2F7, E2F1, UBE2C, IQGAP3, NEK2, ORC1, CDK1, CNTD2, CDKN2A, CCNI2 |
| cellular developmental process | 126 | 3757 | 0.0247 | YBX2, CELSR3, DMRT3, DLL3, SIX4, CYP24A1, AURKA, SRMS, AMH, CD79A, CSF3, SLC44A4, TLX2, VAX2, GSC, RDH10, TCF15, SIX1, GNGT1, MNX1, C1QL1, BARX1, ALPK3, ITPKA, DSG2, CDKN2C, ASF1B, EPCAM, LAMC2, SOX30, LGR5, ZIC5, HES6, MYRF, CCNO, BMP3, SIM2, M1AP, TFF1, PTCHD2, TRIM54, TLX3, GBX1, BATF2, CCL11, GBX2, PCSK9, FOXA3, GPRIN1, ALCAM, HOXC10, TCP11, UGT8, NEFH, SYNE4, E2F7, FOXJ1, FOXL1, FOXC2, HOXD1, KRT7, ASCL2, ZPBP2, HMX2, HIST1H1T, NKX2-3, NKX2-4, E2F1, RBM38, TFRC, KRT19, C1orf186, KIF14, SHCBP1L, ROS1, PRDM13, ADRB1, ARHGAP4, COL11A1, FOXD3, MMP9, CDC20, BMP8B, TAF7L, TREM2, VSTM2L, LHX2, SPDEF, NCMAP, EPHB2, PAX7, ZIC2, TRIM15, UBD, POU4F1, EGFL6, RSAD2, CSPG5, KRT18, ZNF541, ECT2, MLF1, SPP1, CLDN3, CDK1, CCK, HNF4G, KRTAP4-1, DMRTA2, MEI1, MDK, C17orf104, TDRD9, ARTN, AGR2, PAX6, TDRD5, CDKN2A, PEG10, ONECUT2, MCIDAS, PROM1, KRT86, PRAME, KRT8, FOXD1 |

| | | | | |
|---|---|---|---|---|
| biological regulation | 338 | 12171 | 0.0279 | YBX2, TNFRSF17, CELSR3, DMRT3, HSD17B2, DLL3, CDC6, RAB26, MISP, MMP11, OSM, USP18, PDXP, SIX4, CYP24A1, AURKA, MYBL2, SALL4, SRMS, AMH, CD79A, SLC5A5, MEST, CSF3, PPM1H, RAD51AP1, SLC44A4, GMNN, LTF, RBP1, TLX2, VAX2, SLC8A2, RARRES1, GSC, RDH10, HOXC13, HOXC6, IL17C, OR2B6, CD70, TCF15, SIX1, SYNGR3, GNGT1, KIAA1244, BST2, GDF15, MNX1, BPIFB1, BARX1, RHPN2, NUP210, NTS, ADAMDEC1, ERN2, GRP, IGF2BP3, ITPKA, GCHFR, DSG2, CHRNB4, CACNG4, PKMYT1, CDKN2C, GRIN2D, EPCAM, LAMC2, CP, SOX30, MCM2, STC2, LAMP3, LGR5, ZIC5, HES6, PLA2G7, SCGB1A1, MYRF, OTX1, CCNO, BMP3, SKA1, ATAD2, TMSNB, IGF2BP1, SIM2, TFF1, TMPRSS3, GRIN2C, PTCHD2, CXCL3, CXCL5, PF4, TRIM54, CAMK2N2, MFI2, TLX3, GBX1, CLDN10, MMP3, MARVELD3, B4GALNT2, JSRP1, CDT1, BATF2, CCL11, GBX2, HOXC9, PCSK9, MZB1, FOXA3, KCNG3, CXCL10, ALCAM, CXCL8, CXCL11, HOXC10, KCNS1, AGR3, TCP11, HOXB9, RMI2, PCP2, SPSB4, NEFH, CLSPN, SHISA2, NQO1, CDC25C, MMP1, E2F7, FOXJ1, FOXL1, FOXC2, CLGN, NOS2, APOBEC3B, TNFRSF18, HOXD1, NPW, RIPPLY3, PTP4A3, ASCL2, NXPH4, ZPBP2, AMTN, HMX2, HIST1H1T, ZNF695, ICAM4, OAS2, IFI6, NKX2-3, PLEKHG7, NKX2-4, E2F1, DUSP9, WARS, C2CD4A, GPR160, RBM38, UBE2C, PIGR, SYCP2, CCL20, DMBX1, TFRC, PLEKHG4, IQGAP3, CXCL9, C16orf59, KRT19, HIST3H2A, TRIM17, DTL, NEK2, G0S2, C1orf186, KIF14, SHCBP1L, FCGR3A, AIM2, ROS1, HIST2H3C, HIST2H2AA, FCGR1A, PRDM13, ADRB1, VTCN1, ALX3, KIAA1324, ELOVL3, ARHGAP4, HENMT1, HMGB3, GBP5, VGLL1, FOXD3, ZBP1, LRRC26, ORC1, IFIT3, STIL, TSPAN1, CEL, KIF2C, MMP9, ASS1, CDC20, WFDC2, PLAU, BMP8B, TAF7L, TREM2, TSPAN15, VSTM2L, MARCKSL1, LHX2, SMPDL3B, SPDEF, NCMAP, EPHB2, PAX7, ZIC2, PTPRH, GPR158, TRIM15, TRIM31, UBD, POU4F1, LBX2, IGFBPL1, CA9, ISG15, IL2RA, C2CD4B, EGFL6, ESM1, TNNI2, SYT8, RSAD2, SLC34A2, DMRT2, CSPG5, GOLM1, KRT18, LILRB4, GNG4, ZNF541, IL24, PNLDC1, RFC4, ECT2, MLF1, TSPAN8, NMB, SP9, SPP1, CLDN3, CDK1, CXCL1, CCK, HNF4G, NOXO1, ELAVL2, LYPD1, IRF7, MUC16, DMRTA2, CYP4F11, DERL3, SAA1, MDK, HIST2H3A, HMSD, C17orf104, TDRD9, ARTN, AGR2, APLN, SLC4A4, CNTD2, HOXC4, PAX6, CTCFL, SERPINA1, ARHGEF38, CFB, IGFALS, ZFP57, CDKN2A, PEG10, ONECUT2, RNF183, HOXB6, MCIDAS, TNIP3, PROM1, NWD1, IDO1, IGLL5, IL32, FCGR3B, CRACR2B, RAB19, GPR19, FAM132B, AUNIP, PRAME, HOXC11, KRT8, ADAM8, NUSAP1, NUPR1L, TCF24, ALOX15, HIST2H4B, HIST2H4A, APOC1, NLRP7, HIST2H2AA3, HIST1H2BJ, TMPRSS4, CCNI2, IGHV3-11, IGHV3-15, CCL18, HIST1H4I, FOXD1, SMIM22, RECQL4, HSH2D, HIST1H2AB, HIST1H3F, HIST1H3G, MUC1, MUC5AC |

| | | | | |
|---|---|---|---|---|
| cell division | 27 | 493 | 0.0288 | CDC6, MISP, AURKA, OIP5, FAM64A, NUF2, CCNO, SKA1, SYCE2, CDT1, CDC25C, CENPA, UBE2C, SYCP2, NEK2, KIF14, SHCBP1L, CENPW, KIF2C, CDC20, KIF4A, ECT2, CDK1, SEPT3, HEPACAM2, CDKN2A, NUSAP1 |
| collagen catabolic process | 7 | 43 | 0.0317 | MMP11, MMP7, MMP13, MMP10, MMP3, MMP1, MMP9 |
| heterochromatin assembly | 8 | 58 | 0.0317 | HIST2H3C, HIST2H3A, CDKN2A, HIST2H4B, HIST2H4A, HIST1H4I, HIST1H3F, HIST1H3G |
| CENP-A containing nucleosome assembly | 7 | 43 | 0.0317 | CENPM, OIP5, CENPA, CENPW, HIST2H4B, HIST2H4A, HIST1H4I |
| microtubule cytoskeleton organization involved in mitosis | 11 | 112 | 0.032 | MISP, AURKA, MYBL2, NUF2, CENPA, NEK2, STIL, CDC20, KIF4A, PAX6, NUSAP1 |
| homologous chromosome segregation | 8 | 59 | 0.0332 | SYCE2, C11orf85, SYCP2, MEI1, C17orf104, RNF212, MEIOB, STAG3 |
| regulation of cellular process | 308 | 10932 | 0.0332 | YBX2, TNFRSF17, CELSR3, DMRT3, DLL3, CDC6, RAB26, MISP, MMP11, OSM, USP18, PDXP, SIX4, CYP24A1, AURKA, MYBL2, SALL4, SRMS, AMH, CD79A, CSF3, RAD51AP1, SLC44A4, GMNN, LTF, TLX2, VAX2, SLC8A2, RARRES1, GSC, RDH10, HOXC13, HOXC6, IL17C, OR2B6, CD70, TCF15, SIX1, GNGT1, KIAA1244, BST2, GDF15, MNX1, BPIFB1, BARX1, RHPN2, NUP210, NTS, ADAMDEC1, ERN2, GRP, IGF2BP3, ITPKA, DSG2, CHRNB4, CACNG4, PKMYT1, CDKN2C, GRIN2D, EPCAM, LAMC2, SOX30, MCM2, STC2, LAMP3, LGR5, ZIC5, HES6, PLA2G7, SCGB1A1, MYRF, OTX1, CCNO, BMP3, SKA1, ATAD2, TMSNB, IGF2BP1, SIM2, TFF1, GRIN2C, PTCHD2, CXCL3, CXCL5, PF4, TRIM54, CAMK2N2, MFI2, TLX3, GBX1, MMP3, MARVELD3, B4GALNT2, JSRP1, CDT1, BATF2, CCL11, GBX2, HOXC9, PCSK9, MZB1, FOXA3, KCNG3, CXCL10, ALCAM, CXCL8, CXCL11, HOXC10, KCNS1, AGR3, TCP11, HOXB9, RMI2, SPSB4, NEFH, CLSPN, SHISA2, NQO1, CDC25C, MMP1, E2F7, FOXJ1, FOXL1, FOXC2, CLGN, NOS2, APOBEC3B, TNFRSF18, HOXD1, NPW, RIPPLY3, PTP4A3, ASCL2, NXPH4, HMX2, HIST1H1T, ZNF695, OAS2, IFI6, NKX2-3, NKX2-4, E2F1, DUSP9, WARS, C2CD4A, GPR160, RBM38, UBE2C, PIGR, SYCP2, CCL20, DMBX1, TFRC, IQGAP3, CXCL9, C16orf59, KRT19, HIST3H2A, DTL, NEK2, G0S2, C1orf186, KIF14, SHCBP1L, FCGR3A, AIM2, ROS1, HIST2H3C, HIST2H2AA, FCGR1A, PRDM13, ADRB1, |

| | | | | |
|---|---|---|---|---|
| | | | | VTCN1, ALX3, KIAA1324, ARHGAP4, HMGB3, GBP5, VGLL1, FOXD3, ZBP1, LRRC26, ORC1, IFIT3, STIL, CEL, KIF2C, MMP9, ASS1, CDC20, WFDC2, PLAU, BMP8B, TAF7L, TREM2, TSPAN15, VSTM2L, MARCKSL1, LHX2, SMPDL3B, SPDEF, NCMAP, EPHB2, PAX7, ZIC2, PTPRH, GPR158, TRIM15, TRIM31, UBD, POU4F1, LBX2, IGFBPL1, CA9, ISG15, IL2RA, C2CD4B, EGFL6, ESM1, TNNI2, SYT8, RSAD2, DMRT2, CSPG5, KRT18, LILRB4, GNG4, ZNF541, IL24, RFC4, ECT2, MLF1, NMB, SP9, SPP1, CLDN3, CDK1, CXCL1, CCK, HNF4G, NOXO1, ELAVL2, LYPD1, IRF7, MUC16, DMRTA2, DERL3, SAA1, MDK, HIST2H3A, HMSD, C17orf104, TDRD9, ARTN, AGR2, APLN, SLC4A4, CNTD2, HOXC4, PAX6, CTCFL, SERPINA1, IGFALS, ZFP57, CDKN2A, PEG10, ONECUT2, RNF183, HOXB6, MCIDAS, TNIP3, PROM1, NWD1, IDO1, IGLL5, IL32, FCGR3B, RAB19, GPR19, FAM132B, AUNIP, PRAME, HOXC11, KRT8, ADAM8, NUSAP1, NUPR1L, TCF24, ALOX15, HIST2H4B, HIST2H4A, APOC1, NLRP7, HIST2H2AA3, HIST1H2BJ, CCNI2, IGHV3-11, IGHV3-15, CCL18, HIST1H4I, FOXD1, SMIM22, HSH2D, HIST1H2AB, HIST1H3F, HIST1H3G, MUC1, MUC5AC |
| meiotic cell cycle | 16 | 220 | 0.0332 | AURKA, NUF2, M1AP, SYCE2, C11orf85, SMC1B, SYCP2, NEK2, SHCBP1L, MEI1, C17orf104, TDRD9, RNF212, MEIOB, CNTD1, STAG3 |
| regulation of transcription by RNA polymerase II | 80 | 2172 | 0.0343 | DMRT3, OSM, SIX4, MYBL2, SALL4, CSF3, TLX2, VAX2, GSC, HOXC13, HOXC6, TCF15, SIX1, MNX1, BARX1, EPCAM, SOX30, ZIC5, HES6, SCGB1A1, MYRF, OTX1, ATAD2, SIM2, PF4, TLX3, GBX1, BATF2, GBX2, HOXC9, FOXA3, CXCL10, HOXC10, HOXB9, E2F7, FOXJ1, FOXL1, FOXC2, HOXD1, RIPPLY3, ASCL2, HMX2, NKX2-3, NKX2-4, E2F1, DMBX1, PRDM13, ALX3, HMGB3, VGLL1, FOXD3, TAF7L, LHX2, SPDEF, PAX7, ZIC2, POU4F1, LBX2, CA9, DMRT2, LILRB4, ZNF541, SP9, HNF4G, IRF7, DMRTA2, APLN, HOXC4, PAX6, CTCFL, CDKN2A, ONECUT2, HOXB6, MCIDAS, TNIP3, HOXC11, NUPR1L, TCF24, FOXD1, MUC1 |
| innate immune response | 34 | 703 | 0.0369 | LTF, BST2, BPIFB1, CCL11, APOL1, NOS2, APOBEC3B, OAS2, IFI6, CCL20, TRIM17, AIM2, FCGR1A, HMGB3, GBP5, ZBP1, IFIT3, ASS1, WFDC2, TREM2, SMPDL3B, TRIM15, TRIM31, UBD, ISG15, RSAD2, IRF7, SAA1, CFB, IGLL5, HIST1H2BJ, IGHV3-11, IGHV3-15, CCL18 |
| positive regulation of transcription by RNA polymerase II | 52 | 1253 | 0.0369 | OSM, SIX4, MYBL2, SALL4, CSF3, TLX2, HOXC13, TCF15, SIX1, BARX1, EPCAM, SOX30, OTX1, ATAD2, SIM2, PF4, TLX3, GBX1, GBX2, HOXC9, FOXA3, CXCL10, HOXC10, HOXB9, E2F7, FOXJ1, FOXC2, ASCL2, NKX2-3, NKX2-4, E2F1, HMGB3, FOXD3, LHX2, SPDEF, PAX7, POU4F1, DMRT2, SP9, HNF4G, IRF7, APLN, HOXC4, PAX6, CTCFL, CDKN2A, ONECUT2, MCIDAS, TNIP3, HOXC11, FOXD1, MUC1 |

| | | | | |
|---|---|---|---|---|
| mitotic nuclear division | 13 | 156 | 0.0369 | MISP, AURKA, MYBL2, NUF2, CDT1, UBE2C, NEK2, KIF14, KIF2C, CDC20, KIF4A, CDK1, NUSAP1 |
| male meiotic nuclear division | 7 | 46 | 0.0414 | M1AP, SYCP2, SHCBP1L, MEI1, C17orf104, TDRD9, MEIOB |
| DNA replication-dependent nucleosome assembly | 6 | 32 | 0.042 | ASF1B, HIST2H4B, HIST2H4A, HIST1H4I, HIST1H3F, HIST1H3G |
| regulation of transcription, DNA-templated | 114 | 3388 | 0.0426 | DMRT3, CDC6, OSM, SIX4, MYBL2, SALL4, AMH, CSF3, GMNN, LTF, TLX2, VAX2, GSC, HOXC13, HOXC6, TCF15, SIX1, MNX1, BARX1, ERN2, EPCAM, SOX30, ZIC5, HES6, SCGB1A1, MYRF, OTX1, ATAD2, SIM2, PF4, TLX3, GBX1, CDT1, BATF2, GBX2, HOXC9, FOXA3, CXCL10, HOXC10, HOXB9, E2F7, FOXJ1, FOXL1, FOXC2, HOXD1, RIPPLY3, PTP4A3, ASCL2, HMX2, HIST1H1T, ZNF695, NKX2-3, NKX2-4, E2F1, DMBX1, TFRC, HIST3H2A, AIM2, HIST2H3C, HIST2H2AA, PRDM13, ALX3, HMGB3, VGLL1, FOXD3, ORC1, TAF7L, LHX2, SPDEF, PAX7, ZIC2, TRIM15, TRIM31, POU4F1, LBX2, CA9, TNNI2, DMRT2, LILRB4, ZNF541, MLF1, SP9, SPP1, HNF4G, ELAVL2, IRF7, DMRTA2, MDK, HIST2H3A, APLN, HOXC4, PAX6, CTCFL, ZFP57, CDKN2A, ONECUT2, HOXB6, MCIDAS, TNIP3, NWD1, PRAME, HOXC11, ADAM8, NUPR1L, TCF24, HIST2H4B, HIST2H4A, HIST2H2AA3, HIST1H4I, FOXD1, HIST1H2AB, HIST1H3F, HIST1H3G, MUC1 |
| cell chemotaxis | 15 | 204 | 0.0426 | CXCL3, CXCL5, PF4, CCL11, CXCL10, CXCL8, CXCL11, HOXB9, CCL20, CXCL9, CXCL1, SAA1, MDK, ADAM8, CCL18 |
| DNA replication | 15 | 205 | 0.0444 | CDC6, RNASEH2A, GMNN, GINS2, GINS1, MCM2, GINS4, CDT1, RMI2, CLSPN, DTL, ORC1, RFC4, CDK1, RECQL4 |
| tissue development | 67 | 1760 | 0.0461 | DLL3, SIX4, SALL4, AMH, MEST, SLC44A4, TLX2, VAX2, GSC, RDH10, HOXC13, TCF15, SIX1, BARX1, ALPK3, MMP13, EPYC, DSG2, EPCAM, LAMC2, STC2, LGR5, CCNO, BMP3, CCL11, GBX2, SYNE4, E2F7, FOXJ1, FOXL1, FOXC2, KRT7, ASCL2, AMTN, NKX2-3, KRT19, ROS1, COL11A1, STIL, MMP9, BMP8B, LHX2, SPDEF, PAX7, TRIM15, POU4F1, CA9, KRT18, SPP1, CLDN3, CDK1, WDR72, KRTAP4-1, DMRTA2, MDK, ARTN, AGR2, HOXC4, PAX6, ONECUT2, MCIDAS, PROM1, KRT86, HOXC11, KRT8, ALOX15, FOXD1 |
| embryonic morphogenesis | 29 | 571 | 0.0461 | SIX4, SALL4, SLC44A4, TLX2, VAX2, GSC, RDH10, SIX1, MYO3A, OTX1, GBX2, HOXC9, HOXC10, HOXB9, FOXC2, HMX2, ALX3, COL11A1, STIL, MMP9, LHX2, EPHB2, TRIM15, SP9, APLN, HOXC4, PAX6, HOXB6, HOXC11 |

| | | | | |
|---|---|---|---|---|
| cell migration | 40 | 896 | 0.0466 | CELSR3, MISP, SIX4, SRMS, SIX1, EPCAM, CXCL3, CXCL5, PF4, S100P, TLX3, CCL11, GBX2, CXCL10, CXCL8, CXCL11, OLR1, HOXB9, SLC16A8, MMP1, FOXJ1, PTP4A3, NKX2-3, PODXL2, CCL20, CXCL9, PLAU, POU4F1, CDK1, CXCL1, CCK, SAA1, MDK, ARTN, PAX6, IGLL5, ADAM8, SLC16A3, IGHV3-11, CCL18 |
| homologous chromosome pairing at meiosis | 7 | 48 | 0.0483 | SYCE2, C11orf85, SYCP2, C17orf104, RNF212, MEIOB, STAG3 |
| interspecies interaction between organisms | 71 | 1899 | 0.0483 | CSF3, LTF, BST2, BPIFB1, NUP210, ACY3, NTS, CMPK2, SCGB1A1, SLFN13, CXCL3, CXCL5, PF4, BATF2, CCL11, CXCL10, CXCL8, CXCL11, APOL1, CDC25C, MMP1, NOS2, APOBEC3B, KRT7, CENPA, OAS2, IFI6, E2F1, TAP1, RBM38, CCL20, TFRC, CXCL9, KRT19, TRIM17, CHIT1, AIM2, FCGR1A, VTCN1, HMGB3, GBP5, IFI44L, ZBP1, IFIT3, ASS1, WFDC2, TREM2, SMPDL3B, TRIM15, TRIM31, UBD, ISG15, RSAD2, KRT18, IL24, CDK1, CXCL1, IRF7, SAA1, APLN, CLDN9, CFB, TNIP3, IDO1, IGLL5, KRT8, NLRP7, HIST1H2BJ, IGHV3-11, IGHV3-15, CCL18 |
| defense response to other organism | 40 | 900 | 0.0491 | LTF, BST2, BPIFB1, SLFN13, PF4, BATF2, CCL11, CXCL10, APOL1, NOS2, APOBEC3B, OAS2, IFI6, CCL20, CXCL9, TRIM17, AIM2, FCGR1A, HMGB3, GBP5, IFI44L, ZBP1, IFIT3, ASS1, WFDC2, TREM2, SMPDL3B, TRIM15, TRIM31, UBD, ISG15, RSAD2, IRF7, SAA1, CFB, IGLL5, HIST1H2BJ, IGHV3-11, IGHV3-15, CCL18 |

**Supplementary Table S6: Details of the GO Biological processes corresponding to the downregulated DEcGs from STRING database**

| GO Term Description | Observed Gene Count | Background Gene Count | FDR corrected p | Matching proteins in network (labels) |
|---|---|---|---|---|
| keratinocyte differentiation | 81 | 268 | 8.42E-34 | TGM1, TGM5, CNFN, WNT16, KRT32, PI3, EREG, KRT13, KLK12, KRT33B, KRT1, KRT6C, KRT6B, DSG1, DSC1, KRT84, CSTA, KRT10, DSC2, CERS3, KRT72, ACER1, SCEL, KRT78, SPRR1B, KRT73, KRT2, LCE1D, RPTN, SPINK6, KRT79, KRT36, KRT76, SPRR3, SPRR4, LCE5A, LCE1F, LCE1A, LCE3A, KLK5, PPL, KRT77, SPINK5, LCE1B, KRTAP1-5, S100A7, LOR, SPRR2G, SPRR2E, SPRR2B, SPRR2D, SPRR1A, IVL, LCE1E, LCE2A, LCE2B, LCE2C, LCE2D, LCE3D, LCE3E, FLG, HRNR, CDSN, ANXA1, SPINK9, DSP, KRT6A, TGM3, KRTAP3-2, KLK14, SPRR2A, KRT34, LIPK, LIPM, LIPN, LCE6A, KRT3, SPRR2F, KRT4, KLK13, LCE1C |
| keratinization | 75 | 226 | 1.35E-33 | TGM1, TGM5, CNFN, KRT32, PI3, KRT13, KLK12, KRT33B, KRT1, KRT6C, KRT6B, DSG1, DSC1, KRT84, CSTA, KRT10, DSC2, CERS3, KRT72, KRT78, SPRR1B, KRT73, KRT2, LCE1D, RPTN, SPINK6, KRT79, KRT36, KRT76, SPRR3, SPRR4, LCE5A, LCE1F, LCE1A, LCE3A, KLK5, PPL, KRT77, SPINK5, LCE1B, KRTAP1-5, LOR, SPRR2G, SPRR2E, SPRR2B, SPRR2D, SPRR1A, IVL, LCE1E, LCE2A, LCE2B, LCE2C, LCE2D, LCE3D, LCE3E, FLG, HRNR, CDSN, SPINK9, DSP, KRT6A, TGM3, KRTAP3-2, KLK14, SPRR2A, KRT34, LIPK, LIPM, LIPN, LCE6A, KRT3, SPRR2F, KRT4, KLK13, LCE1C |
| epidermis development | 97 | 419 | 2.07E-33 | TGM1, TGM5, CNFN, WNT16, KRT32, GLI1, IGFBP5, PI3, EREG, KRT13, KLK12, KRT33B, KRT1, KRT6C, KRT6B, EMP1, DSG1, DSC1, KRT84, CSTA, FGF10, FGF7, KRT10, DSC2, CERS3, HOXB13, KRT72, ACER1, SCEL, SOSTDC1, KRT78, SPRR1B, KRT73, KRT2, LCE1D, RPTN, SPINK6, KRT79, KRT36, KRT76, SPRR3, SPRR4, LCE5A, LCE1F, LCE1A, LCE3A, KLK5, KRTDAP, PPL, COL17A1, KRT77, SPINK5, LCE1B, KRTAP1-5, S100A7, LOR, SPRR2G, SPRR2E, SPRR2B, SPRR2D, SPRR1A, IVL, LCE1E, C1orf68, LCE2A, LCE2B, LCE2C, LCE2D, LCE3D, LCE3E, FLG, HRNR, KLF4, CDSN, ANXA1, SPINK9, DSP, KRT6A, TGM3, FLG2, KRTAP3-2, KLK14, KLK7, SPRR2A, KRT34, EDA2R, LIPK, LIPM, LIPN, SVEP1, LCE6A, KRT3, SPRR2F, KRT4, KLK13, LCE1C, CRABP2 |
| cornification | 56 | 113 | 8.14E-32 | TGM1, TGM5, KRT32, PI3, KRT13, KLK12, KRT33B, KRT1, KRT6C, KRT6B, DSG1, DSC1, KRT84, CSTA, KRT10, DSC2, CERS3, KRT72, KRT78, SPRR1B, KRT73, KRT2, RPTN, SPINK6, KRT79, KRT36, KRT76, SPRR3, LCE1A, KLK5, PPL, KRT77, SPINK5, LOR, SPRR2G, SPRR2E, SPRR2B, SPRR2D, SPRR1A, IVL, LCE3D, FLG, CDSN, SPINK9, DSP, KRT6A, KLK14, SPRR2A, KRT34, LIPK, LIPM, LIPN, KRT3, SPRR2F, KRT4, KLK13 |
| epidermal cell differentiation | 83 | 315 | 9.48E-32 | TGM1, TGM5, CNFN, WNT16, KRT32, GLI1, PI3, EREG, KRT13, KLK12, KRT33B, KRT1, KRT6C, KRT6B, DSG1, DSC1, KRT84, CSTA, KRT10, DSC2, CERS3, KRT72, ACER1, SCEL, KRT78, SPRR1B, KRT73, KRT2, LCE1D, RPTN, SPINK6, KRT79, KRT36, KRT76, SPRR3, SPRR4, LCE5A, LCE1F, LCE1A, LCE3A, KLK5, PPL, KRT77, SPINK5, LCE1B, KRTAP1-5, S100A7, LOR, SPRR2G, SPRR2E, SPRR2B, SPRR2D, SPRR1A, IVL, LCE1E, LCE2A, LCE2B, LCE2C, LCE2D, LCE3D, LCE3E, FLG, HRNR, KLF4, CDSN, ANXA1, SPINK9, DSP, KRT6A, TGM3, KRTAP3-2, KLK14, SPRR2A, KRT34, LIPK, LIPM, LIPN, LCE6A, KRT3, SPRR2F, KRT4, KLK13, LCE1C |

| | | | | |
|---|---|---|---|---|
| skin development | 90 | 382 | 1.52E-31 | TGM1, TGM5, CNFN, WNT16, KRT32, IGFBP5, PI3, EREG, KRT13, KLK12, ALOX12, KRT33B, KRT1, KRT6C, KRT6B, DSG1, DSC1, KRT84, CSTA, FGF10, FGF7, KRT10, DSC2, CERS3, KRT72, ACER1, SCEL, SOSTDC1, KRT78, SPRR1B, KRT73, KRT2, ALOXE3, ALOX12B, ASPRV1, LCE1D, RPTN, SPINK6, KRT79, KRT36, KRT76, SPRR3, SPRR4, LCE5A, LCE1F, LCE1A, LCE3A, KLK5, PPL, KRT77, SPINK5, LCE1B, KRTAP1-5, S100A7, LOR, SPRR2G, SPRR2E, SPRR2B, SPRR2D, SPRR1A, IVL, LCE1E, LCE2A, LCE2B, LCE2C, LCE2D, LCE3D, LCE3E, FLG, HRNR, CDSN, ANXA1, SPINK9, DSP, KRT6A, TGM3, FLG2, KRTAP3-2, KLK14, SPRR2A, KRT34, LIPK, LIPM, LIPN, LCE6A, KRT3, SPRR2F, KRT4, KLK13, LCE1C |
| tissue development | 178 | 1760 | 7.08E-22 | DCN, TGM1, TGFBR3, TGM5, SFRP1, CNFN, WNT16, KRT32, CCND1, DGAT2, GLI1, FGF6, GPLD1, ISL1, IGFBP5, DUSP1, EGR1, PI3, PTGIS, EREG, KRT13, STATH, KLK12, SALL1, KRT33B, KRT1, KRT6C, KRT6B, EMP1, DSG1, DSC1, SEMA6A, KRT84, TMOD1, SGCA, MEOX2, CHRDL2, CSTA, FGF2, FGF10, FGF7, RHCG, KRT10, GDF7, RSPO2, DSC2, BARX2, GJA1, CERS3, HOXB13, ACTC1, KRT72, TCF23, CLEC3B, BMPER, COL14A1, ACER1, SCEL, IGF1, SEMA3E, SOSTDC1, FOS, KRT78, SPRR1B, KRT73, SULT1B1, TBX5, KRT2, LCE1D, GCNT4, RPTN, SEMA6D, SPINK6, PGR, NOG, KRT79, KRT36, KRT76, SPRR3, SPRR4, SLC24A3, LCE5A, LCE1F, LCE1A, PDE2A, LCE3A, EDN3, KLK5, ZBTB16, KRTDAP, PPL, COL17A1, EYA1, KRT77, APOD, ARC, PLN, MYOZ1, HAND2, SPINK5, LCE1B, TNNI1, KRTAP1-5, LGR6, TCF21, S100A7, LOR, SPRR2G, SPRR2E, SPRR2B, SPRR2D, SPRR1A, IVL, LCE1E, C1orf68, LCE2A, LCE2B, LCE2C, LCE2D, LCE3D, LCE3E, FLG, HRNR, EPHA7, AKNAD1, KLF4, AR, CDSN, KLK6, ANXA1, SPINK9, NPHS1, NR0B1, RBM24, VIT, CYP1A1, DSP, KRT6A, ADRA1A, BNC2, TGM3, GJB2, FLG2, LAMB4, KRTAP3-2, KLK14, KLK7, SPRR2A, KRT34, NDRG4, FAT4, AREG, EDA2R, MYH11, SOX10, LIPK, LIPM, LIPN, SVEP1, FOXP2, DLX3, CYR61, MYOCD, ESR1, LCE6A, KRT3, SPRR2F, KCNAB1, ACTN2, KRT4, TPPP3, GREB1L, KLK13, LCE1C, GREM1, UPK1A, NR4A3, CRABP2 |
| epithelial cell differentiation | 101 | 673 | 7.55E-22 | TGM1, TGM5, CNFN, WNT16, KRT32, GLI1, PI3, EREG, KRT13, KLK12, SALL1, KRT33B, KRT1, KRT6C, KRT6B, DSG1, DSC1, KRT84, TMOD1, CSTA, FGF10, RHCG, KRT10, GDF7, DSC2, CERS3, HOXB13, KRT72, ACER1, SCEL, KRT78, SPRR1B, KRT73, SULT1B1, KRT2, LCE1D, RPTN, SPINK6, PGR, KRT79, KRT36, KRT76, SPRR3, SPRR4, LCE5A, LCE1F, LCE1A, PDE2A, LCE3A, KLK5, PPL, KRT77, SPINK5, LCE1B, KRTAP1-5, TCF21, S100A7, LOR, SPRR2G, SPRR2E, SPRR2B, SPRR2D, SPRR1A, IVL, LCE1E, LCE2A, LCE2B, LCE2C, LCE2D, LCE3D, LCE3E, FLG, HRNR, KLF4, AR, CDSN, ANXA1, SPINK9, NPHS1, NR0B1, CYP1A1, DSP, KRT6A, TGM3, KRTAP3-2, KLK14, SPRR2A, KRT34, LIPK, LIPM, LIPN, DLX3, ESR1, LCE6A, KRT3, SPRR2F, KRT4, KLK13, LCE1C, GREM1, UPK1A |

| | | | | |
|---|---|---|---|---|
| system development | 318 | 4426 | 9.69E-19 | OTC, DCN, NNAT, CTTNBP2, ACTL6B, SPP2, TGM1, TGFBR3, PLA2G3, TGM5, SFRP1, CRX, CNFN, WNT16, KRT32, EFNB3, CCND1, DGAT2, GLI1, FGF6, AICDA, GPLD1, ISL1, GNAT1, IGFBP5, ANGPTL1, C2orf40, MYOT, EGR1, PI3, PTGIS, EREG, KRT13, FLRT1, STATH, KLK12, CMA1, SALL1, ALOX12, KRT33B, KRT1, KRT6C, KRT6B, EPO, DCLK1, DSG1, DSC1, SEMA6A, KRT84, TMOD1, USP2, SLC6A4, SGCA, MEOX2, PCSK2, RPS6KA6, ASPA, CHRDL2, LHX4, GABRA4, CSTA, FGF2, FGF10, KCNC1, SRRM4, FGF7, ASXL3, SERPINB12, KRT10, SLC6A3, PROK1, GDF7, STAR, RSPO2, LRRC4C, SCN2B, GPM6A, IL18, DSC2, BARX2, GRID2, GJA1, CERS3, ADAMTS5, TIMP4, HOXB13, ACTC1, KRT72, LHCGR, TCF23, CLEC3B, HPGD, BMPER, CSMD3, COL14A1, FIGF, NELL1, NYAP1, ACER1, SCEL, IGF1, SEMA3E, SOSTDC1, FOS, KRT78, SPRR1B, KRT73, CTNND2, F2, RBFOX1, TBX5, FOSL1, KRT2, MAL, WFIKKN2, SNTG2, MOBP, PDLIM2, CHRNA9, ALOXE3, ALOX12B, ASPRV1, LCE1D, GCNT4, RPTN, EGR3, GHRHR, RIT2, FAIM2, SEMA6D, SPINK6, PGR, NOG, KRT79, KRT36, PCP4, KRT76, SPRR3, CNTN2, OPCML, BCAN, SPRR4, SLIT3, SPNS2, SLC24A3, LCE5A, LCE1F, LCE1A, PDE2A, LCE3A, CXCL14, EDN3, KLK5, ZBTB16, PPL, STMN4, EYA1, KRT77, PRELP, APOD, GFRA1, BTG4, GJB6, ARC, ARG1, ANK2, COL9A1, PLN, RFX4, MYOZ1, HAND2, SPINK5, LCE1B, TNNI1, NPY1R, NRG2, KRTAP1-5, LGR6, BRINP3, PAPPA2, DNM3, TCF21, APOA2, SPTA1, NKX6-2, S100A7, LOR,<br>SPRR2G, SPRR2E, SPRR2B, SPRR2D, SPRR1A, IVL, LCE1E, LCE2A, LCE2B, LCE2C, LCE2D, LCE3D, LCE3E, FLG, HRNR, EPHA7, RBM20, AKNAD1, BAI3, CHRDL1, NOX1, NAP1L2, PCDH15, SLC46A2, KLF4, AR, ZC4H2, F7, C6orf25, CDSN, ANGPTL7, KLK6, ANXA1, HIF3A, SPINK9, MAOB, NPHS1, NR0B1, CNTFR, RBM24, VIT, PTCHD1, CYP1A1, DSP, KRT6A, ALOX15B, TUBB2A, ADRA1A, BNC2, PTPRD, TGM3, GJB2, CPNE9, FLG2, LAMB4, STAB2, EPHA6, KRTAP3-2, KLK14, KLK8, SPRR2A, KRT34, FHL1, NDRG4, FAT4, SYNPO2L, TRPS1, AREG, AFP, CXCL12, MYH11, SOX10, NAV3, GRIK1, LIPK, LIPM, LIPN, POU6F2, CTNNA2, ABCB5, SVEP1, NRXN1, FOXP2, DLX3, ATCAY, KIF5C, MYT1L, NTF3, CYR61, CHRM2, MYO16, MYOCD, TENM1, ESR1, GAS7, LCE6A, EPGN, FREM1, KRT3, SPRR2F, BOC, LSAMP, EVX1, KCNAB1, GABRA2, DIO3, TENM2, CRYAB, ZIC4, TRPM1, CPNE6, NR4A1, ACTN2, KRT4, PPFIA2, TPPP3, ADCYAP1, RBFOX3, GDF10, GREB1L, PLK5, KLK13, LCE1C, NOS1, GREM1, NCAM1, EPHA5, PLP1, NR4A3, CRABP2, CDHR1 |
| anatomical structure development | 362 | 5402 | 1.84E-17 | OTC, DCN, NNAT, CTTNBP2, ACTL6B, SPP2, ROPN1, TGM1, TGFBR3, PLA2G3, TGM5, SFRP1, CRX, CNFN, WNT16, KRT32, EFNB3, GNRHR, CCND1, DGAT2, GLI1, FGF6, AICDA, GPLD1, ISL1, GNAT1, IGFBP5, ANGPTL1, C2orf40, DUSP1, MYOT, EGR1, PI3, PTGIS, EREG, KRT13, FLRT1, STATH, KLK12, CMA1, SALL1, ALOX12, KRT33B, KRT1, KRT6C, KRT6B, EPO, DCLK1, LYVE1, EMP1, DSG1, DSC1, SEMA6A, KRT84, TMOD1, CCL21, PI15, USP2, SLC6A4, SGCA, MEOX2, CDH19, PCSK2, RPS6KA6, ASPA, CHRDL2, LHX4, GABRA4, CSTA, FGF2, FGF10, KCNC1, PLCZ1, SRRM4, FGF7, RHCG, ASXL3, SERPINB12, KRT10, DMRTC2, SLC6A3, PROK1, GDF7, STAR, RSPO2, LRRC4C, SCN2B, GPM6A, IL18, DSC2, BARX2, GRID2, GJA1, CERS3, ADAMTS5, WIF1, TIMP4, HOXB13, ACTC1, KRT72, LHCGR, TCF23, CLEC3B, HPGD, BMPER, CSMD3, COL14A1, FIGF, SLC18A2, NELL1, SPATA19, TBATA, NYAP1, ACER1, SCEL, IGF1, SEMA3E, SOSTDC1, FOS, KRT78, SPRR1B, KRT73, CTNND2, F2, SULT1B1, RBFOX1, TBX5, FOSL1, KRT2, MAL, WFIKKN2, SNTG2, MOBP, EVX2, PDLIM2, CHRNA9, ALOXE3, ALOX12B, ASPRV1, LCE1D, GCNT4, RPTN, EGR3, FAM9B, GHRHR, RIT2, FAIM2, SEMA6D, SPINK6, PGR, TNP2, NOG, XKR4, KRT79, KRT36, PCP4, KRT76, SPRR3, CNTN2, OPCML, BCAN, SPRR4, SLIT3, SPNS2, SLC24A3, LCE5A, LCE1F, LCE1A, PDE2A, LCE3A, CXCL14, EDN3, KLK5, ZBTB16, KRTDAP, PPL, ELSPBP1, COL17A1, STMN4, EYA1, KRT77, PRELP, APOD, GFRA1, BTG4, SUN5, GJB6, ARC, ARG1, ANK2, COL9A1, PLN, RFX4, MYOZ1, HAND2, RXRG, SPINK5, LCE1B, TNNI1, NPY1R, |

| | | | | |
|---|---|---|---|---|
| | | | | NRG2, KRTAP1-5, LGR6, LMOD1, BRINP3, PAPPA2, DNM3, TCF21, APOA2, SPTA1, NKX6-2, S100A7, LOR, SPRR2G, SPRR2E, SPRR2B, SPRR2D, SPRR1A, IVL, LCE1E, C1orf68, LCE2A, LCE2B, LCE2C, LCE2D, LCE3D, LCE3E, FLG, HRNR, EPHA7, RBM20, AKNAD1, BAI3, CHRDL1, NOX1, NAP1L2, PCDH15, SLC46A2, KLF4, AR, ZC4H2, F7, C6orf25, CDSN, ANGPTL7, KLK6, ANXA1, HIF3A, SPINK9, MAOB, NPHS1, NR0B1, CNTFR, RBM24, VIT, PTCHD1, CYP1A1, DSP, KRT6A, ALOX15B, TUBB2A, ADRA1A, BNC2, PTPRD, TGM3, CATSPERD, GJB2, CPNE9, FLG2, LAMB4, STAB2, EPHA6, KRTAP3-2, KLK14, KLK8, KLK7, SPRR2A, KRT34, FHL1, NDRG4, FAT4, SYNPO2L, PAQR5, TRPS1, AREG, AFP, CXCL12, EDA2R, MYH11, SOX10, NAV3, GRIK1, JAM2, LIPK, LIPM, LIPN, POU6F2, CTNNA2, ABCB5, SVEP1, NRXN1, FOXP2, NECAB1, DLX3, ATCAY, KIF5C, MYT1L, NTF3, CYR61, CHRM2, MYO16, MYOCD, DDC, TENM1, ESR1, GAS7, LCE6A, EPGN, FREM1, KRT3, SPARCL1, SPRR2F, BOC, LSAMP, EVX1, KCNAB1, GABRA2, DDX4, CDH18, SPINK2, DIO3, TENM2, SCUBE2, FAT3, CRYAB, ZIC4, TRPM1, CPNE6, NR4A1, ACTN2, RAI2, KRT4, PPFIA2, HOPX, TPPP3, ADCYAP1, RBFOX3, GDF10, GREB1L, ACSBG2, PLK5, KLK13, LCE1C, NOS1, GREM1, UPK1A, NCAM1, EPHA5, PLP1, NR4A3, CRABP2, CDHR1 |
| epithelium development | 124 | 1109 | 1.89E-17 | TGM1, TGM5, SFRP1, CNFN, WNT16, KRT32, CCND1, GLI1, IGFBP5, PI3, EREG, KRT13, KLK12, SALL1, KRT33B, KRT1, KRT6C, KRT6B, DSG1, DSC1, KRT84, TMOD1, MEOX2, CSTA, FGF2, FGF10, FGF7, RHCG, KRT10, GDF7, RSPO2, DSC2, GJA1, CERS3, HOXB13, KRT72, BMPER, ACER1, SCEL, SEMA3E, SOSTDC1, KRT78, SPRR1B, KRT73, SULT1B1, TBX5, KRT2, LCE1D, RPTN, SPINK6, PGR, NOG, KRT79, KRT36, KRT76, SPRR3, SPRR4, LCE5A, LCE1F, LCE1A, PDE2A, LCE3A, KLK5, PPL, EYA1, KRT77, HAND2, SPINK5, LCE1B, KRTAP1-5, TCF21, S100A7, LOR, SPRR2G, SPRR2E, SPRR2B, SPRR2D, SPRR1A, IVL, LCE1E, LCE2A, LCE2B, LCE2C, LCE2D, LCE3D, LCE3E, FLG, HRNR, EPHA7, KLF4, AR, CDSN, ANXA1, SPINK9, NPHS1, NR0B1, CYP1A1, DSP, KRT6A, TGM3, FLG2, KRTAP3-2, KLK14, SPRR2A, KRT34, NDRG4, FAT4, AREG, SOX10, LIPK, LIPM, LIPN, DLX3, CYR61, ESR1, LCE6A, KRT3, SPRR2F, KRT4, GREB1L, KLK13, LCE1C, GREM1, UPK1A |
| multicellular organism development | 341 | 5023 | 7.54E-17 | OTC, DCN, NNAT, CTTNBP2, ACTL6B, SPP2, TGM1, TGFBR3, PLA2G3, TGM5, SFRP1, CRX, CNFN, WNT16, KRT32, EFNB3, GNRHR, CCND1, DGAT2, GLI1, FGF6, AICDA, GPLD1, ISL1, GNAT1, IGFBP5, ANGPTL1, C2orf40, DUSP1, MYOT, EGR1, PI3, PTGIS, EREG, KRT13, FLRT1, STATH, KLK12, CMA1, SALL1, ALOX12, KRT33B, KRT1, KRT6C, KRT6B, EPO, DCLK1, DSG1, DSC1, SEMA6A, KRT84, TMOD1, PI15, USP2, SLC6A4, SGCA, MEOX2, CDH19, PCSK2, RPS6KA6, ASPA, CHRDL2, LHX4, GABRA4, CSTA, FGF2, FGF10, KCNC1, PLCZ1, SRRM4, FGF7, ASXL3, SERPINB12, KRT10, SLC6A3, PROK1, GDF7, STAR, RSPO2, LRRC4C, SCN2B, GPM6A, IL18, DSC2, BARX2, GRID2, GJA1, CERS3, ADAMTS5, WIF1, TIMP4, HOXB13, ACTC1, KRT72, LHCGR, TCF23, CLEC3B, HPGD, BMPER, CSMD3, COL14A1, FIGF, SLC18A2, NELL1, SPATA19, TBATA, NYAP1, ACER1, SCEL, IGF1, SEMA3E, SOSTDC1, FOS, KRT78, SPRR1B, KRT73, CTNND2, F2, RBFOX1, TBX5, FOSL1, KRT2, MAL, WFIKKN2, SNTG2, MOBP, EVX2, PDLIM2, CHRNA9, ALOXE3, ALOX12B, ASPRV1, LCE1D, GCNT4, RPTN, EGR3, GHRHR, RIT2, FAIM2, SEMA6D, SPINK6, PGR, TNP2, NOG, KRT79, KRT36, PCP4, KRT76, SPRR3, CNTN2, OPCML, BCAN, SPRR4, SLIT3, SPNS2, SLC24A3, LCE5A, LCE1F, LCE1A, PDE2A, LCE3A, CXCL14, EDN3, KLK5, ZBTB16, PPL, STMN4, EYA1, KRT77, PRELP, APOD, GFRA1, BTG4, GJB6, ARC, ARG1, ANK2, |

| | | | | |
|---|---|---|---|---|
| | | | | COL9A1, PLN, RFX4, MYOZ1, HAND2, SPINK5, LCE1B, TNNI1, NPY1R, NRG2, KRTAP1-5, LGR6, BRINP3, PAPPA2, DNM3, TCF21, APOA2, SPTA1, NKX6-2, S100A7, LOR, SPRR2G, SPRR2E, SPRR2B, SPRR2D, SPRR1A, IVL, LCE1E, LCE2A, LCE2B, LCE2C, LCE2D, LCE3D, LCE3E, FLG, HRNR, EPHA7, RBM20, AKNAD1, BAI3, CHRDL1, NOX1, NAP1L2, PCDH15, SLC46A2, KLF4, AR, ZC4H2, F7, C6orf25, CDSN, ANGPTL7, KLK6, ANXA1, HIF3A, SPINK9, MAOB, NPHS1, NR0B1, CNTFR, RBM24, VIT, PTCHD1, CYP1A1, DSP, KRT6A, ALOX15B, TUBB2A, ADRA1A, BNC2, PTPRD, TGM3, CATSPERD, GJB2, CPNE9, FLG2, LAMB4, STAB2, EPHA6, KRTAP3-2, KLK14, KLK8, SPRR2A, KRT34, FHL1, NDRG4, FAT4, SYNPO2L, PAQR5, TRPS1, AREG, AFP, CXCL12, EDA2R, MYH11, SOX10, NAV3, GRIK1, LIPK, LIPM, LIPN, POU6F2, CTNNA2, ABCB5, SVEP1, NRXN1, FOXP2, NECAB1, DLX3, ATCAY, KIF5C, MYT1L, NTF3, CYR61, CHRM2, MYO16, MYOCD, DDC, TENM1, ESR1, GAS7, LCE6A, EPGN, FREM1, KRT3, SPRR2F, BOC, LSAMP, EVX1, KCNAB1, GABRA2, DDX4, CDH18, DIO3, TENM2, SCUBE2, FAT3, CRYAB, ZIC4, TRPM1, CPNE6, NR4A1, ACTN2, RAI2, KRT4, PPFIA2, HOPX, TPPP3, ADCYAP1, RBFOX3, GDF10, GREB1L, ACSBG2, PLK5, KLK13, LCE1C, NOS1, GREM1, NCAM1, EPHA5, PLP1, NR4A3, CRABP2, CDHR1 |
| | | | | OTC, DCN, NNAT, CTTNBP2, ACTL6B, SPP2, ROPN1, TGM1, TGFBR3, PLA2G3, ACR, TGM5, SFRP1, TULP2, CYP4F2, CRX, CNFN, WNT16, KRT32, EFNB3, GNRHR, CCND1, DGAT2, SYT10, GLI1, FGF6, AICDA, GPLD1, ISL1, GNAT1, IGFBP5, ANGPTL1, C2orf40, DUSP1, MYOT, EGR1, KIAA1045, PI3, PTGIS, EREG, MYH2, FOSB, SLURP1, KRT13, FLRT1, STATH, KLK12, CMA1, SALL1, ALOX12, KRT33B, KRT1, KRT6C, KRT6B, EPO, CRP, DCLK1, DSG1, DSC1, SEMA6A, KRT84, TMOD1, PI15, USP2, HTR3B, GUCY2C, SLC6A4, SGCA, MEOX2, CDH19, PCSK2, RPS6KA6, ASPA, CHRDL2, LHX4, GABRA4, CSTA, FGF2, FGF10, KCNC1, PLCZ1, SRRM4, FGF7, SPACA3, ASXL3, SERPINB12, KRT10, DMRTC2, SLC6A3, PROK1, HMCN1, GDF7, CORIN, CRHBP, STAR, RSPO2, LRRC4C, SCN2B, GPM6A, IL18, DSC2, BARX2, GRID2, GRIA4, GJA1, SCN3A, CERS3, ADAMTS5, WIF1, PPEF2, TIMP4, HOXB13, ACTC1, CSTB, KRT72, LHCGR, KCNJ3, TCF23, CLEC3B, HPGD, BMPER, CSMD3, COL14A1, FIGF, SLC18A2, NELL1, SPATA19, TBATA, NYAP1, LOXHD1, ACER1, SOAT2, SCEL, IGF1, SEMA3E, TACR1, SOSTDC1, FOS, KRT78, SPRR1B, KRT73, CTNND2, MYL1, TMPRSS11E, F2, GP6, RBFOX1, TBX5, FOSL1, KRT2, MAL, WFIKKN2, ARR3, SNTG2, MOBP, EVX2, PDLIM2, CHRNA9, KLK2, KLK3, ALOXE3, ALOX12B, ASPRV1, LCE1D, GCNT4, RPTN, EGR3, FAM9B, PSG11, OR4N2, CACNB2, GHRHR, CIDEA, RIT2, FAIM2, |

| | | | | |
|---|---|---|---|---|
| multicellular organismal process | 426 | 6933 | 2.21E-15 | SEMA6D, SPINK6, PGR, TNP2, NOG, KRT79, KRT36, PCP4, KRT76, SPRR3, CNTN2, OPCML, BCAN, SPRR4, SLIT3, IGDCC3, SPNS2, SLC24A3, LCE5A, TAC4, LCE1F, LCE1A, PDE2A, LCE3A, CXCL14, EDN3, KLK5, ZBTB16, PPL, ELSPBP1, STMN4, EYA1, KRT77, ADCY2, PRELP, APOD, GFRA1, BTG4, SUN5, GJB6, PTGER3, ARC, ARG1, ANK2, COL9A1, PLN, GRM7, RFX4, CACNA1G, MYOZ1, HAND2, SPINK5, LCE1B, PRSS3, TNNI1, NPY1R, NRG2, KRTAP1-5, SLC22A2, VIP, LGR6, LMOD1, BRINP3, PAPPA2, DNM3, TCF21, APOA2, SPTA1, NKX6-2, S100A7, S100A12, LOR, SPRR2G, SPRR2E, SPRR2B, SPRR2D, SPRR1A, IVL, LCE1E, LCE2A, LCE2B, LCE2C, LCE2D, LCE3D, LCE3E, FLG, HRNR, ANXA9, EPHA7, RBM20, AKNAD1, BAI3, PTGFR, CHRDL1, DUSP13, NOX1, PGC, NAP1L2, PCDH15, DES, SLC46A2, TRIM63, KLF4, AR, ZC4H2, SPIN2A, F10, F7, C6orf25, DLG2, CDSN, ANGPTL7, KLK6, ANXA1, TRPM3, CA6, HIF3A, SPINK9, MAOB, NPHS1, NR0B1, CNTFR, RBM24, VIT, PTCHD1, CYP1A1, DSP, KRT6A, ALOX15B, TUBB2A, ADRA1A, BNC2, PTPRD, TGM3, CATSPERD, KCNIP4, GJB2, CPNE9, FLG2, LAMB4, STAB2, RNF180, EPHA6, KRTAP3-2, KLK14, KLK8, SPRR2A, KRT34, FHL1, NDRG4, FAT4, SYNPO2L, MMRN1, PAQR5, TRPS1, AREG, AFP, CXCL12, EDA2R, MYH11, SOX10, NAV3, CRHR1, GRIK1, JAM2, LIPK, LIPM, LIPN, POU6F2, CTNNA2, ABCB5, SVEP1, CHRNA2, NRXN1, DGKB, FOXP2, SCN7A, NECAB1, CTNNA3, DLX3, ATCAY, SEPT5, LHFPL3, KIF5C, MYT1L, NTF3, ENDOU, CYR61, CHRM2, OTOGL, MYO16, MYOCD, DDC, TENM1, CKMT2, ESR1, GAS7, LCE6A, EPGN, FREM1, KRT3, SPRR2F, BOC, LSAMP, FAM107A, EVX1, KCNAB1, GABRA2, DDX4, CDH18, SPINK2, DIO3, GRIA1, TENM2, SCUBE2, CSMD1, FAT3, CRYAB, ZIC4, TRPM1, TRDN, CPNE6, NR4A1, ACTN2, RAI2, KRT4, RDH12, PPFIA2, HOPX, C15orf59, SPATA22, TPPP3, ADCYAP1, RBFOX3, GDF10, GREB1L, ACSBG2, PLK5, KLK13, LCE1C, NOS1, GREM1, GPIHBP1, NCAM1, EPHA5, PLP1, FXYD1, GRIA3, NR4A3, CRABP2, CDHR1 |
| animal organ development | 240 | 3197 | 8.04E-15 | OTC, DCN, NNAT, CTTNBP2, ACTL6B, TGM1, TGFBR3, TGM5, SFRP1, CRX, CNFN, WNT16, KRT32, CCND1, DGAT2, GLI1, FGF6, AICDA, GPLD1, ISL1, GNAT1, IGFBP5, EGR1, PI3, PTGIS, EREG, KRT13, STATH, KLK12, CMA1, SALL1, ALOX12, KRT33B, KRT1, KRT6C, KRT6B, EPO, DCLK1, DSG1, DSC1, SEMA6A, KRT84, TMOD1, USP2, SLC6A4, SGCA, MEOX2, CHRDL2, LHX4, CSTA, FGF2, FGF10, KCNC1, FGF7, ASXL3, SERPINB12, KRT10, SLC6A3, GDF7, STAR, RSPO2, GPM6A, DSC2, BARX2, GRID2, GJA1, CERS3, ADAMTS5, HOXB13, ACTC1, KRT72, LHCGR, TCF23, CLEC3B, HPGD, BMPER, COL14A1, ACER1, SCEL, IGF1, SEMA3E, SOSTDC1, FOS, KRT78, SPRR1B, KRT73, TBX5, FOSL1, KRT2, PDLIM2, CHRNA9, ALOXE3, ALOX12B, ASPRV1, LCE1D, GCNT4, RPTN, EGR3, GHRHR, FAIM2, SEMA6D, SPINK6, PGR, NOG, KRT79, KRT36, KRT76, SPRR3, CNTN2, BCAN, SPRR4, SLIT3, SPNS2, SLC24A3, LCE5A, LCE1F, LCE1A, PDE2A, LCE3A, CXCL14, EDN3, KLK5, ZBTB16, PPL, EYA1, KRT77, APOD, GJB6, ARG1, ANK2, COL9A1, PLN, RFX4, MYOZ1, HAND2, SPINK5, LCE1B, TNNI1, NPY1R, NRG2, KRTAP1-5, PAPPA2, TCF21, APOA2, SPTA1, NKX6-2, S100A7, LOR, SPRR2G, SPRR2E, SPRR2B, SPRR2D, SPRR1A, IVL, LCE1E, LCE2A, LCE2B, LCE2C, LCE2D, LCE3D, LCE3E, FLG, HRNR, EPHA7, RBM20, AKNAD1, CHRDL1, PCDH15, SLC46A2, KLF4, AR, ZC4H2, F7, C6orf25, CDSN, ANGPTL7, ANXA1, SPINK9, MAOB, NPHS1, NR0B1, CNTFR, RBM24, VIT, PTCHD1, CYP1A1, DSP, KRT6A, ALOX15B, ADRA1A, BNC2, TGM3, GJB2, FLG2, LAMB4, KRTAP3-2, KLK14, SPRR2A, KRT34, FHL1, NDRG4, FAT4, SYNPO2L, AREG, AFP, CXCL12, MYH11, SOX10, LIPK, LIPM, LIPN, CTNNA2, ABCB5, NRXN1, FOXP2, DLX3, CYR61, MYO16, MYOCD, ESR1, LCE6A, FREM1, KRT3, SPRR2F, EVX1, KCNAB1, DIO3, CRYAB, TRPM1, ACTN2, KRT4, TPPP3, ADCYAP1, GREB1L, KLK13, LCE1C, GREM1, EPHA5, PLP1, NR4A3 |

| | | | | |
|---|---|---|---|---|
| developmental process | 368 | 5841 | 1.26E-13 | OTC, DCN, NNAT, CTTNBP2, ACTL6B, SPP2, ROPN1, TGM1, TGFBR3, FGF22, PLA2G3, TGM5, SFRP1, CRX, CNFN, WNT16, KRT32, EFNB3, GNRHR, CCND1, DGAT2, GLI1, FGF6, AICDA, GPLD1, ISL1, GNAT1, IGFBP5, ANGPTL1, C2orf40, DUSP1, MYOT, EGR1, PI3, PTGIS, EREG, KRT13, FLRT1, STATH, KLK12, CMA1, SALL1, ALOX12, KRT33B, KRT1, KRT6C, KRT6B, EPO, DCLK1, LYVE1, EMP1, DSG1, DSC1, SEMA6A, KRT84, TMOD1, CCL21, PI15, USP2, UNC13C, SLC6A4, SGCA, MEOX2, CDH19, PCSK2, RPS6KA6, ASPA, CHRDL2, LHX4, GABRA4, CSTA, FGF2, FGF10, KCNC1, PLCZ1, SRRM4, FGF7, RHCG, ASXL3, SERPINB12, KRT10, DMRTC2, SLC6A3, PROK1, GDF7, STAR, RSPO2, LRRC4C, SCN2B, GPM6A, IL18, DSC2, BARX2, GRID2, GJA1, CERS3, ADAMTS5, WIF1, TIMP4, HOXB13, ACTC1, KRT72, LHCGR, TCF23, CLEC3B, HPGD, BMPER, CSMD3, COL14A1, FIGF, SLC18A2, NELL1, SPATA19, TBATA, NYAP1, ACER1, SOAT2, SCEL, IGF1, SEMA3E, SOSTDC1, FOS, KRT78, SPRR1B, KRT73, CTNND2, F2, SULT1B1, RBFOX1, DAPL1, TBX5, FOSL1, KRT2, MAL, WFIKKN2, SNTG2, MOBP, EVX2, PDLIM2, CHRNA9, ALOXE3, ALOX12B, ASPRV1, LCE1D, GCNT4, RPTN, EGR3, FAM9B, GHRHR, RIT2, FAIM2, SEMA6D, SPINK6, PGR, TNP2, NOG, XKR4, KRT79, KRT36, PCP4, KRT76, SPRR3, CNTN2, OPCML, BCAN, SPRR4, SLIT3, SPNS2, SLC24A3, LCE5A, LCE1F, LCE1A, PDE2A, LCE3A, CXCL14, EDN3, KLK5, ZBTB16, KRTDAP, PPL, ELSPBP1, COL17A1, STMN4, EYA1, KRT77, PRELP, APOD, GFRA1, BTG4, SUN5, GJB6, ARC, ARG1, ANK2, COL9A1, PLN, RFX4, MYOZ1, HAND2, RXRG, SPINK5, LCE1B, TNNI1, NPY1R, NRG2, KRTAP1-5, LGR6, LMOD1, BRINP3, PAPPA2, DNM3, TCF21, APOA2, SPTA1, NKX6-2, S100A7, LOR, SPRR2G, SPRR2E, SPRR2B, SPRR2D, SPRR1A, IVL, LCE1E, C1orf68, LCE2A, LCE2B, LCE2C, LCE2D, LCE3D, LCE3E, FLG, HRNR, EPHA7, RBM20, AKNAD1, BAI3, CHRDL1, DUSP13, NOX1, NAP1L3, NAP1L2, PCDH15, SLC46A2, KLF4, AR, ZC4H2, F7, C6orf25, CDSN, ANGPTL7, KLK6, ANXA1, HIF3A, SPINK9, MAOB, NPHS1, NR0B1, CNTFR, RBM24, VIT, PTCHD1, CYP1A1, DSP, KRT6A, ALOX15B, TUBB2A, ADRA1A, BNC2, PTPRD, TGM3, CATSPERD, GJB2, CPNE9, FLG2, LAMB4, STAB2, EPHA6, KRTAP3-2, KLK14, KLK8, KLK7, SPRR2A, KRT34, FHL1, NDRG4, FAT4, SYNPO2L, PAQR5, TRPS1, AREG, AFP, CXCL12, EDA2R, MYH11, SOX10, NAV3, GRIK1, JAM2, LIPK, LIPM, LIPN, POU6F2, CTNNA2, ABCB5, SVEP1, NRXN1, FOXP2, NECAB1, DLX3, ATCAY, KIF5C, MYT1L, NTF3, CYR61, CHRM2, MYO16, MYOCD, DDC, TENM1, ESR1, GAS7, LCE6A, EPGN, FREM1, KRT3, SPARCL1, SPRR2F, BOC, LSAMP, EVX1, KCNAB1, GABRA2, DDX4, CDH18, SPINK2, DIO3, TENM2, SCUBE2, FAT3, CRYAB, ZIC4, TRPM1, CPNE6, NR4A1, ACTN2, RAI2, KRT4, PPFIA2, HOPX, TPPP3, ADCYAP1, RBFOX3, GDF10, GREB1L, ACSBG2, PLK5, KLK13, LCE1C, NOS1, GREM1, UPK1A, NCAM1, EPHA5, PLP1, NR4A3, CRABP2, CDHR1 |

| | | | | |
|---|---|---|---|---|
| cell differentiation | 259 | 3702 | 9.27E-13 | NNAT, ACTL6B, ROPN1, TGM1, TGFBR3, FGF22, PLA2G3, TGM5, SFRP1, CRX, CNFN, WNT16, KRT32, EFNB3, CCND1, GLI1, FGF6, AICDA, GPLD1, ISL1, GNAT1, IGFBP5, MYOT, EGR1, PI3, EREG, KRT13, FLRT1, KLK12, SALL1, KRT33B, KRT1, KRT6C, KRT6B, EPO, DCLK1, DSG1, DSC1, SEMA6A, KRT84, TMOD1, CCL21, SLC6A4, ASPA, CHRDL2, LHX4, CSTA, FGF2, FGF10, SRRM4, RHCG, SERPINB12, KRT10, DMRTC2, GDF7, STAR, RSPO2, LRRC4C, GPM6A, DSC2, BARX2, GRID2, GJA1, CERS3, HOXB13, ACTC1, KRT72, TCF23, CSMD3, COL14A1, FIGF, NELL1, SPATA19, TBATA, NYAP1, ACER1, SOAT2, SCEL, IGF1, SEMA3E, FOS, KRT78, SPRR1B, KRT73, CTNND2, F2, SULT1B1, DAPL1, TBX5, KRT2, MAL, WFIKKN2, ALOXE3, LCE1D, RPTN, FAM9B, GHRHR, RIT2, FAIM2, SEMA6D, SPINK6, PGR, TNP2, NOG, KRT79, KRT36, PCP4, KRT76, SPRR3, CNTN2, OPCML, SPRR4, SLIT3, LCE5A, LCE1F, LCE1A, PDE2A, LCE3A, EDN3, KLK5, ZBTB16, KRTDAP, PPL, ELSPBP1, STMN4, EYA1, KRT77, APOD, GFRA1, BTG4, SUN5, ARC, ANK2, MYOZ1, HAND2, RXRG, SPINK5, LCE1B, KRTAP1-5, LGR6, LMOD1, BRINP3, DNM3, TCF21, SPTA1, NKX6-2, S100A7, LOR, SPRR2G, SPRR2E, SPRR2B, SPRR2D, SPRR1A, IVL, LCE1E, LCE2A, LCE2B, LCE2C, LCE2D, LCE3D, LCE3E, FLG, HRNR, EPHA7, AKNAD1, BAI3, CHRDL1, NAP1L3, NAP1L2, KLF4, AR, ZC4H2, C6orf25, CDSN, KLK6, ANXA1, SPINK9, NPHS1, NR0B1, RBM24, VIT, CYP1A1, DSP, KRT6A, TUBB2A, ADRA1A, PTPRD, TGM3, CATSPERD, CPNE9, LAMB4, EPHA6, KRTAP3-2, KLK14, KLK8, SPRR2A, KRT34, FHL1, NDRG4, FAT4, SYNPO2L, PAQR5, AREG, CXCL12, EDA2R, MYH11, SOX10, NAV3, JAM2, LIPK, LIPM, LIPN, POU6F2, CTNNA2, ABCB5, NRXN1, DLX3, ATCAY, KIF5C, MYT1L, NTF3, CYR61, MYO16, MYOCD, TENM1, ESR1, GAS7, LCE6A, KRT3, SPRR2F, BOC, EVX1, KCNAB1, DDX4, SPINK2, DIO3, TENM2, TRPM1, CPNE6, NR4A1, ACTN2, KRT4, PPFIA2, HOPX, ADCYAP1, GDF10, ACSBG2, PLK5, KLK13, LCE1C, NOS1, GREM1, UPK1A, NCAM1, EPHA5, PLP1, NR4A3, CRABP2, CDHR1 |
| cellular developmental process | 261 | 3757 | 1.45E-12 | NNAT, ACTL6B, ROPN1, TGM1, TGFBR3, FGF22, PLA2G3, TGM5, SFRP1, CRX, CNFN, WNT16, KRT32, EFNB3, CCND1, GLI1, FGF6, AICDA, GPLD1, ISL1, GNAT1, IGFBP5, C2orf40, MYOT, EGR1, PI3, EREG, KRT13, FLRT1, KLK12, SALL1, KRT33B, KRT1, KRT6C, KRT6B, EPO, DCLK1, DSG1, DSC1, SEMA6A, KRT84, TMOD1, CCL21, SLC6A4, ASPA, CHRDL2, LHX4, CSTA, FGF2, FGF10, SRRM4, RHCG, SERPINB12, KRT10, DMRTC2, GDF7, STAR, RSPO2, LRRC4C, GPM6A, DSC2, BARX2, GRID2, GJA1, CERS3, HOXB13, ACTC1, KRT72, TCF23, CSMD3, COL14A1, FIGF, NELL1, SPATA19, TBATA, NYAP1, ACER1, SOAT2, SCEL, IGF1, SEMA3E, FOS, KRT78, SPRR1B, KRT73, CTNND2, F2, SULT1B1, DAPL1, TBX5, KRT2, MAL, WFIKKN2, ALOXE3, LCE1D, RPTN, FAM9B, GHRHR, RIT2, FAIM2, SEMA6D, SPINK6, PGR, TNP2, NOG, KRT79, KRT36, PCP4, KRT76, SPRR3, CNTN2, OPCML, SPRR4, SLIT3, LCE5A, LCE1F, LCE1A, PDE2A, LCE3A, EDN3, KLK5, ZBTB16, KRTDAP, PPL, ELSPBP1, STMN4, EYA1, KRT77, PRELP, APOD, GFRA1, BTG4, SUN5, ARC, ANK2, MYOZ1, HAND2, RXRG, SPINK5, LCE1B, KRTAP1-5, LGR6, LMOD1, BRINP3, DNM3, TCF21, SPTA1, NKX6-2, S100A7, LOR, SPRR2G, SPRR2E, SPRR2B, SPRR2D, SPRR1A, IVL, LCE1E, LCE2A, LCE2B, LCE2C, LCE2D, LCE3D, LCE3E, FLG, HRNR, EPHA7, AKNAD1, BAI3, CHRDL1, NAP1L3, NAP1L2, KLF4, AR, ZC4H2, C6orf25, CDSN, KLK6, ANXA1, SPINK9, NPHS1, NR0B1, RBM24, VIT, CYP1A1, DSP, KRT6A, TUBB2A, ADRA1A, PTPRD, TGM3, CATSPERD, CPNE9, LAMB4, EPHA6, KRTAP3-2, KLK14, KLK8, SPRR2A, KRT34, FHL1, NDRG4, FAT4, SYNPO2L, PAQR5, AREG, CXCL12, EDA2R, MYH11, SOX10, NAV3, JAM2, LIPK, LIPM, LIPN, POU6F2, CTNNA2, ABCB5, NRXN1, DLX3, ATCAY, KIF5C, MYT1L, NTF3, CYR61, MYO16, MYOCD, TENM1, ESR1, GAS7, LCE6A, KRT3, SPRR2F, BOC, EVX1, KCNAB1, DDX4, SPINK2, DIO3, TENM2, TRPM1, CPNE6, NR4A1, ACTN2, KRT4, PPFIA2, HOPX, ADCYAP1, GDF10, ACSBG2, PLK5, KLK13, LCE1C, NOS1, GREM1, UPK1A, NCAM1, EPHA5, PLP1, NR4A3, CRABP2, CDHR1 |
| peptide cross-linking | 18 | 34 | 1.15E-09 | DCN, TGM1, TGM5, PI3, KRT1, CSTA, KRT10, TGM4, SPRR1B, KRT2, LOR, SPRR2E, SPRR1A, IVL, FLG, ANXA1, DSP, TGM3 |

| | | | | |
|---|---|---|---|---|
| cell death | 91 | 1091 | 6.04E-06 | TGM1, TGM5, GML, LGALS13, KRT32, PI3, PTGIS, KRT13, KLK12, KRT33B, KRT1, KRT6C, KRT6B, EPO, EMP1, DSG1, DSC1, SEMA6A, KRT84, CSTA, KRT10, DSC2, GJA1, CERS3, KRT72, GSDMA, KRT78, SPRR1B, KRT73, DAPL1, KRT2, MAL, RPTN, CIDEA, FAIM2, SPINK6, PEG3, XKR4, KRT79, KRT36, KRT76, SPRR3, SLIT3, LCE1A, KLK5, ZBTB16, PPL, KRT77, GJB6, PTGER3, HAND2, SPINK5, LOR, SPRR2G, SPRR2E, SPRR2B, SPRR2D, SPRR1A, IVL, LCE3D, FLG, BNIPL, EPHA7, CDSN, HIF3A, SPINK9, DSP, KRT6A, ALOX15B, ADRA1A, CIDEC, KLK14, KLK8, SPRR2A, KRT34, EDA2R, LIPK, LIPM, LIPN, NSG1, ATCAY, CYR61, KRT3, SPRR2F, DIO3, TP53AIP1, CRYAB, KRT4, BLID, KLK13, GREM1 |
| programmed cell death | 88 | 1054 | 9.99E-06 | TGM1, TGM5, GML, LGALS13, KRT32, PI3, PTGIS, KRT13, KLK12, KRT33B, KRT1, KRT6C, KRT6B, EPO, DSG1, DSC1, SEMA6A, KRT84, CSTA, KRT10, DSC2, GJA1, CERS3, KRT72, GSDMA, KRT78, SPRR1B, KRT73, DAPL1, KRT2, MAL, RPTN, CIDEA, FAIM2, SPINK6, PEG3, XKR4, KRT79, KRT36, KRT76, SPRR3, SLIT3, LCE1A, KLK5, ZBTB16, PPL, KRT77, GJB6, HAND2, SPINK5, LOR, SPRR2G, SPRR2E, SPRR2B, SPRR2D, SPRR1A, IVL, LCE3D, FLG, BNIPL, EPHA7, CDSN, HIF3A, SPINK9, DSP, KRT6A, ALOX15B, ADRA1A, CIDEC, KLK14, SPRR2A, KRT34, EDA2R, LIPK, LIPM, LIPN, NSG1, ATCAY, CYR61, KRT3, SPRR2F, DIO3, TP53AIP1, CRYAB, KRT4, BLID, KLK13, GREM1 |
| cell adhesion | 77 | 925 | 9.39E-05 | MYOT, LYPD3, SLURP1, FLRT1, LYVE1, DSG1, DSC1, CCL21, CDH19, PCDH10, CSTA, HMCN1, CRNN, LRRC4C, DSC2, GRID2, CLDN17, COL14A1, LRFN5, CTNND2, SPACA4, SIGLECL1, CNTN2, OPCML, BCAN, CRISP2, COL17A1, DPT, CADM3, WISP3, ANXA9, WISP2, PCDH11X, PCDH15, OMD, DLG2, ITGBL1, CDSN, ANXA1, PCDH9, NPHS1, DSP, PTPRD, FLG2, LAMB4, STAB2, FAT4, PCDHGA2, MMRN1, CXCL12, COL28A1, CLDN8, JAM2, CADM2, CTNNA2, SVEP1, NRXN1, PCDH20, CTNNA3, CYR61, CNTNAP5, TENM1, TNXB, XG, FREM1, SPARCL1, BOC, LSAMP, CDH18, PCDHGB6, TENM2, FAT3, ACTN2, PPFIA2, CNTNAP4, NCAM1, CDHR1 |
| response to ketone | 28 | 199 | 0.00019 | TGFBR3, SFRP1, CCND1, DUSP1, FOSB, EPO, DSG1, CCL21, STAR, HOXB13, FOS, FOSL1, FIBIN, DEFB104B, SLIT3, ADCY2, ARG1, PLN, PTGFR, P2RY4, KLF4, AR, F7, MAOB, CATSPERD, GJB2, SOX10, FOXP2 |
| multicellular organism process | 28 | 213 | 0.00063 | ACR, IGFBP5, PTGIS, FOSB, EPO, DSG1, SLC6A4, CORIN, CRHBP, GJA1, TCF23, HPGD, FOS, FOSL1, PSG11, PGR, TNP2, ARG1, PTGFR, AR, CYP1A1, GJB2, KLK14, CRHR1, ENDOU, ESR1, TPPP3, ADCYAP1 |
| anatomical structure morphogenesis | 141 | 2165 | 0.00071 | DCN, TGFBR3, PLA2G3, SFRP1, CRX, WNT16, EFNB3, GLI1, FGF6, GPLD1, ISL1, GNAT1, IGFBP5, ANGPTL1, DUSP1, MYOT, EREG, FLRT1, SALL1, DCLK1, LYVE1, SEMA6A, TMOD1, SLC6A4, MEOX2, CDH19, LHX4, FGF2, FGF10, FGF7, ASXL3, SLC6A3, PROK1, GDF7, RSPO2, GPM6A, IL18, BARX2, GRID2, GJA1, ADAMTS5, HOXB13, ACTC1, HPGD, BMPER, COL14A1, FIGF, NYAP1, SEMA3E, SOSTDC1, CTNND2, TBX5, EVX2, CHRNA9, GCNT4, EGR3, FAIM2, SEMA6D, PGR, NOG, CNTN2, SLIT3, KLK5, ZBTB16, EYA1, APOD, GFRA1, GJB6, ARC, ARG1, ANK2, COL9A1, MYOZ1, HAND2, TNNI1, NPY1R, LGR6, LMOD1, PAPPA2, TCF21, SPTA1, S100A7, EPHA7, BAI3, NOX1, KLF4, AR, C6orf25, CDSN, HIF3A, NPHS1, VIT, DSP, KRT6A, TGM3, FLG2, LAMB4, STAB2, EPHA6, KLK14, KLK8, FHL1, NDRG4, FAT4, SYNPO2L, AREG, CXCL12, MYH11, SOX10, JAM2, CTNNA2, SVEP1, NRXN1, DLX3, KIF5C, NTF3, CYR61, MYO16, MYOCD, ESR1, GAS7, EPGN, FREM1, BOC, CDH18, SPINK2, DIO3, TENM2, CRYAB, TRPM1, NR4A1, ACTN2, HOPX, GREB1L, NOS1, GREM1, NCAM1, EPHA5, NR4A3, CRABP2, CDHR1 |

| | | | | |
|---|---|---|---|---|
| cell-cell signaling | 86 | 1145 | 0.0008 | SFRP1, WNT16, EFNB3, CCND1, SYT10, GLI1, C2orf40, EREG, CCL21, HTR3B, UNC13C, TRHDE, GABRA4, FGF2, FGF10, FGF7, SLC6A3, CRHBP, RSPO2, SCN2B, IL18, GRID2, GRIA4, GJA1, WIF1, KCNJ3, SLC18A2, SOSTDC1, CTNND2, TBX5, CHRNA9, DLGAP1, EGR3, CACNB2, RIT2, SYT9, PGR, CXCL14, EDN3, GJB6, RSPO1, ANK2, GRM7, CACNA1G, SLC22A2, VIP, LGR6, WISP3, S100A12, ANXA9, WISP2, KLF4, AR, DLG2, ANXA1, PTCHD1, ADRA1A, PTPRD, GJB2, RPH3A, AREG, CRHR1, GRIK1, CHRNA2, NRXN1, NSG1, SEPT5, NTF3, CHRM2, GABRA2, GRIA1, TENM2, NOVA1, RIMS4, CPNE6, GRM4, PPFIA2, C15orf59, ADCYAP1, TRABD2B, NOS1, CCL16, GREM1, PLP1, CCL23, GRIA3 |
| trans-synaptic signaling | 43 | 436 | 0.0011 | EFNB3, SYT10, HTR3B, UNC13C, GABRA4, SLC6A3, CRHBP, SCN2B, GRID2, GRIA4, SLC18A2, CHRNA9, DLGAP1, EGR3, CACNB2, RIT2, SYT9, GRM7, CACNA1G, SLC22A2, ANXA9, DLG2, PTCHD1, PTPRD, RPH3A, GRIK1, CHRNA2, NRXN1, NSG1, SEPT5, CHRM2, GABRA2, GRIA1, TENM2, NOVA1, RIMS4, CPNE6, GRM4, PPFIA2, C15orf59, NOS1, PLP1, GRIA3 |
| fatty acid derivative metabolic process | 23 | 162 | 0.0016 | PLA2G3, CYP4F2, DGAT2, PTGIS, ALOX12, CYP4F22, CYP2C18, HPGD, CYP2A7, ALOXE3, ALOX12B, CYP2F1, THEM5, ELOVL4, PTGDS, CYP1A1, ALOX15B, PLA2G4F, HMGCLL1, AWAT2, ACSS3, ACSM2A, ACSBG2 |
| response to lipid | 68 | 858 | 0.0019 | DCN, TGFBR3, ACR, SFRP1, CCND1, DGAT2, AICDA, GPLD1, ISL1, DUSP1, FOSB, ALOX12, EPO, DSG1, IL36A, IL36B, CCL21, SLC6A4, FGF10, CRHBP, STAR, IL18, GJA1, TIMP4, HOXB13, HPGD, FOS, FOSL1, MLC1, DYNAP, GHRHR, FIBIN, DEFB104B, PGR, TPH2, SLIT3, ADCY2, GJB6, ARG1, PLN, RXRG, BRINP3, APOA2, S100A7, PTGFR, TRIM63, P2RY4, KLF4, AR, F7, ANXA1, MAOB, NR0B1, CYP1A1, CATSPERD, GJB2, IL1F10, IL36RN, AREG, SOX10, FOXP2, ESR1, FAM107A, CRYAB, NR4A1, ADCYAP1, NOS1, NR4A3 |
| hormone metabolic process | 25 | 194 | 0.0026 | DGAT2, CMA1, PCSK2, ADH4, CYP11A1, CORIN, CRHBP, STAR, CYP2C18, ADH1B, SULT1B1, GCNT4, GHRHR, CYP3A4, BCO2, AKR1B10, DIO1, KLK6, CYP1A1, AFP, ESR1, AWAT2, DIO3, RDH12, CRABP2 |
| response to glucocorticoid | 21 | 147 | 0.0036 | CCND1, ISL1, DUSP1, FOSB, EPO, STAR, FOS, FOSL1, GHRHR, FIBIN, TPH2, SLIT3, ARG1, APOA2, TRIM63, ANXA1, MAOB, GJB2, AREG, FAM107A, ADCYAP1 |
| cell-cell adhesion via plasma-membrane adhesion molecules | 29 | 257 | 0.0045 | MYOT, DSG1, DSC1, CDH19, PCDH10, HMCN1, LRRC4C, DSC2, GRID2, CLDN17, LRFN5, CNTN2, CADM3, PCDH11X, PCDH15, PCDH9, PTPRD, FAT4, PCDHGA2, CLDN8, NRXN1, PCDH20, TENM1, SPARCL1, CDH18, PCDHGB6, TENM2, FAT3, CDHR1 |
| response to bacterium | 53 | 634 | 0.0053 | DCN, AICDA, PI3, STATH, EPO, CRP, IL36A, IL36B, FGF10, SPACA3, IL22RA1, HMCN1, STAR, GPM6A, IL18, GJA1, ADAMTS5, TIMP4, PGLYRP3, HPGD, FIGF, GSDMA, FOS, F2, KLK3, DEFB104B, NLRP10, LCE3A, GJB6, ARG1, PGLYRP4, VIP, S100A7, S100A12, GBP6, PTGFR, C10orf99, EPPIN, WFDC12, WFDC5, LY6G6C, MAOB, CYP1A1, KRT6A, GJB2, STAB2, IL1F10, IL36RN, DEFB115, DGKB, SCN7A, TRDN, NOS1 |
| chemical synaptic transmission | 39 | 418 | 0.0083 | SYT10, HTR3B, UNC13C, GABRA4, SLC6A3, CRHBP, SCN2B, GRID2, GRIA4, SLC18A2, CHRNA9, DLGAP1, EGR3, CACNB2, RIT2, SYT9, GRM7, CACNA1G, SLC22A2, ANXA9, DLG2, PTCHD1, RPH3A, GRIK1, CHRNA2, NRXN1, NSG1, SEPT5, CHRM2, GABRA2, GRIA1, NOVA1, RIMS4, CPNE6, GRM4, PPFIA2, C15orf59, PLP1, GRIA3 |

| | | | | |
|---|---|---|---|---|
| cell-cell adhesion | 44 | 505 | 0.011 | MYOT, DSG1, DSC1, CDH19, PCDH10, CSTA, HMCN1, CRNN, LRRC4C, DSC2, GRID2, CLDN17, COL14A1, LRFN5, CTNND2, CNTN2, CRISP2, CADM3, ANXA9, PCDH11X, PCDH15, DLG2, CDSN, ANXA1, PCDH9, DSP, PTPRD, FAT4, PCDHGA2, CLDN8, JAM2, CTNNA2, NRXN1, PCDH20, CTNNA3, CYR61, TENM1, XG, SPARCL1, CDH18, PCDHGB6, TENM2, FAT3, CDHR1 |
| muscle contraction | 27 | 248 | 0.0138 | MYOT, MYH2, TMOD1, SGCA, SCN2B, GJA1, ACTC1, KCNJ3, MYL1, CACNB2, EDN3, PTGER3, ANK2, CACNA1G, TNNI1, LMOD1, DES, TRIM63, ADRA1A, MYH11, SCN7A, CKMT2, CRYAB, TRDN, ACTN2, NOS1, FXYD1 |
| muscle structure development | 42 | 479 | 0.0138 | DCN, TGFBR3, FGF6, ISL1, IGFBP5, EGR1, TMOD1, USP2, SGCA, MEOX2, FGF10, BARX2, ACTC1, TCF23, IGF1, FOS, TBX5, WFIKKN2, PDLIM2, EGR3, NOG, ANK2, MYOZ1, TNNI1, LMOD1, TCF21, BAI3, NPHS1, CNTFR, DSP, ADRA1A, FHL1, SYNPO2L, MYH11, JAM2, FOXP2, MYOCD, KCNAB1, CRYAB, ACTN2, NOS1, GREM1 |
| muscle system process | 30 | 297 | 0.0177 | MYOT, MYH2, TMOD1, SGCA, SCN2B, GJA1, ACTC1, KCNJ3, IGF1, MYL1, CACNB2, EDN3, PTGER3, ANK2, PLN, CACNA1G, MYOZ1, TNNI1, LMOD1, DES, TRIM63, ADRA1A, MYH11, SCN7A, CKMT2, CRYAB, TRDN, ACTN2, NOS1, FXYD1 |
| glycosaminoglycan catabolic process | 12 | 61 | 0.0177 | DCN, HYAL4, LYVE1, OGN, FGF2, PGLYRP3, BCAN, PRELP, PGLYRP4, HPSE2, OMD, STAB2 |
| fatty acid biosynthetic process | 17 | 118 | 0.0177 | PTGIS, ALOX12, HPGD, ALOXE3, ALOX12B, ACSM2B, CYP3A4, ELOVL4, PTGDS, OLAH, CYP1A1, ALOX15B, PLA2G4F, ACSM2A, ACSBG2, PLP1, FADS6 |
| female pregnancy | 22 | 183 | 0.0177 | IGFBP5, PTGIS, FOSB, EPO, DSG1, CORIN, CRHBP, GJA1, TCF23, HPGD, FOS, FOSL1, PSG11, PGR, ARG1, AR, GJB2, CRHR1, ENDOU, ESR1, TPPP3, ADCYAP1 |
| regulation of membrane potential | 39 | 440 | 0.0177 | DCN, NALCN, ALOX12, HTR3B, GABRA4, KCNC1, SCN2B, DSC2, GRID2, GRIA4, GJA1, SCN3A, KCNJ3, CHRNA9, CACNB2, KCNK7, ANK2, PLN, BCO2, CACNA1G, WISP3, DSP, ADRA1A, FHL1, GRIK1, ABCB5, CHRNA2, NRXN1, SCN7A, CTNNA3, GABRA2, GRIA1, TRDN, RIMS4, ACTN2, C15orf59, ADCYAP1, FXYD1, GRIA3 |
| long-chain fatty acid biosynthetic process | 9 | 33 | 0.0177 | ALOX12, HPGD, ALOXE3, ALOX12B, CYP3A4, CYP1A1, ALOX15B, ACSBG2, PLP1 |
| response to steroid hormone | 32 | 328 | 0.0177 | ACR, CCND1, ISL1, DUSP1, FOSB, EPO, DSG1, STAR, FOS, FOSL1, GHRHR, FIBIN, PGR, TPH2, SLIT3, ARG1, RXRG, APOA2, TRIM63, AR, ANXA1, MAOB, NR0B1, CATSPERD, GJB2, AREG, SOX10, ESR1, FAM107A, NR4A1, ADCYAP1, NR4A3 |
| long-chain fatty acid metabolic process | 16 | 111 | 0.0255 | CYP4F2, PTGIS, ALOX12, SLC27A6, CYP2C18, HPGD, CYP2A7, ALOXE3, ALOX12B, CYP2F1, CYP3A4, PTGDS, CYP1A1, ALOX15B, ACSBG2, PLP1 |
| regulation of smooth muscle cell proliferation | 18 | 136 | 0.0255 | IGFBP5, EREG, ALOX12, CNN1, OGN, FGF2, IL18, GJA1, HPGD, IGF1, APOD, VIP, NOX1, KLF4, NDRG4, MYOCD, PDE1A, NR4A3 |
| regulation of water loss via skin | 8 | 28 | 0.0313 | ALOX12, KRT1, ACER1, ALOXE3, ALOX12B, FLG, HRNR, FLG2 |
| arachidonic acid metabolic process | 11 | 57 | 0.0338 | CYP4F2, PTGIS, ALOX12, CYP2C18, CYP2A7, ALOXE3, ALOX12B, CYP2F1, PTGDS, CYP1A1, ALOX15B |

| | | | | |
|---|---|---|---|---|
| response to organic cyclic compound | 65 | 911 | 0.0346 | OTC, ACR, SFRP1, CCND1, GPLD1, ISL1, IGFBP5, DUSP1, EGR1, FOSB, EPO, DSG1, HTR3B, SLC6A4, FGF10, SLC6A3, CRHBP, STAR, IL18, GJA1, HOXB13, HPGD, FOS, HMP19, FOSL1, MLC1, DYNAP, GHRHR, CIDEA, FIBIN, PGR, TPH2, SLIT3, PDE2A, ADCY2, ARG1, PLN, RXRG, MT-ND6, APOA2, PTGFR, TRIM63, P2RY4, KLF4, AR, F7, ANXA1, MAOB, NR0B1, CYP1A1, CATSPERD, GJB2, RYR3, AREG, SOX10, PLA2G4F, FOXP2, NSG1, CHRM2, ESR1, FAM107A, CRYAB, NR4A1, ADCYAP1, NR4A3 |
| monocarboxylic acid biosynthetic process | 21 | 183 | 0.0378 | DCN, PTGIS, ALOX12, STAR, HPGD, ACOX2, ALOXE3, ALOX12B, ACSM2B, BCAN, CYP3A4, ELOVL4, PTGDS, OLAH, CYP1A1, ALOX15B, PLA2G4F, ACSM2A, ACSBG2, PLP1, FADS6 |
| chemotaxis | 44 | 545 | 0.0381 | EFNB3, ISL1, MYOT, SEMA6A, CCL21, LHX4, FGF2, FGF10, FGF7, GDF7, FIGF, SEMA3E, FOSL1, EGR3, DEFB104B, SEMA6D, NOG, CNTN2, SLIT3, CXCL14, EDN3, CMTM5, GFRA1, LGR6, SPTA1, S100A12, EPHA7, C10orf99, ANXA1, EPHA6, CXCL12, NRXN1, KIF5C, NTF3, CYR61, BOC, TENM2, NR4A1, CCL16, CCL14, NCAM1, EPHA5, CCL23, NR4A3 |
| morphogenesis of a branching structure | 20 | 170 | 0.0388 | SFRP1, SALL1, FGF2, FGF10, FGF7, GDF7, RSPO2, HOXB13, SEMA3E, PGR, NOG, EYA1, TCF21, EPHA7, AR, FAT4, AREG, SOX10, ESR1, GREB1L |
| response to hormone | 61 | 849 | 0.0429 | OTC, TGFBR3, ACR, SFRP1, GNRHR, CCND1, GPLD1, ISL1, IGFBP5, DUSP1, EGR1, EREG, FOSB, MAS1, EPO, DSG1, CCL21, GLP2R, CYP11A1, ATP6V1C2, CRHBP, STAR, GJA1, TIMP4, LHCGR, FOS, FOSL1, GHRHR, FIBIN, PGR, TPH2, SLIT3, CRHR2, ADCY2, ARG1, PLN, RXRG, APOA2, PTGFR, TRIM63, P2RY4, AR, F7, ANXA1, MAOB, NR0B1, ADRA1A, CATSPERD, GJB2, AREG, CXCL12, SOX10, CRHR1, ESR1, FAM107A, NR4A1, ACTN2, CPEB2, ADCYAP1, NOS1, NR4A3 |
| olefinic compound metabolic process | 15 | 106 | 0.0429 | CYP4F2, PTGIS, ALOX12, CYP11A1, STAR, CYP2C18, CYP2A7, ALOXE3, ALOX12B, CYP2F1, BCO2, PTGDS, CYP1A1, ALOX15B, AFP |
| system process | 118 | 1942 | 0.0441 | TULP2, CYP4F2, CRX, SYT10, GNAT1, MYOT, KIAA1045, MYH2, SLURP1, STATH, CMA1, ALOX12, EPO, CRP, TMOD1, HTR3B, SLC6A4, SGCA, MEOX2, GABRA4, SRRM4, SLC6A3, HMCN1, CORIN, CRHBP, SCN2B, GRID2, GRIA4, GJA1, SCN3A, PPEF2, ACTC1, LHCGR, KCNJ3, LOXHD1, SOAT2, IGF1, FOS, MYL1, TMPRSS11E, FOSL1, ARR3, CHRNA9, KLK2, KLK3, ALOXE3, LCE1D, OR4N2, CACNB2, CNTN2, IGDCC3, TAC4, PDE2A, EDN3, EYA1, ADCY2, GJB6, PTGER3, ARC, ANK2, PLN, GRM7, CACNA1G, MYOZ1, TNNI1, NPY1R, SLC22A2, VIP, LMOD1, NKX6-2, ANXA9, BAI3, NOX1, PCDH15, DES, TRIM63, AR, DLG2, TRPM3, CA6, NPHS1, PTCHD1, ADRA1A, GJB2, KLK8, NDRG4, CXCL12, MYH11, CRHR1, GRIK1, POU6F2, CTNNA2, SVEP1, CHRNA2, NRXN1, FOXP2, SCN7A, LHFPL3, NTF3, CHRM2, OTOGL, CKMT2, FAM107A, KCNAB1, GABRA2, GRIA1, CSMD1, CRYAB, TRPM1, TRDN, ACTN2, RDH12, C15orf59, ADCYAP1, NOS1, FXYD1, GRIA3, NR4A3 |
| morphogenesis of a branching epithelium | 19 | 159 | 0.0441 | SFRP1, SALL1, FGF2, FGF10, FGF7, GDF7, RSPO2, HOXB13, SEMA3E, PGR, NOG, EYA1, TCF21, AR, FAT4, AREG, SOX10, ESR1, GREB1L |

| | | | | |
|---|---|---|---|---|
| neuron differentiation | 70 | 1019 | 0.0467 | NNAT, ACTL6B, SFRP1, WNT16, EFNB3, ISL1, GNAT1, MYOT, FLRT1, SALL1, DCLK1, SEMA6A, LHX4, SRRM4, GDF7, RSPO2, GPM6A, GRID2, GJA1, FIGF, NYAP1, SEMA3E, CTNND2, FAIM2, SEMA6D, NOG, CNTN2, OPCML, SLIT3, EDN3, STMN4, EYA1, APOD, GFRA1, BTG4, ARC, HAND2, SPINK5, LGR6, SPTA1, NKX6-2, EPHA7, BAI3, ZC4H2, PTPRD, EPHA6, KLK8, FAT4, AREG, CXCL12, CTNNA2, NRXN1, ATCAY, KIF5C, MYT1L, NTF3, MYO16, TENM1, GAS7, BOC, EVX1, DIO3, TENM2, TRPM1, ADCYAP1, NCAM1, EPHA5, PLP1, NR4A3, CDHR1 |
| nervous system development | 139 | 2371 | 0.0482 | NNAT, CTTNBP2, ACTL6B, PLA2G3, SFRP1, CRX, WNT16, EFNB3, GLI1, ISL1, GNAT1, C2orf40, MYOT, FLRT1, CMA1, SALL1, EPO, DCLK1, SEMA6A, SLC6A4, PCSK2, RPS6KA6, ASPA, LHX4, GABRA4, FGF2, FGF10, KCNC1, SRRM4, SLC6A3, GDF7, STAR, RSPO2, LRRC4C, SCN2B, GPM6A, GRID2, GJA1, TIMP4, CSMD3, FIGF, NELL1, NYAP1, SEMA3E, FOS, CTNND2, F2, RBFOX1, MAL, SNTG2, MOBP, EGR3, GHRHR, RIT2, FAIM2, SEMA6D, SPINK6, NOG, PCP4, CNTN2, OPCML, BCAN, SLIT3, EDN3, ZBTB16, STMN4, EYA1, APOD, GFRA1, BTG4, ARC, ANK2, RFX4, HAND2, SPINK5, NRG2, LGR6, BRINP3, DNM3, SPTA1, NKX6-2, EPHA7, BAI3, CHRDL1, NAP1L2, KLF4, ZC4H2, KLK6, ANXA1, MAOB, NR0B1, CNTFR, VIT, PTCHD1, TUBB2A, PTPRD, CPNE9, EPHA6, KLK8, NDRG4, FAT4, AREG, CXCL12, SOX10, NAV3, GRIK1, POU6F2, CTNNA2, NRXN1, FOXP2, ATCAY, KIF5C, MYT1L, NTF3, CHRM2, MYO16, TENM1, GAS7, BOC, LSAMP, EVX1, KCNAB1, GABRA2, DIO3, TENM2, ZIC4, TRPM1, CPNE6, PPFIA2, ADCYAP1, RBFOX3, PLK5, NOS1, NCAM1, EPHA5, PLP1, NR4A3, CRABP2, CDHR1 |
| regulation of antibacterial peptide production | 4 | 4 | 0.0487 | KLK5, SPINK5, PGC, KLK7 |
| response to alcohol | 24 | 233 | 0.0487 | TGFBR3, SFRP1, CCND1, GPLD1, FOSB, CCL21, SLC6A3, CRHBP, STAR, ACTC1, HPGD, FOS, FOSL1, MLC1, DYNAP, DEFB104B, SLIT3, ADCY2, PTGFR, P2RY4, KLF4, F7, MAOB, ADCYAP1 |
| icosanoid metabolic process | 15 | 109 | 0.0489 | PLA2G3, CYP4F2, PTGIS, ALOX12, CYP4F22, CYP2C18, HPGD, CYP2A7, ALOXE3, ALOX12B, CYP2F1, PTGDS, CYP1A1, ALOX15B, PLA2G4F |
| regulation of system process | 46 | 592 | 0.0492 | IGFBP5, C2orf40, CNN1, SGCA, FGF10, CORIN, CRHBP, SCN2B, DSC2, GJA1, KCNJ3, IGF1, TACR1, CACNB2, TAC4, EDN3, PTGER3, ANK2, PLN, CACNA1G, HAND2, TNNI1, VIP, APOA2, PI16, DES, TRIM63, KLF4, NPHS1, DSP, ADRA1A, KCNIP4, RYR3, KLK8, SOX10, CRHR1, JAM2, NRXN1, CTNNA3, CHRM2, MYOCD, TRDN, HSPB6, NOS1, FXYD1, NR4A3 |
| epidermis morphogenesis | 8 | 32 | 0.0497 | IGFBP5, FGF10, FGF7, SOSTDC1, KLF4, TGM3, FLG2, KLK14 |

**Supplementary Table S7: Differential correlation of the noncoding:coding gene pairs among the HPV16 positive CaCx patients as compared to HPV negative normal individuals**

.

| S. No | Noncoding genes log2(Fold change) | Corresponding Coding genes log2(Fold change) | Correlation between noncoding: coding gene pairs among HPV16 positive CaCx patients; Pearson's Correlation coefficient (r1) (FDR corrected p value) | Correlation between noncoding: coding gene pairs among HPV negative normal individuals; Pearson's Correlation coefficient (r2) (FDR corrected p value) | Fisher r to z transformation of the Pearson's correlation coefficients of the noncoding:coding gene pairs among the HPV16 positive CaCx patients (r1) as compared to HPV negative normal individuals (r2); z score (FDR corrected p value) |
|---|---|---|---|---|---|
| 1. | RP11-326C3.2 (3.277) | ATHL1 (2.275) | 0.9969413 **(3.73E-16)** | 0.8540564 **(2.18E-10)** | -8.28 **(1.01E-14)** |
| 2. | RP11-284F21.7 (3.031) | BCAN (-2.114) | 0.887049945 **(1.75E-15)** | 0.994559889 **(8.62E-32)** | 6.49 **(3.43E-09)** |
| 3. | TEX26-AS1 (-2.606) | MEDAG (-2.103) | 0.998736758 **(9.49E-55)** | 0.977160967 **(2.22E-22)** | -6.1 **(2.72E-08)** |
| 4. | FLG-AS1 (-2.795) | FLG2 (-7.825) | 0.912714333 **(1.25E-17)** | 0.992659053 **(7.34E-30)** | 5.29 **(2.46E-06)** |
| 5. | RP11-687M24.7 (-2.78) | PKNOX2 (-2.841) | 0.996890386 **(7.66E-47)** | 0.964218559 **(2.03E-19)** | -5.17 **(3.77E-06)** |
| 6. | AC053503.6 (-3.064) | DES (-2.348) | 0.9995801 **(2.59E-64)** | 0.995585906 **(3.83E-33)** | -4.95 **(9.96E-06)** |
| 7. | TMPRSS4-AS1 (-2.263) | TMPRSS4 (-2.263) | 0.976401388 **(6.05E-29)** | 0.794670658 **(2.64E-08)** | -4.75 **(2.08E-05)** |
| 8. | RP11-523H20.3 (2.245) | LBX2 (2.304) | 0.974402311 **(3.11E-28)** | 0.779437491 **(7.14E-08)** | -4.74 **(2.08E-05)** |
| 9. | MNX1-AS1 (3.055) | MNX1 (3.233) | 0.620896272 **(7.64E-06)** | 0.949331214 **(4.35E-17)** | 4.62 **(3.44E-05)** |
| 10. | CTA-384D8.31 | KLHDC7B | 0.9589518 | 0.7216236 | -4.29 |

| | | | | | |
|---|---|---|---|---|---|
| | (6.226) | (5.55) | **(3.65E-16)** | **(1.71E-06)** | **(1.39E-04)** |
| 11. | AC019349.5 (-2.791) | KRT13 (-3.983) | 0.997854146 **(4.80E-50)** | 0.983926805 **(1.50E-24)** | -4.24 **(1.5E-04)** |
| 12. | CTC-518B2.9 (-2.983) | KLK12 (-5.026) | 0.997530417 **(7.32E-49)** | 0.984474048 **(9.72E-25)** | -3.88 **(0.001)** |
| 13. | GRIK1-AS1 (-2) | GRIK1 (-4.476) | 0.836279929 **(2.12E-12)** | 0.97179263 **(5.56E-21)** | 3.84 **(0.001)** |
| 14. | RP11-1L12.3 (2.303) | BBOX1 (-3.602) | 0.995209083 **(4.95E-43)** | 0.974180223 **(1.45E-21)** | -3.56 (**0.002)** |
| 15. | ADAMTS19-AS1 (-2.455) | ADAMTS19 (-2.48) | 0.962903031 **(5.61E-25)** | 0.817100378 **(5.18E-09)** | -3.51 **(0.002)** |
| 16. | RP4-792G4.2 (4.173) | FOXD3 (2.287) | 0.961772226 **(8.94E-25)** | 0.817281162 **(5.19E-09)** | -3.45 **(0.003)** |
| 17. | RP11-94C24.11 (-2.484) | CACNA1G (-2.083) | 0.913608245 **(1.04E-17)** | 0.672532914 **(1.50E-05)** | -3.08 **(0.01)** |
| 18. | AC103563.8 (-3.868) | MAL (-6.813) | 0.953087569 **(5.29E-23)** | 0.815968801 **(5.57E-09)** | -3.02 **(0.011)** |
| 19. | RNF219-AS1 (-2.236) | POU4F1 (2.484) | 0.8405644 **(1.31E-12)** | 0.9583 **(6.52E-16)** | 2.95 **(0.013)** |
| 20. | RP11-483C6.1 (-2.871) | NOVA1 (-2.831) | 0.984787566 **(9.49E-33)** | 0.93981585 **(5.43E-16)** | -2.94 **(0.013)** |
| 21. | RP11-845M18.6 (2.535) | KRT86 (2.191) | 0.59416069 **(2.28E-05)** | 0.874886066 **(2.26E-11)** | 2.81 **(0.018)** |
| 22. | CTD-3214H19.6 (2.17) | PCP2 (2.068) | 0.972681202 **(1.14E-27)** | 0.901638209 **(6.68E-13)** | -2.77 **(0.02)** |
| 23. | RP11-64D24.2 (-3.131) | ZBTB16 (-3.274) | 0.462637338 **(1.67E-03)** | 0.801612368 **(1.63E-08)** | 2.53 **(0.039)** |
| 24. | RP11-74H8.1 (-2.403) | CACNG4 (2.428) | 0.915291529 **(7.16E-18)** | 0.745802783 **(5.07E-07)** | -2.5 **(0.041)** |
| 25. | EVX1-AS (-2.394) | EVX1 (-2.012) | 0.992813687 **(2.14E-39)** | 0.978045795 **(1.36E-22)** | -2.36 (0.057) |

| | | | | | |
|---|---|---|---|---|---|
| 26. | CDKN2B-AS1 (3.196) | CDKN2A (5.426) | 0.867824196 **(3.69E-14)** | 0.647170274 **(3.96E-05)** | -2.33 (0.061) |
| 27. | RP11-160H12.2 (-3.449) | DLG2 (-2.639) | 0.919943349 **(2.35E-18)** | 0.782333456 **(6.00E-08)** | -2.26 (0.07) |
| 28. | RP11-746B8.1 (-3.616) | CCDC178 (-3.162) | 0.980828136 **(1.02E-30)** | 0.947278251 **(7.45E-17)** | -2.16 (0.087) |
| 29. | HOXC-AS2 (2.821) | HOXC9 (3.641) | 0.97132462 **(2.98E-27)** | 0.927942209 **(7.05E-15)** | -1.98 (0.121) |
| 30. | RP11-44K6.2 (3.9) | IDO1 (5.532) | 0.897092724 **(3.03E-16)** | 0.755281002 **(3.02E-07)** | -1.98 (0.121) |
| 31. | ZFHX4-AS1 (-3.55) | ZFHX4 (-3.027) | 0.837861246 **(1.79E-12)** | 0.630512017 **(7.01E-05)** | -1.98 (0.121) |
| 32. | RP11-2O17.2 (-3.401) | C5orf64 (-2.242) | 0.990522419 **(5.73E-37)** | 0.996169513 **(5.31E-34)** | 1.91 (0.139) |
| 33 | RNF157-AS1 (2.172) | FOXJ1 (4.289) | 0.8749213 **(1.27E-14)** | 0.723049 **(1.62E-06)** | -1.85 (0.151) |
| 34. | RP5-1120P11.1 (2.312) | C6orf223 (4.37) | 0.962748189 **(5.87E-25)** | 0.912659393 **(1.22E-13)** | -1.84 (0.151) |
| 35. | FGF10-AS1 (-3.557) | FGF10 (-3.211) | 0.952303463 **(7.21E-23)** | 0.978433014 **(1.10E-22)** | 1.7 (0.203) |
| 36. | RP11-302F12.10 (-3.347) | SEL1L3 (2.262) | 0.800245582 **(8.93E-11)** | 0.904112312 **(4.75E-13)** | 1.66 (0.213) |
| 37. | RP11-485G7.5 (2.214) | RMI2 (2.37) | 0.852320312 **(3.12E-13)** | 0.929758202 **(4.90E-15)** | 1.65 (0.213) |
| 38. | AP000697.6 (2.309) | SIM2 (3.186) | 0.941278075 **(4.95E-21)** | 0.880707098 **(1.15E-11)** | -1.56 (0.249) |
| 39. | RP11-1081L13.4 (-2.241) | PTPN5 (-2.841) | 0.954278375 **(3.22E-23)** | 0.90795516 **(2.60E-13)** | -1.52 (0.26) |
| 40. | CACNA1G-AS1 (-2.242) | CACNA1G (-2.083) | 0.955596018 **(1.82E-23)** | 0.914168171 **(9.89E-14)** | -1.43 (0.298) |
| 41. | ST6GAL2-IT1 (-2.86) | ST6GAL2 (-2.022) | 0.774550291 | 0.878765378 | 1.42 (0.298) |

| | | | | | |
|---|---|---|---|---|---|
| | | | **(8.52E-10)** | **(1.43E-11)** | |
| 42. | RP11-1081L13.4 (-2.241) | IGSF22 (-2.043) | 0.902887947 **(1.01E-16)** | 0.819296024 **(4.49E-09)** | -1.4 (0.304) |
| 43. | CTB-92J24.3 (-2.435) | ZNF726 (-2.303) | 0.988705232 **(2.05E-35)** | 0.978759496 **(9.37E-23)** | -1.34 (0.333) |
| 44. | CTC-518B2.12 (-3.191) | KLK10 (-2.906) | 0.981361959 **(6.07E-31)** | 0.966014855 **(9.47E-20)** | -1.28 (0.361) |
| 45. | RP11-464F9.21 (-2.47) | SYNPO2L (-4.381) | 0.830163545 **(4.17E-12)** | 0.902904791 **(5.61E-13)** | 1.26 (0.367) |
| 46. | RP11-475O23.2 (-2.918) | FAM107A (-2.185) | 0.923152361 **(1.08E-18)** | 0.86742209 **(5.29E-11)** | -1.21 (0.39) |
| 47. | RP3-525N10.2 (-2.977) | BAI3 (-3.035) | 0.851467077 **(3.43E-13)** | 0.913020935 **(1.18E-13)** | 1.19 (0.391) |
| 48. | RP11-532N4.2 (-3.124) | TRDN (-2.764) | 0.996181766 **(4.86E-45)** | 0.997815693 **(1.01E-37)** | 1.17 (0.395) |
| 49. | AC098617.1 (-2.431) | SDPR (-2.408) | 0.438953338 **(2.98E-03)** | 0.631587815 **(6.84E-05)** | 1.15 (0.4) |
| 50. | HOXC-AS2 (2.821) | HOXC6 (3.202) | 0.962557663 **(6.27E-25)** | 0.936484941 **(1.17E-15)** | -1.14 (0.4) |
| 51. | RFPL1S (2.341) | RFPL1 (-3.096) | 0.932847029 **(7.41E-20)** | 0.887649918 **(4.83E-12)** | -1.13 (0.4) |
| 52. | CTD-2530H12.2 (-2.051) | DGAT2 (-2.407) | 0.768719486 **(1.35E-09)** | 0.848579193 **(3.63E-10)** | 0.98 (0.495) |
| 53. | RP3-333A15.2 (-3.45) | PTGER3 (-3.504) | 0.660972439 **(1.22E-06)** | 0.771550744 **(1.16E-07)** | 0.96 (0.499) |
| 54. | RP11-326E22.1 (-2.57) | EYA1 (-3.36) | 0.896432153 **(3.37E-16)** | 0.846151518 **(4.51E-10)** | -0.89 (0.538) |
| 55. | RP1-118J21.25 (2.702) | BMP8B (2.348) | 0.597016075 **(2.06E-05)** | 0.716700211 **(2.15E-06)** | 0.89 (0.538) |
| 56. | JAZF1-AS1 (-2.874) | JAZF1 (-2.114) | 0.755288095 **(3.77E-09)** | 0.82782832 **(2.25E-09)** | 0.82 (0.579) |

| | | | | | |
|---|---|---|---|---|---|
| 57. | HOXC-AS1 (3.02) | HOXC9 (3.641) | 0.906239938 **(5.13E-17)** | 0.935080586 **(1.59E-15)** | 0.8 (0.584) |
| 58. | <u>CTB-92J24.2</u> (-2.248) | ZNF726 (-2.303) | 0.901703822 **(1.24E-16)** | 0.931450919 **(3.57E-15)** | 0.79 (0.585) |
| 59. | RP11-284F21.10 (2.816) | BCAN (-2.114) | 0.926995608 **(3.93E-19)** | 0.948474248 **(5.42E-17)** | 0.76 (0.603) |
| 60. | <u>CTD-3080F16.3</u> (-2.87) | SFRP1 (-2.228) | 0.446683003 **(2.49E-03)** | 0.564664919 **(5.22E-04)** | 0.67 (0.663) |
| 61 | POU6F2-AS1 (-2.017) | POU6F2 (-2.273) | 0.853496373 **(2.72E-13)** | 0.889043326 **(4.09E-12)** | 0.62 (0.69) |
| 62. | HOXD-AS1 (2.692) | HOXD1 (2.968) | 0.84521667 **(7.52E-13)** | 0.880954096 **(1.14E-11)** | 0.59 (0.706) |
| 63. | CTA-134P22.2 (-3.064) | CADM3 (-3.644) | 0.881761465 **(4.25E-15)** | 0.85220762 **(2.58E-10)** | -0.5 (0.772) |
| 64. | AC008067.2 (-2.926) | CTNNA2 (-2.715) | 0.991392032 **(8.43E-38)** | 0.99282923 **(5.89E-30)** | 0.39 (0.842) |
| 65. | CTD-2314G24.2 (-2.354) | ISL1 (-2.28) | 0.895116562 **(4.08E-16)** | 0.911916929 **(1.36E-13)** | 0.39 (0.842) |
| 66. | HOXC-AS1 (3.02) | HOXC6 (3.202) | 0.902833159 **(1.00E-16)** | 0.917935217 **(5.10E-14)** | 0.37 (0.842) |
| 67. | AC118754.4 (-2.411) | SPNS2 (-2.39) | 0.802765065 **(7.14E-11)** | 0.83170734 **(1.64E-09)** | 0.37 (0.842) |
| 68 | CTC-518B2.9 (-2.983) | KLK11 (-3.288) | 0.655434378 **(1.59E-06)** | 0.61084622 **(1.33E-04)** | -0.31 (0.876) |
| 69 | HAND2-AS1 (-2.439) | HAND2 (-2.323) | 0.961994322 **(8.22E-25)** | 0.966937605 **(6.46E-20)** | 0.3 (0.877) |
| 70. | RP11-489D6.2 (-2.617) | RYR3 (-2.153) | 0.92305551 **(1.08E-18)** | 0.930018794 **(4.77E-15)** | 0.21 (0.937) |
| 71. | CTC-575N7.1 (-3.331) | ADAMTS19 (-2.48) | 0.979367974 **(4.14E-30)** | 0.981112464 **(1.75E-23)** | 0.19 (0.937) |
| 72. | AC007563.5 (-2.121) | IGFBP5 (-2.409) | 0.979366245 | 0.977497869 | -0.18 (0.937) |

| | | | | | |
|---|---|---|---|---|---|
| | | | **(3.92E-30)** | **(1.87E-22)** | |
| 73. | RP11-49I11.1<br>(2.097) | MOCOS<br>(2.601) | 0.831277548<br>**(3.73E-12)** | 0.842614538<br>**(6.20E-10)** | 0.16<br>(0.944) |
| 74. | TBX5-AS1<br>(-2.758) | TBX5<br>(-2.491) | 0.959430409<br>**(2.94E-24)** | 0.956616259<br>**(4.00E-18)** | -0.14<br>(0.944) |
| 75. | CTB-174D11.1<br>(-3.308) | SLIT3<br>(-2.777) | 0.980493037<br>**(1.37E-30)** | 0.97928941<br>**(6.85E-23)** | -0.13<br>(0.944) |
| 76. | RP3-323P13.2<br>(-2.554) | TCF21<br>(-2.187) | 0.3535263<br>**(1.90E-02)** | 0.3774077<br>**(2.81E-02)** | 0.12<br>(0.944) |
| 77. | RP11-573D15.3<br>(-2.374) | RFC4<br>(2.499) | 0.712436865<br>**(6.90E-08)** | 0.723662653<br>**(1.59E-06)** | 0.1<br>(0.946) |
| 78. | RP11-253M7.6<br>(-3.504) | PAQR5<br>(-2.769) | 0.62673611<br>**(5.97E-06)** | 0.617603044<br>**(1.07E-04)** | -0.06<br>(0.957) |
| 79. | CTC-321K16.1<br>(-2.798) | CXCL14<br>(-2.049) | 0.998921304<br>**(5.17E-56)** | 0.998948491<br>**(1.69E-42)** | 0.05<br>(0.957) |
| 80. | NOVA1-AS1<br>(-2.833) | NOVA1<br>(-2.831) | 0.195333638<br>(2.06E-01) | 0.571745978<br>**(4.33E-04)** | -- |
| 81. | RP1-40E16.11<br>(-3.065) | TUBB2A<br>(-2.131) | 0.009467543<br>(9.40E-01) | 0.4970141<br>**(2.86E-03)** | -- |
| 82. | RP11-96C23.10<br>(-4.002) | FAM25A<br>(-4.007) | 0.018765058<br>(9.04E-01) | 0.090396983<br>(6.11E-01) | -- |
| 83. | RP11-256L6.3<br>(-2.338) | HAL<br>(-4.788) | 0.651107753<br>**(1.94E-06)** | 0.085489689<br>(6.23E-01) | **--** |

| S. No | Noncoding genes log2(Fold change) | Corresponding Noncoding gene log2(Fold change) | Correlation between noncoding: noncoding gene pair among HPV16 positive CaCx patients; Pearson's Correlation coefficient (r1) (p value) | Correlation between noncoding: noncoding gene pair among HPV negative normal individuals; Pearson's Correlation coefficient (r2) (p value) | Fisher r to z transformation of the Pearson's correlation coefficient of the noncoding:noncoding gene pair among the HPV16 positive CaCx patients (r1) as compared to HPV negative normal individuals (r2); z score (p value) |
|---|---|---|---|---|---|
| 1. | RP11-149I2.4 (3.268) | CDKN2B-AS1 (3.196) | 0.751523348 **(4.141 E-09)** | 0.587204540 **(0.002)** | -1.27 (0.204) |

**NB:**

- **The noncoding genes that are underlined are sense intronic genes whereas the rest of the noncoding genes are antisense (ncNATs)**
- **The p values marked in bold are significant (p<0.05) and represent the FDR corrected p values determined after BH correction using R from the crude p values**

**Supplementary Table S8: Details of the gene clusters, that excluded DEcGs of significantly correlated ncNAT/sense intronic: coding gene pairs, identified through PPI network analysis of all DEcGs employing the MCODE plug-in of Cytoscape, along with their corresponding functions determined through DAVID**

| Cluster Number | MCODE score | # nodes | Common functions of the cluster (using DAVID) | Names of the genes forming the cluster |
|---|---|---|---|---|
| Cluster 1 | 32 | 32 | Involved in keratinisation, cornification, epidermis development, peptide crosslinking | LCE1F, SPRR1B, RPTN, LCE3E, LCE3A, IVL, LCE3D, PI3, SPRR2E, LCE2A, LCE1C, SPRR2G, SPRR2B, LCE6A, LCE1B, LCE2C, SPRR2A, LCE1A, SPRR3, SPRR2D, LCE1D, SPRR2F, TGM1, CDSN, LCE1E, PPL, LCE5A, SPRR1A, LCE2D, LCE2B, CSTA, LOR |
| Cluster 4 | 16.882 | 35 | Involved in cell cycle related processes like cell division, mitotic splindle organisation and attachment to kinetochore, DNA replication, chromosome segreagation, G2-M transition, nucleosome assembly, DNA damage response, cytokinesis | CENPA, CDK1, AURKA, CDC20, KIF2C, NUSAP1, UBE2C, ASF1B, KIF4A, OIP5, DTL, GINS2, CDC6, MCM2, CENPM, NEK2, NUF2, HIST1H2BJ, RAD51AP1, KIF14, RFC4, CDC25C, HIST1H3F, HIST1H3G, HIST1H4I, HIST2H4A, HIST2H4B, CDT1, HIST1H2AB, HIST2H2AA, HIST2H2AA3, HIST2H3A, HIST2H3C, HIST3H2A, FAM64A |
| Cluster 5 | 14.286 | 43 | Involved in cellular protein metabolic processes and post translational cellular protein modifications, GPCR pathway, signal transduction | AFP, APOA2, GPIHBP1, SPP1, ALPPL2, AMTN, APOL1, ART4, CHRDL1, CP, CYR61, FCGR3B, GOLM1, GP2, GPLD1, IGFBP5, LSAMP, LY6G6C, LY6K, LYPD1, LYPD3, OPCML, PCSK9, PRSS21, SERPINA1, SPACA4, SPARCL1, SPP2, STC2, TEX101, TMEM132A, XPNPEP2, F2, NTS, ADRA1A, CCK, EDN3, GNRHR, GRP, NMB, PROK1, PTGFR, TACR1 |

| | | | | |
|---|---|---|---|---|
| Cluster 6 | 9.111 | 10 | Involved in Type I Interferon signaling pathway, defence response to virus, regulation of viral genome replication, innate immune response, positive regulation of IFN-beta production | IFI6, IFIT3, IRF7, ISG15, OAS2, RSAD2, BST2, EGR1, IFI44L, USP18 |
| Cluster 7 | 8 | 8 | Involved in negative regulation of inflammatory response to antigenic stimulus, involved in various pathways like adenylate cyclase activity and GPCR signaling pathway, hormone mediated signaling pathway, positive regulation of cellular proliferation | ADCYAP1, ADRB1, CRHR1, CRHR2, GHRHR, GLP2R, LHCGR, VIP |
| Cluster 10 | 5.333 | 10 | Involved in fibroblast growth factor receptor signaling pathway, regulation of cell motility, migration and extracellular matrix organisation, positive regulation of protein phosphorylation, MAPK cascade | FGF2, FGF10, FGF22, FGF6, FGF7, FLRT1, MMP1, MMP13, MMP3, MMP7 |
| Cluster 11 | 5.2 | 6 | Involved in Interferon gamma signaling pathway | FCGR1A, GBP5, GBP6, NCAM1, TRIM17, TRIM31 |
| Cluster 13 | 5.111 | 10 | Involved in arachidonic acid metabolism, lipoxygenase activity, lipid oxidation, lipid metabolic processes | ALOX12B, CYP4F22, NIPAL4, PNPLA1, ALOX12, ALOX15, ALOX15B, ALOXE3, CYP4F2, LIPN |
| Cluster 14 | 5 | 5 | no sig pathways | DNM3, SNAP91, SYT8, SYT9, TFRC |
| Cluster 15 | 5 | 5 | Involved in keratan sulphate biosynthetic processes | B3GNT7, CHST6, OGN, OMD, PRELP |
| Cluster 16 | 5 | 5 | Involved in epinephrine receptor signaling pathway, positive regulation of kinase activity, peptidyl tyrosine phosphorylation, cell axon guidance | EFNB3, EPHA5, EPHA6, EPHA7, EPHB2 |

**Supplementary Table S9: The details of the hub coding genes of the clusters incorporating the coding genes of the ncNATs:coding gene pairs portraying significant altered correlated co-expression in CaCx patients and their association with overall patient survival.**

| Cluster No | Name of Hub genes | Degree of connectivity | Survival associated Hub genes |
|---|---|---|---|
| 2 | ANXA1, APLN, CCL16, CCL20, CCL21, CHRM2, CXCL1, CXCL10, CXCL11, CXCL12, CXCL3, CXCL5, CXCL8, CXCL9, GNG4, GNGT1, GRM4, GRM7, NPW, NPY1R, NPY4R, P2RY4, PF4, PPY, PTGER3, SAA1, ADCY2, PCP2 | 27 | PPY, CXCL1, GRM4,GRM7, GNG4, NPW |
| 3 | KRT18, KRT1, KRT8, KRT10 | 25 | KRT1 |
| 8 | MMP9, COL14A1 | 8 | None |
| 9 | ACTN2 | 9 | ACTN2 |
| 12 | GRIN2D, GRIN2C | 7 | GRIN2D |
